# Transcriptomic analysis of the BDNF-induced JAK/STAT pathway in neurons: a window into epilepsy-associated gene expression

**DOI:** 10.1101/577627

**Authors:** Kathryn M Hixson, Meaghan Cogswell, Amy R Brooks-Kayal, Shelley J Russek

## Abstract

**Background:** Brain-derived neurotrophic factor (BDNF) is a major signaling molecule that the brain uses to control a vast network of intracellular cascades fundamental to properties of learning and memory, and cognition. While much is known about BDNF signaling in the healthy nervous system where it controls the mitogen activated protein kinase (MAPK) and cyclic-AMP pathways, less is known about its role in multiple brain disorders where it contributes to the dysregulated neuroplasticity seen in epilepsy and traumatic brain injury (TBI). We previously found that neurons respond to prolonged BDNF exposure (both *in vivo* (in models of epilepsy and TBI) and *in vitro* (in BDNF treated primary neuronal cultures)) by activating the Janus Kinase/Signal Transducer and Activator of Transcription (JAK/STAT) signaling pathway. This pathway is best known for its association with inflammatory cytokines in non-neuronal cells.

**Results:** Here, using deep RNA-sequencing of neurons exposed to BDNF in the presence and absence of well characterized JAK/STAT inhibitors, and without non-neuronal cells, we determine the BDNF transcriptome that is specifically reliant on JAK/STAT signaling. Surprisingly, the transcriptome contains ion channels and neurotransmitter receptors coming from all the major classes expressed in the brain, along with key modulators of synaptic plasticity, neurogenesis, and axonal remodeling. Analysis of this dataset has also provided a window on the unique mechanism of JAK/STATs in neurons as differential gene expression mediated by STAT3 does not appear to be dependent upon phosphorylation at residue 705.

**Conclusions:** Our findings strengthen and expand the role that BDNF plays in the regulation of brain excitability at the transcriptional level. They also suggest that a majority of such signaling in neurons is tied to the activation of the JAK/STAT pathway which may be non-canonical, not based on phosphorylation of STAT3 proteins at Tyrosine 705.

## Background

In the injured brain, it has been suggested that neurogenesis (1), mossy fiber sprouting (2), and hippocampal cell death (3) contribute to aberrant neuronal circuit reorganization. One manifestation of this reorganization is thought to be altered long-term potentiation (LTP)-induced synaptic plasticity (4), which is associated with altered levels of particular neurotransmitter receptors and ion channels (5,6). As the functional properties of neurons change, a state of unchecked overexcitation may occur and is accompanied by decreased GABA-mediated inhibition (7). Many believe that a major mechanism for this altered brain plasticity is the increased synthesis and release of Brain-derived Neurotrophic Factor (BDNF), a major brain signaling molecule that plays an important role throughout life. BDNF is a neurotrophin ubiquitously expressed in the whole brain with a major role in activity-dependent alterations in neuronal morphology and synaptogenesis. Protein and mRNA levels of BDNF are markedly increased after seizures in brain regions involved in epileptogenesis (8,9) and mutations in *BDNF* can reduce the development of spontaneous seizures in animal models (10). Moreover, a conditional deletion of the high-affinity BDNF receptor TrkB in a subset of neurons, is sufficient to completely eliminate all behavioral evidence of the progression of epilepsy in the kindling model (11) and it has been shown that enhanced TrkB signaling can exacerbate epileptogenesis (12).

In addition to the TrkB-mediated BDNF response, the p75 Neurotrophin Receptor (p75NTR, also referred to as NGFR) is a low-affinity receptor for the cleaved mature form of BDNF while a high-affinity receptor for the uncleaved form, proBDNF. p75NTR levels are elevated after SE in animals and increased activation by proBDNF, like mature BDNF, increases susceptibility to seizures (13,14). Interestingly, our labs have shown that increased synthesis of proBDNF, and not mature BDNF, is the first response of the injured brain in the pilocarpine (PILO) model of epilepsy prior to SE-induced increases of p75NTR (15). This finding suggests that high levels of proBDNF may exert their effects through TrkB rather than p75NTR.

In the normal brain, BDNF activation of its receptors regulates multiple signaling pathways: mitogen-activated protein kinase/extracellular signal-regulated protein kinase (MAPK/ERK), phospholipase Cγ (PLCγ), phosphoinositide 3-kinase (PI3K), c-Jun N-terminal kinase (JNK), and NFkB (16). It is well established that BDNF modulates long-term potentiation (LTP) at Schaffer collateral-CA1 and mossy fiber-CA3 hippocampal synapses by acting through TrkB (17–21). Although p75NTR is not implicated in LTP, it plays a significant role in learning and memory through its modulation of hippocampal long-term depression (LTD) by altering AMPA receptor expression (22).

Studies from our laboratories demonstrate that in addition to the signaling pathways described above, increased levels of BDNF activate the Janus kinase/signal transducer and activator of transcription (JAK/STAT) pathway both *in vivo* in the rat PILO model of epilepsy and in vitro in BDNF-treated primary cultured neurons (23). The JAK/STAT pathway is a signaling cascade that has a prominent role in immune function and cancer development. The canonical pathway involves the binding of JAK to its target (such as a cytokine or hormone receptor), subsequent JAK phosphorylation that stimulates the recruitment and activation of STAT proteins (phospho-STAT (p-STAT)) and their movement as dimers into the nucleus where they function as transcriptional activators. Although JAK/STATs regulate transcription through this canonical pathway, more recently, non-canonical JAK/STAT signaling has been discovered where JAKs can act independently of STATs to regulate transcription in the nucleus, and STATs can function independently of their phosphorylation state to alter genome stability (24). While the role, and mechanism, of JAK/STAT signaling in the brain is still not completely understood, it is clear that it is an important part of how the brain regulates its synaptic connections (25).

Research from our laboratory has demonstrated that activation of the JAK/STAT pathway results in the expression of the inducible cAMP early repressor (ICER) via STAT3-mediated gene regulation. Moreover, we show that ICER represses the gene (*Gabra1/GABRA1*) coding for the α1 subunit of the type A γ-aminobutyric acid (GABA) receptor (GABAR), the major inhibitory neurotransmitter receptor in the brain, and that such repression reduces the number of α1 containing GABARs in neurons (26). Most importantly, reduced α1 subunit gene expression also occurs in epilepsy patients (5) as well as in other disorders of the nervous system (27–29). In addition, use of the JAK/STAT inhibitor, WP1066, in the PILO rat model of epilepsy reduces the number of spontaneous seizures after the latent period (30). We now hypothesize that in addition to controlling the gene regulation of *Gabra1/GABRA1*, the BDNF-induced JAK/STAT pathway controls the expression of diverse gene products that are involved in synaptic plasticity as well as in the epileptogenic process that follows brain injury. We also hypothesize that attenuating this pathway may provide a gateway to new treatments for refractory epilepsy and other brain disorders that display dysregulated neuroplasticity.

To test these hypotheses, and to gain a comprehensive understanding of the genome impacted by BDNF-induced JAK/STAT activation, we exposed primary cultured neurons to BDNF with and without JAK/STAT inhibitors and performed deep RNA-sequencing (RNA-seq) in order to determine the gene set within the BDNF response that could be ascribed to JAK/STAT activation (as defined by WP1066 and Ruxolitinib (Ruxo), a potent inhibitor of JAK1/2 activation (31)). We now report evidence that suggests BDNF acts through the JAK/STAT pathway to execute changes in: the expression of ion channels, neurotransmitter and GPCR receptors, and the expression of key modulators of synaptic plasticity, neurogenesis, and axonal remodeling. Many of these functions were previously ascribed to BDNF, however, the role of the JAK/STAT pathway in their manifestation has never been previously explored. Our studies also reveal the presence of many new BDNF target genes providing a broader landscape for BDNF mediated gene regulation of the healthy and impaired nervous system.

## Results

### BDNF signaling and the identification of epilepsy-associated gene networks

Primary cultured neurons were treated with BDNF (0.7nM, a physiologically relevant level of normal brain (15)) or vehicle control (water) to determine the genome response to prolonged receptor activation (4 hours) (Fig. 1). Using RNA-sequencing for open discovery, we identified a total of 2869 differentially expressed genes (DEGs) whose levels change (increase or decrease) in response to BDNF signaling (Fig. 2A) (CTRL vs V+B gene list, additional excel sheet file). Inspection of the gene set using Ingenuity Pathway Analysis (IPA) (32) revealed that only 83 of these genes have been previously associated with BDNF (as determined using the Ingenuity Knowledge Base, a curated database of all historical scientific information in IPA databases), suggesting that the majority of genes identified in this study are novel in their association with the BDNF signaling pathway. A subset of 194 potential BDNF regulated genes are associated with “seizure disorders” (BDNF and epilepsy, additional excel sheet file), an umbrella category in IPA that includes all epilepsy classes as represented in the curated knowledge base. Moreover, seizure disorder is the primary most significant neurological disorder identified by enrichment analysis of the gene pool (p= 3.29E-27).

**Figure 1.**
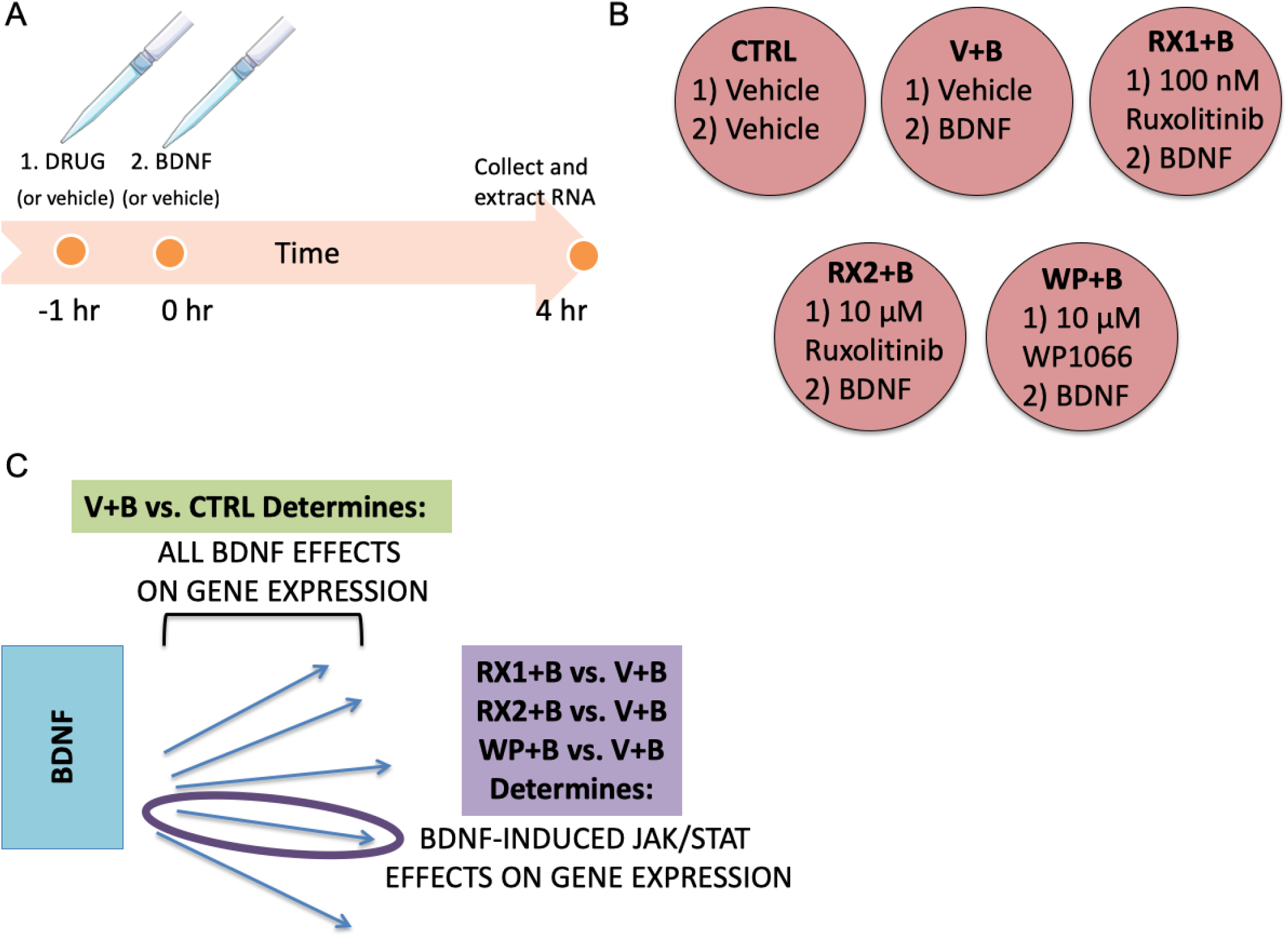
Schematic of experimental design and analysis. (A) Timeline representation of the drug treatment protocol, including the 1 hr drug or DMSO (vehicle, V) pretreatment (initiated at −1 hr), administration of BDNF (B) or water (vehicle) treatment (Time=0 hr) and time of cell collection and RNA extraction (4 hr). (B) 6-well plate showing each one of the 5 treatment groups and the abbreviations by which they will be referred to in the text: DMSO+Water (CTRL), DMSO+BDNF (V+B), 100nM Ruxolitinib+BDNF (RX1+B), 10µM Ruxolitinib+BDNF (RX2+B), 10µM WP1066+BDNF (WP+B). (C) Diagram represents the approach taken to identify differential gene expression (DEG) in response to JAK/STAT pathway inhibition. BDNF through its receptors activates multiple signaling pathways (represented by blue arrows) that impact transcription. Differential expression analysis comparing V+B vs. CTRL treatment groups (green box) will reveal the total set of genes that are regulated by BDNF-induced intracellular signaling pathways. To identify BDNF DEGs that are specific to JAK/STAT signaling, comparisons are made between the groups pretreated with JAK inhibitors (RX1+B, RX2+B, WP+B) and the group pretreated with vehicle (V+B) (purple box); DEGs from this comparison (where JAK/STAT inhibition reverses BDNF stimulation or inhibition) are thought to be associated with the JAK/STAT pathway.

**Figure 2.**
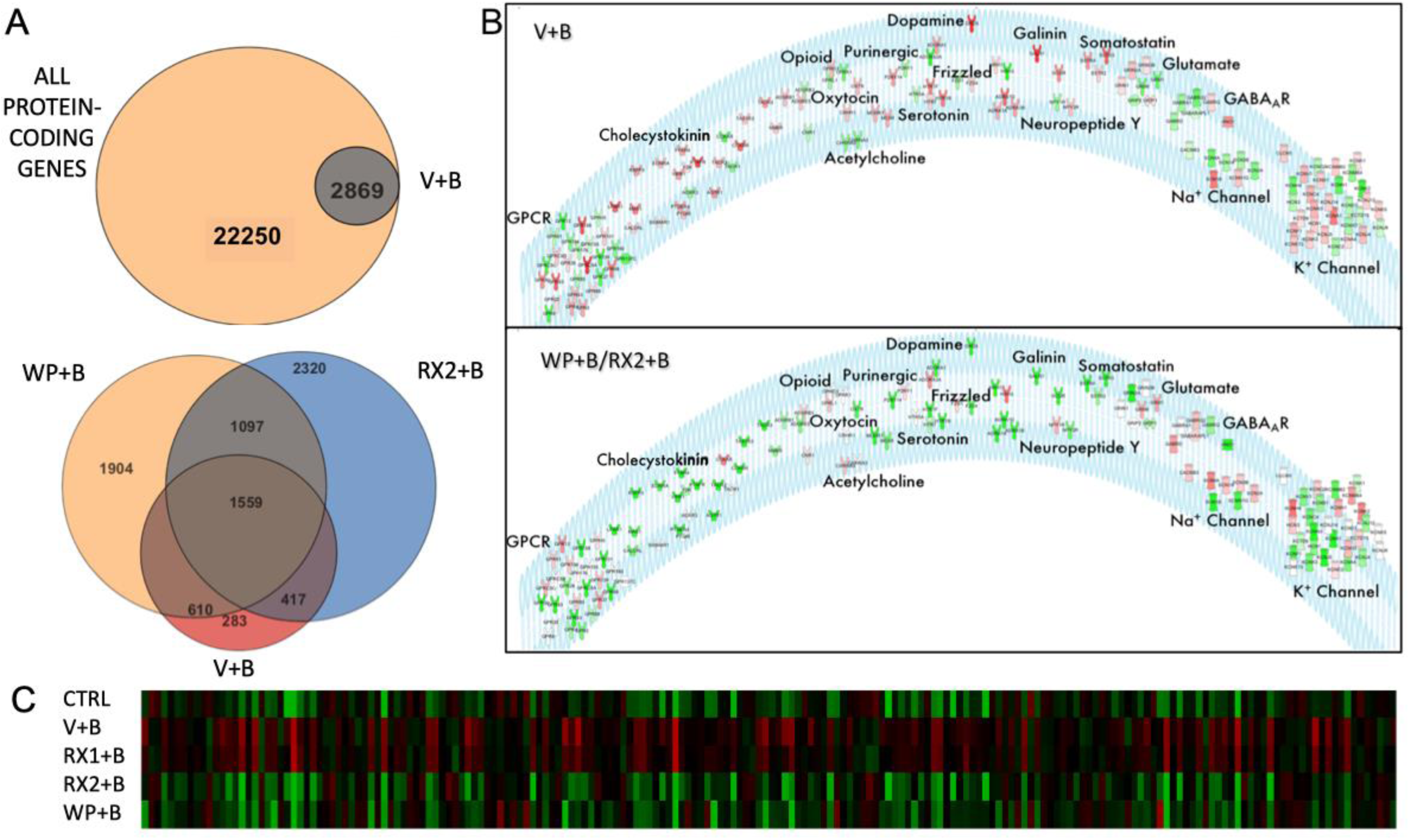
JAK/STAT inhibitors reverse BDNF-induced gene expression. (A) TOP: Venn diagram representation of all protein-coding genes in the rat genome (in accordance with Strand NGS notation) with all differentially expressed genes (DEGs) between primary neurons treated with DMSO+BDNF vs. DMSO+Water. BOTTOM: Venn diagram representation of all DEGs in DMSO+Water vs. DMSO+BDNF (BDNF, Red, 2869), WP1066+BDNF vs. DMSO+BDNF (WP1066,Orange, 5170 total) and 10µM Ruxo+BDNF vs. DMSO+BDNF (RX2, Blue, 5393 total genes). (B) Receptor and Ion Channel expression is altered by BDNF and rescued by JAK inhibition. TOP: List of Ion Channels and Receptor Subunits whose expression is altered by 4-hr BDNF treatment. Color represents direction and degree of fold change (Red: up, Green: down) sorted by receptor or channel type. BOTTOM: Receptor and Ion Channel reversal in expression by addition of WP or RX2 (white receptors are not affected by JAK/STAT inhibitors). Response to BDNF in presence of WP was used as the basis for coloring receptors depicted in the diagram. Red: Upregulated, Green: Downregulated. (C) Heatmap of all DEGs (columns) associated with Epilepsy (IPA) that are affected by exposure to BDNF. DW: DMSO+Water, DB: DMSO+BDNF, RX1: 100nM Ruxo+BDNF, RX2:10µM Ruxo+BDNF, WP: 10µM WP1066+BDNF. Green: low expression, red: high expression.

In addition to the 194 epilepsy-associated genes, pathway analysis suggests that increased levels of BDNF alters the expression of essential neurobiological gene sets that are required for brain development and that are dysregulated during epileptogenesis in animal models and human patients. For instance, the differentially expressed set of transcripts from BDNF-treated primary neurons is highly enriched for genes essential to axonal guidance (p=1.09E-11); further, there is a striking enrichment that is specific to all major classes of ion channels and neurotransmitter/neuropeptide receptors, with 133 of 599 total genes related to one of these categories regulated by prolonged BDNF exposure (4 hr) (p=2.66E-11) (Fig. 2B). Specifically, the differentially expressed transcripts for receptors include those activated by GABA, glutamate, acetylcholine, dopamine, opioids, serotonin, galanin, and neuropeptide Y, covering both metabotropic and ionotropic receptor subtypes. Most classes of ion channel genes are also represented, including those for potassium, sodium, and calcium, where mutations have been shown to underlie specific genetic epilepsies (33–35). There is also a specific decrease (1.73 fold) in the expression of *Gabra1* that is a member of the GABAR gene cluster which we have previously demonstrated to be regulated by BDNF using qRT-PCR in extracts of BDNF-treated primary neurons and hippocampal tissue extracted from the *in vivo* PILO model of temporal lobe epilepsy (23). As expected, the neuronal RNA-seq dataset revealed that prolonged exposure to BDNF increases BDNF transcript levels, a finding that is well established in the literature (36). Given our previous identification of a novel JAK/STAT pathway in neurons that is regulated by BDNF, we asked whether any members of the JAK/STAT pathway might be regulated at a transcriptional level in response to BDNF receptor activation. Our dataset indicates significant enrichment for transcripts linked to JAK/STAT signaling (p=6.03E-5) and in particular, an enrichment of *Jak1* (1.6-fold enrichment) and *Jak2* (1.8-fold enrichment).

### Neuropharmacological dissection of BDNF-induced JAK/STAT signaling reveals a non-canonical pathway in neurons

Primary cultured neurons were pre-treated with agents known to inhibit the JAK/STAT pathway (Fig. 1) by interfering with JAK kinase activity, either directly by inhibiting JAK phosphorylation or stimulating JAK degradation. Two different antagonists were employed in these studies: the small molecule WP1066 that has been shown to inhibit JAK2/STAT3 signaling and degrade total JAK2 (37) and Ruxolitinib (Ruxo), currently in use as a cancer therapeutic, which acts on JAK1 and JAK2 by forming two hydrogen bonds with the hinge region of the JAK protein (the segment that connects the N-lobe to the C-lobe of the kinase domain) and reduces phosphorylation of the JAK2 protein at Tyr1007/1008 (31). Based on our earlier discovery that siRNA knockdown of STAT3 reverses BDNF-induced downregulation in the levels of *Gabra1* subunit mRNAs (23), as does WP1066, we used RNA-seq to identify the full set of genes responsive to JAK inhibition. This strategy enables us to unmask those genes in the BDNF transcriptome that may be JAK/STAT dependent.

Comparing the transcriptomes BDNF (V+B) or 100nM Ruxo + BDNF (RX1+B), we were surprised to find that there was no difference between these two transcriptomes, as statistically assessed using DSEQ2 (0 DEGs) (FDR=0.05, Wald test). To make sure that Ruxo treatment at 100nM was active in the primary cultures of our studies, Western blot was performed using the same drug stock as used in cultures that generated the RNA-seq libraries. In this control experiment, Ruxo (100nM) potently inhibited STAT3 phosphorylation at 30 min (80% +-3.07 reduction of BDNF signal) and at 4 hr (73% reduction of BDNF signal, Fig. 3). Therefore, despite reducing STAT3 phosphorylation at Tyr, our data taken together demonstrate that 100nM Ruxo does not alter BDNF-induced gene regulation, including changes in ICER expression (Fig. 3C). This finding suggests that the BDNF-induced JAK/STAT pathway is not STAT3 phosphorylation-dependent even though knockdown of STAT3 with siRNAs prevents ICER induction and *Gabra1* downregulation (23). Note that in our current studies BDNF does not significantly increase STAT3 phosphorylation but Ruxo still inhibits the basal state of its activation. These observations favor the hypothesis that neurons use a non-canonical mechanism of JAK/STAT signaling which may be relevant to epilepsy, and potentially to learning and memory, where BDNF is an essential signaling molecule.

**Figure 3.**
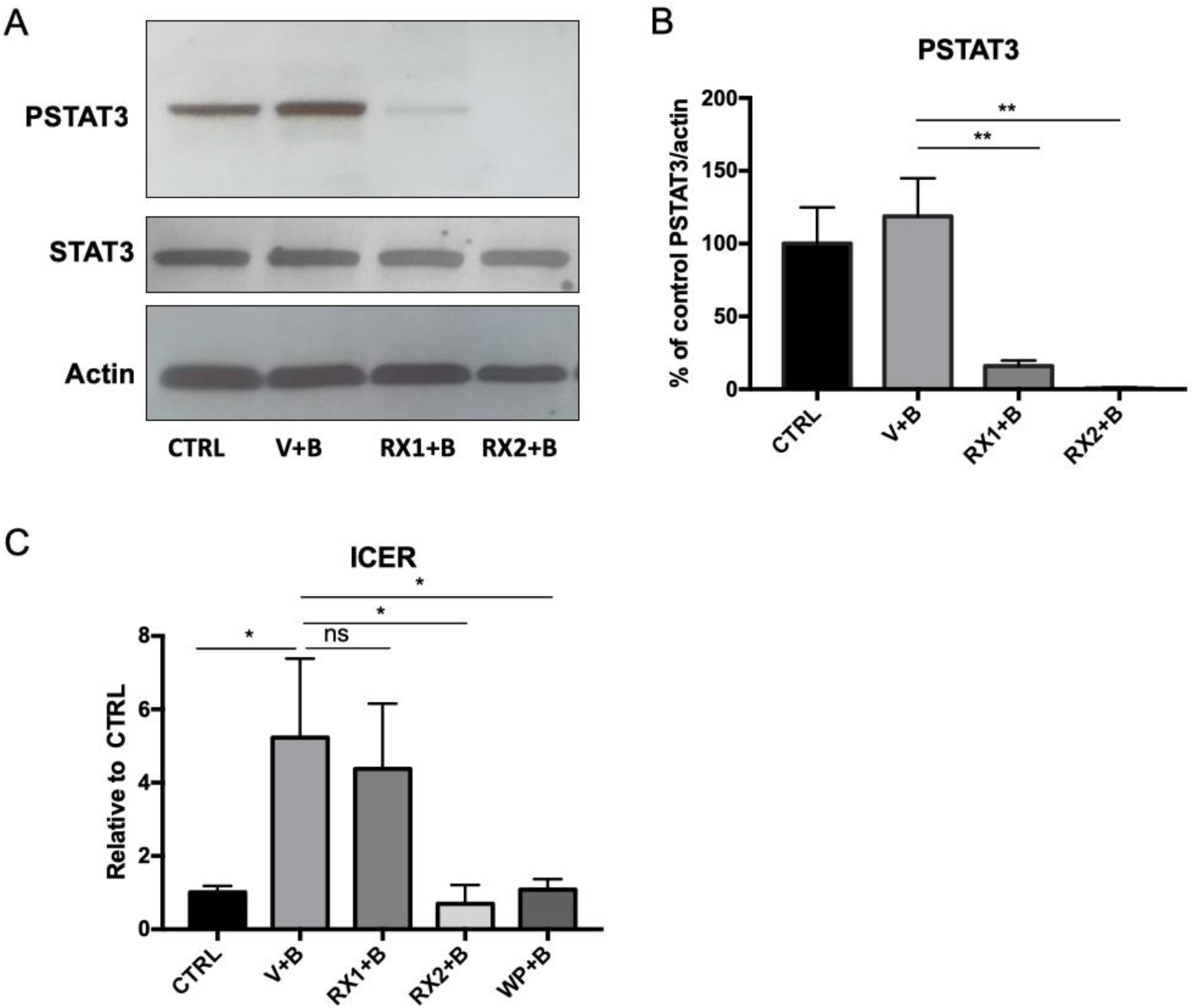
RX1 and RX2 reduce levels of phospho-STAT3 but only RX2 blocks ICER induction. (A) Representative Western Blot analysis of whole-cell protein extracts from primary neurons 9 DIV pretreated with100nM Ruxolitinib (RX1), 10uM Ruxo (RX2), or DMSO for 1 hr before the addition of BDNF. Cells were collected 30 min after BDNF administration and probed with anti pSTAT3, STAT3 and β-Actin. (B) Quantitation of signals from densitometry is displayed as mean percent change (±SEM) relative to the DMSO+Water control group (**p<0.01). Ruxolitinib significantly reduces the levels of P-STAT3 with no change in total STAT3. (C) Graphical representation of results from real time PCR analysis using specific Taqman probe and primers. RNA was extracted from cells collected 4 hr after BDNF administration. Transcript levels are shown as the mean values (±SEM) of the ratio relative to the DMSO+Water control group for *Icer* n=6. Note, that while RX1 blocks pSTAT3, it does not block the effect of BDNF treatment on *ICER* induction while WP and RX2 do.

### Using pharmacology to reduce the dimensionality of the dataset and identify potential BDNF-induced JAK/STAT target genes

While we discovered that co-treatment of neurons with BDNF (V+B) vs. RX1+B produced similar transcriptomes, we found that a 10-fold increase in the concentration of Ruxo (10µM) plus BDNF (RX2+B) attenuated the expression of multiple genes that were altered in response to BDNF, and most importantly, those that we previously identified in the PILO model of epilepsy. We next compared the RX2 DEG as determined by DESEQ2 (FDR=0.05, Wald test) to those generated from cultures treated with 10µM WP1066 plus BDNF (WP+B) and generated a subset of genes whose BDNF-induced expression was reversed back to vehicle control by both drug treatments. Using these JAK/STAT inhibitors as probes for unmasking potential genes that are regulated by the BDNF-induced JAK/STAT pathway, we identified 1559 of the 2659 BDNF target genes that may be most associated with JAK/STAT signaling in neurons. Interrogation of IPA networks specific to this shared gene set revealed that 131 genes are epilepsy-related and seizure disorder is still the top most significant neurological disease (P=7.31E-22). Considering that 131 of the 194 BDNF-regulated epilepsy-linked genes are reversed by JAK/STAT inhibitors (68%), it suggests that JAK/STAT signaling may be a large component of BDNF-related epileptogenesis. This can be best appreciated by looking at the heat map displayed in Figure 2C which depicts the expression levels of all 194 epilepsy-linked genes that are changed by BDNF treatment. The results clearly show that WP+B (10µM) and Ruxo (RX2+B, 10µM) block BDNF-induced changes in neuronal gene expression for epilepsy-linked genes and both appear to have a similar gene expression signature to that of DMSO+Water control. Note that Ruxo at its lower concentration (RX1+B, 100nM) looks similar to BDNF alone.

### Gene Ontology and Pathway Analysis

Gene Ontology (GO) term analysis of the WP+B/RX2+B dataset, performed using EnrichR (38,39), reveals that the 1559 overlapping genes are significantly associated with molecular functions related to receptor and channel activity (found in 8/10 top molecular functions) (Fig. 4A, top panel). All of the top ten biological process GO terms include the words “synaptic transmission” (Fig. 4A, middle panel). These consistent findings strongly suggest that both WP and Ruxo have the capacity to regulate synaptic transmission at the gene expression level in the brain. When compared with the total list of receptor/ion channel related genes that were regulated by BDNF (as depicted in Figure 2B, top panel), 107/133 genes (80%) were reversed by treatment with JAK/STAT inhibitors (Fig. 2B, bottom panel). This is best seen by the color reversal in the receptors for all but a few that were not significantly affected by pre-treatment with the inhibitor prior to BDNF activation (displayed in white).

**Figure 4.**
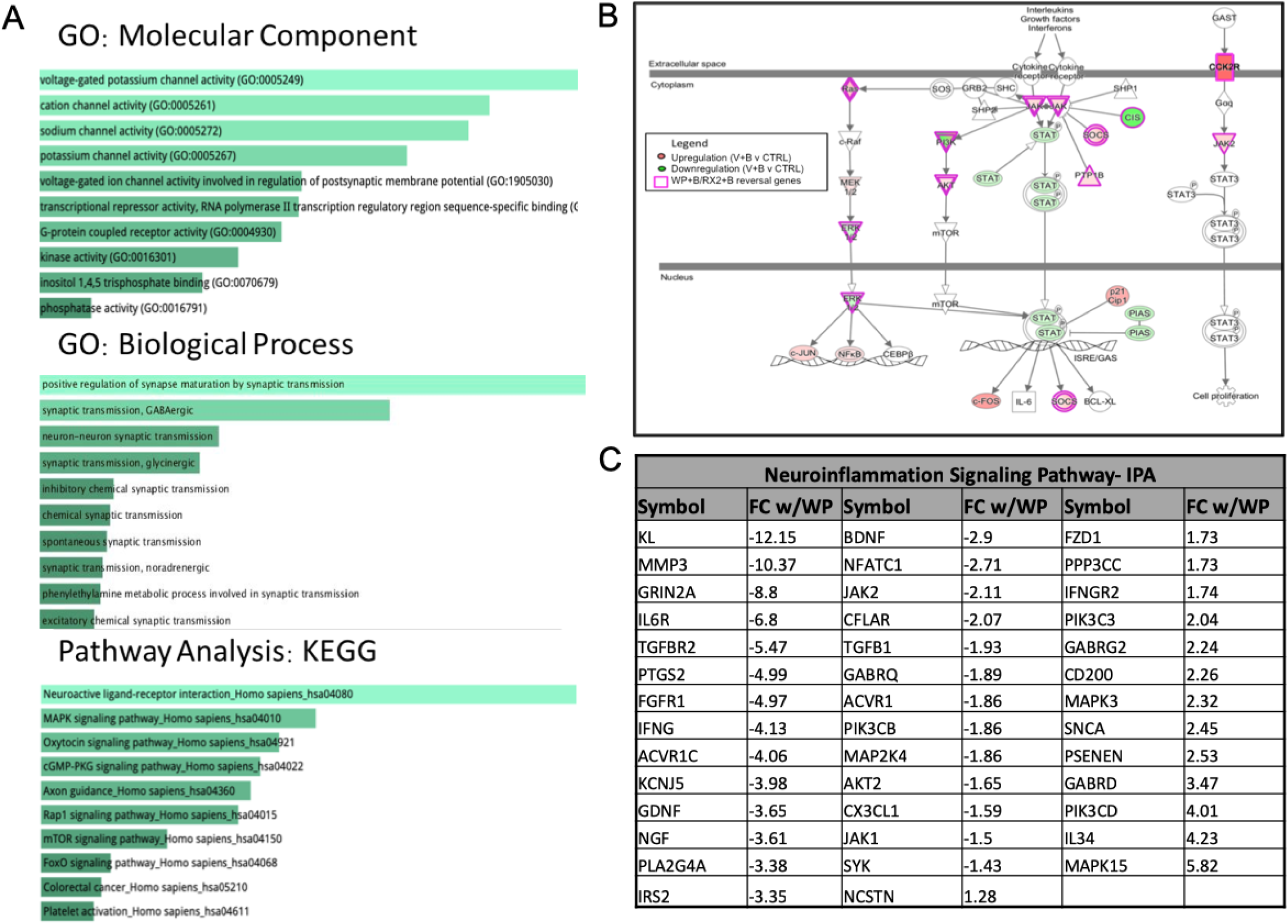
Pathway Enrichment and Gene Ontology analysis of the overlapping RX2/WP dataset. (A) Pathways and functions enriched in the list of 1559 genes altered by WP and 10µM Ruxo (RX2). Gene Ontology and KEGG enrichment analysis performed using EnrichR. (B) Genes involved in the canonical JAK/STAT signaling pathway with functional representation. Genes are colored by degree of fold change 9 with BDNF alone (see panel legend). Purple border signifies that WP+B and RX2+B reverse effects of BDNF on gene expression. (C) List of DEGs involved in neuroinflammatory signaling and degree of fold change of WP+B vs V+B. Listed in order of FC most negative to most positive.

In addition, neurotransmitter receptors, axonal guidance, GPCR and c-AMP signaling were found to be amongst the top KEGG pathways associated with the shared gene set of WP+B and RX2+B (Figure 4A, bottom panel). Members of the JAK/STAT signaling pathway were also identified as significant gene targets (P=1.044E-3) suggesting that BDNF may also participate in the feedback regulation of the pathway. Figure 4B is a schematic representation showing many of the component molecules that make up the JAK/STAT pathway, with expression of all the colored genes being affected by BDNF treatments and all the ones with a purple border reversed by JAK/STAT inhibitor treatments WP+B and RX2+B.

In addition to the significant relationship to epilepsy identified by IPA (see Additional Table 1), the overlapping WP/RX2 gene set had a significant link to neuroinflammation despite the neuronal cultures being largely devoid of any glial cells, 41 genes of the BDNF-induced set of 56 neuroinflammation-linked genes were reversed by WP+B or RX2+B treatment (73% of the gene set) (Fig. 4C).

### Functional relationships between genes as described in the literature

Pathway and GO analysis identified the following epileptogenic processes, in addition to inflammation, as being enriched in the WP+B/RX2+B overlapping data set: synaptic plasticity, receptors and ion channels, epilepsy, transcription factors, proliferation, and neurogenesis. In order to visualize the functional connectivity of the genes involved in epileptogenesis, we created a functional network of the most significant genes as determined by a p-value of <1×10^10 and a fold-change >2 (Additional Figure 1). Genes that met these conditions were separated into the lists of the epileptogenic processes, as described above. In order to relate these genes to one another and their epileptogenic functions, we created a network that portrays their function, expression fold change, direction, and functional protein relationships in Cytoscape (40)(Fig. 5A). Using GENEMania, a Cytoscape tool containing a database of experimental data showing protein-protein interactions (41), we determined potential functional relationships. Each node in the network represents a gene while the edges represents a protein interaction. Node color is determined by the epileptogenic category. Due to the interplay between these functions, many genes within one category overlap with another category. For instance, Egr3 plays a role in 5 of the 6 categories: synaptic plasticity, epilepsy, transcription factors, proliferation, and neurogenesis. The color in this case is determined by whichever category comes first in this list: synaptic plasticity, receptors and ion channels, epilepsy, transcription factors, proliferation, and neurogenesis, where the order is determined by the number of genes in that category from smallest to largest. This was done to avoid having the majority of the nodes colored for the largest category (neurogenesis). For Egr3, it is yellow representing synaptic plasticity, however, its location in the network is near the center because the proximity of the node to other categories represents its link to those secondary categories. In addition, the Venn diagram in 5B also shows the degree of overlap for each category. For instance, there are 108 total genes related to receptors and ion channels. In the list, 61 of those did not belong to any other category, 11 were also linked to neurogenesis, 5 to receptors, neurogenesis and proliferation,14 to receptors and epilepsy, 6 to receptors, epilepsy and neurogenesis, and 2 to receptors, neurogenesis, epilepsy and proliferation.

**Figure 5.**
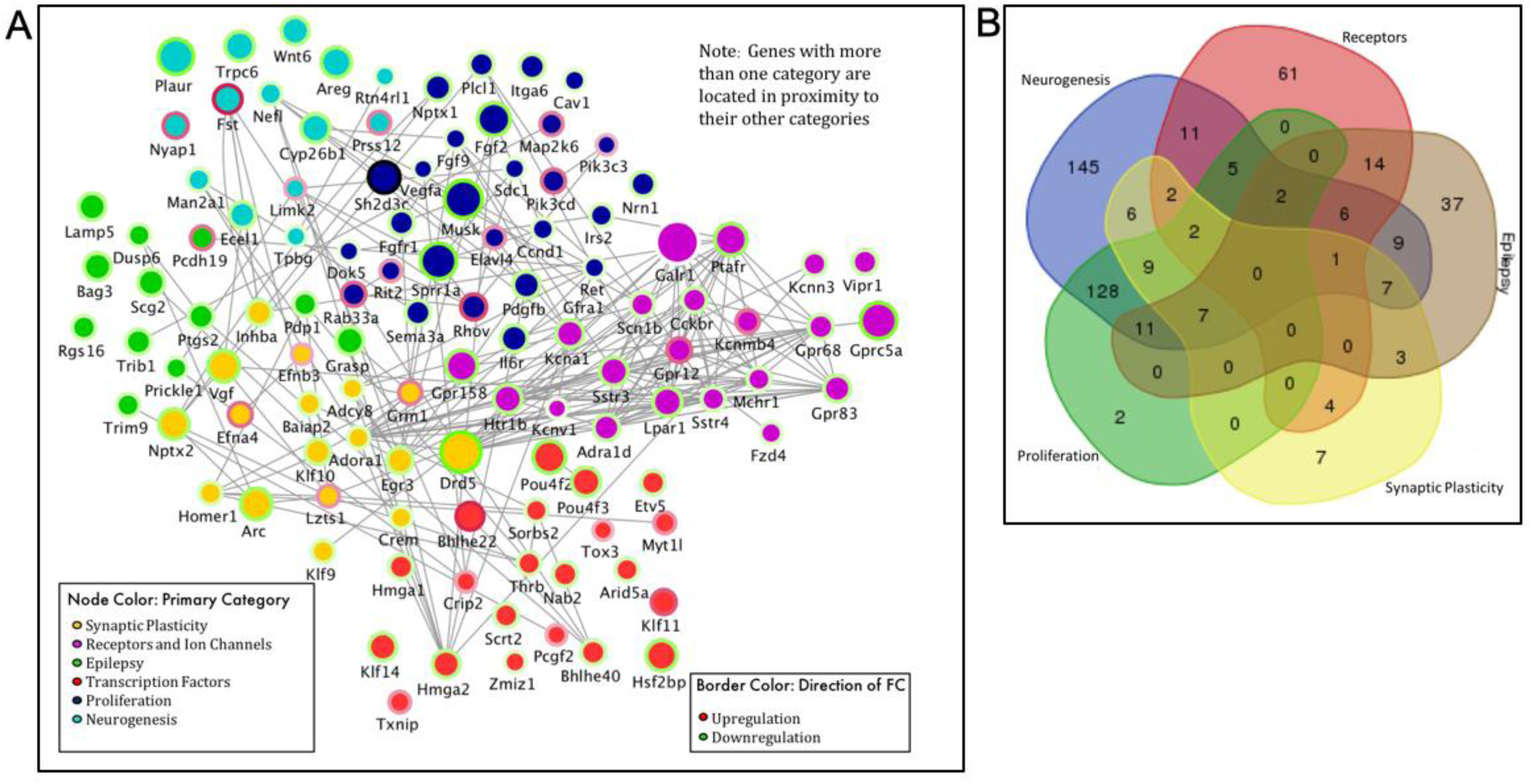
Functional Network of DEGs involved in epileptogenic-related processes. (A) Node color represents primary category, determined by order indicated in panel legend. Node size indicates degree of fold change (FC). Border color indicates direction of FC. Gray lines show functional connectivity determined by GeneMania in Cytoscape. Location in network roughly determined by category and multiple associations with neighboring and other categories. (Genes located near the center are involved in multiple categories). (B) Venn diagram of genes contained in network diagram in A demonstrating overlap of the genes in each category, neurogenesis (blue), receptors (red), epilepsy (gray), synaptic plasticity (yellow), and proliferation (green). Gene associations provided by IPA.

### Differentially expressed genes exclusive to BDNF/RX2 contain a rich set of epilepsy-associated targets not present in the WP1066 exclusive list

To further dissect the regulatory contribution of the two JAK/STAT inhibitors, we examined the list of DEGs (as determined by DESEQ2, FDR=0.05, Wald test) that differed between WP+B and RX2+B treatments. 417 DEGs are reversed by RX2+B treatment that are absent with WP+B and 610 DEGs are reversed by WP+B that are not by RX2+B (Fig. 2A, bottom panel). Interestingly, the genes whose expression is exclusively reversed by RX2+B are significantly related to neurodegeneration, and brain-related movement disorders (i.e., Huntington’s disease) as well as to epilepsy. On the other hand, those exclusively regulated by WP+B were not highly associated with epilepsy or other brain disorders. Additional Figure 2 contains the EnrichR generated KEGG pathway analysis for this gene set (Additional Figure 2A). 9 of the 10 top related canonical pathways differ from the KEGG list of overlapping WP/RX2 DEGs (Fig. 4B). Top genes involved in the KEGG pathways listed are also shown in a clustergram (Additional Fig. 2B). The list of genes exclusively regulated by RX2+B that are linked to epilepsy, Huntington’s disease, and neurodegeneration (E) are also presented (Additional Fig. 2C-E).

### qRT-PCR gene validation of datasets

Many of the genes identified in these RNA-seq studies were originally reported by us in the literature (*ICER, Gabra1, Egr3*), using a variety of molecular approaches (23,30,42). To further validate the new datasets, we generated an additional set of cultures, originating from the embryos of multiple pregnant rats, and chose the following genes for qRT-PCR because of their interest in the field of epilepsy: Dopamine Receptor D5 (*Drd5)*, Galinin Receptor 1 *(Galr1)*, γ2 subunit of the GABAR (*Gabrg2)*, Glutamate metabotropic receptor 1 (*Grm1)*, and their relationship to the JAK/STAT pathway: Inducible cAMP early repressor (*Icer)*, myeloid leukemia cell differentiation protein *(Mcl1)*, Suppressor Of Cytokine Signaling 3 *(Socs3)*, Cyclin D1 *(Ccnd1)*, and C-X-C Motif Chemokine Receptor 4 *(Cxcr4)* (Fig. 6). In most cases, findings from qRT-PCR were consistent with those of RNA-seq. Quantitated results from select validations are shown (Figure 6A-6D). A results table for all candidates where validation was performed is included in Figure 6E. All significance was determined using One-Way ANOVA followed by Tukey’s test for multiple comparisons.

**Figure 6.**
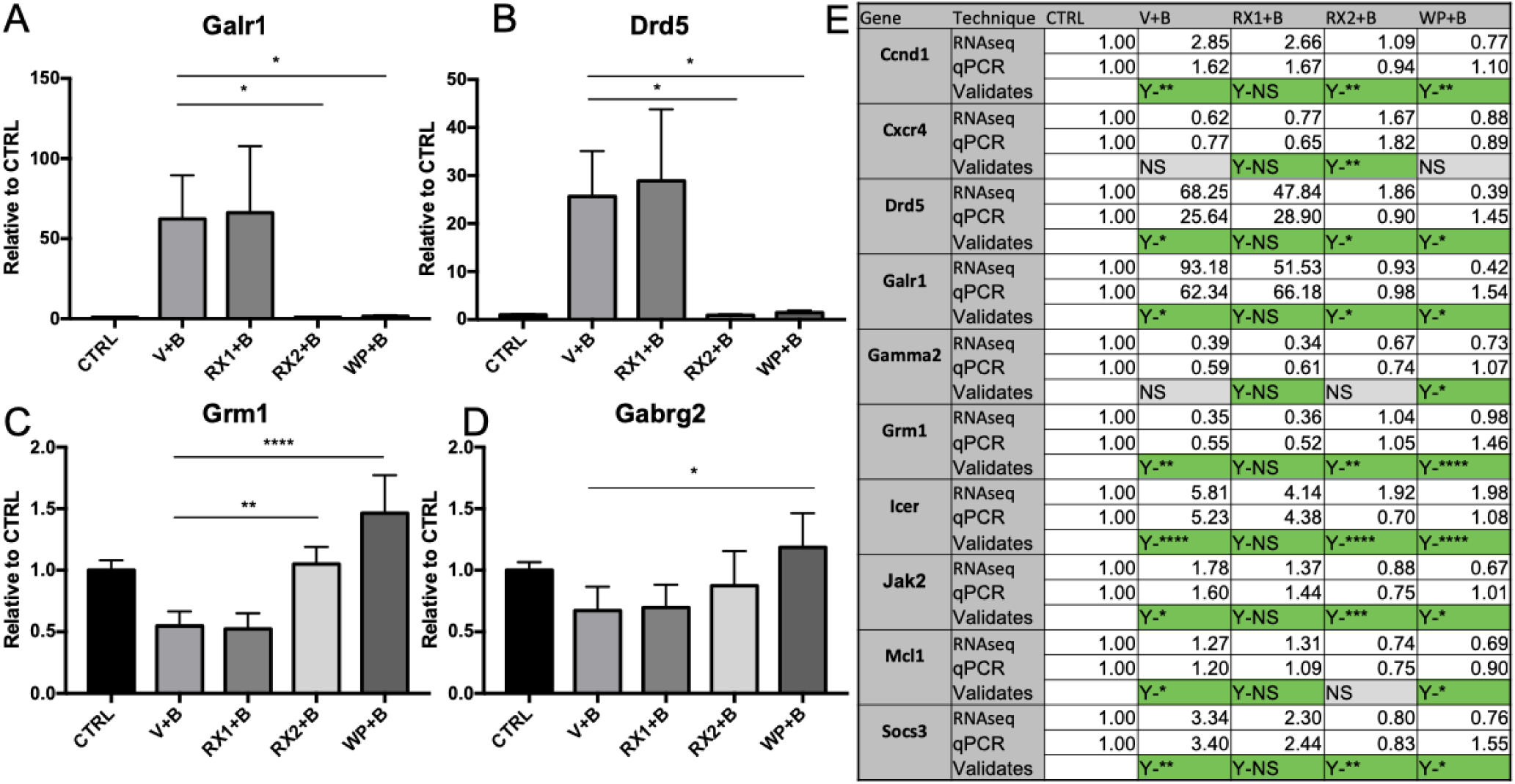
qRT-PCR validation of BDNF regulated genes in presence of JAK/STAT inhibitors. Primary neurons 9 DIV were pretreated with100nM Ruxolitinib (RX1), 10uM Ruxo (RX2), 10uM WP1066 (WP) or DMSO for 1 hr before the addition of BDNF. (A-D) Graphical representation of selected results of real time PCR analysis using Taqman probe and primers. RNA was extracted from cells collected 4 hr after BDNF administration. mRNA levels are shown as the mean values (±SEM) of the ratio relative to DMSO+Water control group for (A) Galr1 n=3, (B) Drd5 n=3, (C) Grm1 n=4, (D) Gabrg2 n=4. (E) Compilation of all genes tested for validation with qRT-PCR represented alphabetically. Values are the ratio relative to DMSO+Water for RNAseq (top row) and qRT-PCR (2^nd^ row) for comparison. Indication of a validating result (3^rd^ row) is marked in green. Results marked with Yes(Y) followed by the level of significance indicate a significant result consistent with RNA-seq. Yes-Not significant (Y-NS) seen in all the RX1 is the expected result consistent with our RNA-seq findings. Measurements that did not reach significance are marked in gray as Not Significant (NS). Significance was tested using One-Way ANOVA followed by Tukey’s test for multiple comparisons. (*p<0.05, **p<0.01, ***p<0.0005, ****p<.0001).

### Comparison of DEGs with ChIP-seq datasets

Using a list obtained from previously published chromatin immunoprecipitation (ChIP)-sequencing (ChIP-seq) of STAT3 target genes, identified in gliablastoma cells (43), we generated a list of 308 genes from the overlapping WP+B/RX2+B dataset that have been previously shown to have functional binding sites for STAT3 (Fig. 7). We were particularly interested that *BDNF*, Calcium Voltage-Gated Channel Auxiliary Subunit Beta 4 (*Cacnb4), Gabrg2, Grm1, Jak1/2*, and Sodium Voltage-Gated Channel Alpha Subunit 1(*Scn1a)* were on the list. They have all been associated with epilepsy models and with human mutations in epilepsy patients (33,44–46).

**Figure 7.**
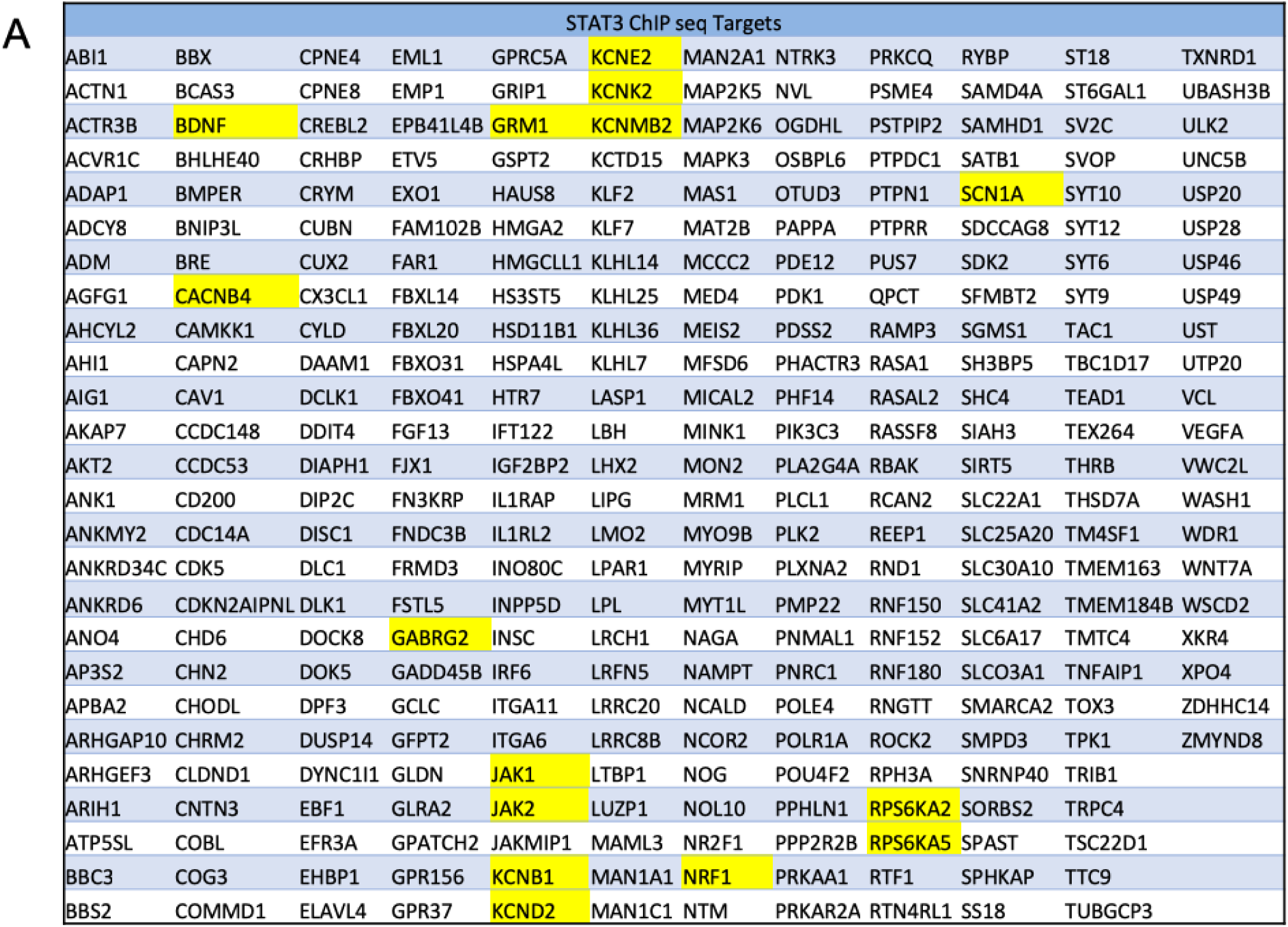
ChIP-seq identifies multiple genes in the WP+B/RX2+B overlapping datasets that contain validated STAT3 binding sites. List of all WP+B/RX2+B genes (308) that are validated STAT3 ChIP-seq targets (as determined by Zhang, et.al, 2013) in alphabetical order. Highlighted in yellow are genes of special interest.

## Discussion

There is a strong body of evidence suggesting that BDNF signaling through its TrkB receptor contributes in a major way to the development of TLE, as suggested by animal models of epilepsy (11,47) and their potential relationship to human epilepsy brain pathology (48). BDNF plays a critical role in the cell biological processes that regulate the structural plasticity of dentate granule cells and the enhanced excitation associated with changes in LTP and compromised GABA-mediated inhibition (17,44,47,49). Although the role BDNF plays in brain disorders is better understood now than at the time of its discovery, many key aspects of its complex regulatory programs remain a mystery. We designed our study to gather a comprehensive list of genes whose expression is either upregulated or downregulated by increased exposure to BDNF, as is believed to occur in response to seizures and other brain insults such as traumatic brain injury (50).

The use of primary rat neurons treated with recombinant BDNF allows us to pinpoint the effects of BDNF without the background of additional signaling pathways that may be activated in TLE models. Our study provides a comprehensive list of the changes in the transcriptome that occur in response to a relatively short but continuous exposure to BDNF (4 hr) at a concentration of 0.7 nM that is relevant to endogenous levels in neurons that activate TrKB receptors and not p75NTR (unpublished data from our laboratory). Many of the BDNF regulated genes within the list have been previously identified both in treated cultures and *in vivo* by our laboratory and others. They include *Gabra1*, the inducible cAMP early repressor (*ICER*) (which negatively regulates *Gabra1)* (23), *BDNF* itself (36), and multiple ion channel genes, as well as genes that code for important markers of neurogenesis and cell proliferation (51,52).

In addition to being consistent with the findings in the literature, our results reveal new BDNF-induced gene targets that are linked to epilepsy. Continuous exposure to BDNF alters the expression of 194 epilepsy-linked genes identified by IPA. The following genes are of particular interest to us because of their marked level of transcriptional alteration and their involvement in epilepsy:

- Dopamine receptor D5 (*Drd5)-A* D1-type dopamine receptor whose knockdown has been shown to be protective against seizures (53). In our dataset, the neuronal expression of Drd5 increases nearly 15-fold in response to BDNF treatment.
- Galinin receptor 1 (*Galr1*)-Galanin is a neuropeptide in the brain that inhibits glutamate release and galanin agonists delivered to the brain can inhibit seizures (54). Our dataset reveals a 35-fold increase in *Galr1* expression, *de*monstrating the complex medley of both protective and pathogenic changes in genome expression that are invoked by increasing levels of neuronal BDNF.
- Glutamate metabotropic receptor 1 (*Grm1*)-*Grm1* has been associated with the alterations in synaptic plasticity and glutamate disturbances that result in neuronal overexcitation, a hallmark of epilepsy (46,55). *Grm1* mutations can also cause spinocerebellar ataxia type 44, a genetic disease that can present with seizures. In our dataset, BDNF treatment led to a 2.45-fold increase in the expression of *Grm1.*
- γ2 subunit of the GABAR (*Gabrg2*)-the *Gabrg2* subunit of GABAR is important for benzodiazepine action and mutations lead to epilepsy (56–59). In this study, RNA-seq revealed a 2.38-fold decrease in the expression of *Gabrg2* in response to BDNF.

Our new results suggest that sustained levels of BDNF at endogenous concentrations (0.7 nM) can dysregulate a major segment of the epilepsy-associated transcriptome. Recent discovery of the role of BDNF in cancer suggests that understanding how BDNF orchestrates its diverse effects on genome dysregulation will also provide a new window on cancer chemotherapies (60–62). Surprisingly, beyond its activation of downstream targets like the cAMP regulatory element binding (CREB) protein, whose relationship to BDNF has been rigorously described (63,64), we still know little about the role of other BDNF controlled transcriptional regulators and their intracellular signaling partners that contribute to disease.

Using the TLE model of epilepsy, we previously showed that during the latent period after pilocarpine-induced SE, there is a marked upregulation in activated CREB and an induction in the expression of the inducible cAMP early repressor (ICER) (23). We also reported that activated CREB interacts with ICER to downregulate the levels of *Gabra1* transcripts leading to a decrease in α1-containing GABARs in hippocampal neurons. In our studies, we discovered that BDNF was linked to the JAK/STAT pathway as 1) STAT3 knockdown blocked ICER induction in response to BDNF, 2) STAT3 was found on the ICER promoter in response to BDNF treatment (as measured by chromatin immunoprecipitation (ChIP), 3) JAK/STAT inhibitors (pyridone 6 and WP1066) blocked ICER induction and *Gabra1* downregulation, and 4) *in vivo* application of WP1066 at the time of pilocarpine-induced SE reduced the frequency of subsequent spontaneous seizure animals from pilocarpine-induced spontaneous seizures after SE (30).

Based on these findings, we asked whether the BDNF transcriptome, as identified in our prior studies, could be partitioned into those where the BDNF response could be blocked by exposure to JAK/STAT inhibitors (Fig. 1). We chose two inhibitors, WP1066 (a JAK2 inhibitor with diverse mechanisms of inhibition (37)), that we showed was active in primary cultures and *in vivo*, and Ruxolitinib (Ruxo), a high-affinity JAK1/2 inhibitor that blocks JAK activation and is currently used in the treatment of patients with high-risk myelofibrosis. To our surprise, when we compared the transcriptome of cells exposed to BDNF, and to low dose Ruxo plus BDNF (RX1,100nM+B), we did not see a significant difference between the two datasets. As we show in Figure 3, this was not due to any issue with the ability of the stock drug to block STAT3 phosphorylation. Phosphorylation-independent STAT3 activity, however, has been described in multiple contexts (65,66) suggesting that JAK/STAT signaling in neurons may act in a non-canonical fashion, at least when activated by prolonged BDNF signaling as studied in our research. A role for unphosphorylated STATs in genome stability and for nuclear JAK2 in the regulation of histone 3 Y41 phosphorylation has been described (67,68) and is the subject of our future studies.

Despite the lack of response of the transcriptome to BDNF in the presence of the high affinity concentration of Ruxo (RX1, 100nM), WP1066, and a ten-fold increase in Ruxo (RX2, 10µM) produced substantial changes in gene expression (Fig. 2A) suggesting the increase in concentration of Ruxo engages a distinct mechanism of action. By comparing the two datasets (WP1066+B and RX2+B) we were able to decrease the dimensionality of the BDNF transcriptome into those most relevant by their association with epilepsy. In fact, using enrichment analysis, we discovered an association with epilepsy that is more pronounced than any other neurological disease, due to the large proportion of differentially expressed epilepsy-associated genes present within the WP+B and RX2+B subset (68%) (see Additional Table 2).

Pathway analysis also revealed that JAK/STAT inhibition had a marked effect on reversing the BDNF-induced transcriptional response for genes organized in pathways that control neurogenesis, neuroinflammation, synaptic plasticity, receptor expression, and synaptic transmission. Alterations in these functions exist in the epileptogenic brain, which is known to have altered neurogenesis, mossy-fiber sprouting, inflammation and neuronal pruning leading to increased plasticity, and altered excitation and inhibition at the synapse (2,3,69). They also expand upon the recent findings that JAK/STAT signaling plays an important role in learning and memory (25). In the future, it will be important to study these changes in the context of an epileptogenic animal model where STAT3 expression is modified in both neurons and glia.

Given that multiple epilepsy-associated genes have been shown to have functional binding sites for STAT3, including BDNF itself (Fig. 7), and considering the substantial relationship between prolonged exposure to BDNF and the ability of JAK/STAT inhibitors to reverse the differential expression of epilepsy associated genes in our studies, we hypothesize that identifying the mechanism that connects BDNF signaling to this novel non-canonical role of JAK/STATs in neurons has the potential to reveal new strategies in the treatment of intractable epilepsy for human patients and provide a new window on how neuropharmacology can be used to identify the complex cross-talk between different intracellular signaling pathways.

## Conclusion

We have employed deep RNA-seq to determine the response of BDNF on the transcriptome of neurons and have employed a neuropharmacological approach to reduce the dimensionality of the datasets to identify BDNF responsive genes whose transcriptional polarity is reversed by two well characterized JAK/STAT inhibitors. Our results show that this subset of genes contains many with previous association to epilepsy in both animal models and human patients. The BDNF-induced JAK/STAT gene set is highly enriched for genes involved in synaptic neurotransmission, and contains targets from all the major classes of ion channels and neurotransmitter receptors of the brain. In particular, the gene set includes those known to play a role in epileptogenesis through the regulation of synaptic plasticity, neurogenesis, transcriptional regulation, neuroinflammation and proliferation. Some of the genes are known to contain functional STAT recognition sites as verified by ENCODE, while others remain to be interrogated. Pharmacological analysis also revealed that phosphorylation of STAT3 at Tyr705 most likely does not control BDNF-induced JAK/STAT regulation of genome expression, suggesting that the mechanism is non-canonical. Most importantly, the RNA-seq datasets highlight the importance of the JAK/STAT pathway in neurons and stimulate further discovery in this area based on its potential relevance to identifying new therapeutic strategies for the treatment of intractable neurological disorders.

## Methods

### Cell culture and treatments

Primary neocortical neurons were dissected from neocortex of 3 E18 Sprague-Dawley rat embryos (Charles River Laboratories). Pregnant dams were euthanized by CO 2 according to approved Boston University Institutional animal care and use (IACUC) protocol (AN14327). Embryonic brains were removed and placed in ice-cold Ca 2+ /Mg 2+ free (CMF) media [Ca 2+ /Mg 2+ free Hanks BSS, 4.2mM sodium bicarbonate,1mM pyruvate, 20mM HEPES, 3mg/mL BSA, pH 7.25-7.3]. Cortices were dissected under a dissection microscope, trypsinized centrifuged, and triturated in plating media [Neurobasal media (Invitrogen), 10% fetal bovine serum (Gibco), 100 U/mL penicillin, 100 μg/mL streptomycin, 200mM glutamax (Gibco)]. Dissociated neurons were plated on Poly-L-lysine coated 6-well plates (1×10^6^ cells/well) and placed in an incubator (37°C/5% CO 2) for attachment to plate. 1hr later, plating media was removed and replaced with 2mL of defined medium [Neurobasal media (Invitrogen), B27 serum-free supplement (Gibco), 100 U/mL penicillin, 100 μg/mL streptomycin, 200mM glutamax (Gibco)]. Neurons were cultured in the incubator for 9 days until use (9 days in vitro [DIV]), cultures are grown under conditions to minimize the growth of non-neuronal cells, >= to 90% neurons or precursor neurons. Treatments: At 9DIV cells were pretreated for 1-hr. with vehicle dimethyl sulfoxide (DMSO) at 0.1%, Ruxolitinib (Sellckchem S1378; 100nM or 10uM), or WP1066 (EMD Millipore 573097; 10uM). Nuclease-free water vehicle or Brain-derived neurotrophic factor was then added (EMD Millipore GF029; 0.7nM) for 4 hours before collection. N=3, each N was cells collected from one pregnant dam. Enough primary cells were collected from each N sufficient for 1 biological replicate of each treatment.

### RNA extraction and library preparation

At the end of the 4-hr treatment period, cells were collected and RNA extracted using the Qiagen RNeasy Micro Kit with DNAase treatment (Cat. 74004). RNA was run in the Agilent Bioanalyzer to determine RNA integrity number (RIN). RIN of all samples was ≥ 9.0. mRNA was selected by using the NEBNext® Poly(A) mRNA Magnetic Isolation Module (E7490S) before using the NEBNext^®^ Ultra™ Directional RNA Library Prep Kit for Illumina^®^ (E7420S) with Agencourt AMPure XP Clean up beads (A63881) to make RNA into strand specific cDNA libraries with multiplexing barcodes from the NEBNext® Multiplex Oligos for Illumina® (Index Primers Set 1) kit (E7335S).

### Illumina Sequencing

Sequencing was performed at the Boston University Microarray Core and University of Chicago Genome Center. Fragmentation and concentration were analyzed on the Agilent Bioanalyzer and Fragment analyzer prior to sample pooling. Runs were either sequenced at 75 bp single-end reads on the Illumina NextSeq sequencer to a depth of 100-120 million reads per sample or at 50 bp single-end reads on the Illumina HiSeq sequencer to a depth of 90-100 reads per sample. All reads were conducted in one lane using barcodes to separate biological replicates.

### RNA-seq bioinformatics

Data analysis was performed using Strand NGS software, Version 3.2, Build 237248. © Strand Life Sciences, Bangalore, India. Proprietary Strand NGS algorithms were used to conduct read alignment to the rat genome build rn6 RefSeq Genes (2016.05.11). Reads were filtered based on the following parameters: Max number of novel splices: 1, Min alignment score: 90, Number of gaps allowed: 5, number of matches to be output for each read: 1, exclude reads with alignment length less than: 25, ignore reads with matches more than: 5, Trim 3’ end with average base quality less than: 10. RNA quantification was performed by the Strand NGS software, which counts the total number of reads that map to each gene and exon in the genome, and is reported as raw counts. Normalization of the raw counts was achieved using the DESeq method, which accounts for difference in the total number of reads between samples and log transforms the data (70). R version 3.1.2 was used with DEseq2 version 1.6.1 to determine differential expression FDR=0.05, Wald test type. The list of significant genes was then used in subsequent analyses. Disease involvement data was analyzed through the use of IPA with proprietary statistical algorithms (QIAGEN Inc., https://www.qiagenbioinformatics.com/products/ingenuitypathway-analysis). Cytoscape 3.5.1 software was used with GeneMANIA for functional network generation with (40,41). EnrichR was used for GO-term analysis and KEGG pathway analysis (38,39).

### Real-time quantitative reverse transcriptase PCR (qRT-PCR)

Primary neurons were prepared, treated, and RNA extracted, as described above for RNA-seq. The Applied Biosystems 1-step RNA to Ct (cat. 4392938) kit was used and protocol was followed as directed: 50 ng of RNA template, 10 μL of 2x mastermix, 0.5 μL of each 40x primer/probe kit (Ppia and the gene of interest), 0.5 μL of the enzyme mix, and nuclease free water was added to each 20 μL reaction. Each reaction was performed with 2 technical replicates in a 384-well plate. All taqman primer/probe kits were ordered from Thermo Fisher Scientific.

| Gene Symbol | Assay ID |
| --- | --- |
| Ppia (housekeeping) | **Rn00690933_m1 VIC-MGB** |
| Galr1 | **Rn04932425_m1 FAM-MGB** |
| Drd5 | **Rn00562768_s1 FAM-MGB** |
| Jak2 | **Rn00676341_m1 FAM-MGB** |
| Gabrg2 | **Rn01464079_m1 FAM-MGB** |
| Grm1 | **Rn00566625_m1 FAM-MGB** |
| Icer | **Rn00569145_m1 FAM-MGB** |
| Ccnd1 | **Rn00432359_m1 FAM-MGB** |
| Icer | **Rn00569145_m1 FAM-MGB** |
| Socs3 | **Rn00585674_s1 FAM-MGB** |
| Mcl1 | **Rn00821024_g1 FAM-MGB** |

The Applied Biosystems 7900 HT Real-Time PCR system was used to run with the following cycling parameters: 48°C 15 min. hold (Reverse Transcription), 95°C 10 min. hold, 40 cycles: 95°C 15 sec, 60°C 1 min. After run completion, the ΔΔCT method was used to calculate the relative values of the transcripts.

### Western Blot

Primary cortical cultures used for Western Blot were prepared and treated with drugs the same as described for RNA-seq. After the 4-hr period of BDNF treatment, cells were washed once with ice-cold PBS solution (PBS+EDTA, 1x PhosStop (Roche) and 1x Protease Inhibitor cocktail (Roche cOmplete protease inhibitor)) and centrifuged in 1.5 mL tubes at 5000RPM for 4 min at 4°C. PBS was removed and cell pellet was resuspended in 1x RIPA buffer (with 1x Protease and Phosphatase inhibitors). Lysates were incubated on a rotator at 4°C for 15 min, then centrifuged at 13000 RPM for 10 min. Supernatant was transferred to a fresh tube and aliquoted for Western Blot. 35μg of sample was diluted to 10μL in RIPA buffer and 10μL of 2x SDS Tris-Glycine loading buffer (Invitrogen LC2676) with 50mM DTT. Samples were boiled at 95°C for 5 min, centrifuged and loaded into a Novex 10% Wedgewell gel (Invitrogen XP00100BOX). The gel was run at 220V for 40 mis and wet transfer was performed at 33V for 2 hrs to a nitrocellulose membrane. Membrane was quickly washed in TBST then blocked in 5% milk in TBST for 1 hr at room temperature. Three 5 min washes in TBST were performed. Membrane was incubated overnight at 4°C in primary antibody then washed 3×10 minutes in TBST. HRP-conjugated secondary was added for one hour at room temperature, then 3 washes in 1% milk were performed prior to imaging the blot with ECL (BioRad Clarity #1705060). Membrane was stripped using the GM Biosciences One Minute Advance Stripping buffer #GM6031. Antibodies were used as follows: 1:1000 *pSTAT3* (CST #9131S) in 5% BSA, 1:5000 anti-rabbit HRP-conjugated secondary antibody (EMD Millipore AP132P) in 1% milk in TBST. 1:2000 total *STAT3* (CST #4904S) in 5% milk in TBST. 1:6000 <u>β-actin</u> (Sigma A5441) in 1% milk in TBST, 1:2500 Anti-mouse HRP-conjugated secondary antibody (Vector Labs PI2000) in 1% milk in TBST.

## Declarations

### Abbreviations

AMPA: α-amino-3-hydroxy-5-methyl-4-isoxazolepropionic acid
BDNF: Brain-derived neurotrophic factor
ChIP: Chromatin Immunoprecipitation
DEG: Differentially expressed genes
*Drd5*: Dopamine receptor D5
ERK: Extracellular-signal-regulated kinase
GABA: Gamma-Aminobutyric Acid
*Gabra1*: Gaba A receptor alpha 1 subunit
*Gabrg2*: Gaba A receptor gamma 2 subunit
*Galr1*: Galinin receptor 1
GO: Gene ontology
GPCR: G-protein-coupled receptor
*Grm1*: Glutamate metabatropic receptor 1
IACUC: Institutional animal care and use
*Icer*: Inducible cyclic-AMP early repressor
IPA: Ingenuity pathway analysis
JAK/STAT: Janus kinases (JAK)
JNK-c: Jun N-terminal kinase
LTD: Long-term depression
LTP: Long-term potentiation
MAPK: Mitogen-activated protein kinase
mRNA: messenger ribonucleic acid
NF-κB: nuclear factor kappa-light-chain-enhancer of activated B cells
NGFR: Nerve growth factor receptor
P75NTR: p75 Neurotrophin receptor
PI3K: Phosphoinositide 3-kinase
PILO: Pilocarpine
PLCγ: Phospholipase C gamma
qRT-PCR: quantitative real-time polymerase chain reaction
RUXO: Ruxolitinib
SE: Status Epilepticus
STAT: Signal Transducer and Activator of Transcription
TBI: Traumatic brain injury
TLE: Temporal lobe epilepsy
TrkB: Tropomyosin receptor kinase B

### Ethics approval and consent to participate

Rats were sacrificed in accordance with the Boston University practice for laboratory animal procedures. The protocol (AN14327) was approved by the Boston University Institutional Animal Care and Use committee (IACUC)

### Consent for publication

All authors consent for publication

### Availability of data and material

All data generated and analyzed during our study which support the conclusions of the article are included in the manuscript with its supplemental files. Full datatsets from RNAseq will be deposited in public information databases upon acceptance for publication.

## Funding

Funding was generously provided by the following grants:

NIGMS T32GM008541

NINDS R01NS051710

NINDS R21 NS083057

CURE Epilepsy Multidisciplinary Award

## Authors’ contributions

KH performed the experiments, analysis, and wrote the manuscript with input from all authors. MC assisted with study design and data analysis. ABK assisted with study design and results interpretation. SR oversaw study design, experimental work, data analysis and read, edited and wrote the final manuscript. All authors read and approved the final manuscript.

## Acknowledgements

Not applicable

---

**Supplemental Table 1.** Genes of interest in the dataset and their relationship to epilepsy. Several genes of particular interest to the field of epilepsy are listed with the degree of FC from DMSO+Water vs. DMSO+BDNF and the FC of DMSO+BDNF vs. WP1066+BDNF. Many have not been associated with the JAK/STAT pathway or BDNF regulation. The name of the gene, the related function, and a reference to support the potential relationship with epilepsy is listed.

| Relationship of Epilepsy Genes with JAK/STAT Inhibition |  |  |  |  |  |
| --- | --- | --- | --- | --- | --- |
| Gene of Interest | FC V+B | FC WP+B | Name | Function | References |
| GALR1 | 34.36 | -115.70 | Galanin receptor 1 | Protective in Epilepsy | Mazarati, 2004 |
| BHLHE22 | -5.1 | 7.78 | Basic Helix Loop Helix E 22 | Repressor involved in neurodevelopment | Joshi, 2008 |
| SOCS3 | 2.89 | -3.29 | Suppressor of Cytokine Signaling 3 | Inhibits JAK signaling, increased in epilepsy | Pernot, 2011 |
| DRD5 | 14.91 | -63.92 | Dopamine Receptor 5 | Knockout protects from seizures | O'Sullivan, 2008 |
| INHBA | 5.26 | -5.27 | Inhibin Beta A | Reduces excitotoxicity | Lau, 2015 |
| NPTX1 | 5.01 | -6.30 | Neuronal Pentraxin 1 | Plays role in epileptogenic synaptic remodeling | Sanabria, 2008 |
| GRIN2A | 2.18 | -8.80 | Glutamate Ionotropic Receptor NMDA Type Subunit 2A | Mutations cause epilepsy, involved in LTP | Carvill, 2013 |
| CXCR4 | -1.53 | 1.77 | CXC Motif Chemokine Receptor 4 | Inhibition protects against epilepsy | Song, 2016 |
| HRH3 | -3.93 | 6.53 | Histamine H3 Receptor | Increased Histamine confers protection in epilepsy | Bhowmik, 2012 |
| RGS4 | 4.46 | -2.77 | Regulator of G protein signaling 4 | Inhibition protects against epilepsy | Chen, 2012 |
| SH2D3C | -11.46 | 14.91 | SH2 Domain Containing 3C | Involved in cell migration | Wang, 2012 |
| ELAVL4 | -2.04 | 2.76 | ELAV Like RNA Binding Protein 4 | Controls mRNA processing, increased in Epilepsy models | Bronicki, 2013 |
| FYN | 1.35 | NA | FYN Proto-Oncogene, Src Family Tyrosine Kinase | Overactivity leads to epilepsy, activates mTORC1 pathway | Kojima, 1996; Wang, 2015 |
| RPS6K(A2, A3, A5, A6, B1, B2) | A2--2.04 | A2-2.59 | Ribosomal Protein S6 Kinase | mTOR activated, involved in proliferation and cell growth | Cho, 2011 |
| ARC | 33.24 | -15.04 | Activity Regulated Cytoskeleton Associated Protein | Known to be seizure and BDNF induced, plasticity | Kulpers, 2016 |
| BDNF | 3.23 | -2.90 | Brain Derived Neurotrophic Factor | Increased in epilepsy, neuroplasticity | Binder, 2001 |
| JAK2 | 1.79 | -2.11 | Janus Kinase 2 | Activated by BDNF and Status Epil. | Lund, 2008 |
| EGR3 | 5.57 | -6.08 | Early Growth Response 3 | Regulates $\alpha 4$ subunit GABA<sub>A</sub>R | Roberts et al, 2006 |

**Supplemental Table 2.**
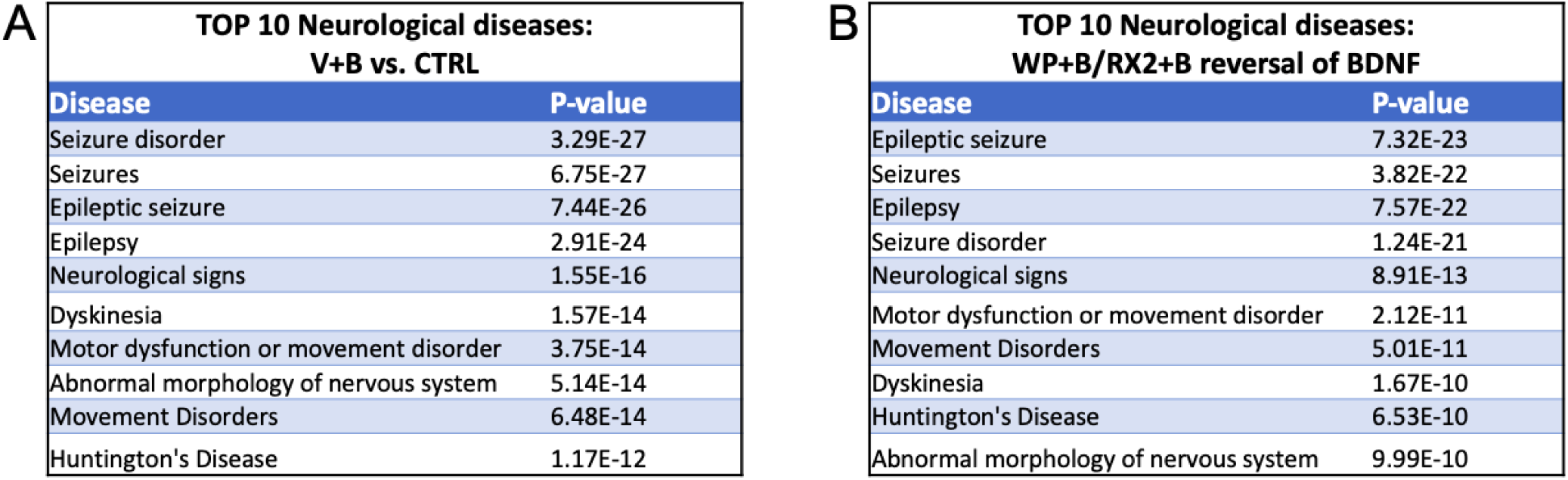
Top Neurological disease relationships determined by IPA. (A) Top 10 significantly enriched neurological diseases in the BDNF transcriptome with their p-value (IPA). (B) Top 10 significantly enriched neurological diseases in the WP/RX2 vs. BDNF set of differentially expressed genes listed with their p-value from IPA

| TOP 10 Neurological diseases:<br>V+B vs. CTRL |  |
| --- | --- |
| Disease | P-value |
| Seizure disorder | 3.29E-27 |
| Seizures | 6.75E-27 |
| Epileptic seizure | 7.44E-26 |
| Epilepsy | 2.91E-24 |
| Neurological signs | 1.55E-16 |
| Dyskinesia | 1.57E-14 |
| Motor dysfunction or movement disorder | 3.75E-14 |
| Abnormal morphology of nervous system | 5.14E-14 |
| Movement Disorders | 6.48E-14 |
| Huntington's Disease | 1.17E-12 |

**B**
| TOP 10 Neurological diseases:<br>WP+B/RX2+B reversal of BDNF |  |
| --- | --- |
| Disease | P-value |
| Epileptic seizure | 7.32E-23 |
| Seizures | 3.82E-22 |
| Epilepsy | 7.57E-22 |
| Seizure disorder | 1.24E-21 |
| Neurological signs | 8.91E-13 |
| Motor dysfunction or movement disorder | 2.12E-11 |
| Movement Disorders | 5.01E-11 |
| Dyskinesia | 1.67E-10 |
| Huntington's Disease | 6.53E-10 |
| Abnormal morphology of nervous system | 9.99E-10 |

**Supplemental Figure 1.**
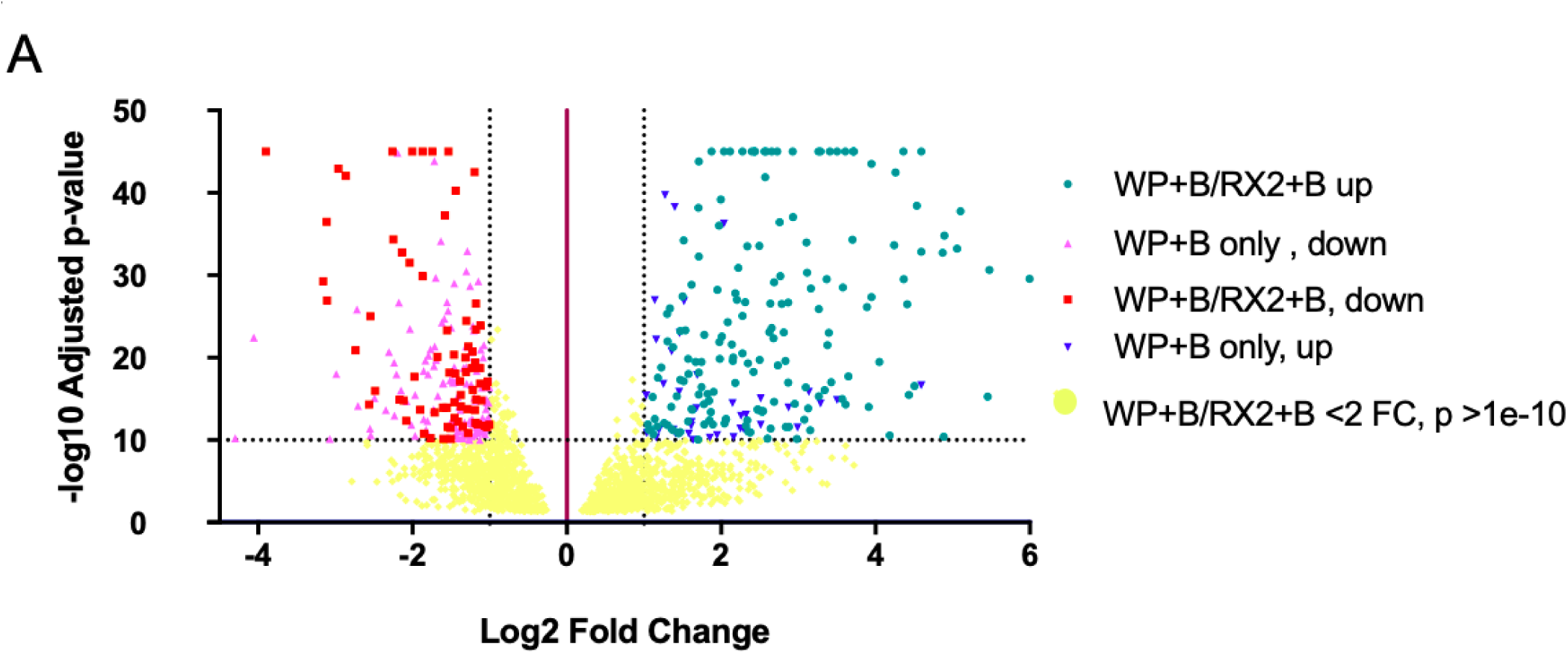
Volcano Plot of BDNF regulated genes whose response is reversed by WP+B/RX2+B. Representation of degree of fold change and significance used to narrow down the list of top genes to include in the Figure 3 network. Teal-genes upregulated by both WP+B and RX2+B; Blue: genes upregulated by WP+B only; Red: genes downregulated by both WP+B and RX2+B; Pink: genes downregulated only by WP+B; Yellow: genes that did not meet cutoffs of >2 FC, p<1e-10. Teal and Red gene list used for Figure 3 network. Maximum –log10 p value detected was 45, p values more significant than 45 are represented as –log10 of 45.

**Supplemental Figure 2.**
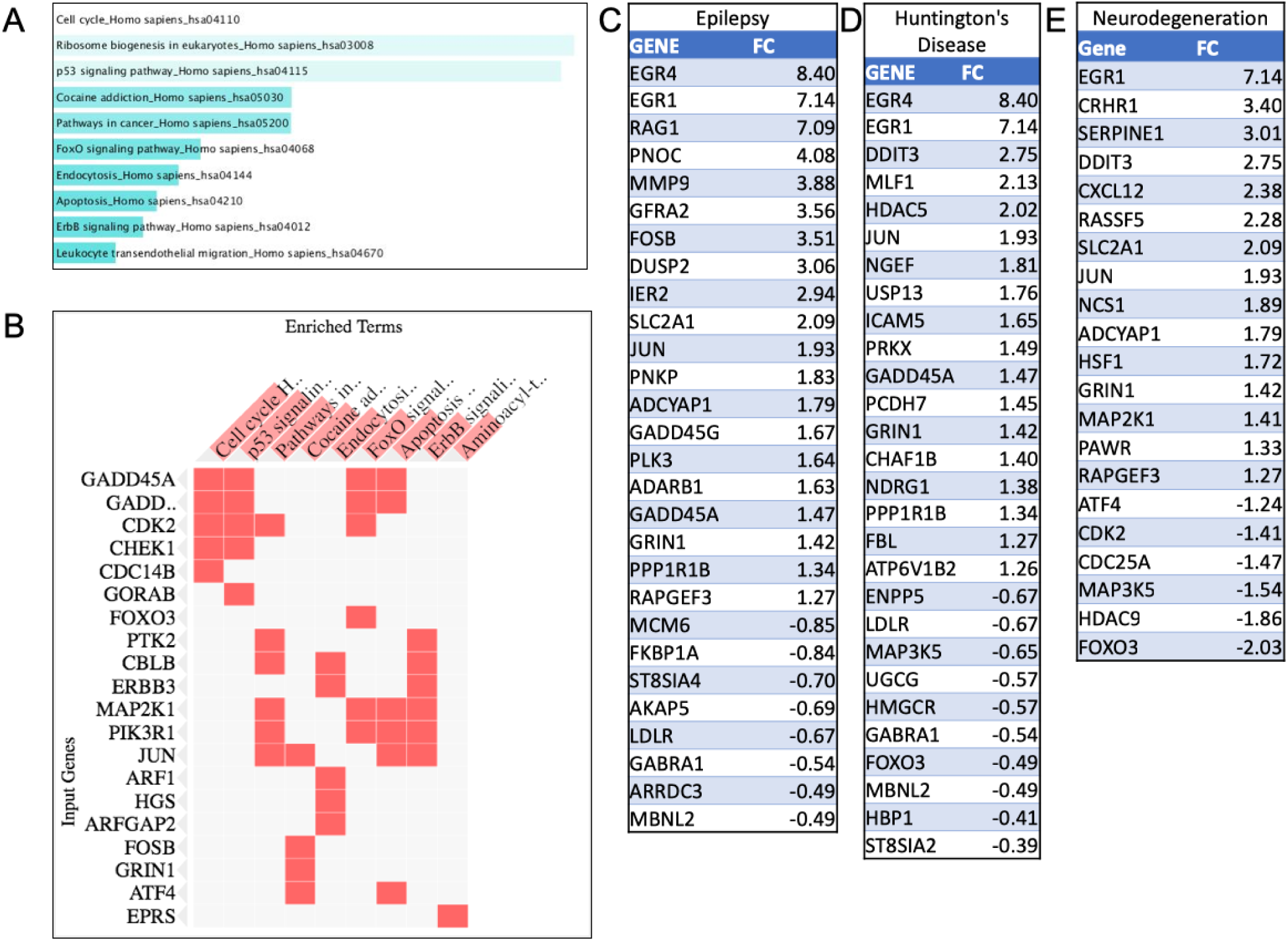
Enrichment Analysis of BDNF regulated genes whose expression is reversed exclusively by 10uM Ruxo. Enrichment analysis was performed on the set of 417 DEGs between RX2+B vs. V+B that were not overlapping with WP+B. (A) KEGG pathway analysis as generated in EnrichR ranked and colored by p value. (B) Top gene clusters involved in the KEGG canonical pathways from A. (C-E) Neurological diseases significantly associated with the RX2 exclusive gene set for (C) Epilepsy, (D) Huntington’s disease, (E) Neurodegeneration.

