## Supplemental Data Tables for "Transcriptomic analysis of the BDNF-induced JAK/STAT pathway in neurons: a window into epilepsy-associated gene expression"

**V+B v CTRL LIST**

| Gene Symbol | log2FC_db_dw | pAdj_db_dw | Aliases | Entrez ID |
|---|---|---|---|---|
| SH2D3C | -3.5186608 | 0 | Nsp3 | 362111 |
| TXNIP | -2.0627642 | 0 | Vdup1 | 117514 |
| KLF11 | -2.0083106 | 0 | Tcfcp2l2 | 313994 |
| CCNG2 | -1.8418735 | 0 | | 29157 |
| HECTD2 | 1.6576949 | 0 | | 309514 |
| BAIAP2 | 1.6914603 | 0 | Irsp53 | 117542 |
| ARHGEF3 | 1.8163428 | 0 | | 290541 |
| TPBG | 1.8295311 | 0 | 5T4\|WAIF1 | 83684 |
| PDP1 | 1.8499978 | 0 | Ppm2c | 54705 |
| THRB | 1.9880408 | 0 | C-erba-beta\|ERBA2\|Nr1a2\|R | 24831 |
| NEFL | 2.0243213 | 0 | NF-L\|Nfl | 83613 |
| SRXN1 | 2.0964212 | 0 | Ab2-390\|Npn3 | 296271 |
| SDC1 | 2.1574545 | 0 | HSPG\|SYNDECA\|Synd1\|Synd | 25216 |
| RGS4 | 2.158761 | 0 | | 29480 |
| BAG3 | 2.1794496 | 0 | | 293524 |
| SPRY4 | 2.1836648 | 0 | | 291610 |
| SMTN | 2.207883 | 0 | | 289734 |
| MAS1 | 2.4278078 | 0 | Mgra\|c-mas | 25153 |
| GFRA1 | 2.4593477 | 0 | | 25454 |
| DUSP6 | 2.5314097 | 0 | Mkp3 | 116663 |
| SCN1B | 2.5327137 | 0 | | 29686 |
| KLF10 | 2.5528164 | 0 | Tieg | 81813 |
| GYPC | 2.6156454 | 0 | GPC | 364837 |
| SMPDL3B | 2.759265 | 0 | | 362619 |
| PDGFB | 2.8178334 | 0 | SIS\|c-sis | 24628 |
| DUSP5 | 2.8205106 | 0 | Cpg21 | 171109 |
| GADD45B | 2.8589942 | 0 | | 299626 |
| SCG2 | 2.8881552 | 0 | Chcg | 24765 |
| LPAR1 | 2.9599757 | 0 | Edg2 | 116744 |
| GRASP | 2.962261 | 0 | | 192254 |
| HAUS8 | 2.999679 | 0 | Ny-sar-48\|RGD1311565 | 290626 |
| EGR1 | 3.0695915 | 0 | Krox-24\|NGFI-A\|Ngf1\|Ngfi\|zi | 24330 |
| ECEL1 | 3.1382005 | 0 | Dine | 60417 |
| LAMP5 | 3.2535648 | 0 | BAD-LAMP\|LAMP-5\|RGD130 | 362220 |
| GPR158 | 3.32403 | 0 | | 291352 |
| RASL11A | 3.382681 | 0 | | 304268 |
| STX11 | 3.3982918 | 0 | | 292483 |
| TRPC6 | 3.5656304 | 0 | Trrp6 | 89823 |
| NPTX2 | 3.5869105 | 0 | NP-II\|NP2\|Narp | 288475 |
| ANKRD33B | 3.6229866 | 0 | RGD1564227 | 310200 |
| FOSB | 3.7941096 | 0 | fra-2 | 100360880 |

| | | | | |
|---|---|---|---|---|
| CYP26B1 | 3.9069061 | 0 | | 312495 |
| EGR2 | 4.0791073 | 0 | Krox20 | 114090 |
| FAM150B | 4.0822096 | 0 | | 679566 |
| VGF | 4.137181 | 0 | | 29461 |
| EMP1 | 4.1743793 | 0 | CL-20\|EMP-1\|ENP1MR\|TMP | 25314 |
| HSF2BP | 4.194514 | 0 | RGD1565879 | 499413 |
| SIM1 | 4.295678 | 0 | | 309888 |
| RRAD | 4.5104423 | 0 | | 83521 |
| PLAUR | 4.754548 | 0 | Par\|Plaur3\|uPAR\|uPAR-2\|uP | 50692 |
| ARC | 5.054995 | 0 | rg3.1 | 54323 |
| GALR1 | 5.1025863 | 0 | Galnr1 | 50577 |
| MMP3 | 5.109894 | 0 | | 171045 |
| MMP13 | 5.2357397 | 0 | | 171052 |
| GPRC5A | 5.4281096 | 0 | Rai3 | 312790 |
| CITED2 | 2.067604 | 1.40E-45 | Mrg1 | 114490 |
| PRSS23 | 2.4334733 | 2.80E-45 | | 308807 |
| CCKBR | 2.7934568 | 2.80E-45 | Cck2r\|Cholrec | 25706 |
| CSRNP1 | 2.6077116 | 3.20E-44 | Axud1 | 363165 |
| GALNT9 | 1.649546 | 7.40E-44 | | 304571 |
| PENK | 3.258971 | 7.60E-44 | Enk\|Penk-rs\|Penk1\|Penk2 | 29237 |
| SPRR1A | 4.405632 | 1.18E-43 | Spr\|Spr1a\|Sprr\|Sprr1\|Sprr1a | 499660 |
| NPTX1 | 2.3235111 | 1.19E-43 | | 266777 |
| SSTR3 | 3.0217788 | 1.51E-43 | SSTR\|Smstr28 | 171044 |
| HOMER1 | 1.6634928 | 5.16E-43 | HOMER1F\|Vesl-1 | 29546 |
| NAB2 | 1.9878603 | 6.96E-43 | | 314910 |
| ALOXE3 | 2.4174693 | 2.52E-42 | e-LOX-3\|eLOX-3 | 287424 |
| SERTAD1 | 2.2239814 | 2.65E-42 | | 361526 |
| GPR68 | 2.5399456 | 2.66E-41 | | 314386 |
| EGR4 | 3.1501546 | 4.21E-41 | Egr4l1\|NGFI-C | 25129 |
| LBH | 1.3661401 | 6.01E-41 | | 683626 |
| DAAM1 | -1.1606426 | 7.91E-41 | | 314212 |
| RT1-S3 | 2.45826 | 5.13E-40 | H2-T23\|RT-BM1\|RT1-BM1\|R | 294228 |
| SSTR4 | 2.0445876 | 6.55E-40 | Smstr4 | 25555 |
| PCDH8 | 1.7111181 | 8.03E-40 | Arcadlin | 64865 |
| PTAFR | 3.4609513 | 1.20E-39 | | 58949 |
| PLCL1 | 1.8863832 | 2.60E-39 | | 84587 |
| DHRS9 | 3.8141215 | 4.01E-38 | Rdhl | 170635 |
| SLCO2A1 | 3.8286128 | 1.77E-37 | Matr1\|Slc21a2 | 24546 |
| KLF9 | 1.8472241 | 4.20E-36 | Bteb\|Bteb1 | 117560 |
| VASP | 1.8119922 | 7.63E-36 | | 361517 |
| ISM1 | 3.1125116 | 8.86E-36 | RGD1562551 | 311760 |
| OSBP2 | 1.3585519 | 9.21E-36 | RGD1309479 | 305475 |
| MEIS2 | 1.2149252 | 1.08E-35 | | 311311 |

| | | | | |
|---|---|---|---|---|
| MYT1L | -1.253483 | 1.69E-35 | Nzf-1\|Nzf1\|Nztf1 | 116668 |
| FAM110C | 3.6874802 | 2.14E-35 | | 500638 |
| MAN1A1 | 2.551768 | 2.79E-35 | Aa2-166\|Man1a | 294410 |
| IRS2 | 1.6023843 | 3.76E-35 | 4PS\|IRS-2 | 29376 |
| SPRY2 | 1.3205673 | 4.32E-35 | | 306141 |
| SPHK1 | 3.0671668 | 6.52E-35 | | 170897 |
| AREG | 3.6564286 | 1.21E-34 | | 29183 |
| RDH10 | 1.7078578 | 1.30E-34 | | 353252 |
| GCNT1 | 3.2552552 | 2.13E-34 | core 2 GlcNAc-T | 64043 |
| MYCL | -2.4887106 | 6.92E-34 | Lmyc1\|Mycl1 | 298506 |
| DUSP4 | 2.4186413 | 1.97E-33 | Mkp-2\|Mkp2 | 60587 |
| INHBA | 2.3958855 | 2.04E-33 | | 29200 |
| ADCY8 | 1.7645378 | 4.52E-33 | Ac8 | 29241 |
| KCNJ4 | 1.2723438 | 5.24E-33 | Hirk2 | 116649 |
| PDXK | 2.2978668 | 1.10E-32 | | 83578 |
| SCN2A | -1.0416238 | 4.14E-32 | NachII\|Nav1.2\|RII/RIIA\|RNSC | 24766 |
| LMNA | 1.6316702 | 4.87E-32 | | 60374 |
| PRIMA1 | 3.1429088 | 9.42E-32 | PRiMA\|Prima1l | 690195 |
| KCNA1 | 2.5377295 | 1.93E-31 | Kcna\|Kcpvd\|Kv1.1 | 24520 |
| IFRD1 | 1.5897458 | 3.00E-31 | Pc4 | 29596 |
| IER5 | 1.8107331 | 4.45E-31 | | 498256 |
| TRIB2 | 1.0827541 | 5.40E-31 | RGD1564451 | 313974 |
| SMAGP | 2.7230294 | 5.71E-31 | | 300236 |
| DRD5 | 3.8985677 | 7.07E-31 | D1B | 25195 |
| IL6R | 2.301657 | 1.60E-30 | IL6R1\|Il6ra | 24499 |
| FAM84A | 1.5672331 | 1.86E-30 | RGD1305779 | 313969 |
| INSC | 3.1004944 | 1.88E-30 | RGD1311379 | 293166 |
| CYP27B1 | 2.0879796 | 3.07E-30 | Cyp40 | 114700 |
| GAL | 3.638903 | 4.39E-30 | Galn | 29141 |
| ITGA6 | 1.8318036 | 4.59E-30 | | 114517 |
| PIK3IP1 | -1.2761761 | 5.60E-30 | RGD1311203\|zgc:66482 | 305472 |
| NPL | 1.8066884 | 9.61E-30 | | 304860 |
| HS3ST5 | 1.6394793 | 1.43E-29 | | 294449 |
| ACAN | 2.3603435 | 1.89E-29 | Agc\|Agc1 | 58968 |
| FST | -2.2953715 | 3.23E-29 | FOL1\|FS\|Fst-288\|RATFOL1 | 24373 |
| FRMD6 | 1.591484 | 3.23E-29 | | 257646 |
| CRHBP | 2.8745425 | 1.69E-28 | Crfbp | 29625 |
| GEM | 2.114901 | 2.35E-28 | | 297902 |
| FOS | 1.8276407 | 3.46E-28 | c-fos | 314322 |
| FGFR1 | 1.739113 | 1.00E-27 | | 79114 |
| NRN1 | 2.0853286 | 1.11E-27 | Nrn | 83834 |
| EHD4 | 1.4695604 | 1.22E-27 | Past2 | 192204 |
| C1QL3 | 1.7291608 | 1.40E-27 | | 680404 |

| | | | | |
|---|---|---|---|---|
| PRICKLE1 | 1.6093031 | 2.63E-27 | Rilp | 315259 |
| MTUS2 | -1.7557187 | 1.02E-26 | RGD1562533 | 498136 |
| ADORA1 | 1.7060324 | 1.44E-26 | | 29290 |
| C1QL2 | 3.2518566 | 1.46E-26 | | 288979 |
| LEXM | 2.7456307 | 1.92E-26 | RGD1559493 | 500516 |
| BHLHE22 | -2.3506196 | 2.87E-26 | Bhlhb5 | 365748 |
| RGS16 | 1.710074 | 3.55E-26 | | 360857 |
| CXXC5 | 1.1755618 | 4.92E-26 | | 291670 |
| ARID5A | 1.7327574 | 7.22E-26 | | 316327 |
| PTPRR | -1.9995065 | 1.02E-25 | PC12-PTP1\|Pcptp1\|R-PTP-R | 94202 |
| ADRA1D | 2.8663747 | 1.21E-25 | Adrd1 | 29413 |
| SEMA3A | 2.0394833 | 1.24E-25 | | 29751 |
| DUSP14 | 1.8628569 | 1.29E-25 | Dusp14l2 | 360580 |
| BHLHE40 | 1.5580747 | 1.70E-25 | Bhlhb2\|Dec1\|SHARP-2\|Sharp | 79431 |
| TBX21 | 3.676611 | 2.03E-25 | | 303496 |
| SP7 | 3.4122097 | 4.62E-25 | Osx | 300260 |
| TMEM150C | -1.1294942 | 5.69E-25 | RGD1306105 | 360916 |
| PLCXD2 | -1.1535157 | 7.99E-25 | RGD1563504 | 363781 |
| HTR1B | 2.5193267 | 8.85E-25 | | 25075 |
| CCND1 | 1.4953775 | 1.45E-24 | | 58919 |
| PTGS2 | 2.4685276 | 2.83E-24 | COX-2\|Cox2 | 29527 |
| TMEM65 | 1.164343 | 6.48E-24 | RGD1563224 | 500874 |
| RASSF8 | 2.6404998 | 6.48E-24 | RGD1308820 | 312846 |
| MTMR12 | 1.8788508 | 1.05E-23 | Pip3ap | 310155 |
| CNGA4 | 3.2149022 | 1.55E-23 | Cgn2\|rOCNC2 | 85258 |
| RTN4RL1 | 1.703219 | 3.08E-23 | Ngrh2 | 303311 |
| MMP10 | 3.6014867 | 3.49E-23 | | 117061 |
| MMP9 | 3.1567497 | 4.16E-23 | | 81687 |
| FAM102B | 1.0435205 | 5.39E-23 | RGD1310037 | 365903 |
| RNF217 | 1.0695094 | 7.63E-23 | Ibrdc1 | 292188 |
| ZYX | 1.2675476 | 1.46E-22 | | 114636 |
| TRIB1 | 1.8991559 | 2.54E-22 | Gig2 | 78969 |
| GPAT3 | 1.9999765 | 2.57E-22 | Agpat9 | 305166 |
| MYO1E | 1.7415842 | 3.58E-22 | MYR5\|Myr3 | 25484 |
| SERPINB2 | 3.5841072 | 5.20E-22 | Pai2a | 60325 |
| DBNDD1 | -1.8708739 | 1.08E-21 | RGD1310008 | 361437 |
| B3GNT7 | -1.2929933 | 1.67E-21 | | 316583 |
| SORBS2 | 1.1322527 | 1.91E-21 | Argbp2 | 114901 |
| DOK5 | 1.6272763 | 3.00E-21 | RGD1562846 | 502694 |
| HSPH1 | 1.0666467 | 4.02E-21 | Hsp105 | 288444 |
| ATG16L1 | 1.066054 | 5.70E-21 | Apg16l\|Wdr30 | 363278 |
| DLGAP2 | 2.4980152 | 6.56E-21 | DAP-2\|SAPAP2 | 116681 |
| DCLK3 | -2.5603716 | 7.79E-21 | Dcamkl3\|RGD1309232 | 316023 |

| | | | | |
|---|---|---|---|---|
| RUNX2 | 2.8620102 | 2.15E-20 | Cbfa1\|OSF-2 | 367218 |
| HMGA1 | 1.9424802 | 2.52E-20 | Hmgi\|Hmgiy | 117062 |
| ZC2HC1A | -1.018401 | 2.60E-20 | Fam164a\|RGD1311970 | 310244 |
| CAV1 | 1.5572658 | 2.61E-20 | Cav | 25404 |
| EFR3A | 1.0692867 | 2.68E-20 | RGD1305976 | 362923 |
| RGS20 | 1.2998077 | 3.26E-20 | | 362477 |
| PTH2R | 2.567532 | 4.37E-20 | Pthr2 | 81753 |
| KCNV1 | 1.3596368 | 4.89E-20 | Kv8.1 | 60326 |
| GPR83 | 2.9704585 | 8.92E-20 | Gir | 140595 |
| ERRFI1 | 1.1447444 | 9.01E-20 | RGD1307599\|Ralt\|Xxx\|gene | 313729 |
| IER5L | 1.5424821 | 1.11E-19 | | 499772 |
| NEDD9 | 1.1884135 | 1.48E-19 | | 291044 |
| WDR43 | 0.78759044 | 1.99E-19 | | 362703 |
| BAALC | 1.0558892 | 2.48E-19 | | 140720 |
| DKK2 | 3.33558 | 3.13E-19 | Dkk4 | 295445 |
| MAP2K6 | -1.2294734 | 3.97E-19 | Mkk6 | 114495 |
| AP1S3 | 1.6560899 | 4.60E-19 | | 367304 |
| PLK2 | 1.2168903 | 4.68E-19 | Snk | 83722 |
| FAM212B | 1.5589373 | 5.27E-19 | RGD1306526 | 310764 |
| FAM196A | 1.2841877 | 5.33E-19 | | 100233213 |
| DNAJB5 | 1.018804 | 6.05E-19 | | 313811 |
| SELE | 2.8674822 | 6.41E-19 | | 25544 |
| 42805 | 1.4264649 | 6.69E-19 | MARCH-XI\|RGD1559945 | 499558 |
| VIPR1 | 1.814615 | 9.38E-19 | PACAP-R-2\|PACAP-R2\|RATVA | 24875 |
| PTPRU | 0.9018883 | 9.45E-19 | | 116680 |
| GCGR | 2.2198486 | 9.56E-19 | | 24953 |
| BAZ1A | 1.8905354 | 1.21E-18 | | 314126 |
| KITLG | 1.0291016 | 1.52E-18 | Kitl\|Mgf\|SCF | 60427 |
| POU4F3 | 2.8643954 | 3.79E-18 | | 364855 |
| SS18 | 0.89652956 | 4.13E-18 | | 361295 |
| ARHGAP10 | 1.32641 | 4.25E-18 | | 688429 |
| RGD1307461 | -1.6390955 | 6.60E-18 | | 300990 |
| SYT10 | 2.6374457 | 7.16E-18 | | 60567 |
| ATP2B1 | 0.7975083 | 7.42E-18 | Pmca1a\|Pmca1b\|Pmca1c | 29598 |
| EGR3 | 2.477239 | 1.16E-17 | | 25148 |
| TMOD1 | 1.247792 | 1.25E-17 | E-Tmod\|Tmod | 25566 |
| BDNF | 1.6924942 | 1.46E-17 | | 24225 |
| CDKN1A | 1.5740019 | 1.68E-17 | Cip1\|UV96\|Waf1 | 114851 |
| PCSK1 | 1.3431313 | 1.73E-17 | BDP\|PC1\|PC3 | 25204 |
| AJAP1 | 1.2560861 | 2.15E-17 | neumo48 | 687031 |
| PPP1R18 | -0.9783594 | 2.15E-17 | Kiaa1949\|RGD1309543 | 361790 |
| SHC4 | 2.0069253 | 4.60E-17 | RGD1309182 | 679845 |
| KLHDC8A | -1.9683065 | 5.18E-17 | RGD1305132 | 305096 |

| | | | | |
|---|---|---|---|---|
| CARS | 0.8850533 | 6.37E-17 | | 293638 |
| PDE9A | 1.5808959 | 9.09E-17 | | 191569 |
| VEGFA | 1.445269 | 9.35E-17 | VEGF-A\|VEGF164\|VPF\|Vegf | 83785 |
| CSPG4 | 1.0248827 | 1.00E-16 | Ng2 | 81651 |
| HACE1 | 1.0541575 | 1.18E-16 | | 361866 |
| ELOVL7 | 2.0444527 | 1.19E-16 | | 361895 |
| DNAJB4 | -0.92410535 | 1.29E-16 | | 295549 |
| ADRA1B | 1.8079491 | 1.36E-16 | | 24173 |
| RBMS1 | 1.556421 | 2.05E-16 | | 362138 |
| MYBPH | 1.4676771 | 3.00E-16 | | 83708 |
| TRIM9 | 1.334747 | 4.48E-16 | | 155812 |
| PIK3CD | -1.2090521 | 4.90E-16 | | 366508 |
| RGD1563365 | 1.0237044 | 4.90E-16 | | 299700 |
| VCL | 1.0615209 | 7.73E-16 | | 305679 |
| PNMA2 | -0.8894666 | 8.17E-16 | | 305977 |
| WDR1 | 0.78475046 | 8.63E-16 | | 360950 |
| PLEKHH2 | 1.9632562 | 8.70E-16 | RGD1304935 | 313866 |
| CCPG1 | -0.99297786 | 9.71E-16 | | 363098 |
| CREM | 1.5828385 | 1.21E-15 | Icer | 25620 |
| PRAGMIN | 1.5849367 | 1.32E-15 | RGD1311793 | 306506 |
| SIK1 | 1.9205624 | 1.63E-15 | Sik\|Snf1lk | 59329 |
| KCNK2 | -1.6440685 | 1.99E-15 | Trek-1\|rTREK1d | 170899 |
| MAP2K3 | 1.8152273 | 2.12E-15 | Mek3\|Mkk3 | 303200 |
| UBASH3B | 1.6640985 | 2.29E-15 | RGD1310357\|Sts-1 | 315579 |
| SIAH3 | 2.1123347 | 3.02E-15 | | 692004 |
| FAM184B | -1.8618681 | 3.59E-15 | RGD1564950 | 289671 |
| RBM12 | 1.0843652 | 3.59E-15 | | 652928 |
| PVR | 1.6094185 | 3.95E-15 | Taa1\|Tage4 | 25066 |
| ADCK3 | -1.4241498 | 4.09E-15 | Cabc1 | 360887 |
| HSPA4L | 1.0953513 | 4.81E-15 | APG-1\|OSP94 | 294993 |
| NGF | 2.332491 | 5.61E-15 | Ngfb\|beta-NGF | 310738 |
| AGTR1A | 1.9648544 | 5.98E-15 | AT1\|AT1A\|AT1R\|Agtr1 | 24180 |
| SLC6A17 | 1.475906 | 6.02E-15 | Ntt4 | 613226 |
| EIF2AK3 | 0.8770304 | 6.95E-15 | PEK | 29702 |
| CDC37L1 | 0.7691417 | 7.22E-15 | Cdc37l | 293886 |
| TIMP1 | 1.1849104 | 7.35E-15 | TIMP-1\|Timp | 116510 |
| PANX1 | 1.0717888 | 8.85E-15 | px1 | 315435 |
| FAM43A | 1.5121964 | 9.13E-15 | RGD1304790 | 288031 |
| ACVR1 | 0.99055094 | 1.10E-14 | | 79558 |
| DDHD1 | 0.7036751 | 1.13E-14 | | 305816 |
| FGF2 | 2.3530617 | 1.22E-14 | Fgf-2\|bFGF | 54250 |
| GREM1 | 1.2579631 | 1.38E-14 | Cktsf1b1\|drm | 50566 |
| ZMIZ1 | 1.3394367 | 1.43E-14 | Rai17 | 361103 |

| | | | | |
|---|---|---|---|---|
| FGF9 | 1.2275751 | 1.46E-14 | | 25444 |
| KLF14 | 2.2850194 | 1.77E-14 | | 312203 |
| RGD1305014 | 0.9552032 | 2.23E-14 | | 309029 |
| RTN4RL2 | 1.2922189 | 2.29E-14 | Ngrh1 | 311169 |
| ETV5 | 1.6104954 | 2.42E-14 | | 303828 |
| ST7 | 0.9344929 | 2.79E-14 | | 296911 |
| ACTN1 | 1.5171064 | 2.86E-14 | | 81634 |
| KLHDC8B | -0.99473333 | 3.35E-14 | | 306589 |
| GABRG2 | -1.2529383 | 4.04E-14 | | 29709 |
| HIC1 | 2.4993727 | 4.96E-14 | | 303310 |
| ERCC1 | 1.079485 | 4.97E-14 | | 292673 |
| MUSK | 2.7192237 | 5.00E-14 | Nsk1 | 81725 |
| SYPL2 | 2.1900775 | 5.78E-14 | Mg29 | 362018 |
| BAI1 | 1.1931818 | 6.54E-14 | Bai1 | 362931 |
| YPEL5 | -0.77750516 | 7.20E-14 | | 298792 |
| JUN | 0.8752457 | 7.57E-14 | | 24516 |
| RRAGD | -1.0506474 | 8.18E-14 | | 297960 |
| PTPRN | 1.0244571 | 8.32E-14 | ICA512\|Ia-2 | 116660 |
| LRRC26 | 2.4362385 | 9.51E-14 | RGD1308398 | 311803 |
| ACSL4 | 0.82943016 | 1.20E-13 | Acs4\|Facl4 | 113976 |
| SCRT2 | 1.8648645 | 1.30E-13 | RGD1564796 | 366229 |
| RPS6KA5 | -0.8596208 | 1.54E-13 | Msk1 | 314384 |
| NARF | -0.8929865 | 1.71E-13 | RGD1310894 | 360681 |
| ANO4 | 1.349041 | 1.84E-13 | Tmem16d | 299714 |
| RAI14 | 0.94001484 | 1.88E-13 | | 294804 |
| TRMT6 | 0.92291427 | 2.07E-13 | RGD1308877 | 311441 |
| MYH4 | 2.5091531 | 2.37E-13 | | 360543 |
| PPHLN1 | 0.81124485 | 2.77E-13 | | 366975 |
| UOX | 2.5953753 | 3.03E-13 | UOX-2\|Uri\|Uri2 | 114768 |
| MTSS1 | -0.67216885 | 3.26E-13 | | 362918 |
| CACNG3 | 1.1780035 | 3.64E-13 | | 140724 |
| FAM46A | 2.0539212 | 3.71E-13 | RGD1311381 | 300870 |
| HRH3 | -1.9751954 | 4.26E-13 | | 85268 |
| FKBP5 | 0.8295817 | 4.26E-13 | | 361810 |
| SYNJ2 | 1.9054892 | 4.92E-13 | | 84018 |
| DLC1 | 2.12959 | 6.61E-13 | Arhgap7\|RhoGAP | 58834 |
| EHBP1L1 | -1.0739992 | 7.41E-13 | | 309169 |
| TMTC4 | -1.059587 | 8.35E-13 | RGD1560183 | 290501 |
| SSTR2 | 0.9105852 | 9.80E-13 | Smstr2 | 54305 |
| UNG | 0.9744515 | 1.04E-12 | | 304577 |
| GPR37 | -1.4318073 | 1.08E-12 | Ednrbl | 117549 |
| STAM | 0.6097905 | 1.26E-12 | RGD1564499 | 498798 |
| JUNB | 1.1255476 | 1.26E-12 | | 24517 |

| | | | | |
|---|---|---|---|---|
| NUB1 | -0.65661305 | 1.29E-12 | BS4\|NYREN18 | 296731 |
| IGSF9 | -1.2876827 | 1.34E-12 | | 304982 |
| CALCA | 2.0663934 | 1.53E-12 | CAL6\|CGRP\|Cal1\|Calc\|RATCA | 24241 |
| TTPAL | 1.0955765 | 1.56E-12 | RGD1305754 | 296349 |
| SIK3 | 1.1485419 | 1.63E-12 | | 684112 |
| NEFM | 1.3457621 | 1.64E-12 | Nef3\|Nfm | 24588 |
| DMP1 | 2.437485 | 2.08E-12 | DENTMAT | 25312 |
| PNMAL1 | -1.2880445 | 2.17E-12 | RGD1310803 | 361515 |
| LYVE1 | 2.454409 | 2.17E-12 | Xlkd1 | 293186 |
| MRM1 | 0.79004884 | 2.17E-12 | RGD1566232 | 363661 |
| LYRM9 | -1.2623076 | 3.30E-12 | RGD1562012 | 497962 |
| CDC42EP4 | -0.9025045 | 3.30E-12 | | 303653 |
| RNF152 | -1.9679434 | 3.43E-12 | | 293561 |
| SLCO3A1 | 1.2057765 | 3.58E-12 | Slc21a11 | 140915 |
| MICAL2 | 0.9951284 | 4.08E-12 | MICAL-2\|RGD1311773 | 365352 |
| EPHA4 | -0.9142841 | 4.79E-12 | RGD1560587 | 316539 |
| HDAC11 | -0.90699375 | 4.79E-12 | | 297453 |
| RGD1305587 | -1.0728447 | 6.45E-12 | | 294499 |
| DPF3 | 1.2705431 | 6.45E-12 | | 299186 |
| SPRED3 | 1.4192659 | 6.57E-12 | RGD1311713 | 308478 |
| SYNPO | 1.3086188 | 6.83E-12 | | 60324 |
| ZFP143 | 1.0058349 | 6.95E-12 | STAF\|Znf143 | 361627 |
| KCNF1 | 1.4807354 | 6.95E-12 | Kh1\|Kv5.1 | 298908 |
| HTR1F | 2.439059 | 7.05E-12 | Htr1eb | 60448 |
| SESN1 | -0.86736435 | 7.19E-12 | | 294518 |
| EZR | 1.0318053 | 7.50E-12 | Vil2 | 54319 |
| ARVCF | -0.6820622 | 9.00E-12 | | 303798 |
| EFNA4 | -1.4177372 | 1.06E-11 | | 310643 |
| P4HB | 0.5871119 | 1.10E-11 | PDI\|PDIR | 25506 |
| IFFO2 | 1.2169538 | 1.11E-11 | | 641315 |
| VSTM2B | 1.2963742 | 1.25E-11 | RGD1306235 | 361560 |
| KCNC4 | 1.3986031 | 1.34E-11 | | 684516 |
| RPS6KA2 | -1.0282438 | 1.41E-11 | | 117269 |
| RGD1561849 | 1.280649 | 1.50E-11 | | 500393 |
| VWA1 | -1.4944415 | 1.56E-11 | RGD1311476 | 298683 |
| MCM3 | 0.88786876 | 1.83E-11 | | 316273 |
| RGD1304810 | 1.324045 | 1.96E-11 | | 306504 |
| WSCD1 | 0.8512739 | 2.08E-11 | RGD1308212 | 287466 |
| TSPAN5 | 0.97393215 | 2.12E-11 | Tm4sf9\|Tspan-5 | 362048 |
| MYC | 0.7217986 | 2.57E-11 | RNCMYC\|c-myc\|mMyc | 24577 |
| FLT1 | 2.370946 | 2.83E-11 | FLT-1\|VEGFR-1 | 54251 |
| NPY1R | -0.9346991 | 2.91E-11 | NPY-1 | 29358 |
| CLMP | -1.2245033 | 3.02E-11 | ACAM\|Asam\|Ol16 | 286939 |

| | | | | |
|---|---|---|---|---|
| GPR12 | -1.6918236 | 3.45E-11 | Gpcr12 | 80840 |
| AK1 | -0.9593049 | 3.91E-11 | Ak 1 | 24183 |
| SEMA6C | -1.6502132 | 3.95E-11 | | 29744 |
| CXCL12 | 1.1677482 | 4.43E-11 | Sdf1 | 24772 |
| HECA | 0.6286029 | 4.48E-11 | | 308624 |
| SLC22A3 | -1.5044653 | 4.74E-11 | | 29504 |
| EDNRA | 1.3215874 | 4.97E-11 | ET-A\|ET-AR\|Endor\|Eta\|RGD1 | 24326 |
| PCED1B | -1.3029708 | 5.22E-11 | Fam113b\|RGD1561028 | 315283 |
| GPR137C | -1.3275257 | 5.98E-11 | RGD1563070 | 305812 |
| PIK3C3 | -0.86102486 | 6.10E-11 | | 65052 |
| FILIP1 | -1.8423249 | 6.32E-11 | Filip | 246776 |
| IPPK | 1.1657133 | 6.91E-11 | C9orf12\|RGD1311271\|rIpk1 | 306808 |
| SLC9A2 | 1.4313039 | 7.30E-11 | Nhe2 | 24783 |
| HSH2D | 2.2285504 | 7.33E-11 | | 100360518 |
| ABHD13 | 0.65878487 | 7.49E-11 | RGD1308317 | 306630 |
| KLHL29 | 0.87309074 | 7.65E-11 | Kbtbd9 | 298867 |
| ARMCX6 | -1.0612724 | 9.60E-11 | | 363496 |
| CYLD | -0.87643355 | 9.95E-11 | LRRGT00003\|Rp1\|Rp1h | 312937 |
| STAC2 | 1.1889721 | 9.95E-11 | | 363674 |
| DIXDC1 | -0.8126806 | 1.38E-10 | Ccd1 | 363062 |
| GUCY1B3 | -0.68394923 | 1.40E-10 | Gucy1b1\|SGC | 25202 |
| NGEF | 0.7739159 | 1.50E-10 | Besh3 | 246217 |
| SLC4A11 | 1.8949499 | 1.59E-10 | | 311423 |
| LOC680663 | 1.8478743 | 1.69E-10 | | 680663 |
| RUNX1 | 2.1376946 | 1.70E-10 | Aml1\|B\|CBF-alpha-2\|Cbfa2\|I | 50662 |
| ZFP238 | -1.0958638 | 1.71E-10 | Rp58\|Zbtb18\|Znf238 | 64619 |
| SLC43A2 | -0.6407087 | 1.72E-10 | RGD1305819 | 287532 |
| BCAR3 | 1.0566084 | 1.86E-10 | | 310838 |
| RND1 | -1.0471507 | 1.98E-10 | | 362993 |
| PHEX | 1.9048415 | 2.01E-10 | PEX | 25512 |
| UNC5B | 1.3878748 | 2.11E-10 | Unc5h2 | 60630 |
| GRWD1 | 0.9444853 | 2.14E-10 | | 308592 |
| LHFPL2 | 0.89294094 | 2.19E-10 | | 294643 |
| CDK17 | 0.75188833 | 2.35E-10 | Pctaire2\|Pctk2 | 314743 |
| MCHR1 | 1.4962466 | 2.37E-10 | Gpr24\|Mch-1r\|Slc1 | 83567 |
| CBLN2 | 1.2851559 | 2.71E-10 | | 291388 |
| UAP1 | 0.9331793 | 2.85E-10 | RGD1561967 | 498272 |
| HES6 | -0.73470825 | 3.25E-10 | | 316626 |
| SDCBP | 0.6194749 | 3.28E-10 | TACIP18\|mda-9 | 83841 |
| CYGB | -1.2168483 | 3.42E-10 | Staap\|Stap | 170520 |
| ME1 | 0.57811105 | 3.42E-10 | MOD1 | 24552 |
| TMEM246 | -0.8830453 | 3.44E-10 | RGD1307218 | 362518 |
| ELMOD1 | -0.77389467 | 3.91E-10 | RGD1564186 | 315670 |

| | | | | |
|---|---|---|---|---|
| EML4 | 0.7334703 | 4.16E-10 | | 313861 |
| NRF1 | 0.98583704 | 5.41E-10 | Nfe2l1_retired | 312195 |
| RBM24 | -1.4161272 | 5.43E-10 | | 690139 |
| ZBTB46 | 1.3315436 | 5.43E-10 | Btbd4 | 311718 |
| POU4F2 | 2.4603977 | 5.44E-10 | Brn3b | 171355 |
| SMAD3 | 1.0452014 | 6.06E-10 | Madh3\|Smad 3\|mad3 | 25631 |
| CREBL2 | -0.84521633 | 7.53E-10 | | 362453 |
| TACR3 | 1.8273549 | 7.79E-10 | Nkr\|Nmkr\|Tac3r | 24808 |
| LIMK2 | -0.8737091 | 8.14E-10 | Limk2b\|Link2 | 29524 |
| RAG1 | 2.31254 | 8.26E-10 | | 84600 |
| NDUFAF4 | 1.0272276 | 8.27E-10 | Hrpap20 | 362495 |
| KCNN1 | -1.3946551 | 9.01E-10 | KCa2.1 | 54261 |
| KRAS | 1.1000634 | 9.01E-10 | Kras2\|c-Ki-ras\|p21 | 24525 |
| FMNL1 | 1.1394138 | 9.39E-10 | | 287746 |
| MAL2 | -0.8264061 | 1.04E-09 | | 362911 |
| PRSS12 | -0.833376 | 1.55E-09 | Nt | 85266 |
| HBEGF | 1.0555133 | 1.62E-09 | Dtr\|GFHB\|Hb-egf\|Hegfl | 25433 |
| RASSF7 | -1.6510895 | 1.65E-09 | RGD1306244 | 293623 |
| SLC25A37 | 1.4660352 | 1.76E-09 | Mscp\|RGD1359361 | 306000 |
| LRRC61 | -0.8664268 | 1.81E-09 | RGD1561800 | 500111 |
| SLITRK4 | -0.9375122 | 2.01E-09 | | 302473 |
| RAB3D | -1.1076238 | 2.07E-09 | | 140665 |
| PSME3 | 0.65283084 | 2.24E-09 | Ab2-371 | 287716 |
| GALNT7 | 1.2785124 | 2.26E-09 | | 29750 |
| LRRC71 | -0.78167814 | 2.38E-09 | RGD1309453 | 310689 |
| CCDC92 | 0.75157523 | 2.62E-09 | | 100036765 |
| ZC4H2 | -0.76520795 | 3.37E-09 | RGD1561708 | 367838 |
| DHX57 | -0.58144784 | 3.59E-09 | | 366532 |
| MCM2 | 0.8999621 | 3.80E-09 | | 312538 |
| SLC41A2 | 1.037418 | 3.80E-09 | | 362861 |
| AOC3 | 1.9076623 | 3.96E-09 | SSAO | 29473 |
| SLC2A3 | 1.2184542 | 4.09E-09 | GLUT3 | 25551 |
| NRIP1 | -0.7432924 | 4.11E-09 | RIP140 | 304157 |
| CLDND1 | 0.586019 | 4.17E-09 | | 288182 |
| BLES03 | -0.8337398 | 4.38E-09 | | 266609 |
| RASGEF1C | -0.9259993 | 4.41E-09 | | 360519 |
| TAGAP | 1.5805598 | 4.44E-09 | | 308097 |
| PRKAR2A | 1.4116378 | 4.84E-09 | | 29699 |
| SPNS2 | 1.150435 | 4.93E-09 | | 100270678 |
| MCM6 | 0.58436936 | 5.17E-09 | Mcmd6 | 29685 |
| TMEM218 | -0.89298445 | 5.23E-09 | RGD1311364 | 300516 |
| ADCY7 | 1.4964327 | 5.23E-09 | | 84420 |
| CFLAR | 1.031613 | 5.25E-09 | Flip | 117279 |

| | | | | |
|---|---|---|---|---|
| ATP1B1 | 0.7771755 | 5.34E-09 | ATPBS | 25650 |
| CCDC86 | 0.79873097 | 5.44E-09 | | 293738 |
| CHN2 | -0.97858006 | 5.54E-09 | | 84031 |
| DOK4 | -0.7422019 | 5.64E-09 | | 361364 |
| SAMD4A | 1.6091677 | 5.70E-09 | Samd4 | 305826 |
| ZFPM1 | 1.1657131 | 6.00E-09 | | 691504 |
| MAN2A1 | 1.3874837 | 6.05E-09 | Mana2\|Manno | 25478 |
| NETO1 | 0.9790565 | 6.11E-09 | RGD1566269 | 307206 |
| SLC4A7 | 1.6119723 | 6.16E-09 | NBC3\|NBCn1 | 117955 |
| ORAI2 | -1.1369072 | 6.24E-09 | RGD1310213 | 304592 |
| LRRN3 | -1.0593017 | 6.28E-09 | Nlrr3 | 81514 |
| FAM89A | 1.0497959 | 6.56E-09 | RGD1309879 | 361441 |
| LOC688613 | -1.1505996 | 7.25E-09 | | 688613 |
| LRRTM2 | -0.8271959 | 7.34E-09 | | 685472 |
| GUCY1A3 | -1.0689547 | 7.49E-09 | Gucy1a1\|SGC | 497757 |
| NCBP1 | 0.5205126 | 7.53E-09 | | 298075 |
| RASGRP2 | -0.6573872 | 7.58E-09 | | 361714 |
| RXRG | -1.9081825 | 7.88E-09 | | 83574 |
| FAR1 | 1.0130634 | 7.96E-09 | Mlstd2 | 293173 |
| RGD1563072 | -1.1282976 | 8.27E-09 | | 313595 |
| TMEM163 | 1.1404581 | 8.67E-09 | RGD1306212\|Sv31 | 360839 |
| BMPER | 1.1796024 | 9.18E-09 | RGD1563373 | 300455 |
| PER2 | 0.7842451 | 9.52E-09 | rPER2 | 63840 |
| KLC4 | -0.75756204 | 1.23E-08 | 1200014p03rik\|Knsl8 | 316226 |
| ANXA11 | 0.9251612 | 1.25E-08 | | 290527 |
| SF1 | 0.58964086 | 1.27E-08 | Zfp162 | 117855 |
| NR1D2 | -0.97861964 | 1.31E-08 | HZF-2 | 259241 |
| KCNN3 | 1.3866298 | 1.32E-08 | KCa2.3\|SK3 | 54263 |
| ARRDC1 | -0.8560828 | 1.37E-08 | | 366001 |
| DDX21 | 0.96944666 | 1.42E-08 | Ddx21a\|Ddx21b | 317399 |
| TM4SF1 | 1.8135058 | 1.55E-08 | | 295061 |
| STK26 | 2.1175604 | 1.66E-08 | Mask\|Mst4\|RGD1563568 | 317589 |
| CPNE8 | -0.8491475 | 1.71E-08 | | 362988 |
| CRHR1 | 1.1218804 | 1.73E-08 | CRFR1 | 58959 |
| KLF7 | -1.2560346 | 1.84E-08 | | 363243 |
| GADD45G | 1.1591874 | 1.88E-08 | | 291005 |
| DLK1 | 1.3777293 | 1.91E-08 | Pref-1\|Zog | 114587 |
| TMEM200C | -0.834058 | 1.91E-08 | RGD1561734 | 501201 |
| SHROOM2 | -1.0640309 | 1.97E-08 | Ab2-404\|Apxl | 317435 |
| PAM | 1.0347998 | 1.97E-08 | | 25508 |
| CCDC129 | 1.7321779 | 1.97E-08 | RGD1561131 | 500139 |
| PRRT4 | -1.488174 | 2.00E-08 | RGD1559885 | 500059 |
| MAP2K4 | 0.80338305 | 2.04E-08 | MKK4\|Sek1 | 287398 |

| | | | | |
|---|---|---|---|---|
| POPDC2 | -1.1629769 | 2.05E-08 | | 360718 |
| CAPN5 | -0.91000926 | 2.26E-08 | Htra3 | 171495 |
| COL4A1 | 1.1251773 | 2.29E-08 | | 290905 |
| HMGCLL1 | -1.0695127 | 2.33E-08 | RGD1565090 | 367112 |
| CTSF | -0.81760997 | 2.34E-08 | | 361704 |
| KLHL24 | -1.0576439 | 2.43E-08 | Dre1 | 303803 |
| NFIB | -0.60702807 | 2.45E-08 | | 29227 |
| KLHL5 | -0.5935278 | 2.49E-08 | | 305351 |
| RGS17 | 0.872459 | 2.49E-08 | | 308118 |
| RGD1564036 | -0.82035017 | 2.69E-08 | | 497895 |
| RGD1561931 | -0.6385058 | 2.73E-08 | KIAA2022 | 302396 |
| KLF5 | 0.81559616 | 2.88E-08 | Bteb2\|IKLF\|bteb2 | 84410 |
| PDSS2 | -0.96159464 | 2.90E-08 | | 365592 |
| M6PR | 0.597658 | 2.90E-08 | | 312689 |
| SLC16A14 | 1.4101831 | 2.90E-08 | | 316578 |
| DENND2C | 1.1995537 | 2.96E-08 | RGD1308197 | 295333 |
| GOLPH3 | 0.58364147 | 2.96E-08 | Gmx33 | 78961 |
| MYH1 | 2.0129728 | 3.14E-08 | MYHC | 287408 |
| IGSF3 | -0.91093475 | 3.42E-08 | | 295325 |
| ALDH1B1 | -0.90188223 | 3.43E-08 | | 298079 |
| RAMP3 | -1.5966079 | 3.50E-08 | | 56820 |
| BSDC1 | -0.55472773 | 3.50E-08 | RGD1311622 | 297890 |
| DUSP2 | 1.0111473 | 3.79E-08 | | 311406 |
| EHBP1 | -0.5281663 | 4.14E-08 | RGD1310604 | 305556 |
| LZTS1 | -0.97071993 | 4.18E-08 | Psdzip70 | 266711 |
| ZFP775 | -0.81265396 | 4.21E-08 | RGD1304910\|Znf775 | 312309 |
| LRRTM3 | 0.59607244 | 4.21E-08 | | 294380 |
| RET | 1.0616585 | 4.21E-08 | | 24716 |
| BVES | 1.2946421 | 4.30E-08 | Popdc1 | 365603 |
| SOCS3 | 1.5331312 | 4.83E-08 | Cish3\|Socs-3\|Ssi-3 | 89829 |
| WNT6 | 1.939307 | 4.89E-08 | | 316526 |
| SPHKAP | -1.3342793 | 5.50E-08 | RGD1311951\|Skip | 316561 |
| NAMPT | 0.7191915 | 5.65E-08 | Pbef\|Pbef1 | 297508 |
| NIPSNAP3B | -0.71185213 | 5.84E-08 | Nipsnap3a | 313211 |
| ELAVL4 | -1.0310686 | 5.85E-08 | HuD | 432358 |
| USO1 | 0.5623836 | 6.19E-08 | TAP\|Vdp | 56042 |
| CITED1 | 1.1248223 | 6.22E-08 | Msg1 | 64466 |
| LOC100911664 | 0.53927714 | 6.60E-08 | | 100911664 |
| CMPK2 | -0.8952171 | 6.66E-08 | Tyki | 314004 |
| PRKG2 | 1.1620419 | 7.14E-08 | GDPKII\|cGKII | 25523 |
| CDCA7L | 0.7639772 | 7.14E-08 | | 619566 |
| MAF1 | -0.6388791 | 7.44E-08 | | 315093 |
| GFPT2 | -1.1508073 | 8.02E-08 | | 360518 |

| | | | | |
|---|---|---|---|---|
| DMKN | 2.0456488 | 8.23E-08 | RGD1561521 | 361548 |
| CAPN2 | 0.5824006 | 8.33E-08 | | 29154 |
| FBXO16 | -1.0601866 | 8.40E-08 | | 305970 |
| TMEM185B | 0.64008254 | 8.72E-08 | RGD1309602 | 304731 |
| RHOV | -1.5801151 | 9.32E-08 | Arhv\|Chp | 171581 |
| TRAFD1 | -0.48449132 | 9.58E-08 | Fln29 | 114635 |
| IER2 | 1.325633 | 9.85E-08 | | 494344 |
| SEC16B | 1.8399274 | 1.10E-07 | Lztr2\|Rgpr | 89868 |
| INHA | -0.918004 | 1.12E-07 | | 24504 |
| LOXL2 | 1.6070518 | 1.12E-07 | | 290350 |
| AK4 | 1.0069963 | 1.17E-07 | AK 4\|Ak3l1\|Ak3l2 | 29223 |
| PFKFB3 | 1.393767 | 1.17E-07 | | 117276 |
| R3HDM1 | 1.0576135 | 1.18E-07 | R3hdm | 304763 |
| SPSB1 | 1.363512 | 1.19E-07 | RGD1309319 | 313722 |
| ARSJ | 1.7609345 | 1.21E-07 | RGD1307640 | 311013 |
| STT3B | 0.8318333 | 1.27E-07 | RGD1311563 | 363160 |
| LRRN1 | -0.65649915 | 1.29E-07 | | 500280 |
| NYAP1 | -0.97091067 | 1.33E-07 | RGD1310964 | 304376 |
| SDCCAG8 | -0.66369975 | 1.33E-07 | | 305002 |
| EIF3B | 0.49294233 | 1.33E-07 | Eif3s9 | 288516 |
| ZC3H18 | 0.51839393 | 1.35E-07 | Nhn1 | 292067 |
| SEMA3D | 1.1440028 | 1.39E-07 | | 246262 |
| MAN1C1 | 1.0923942 | 1.43E-07 | | 362625 |
| UTP4 | 0.62835366 | 1.46E-07 | Cirh1a\|Cirhin\|Naic\|Teg-292\| | 291987 |
| IL34 | -1.3838671 | 1.49E-07 | | 498951 |
| PCDH19 | -0.9742993 | 1.51E-07 | RGD1565392 | 317183 |
| GPR50 | 2.0853536 | 1.51E-07 | H9 | 117097 |
| GMEB1 | 0.9823674 | 1.53E-07 | RGD1563208 | 500558 |
| FHL2 | 0.9380043 | 1.53E-07 | | 63839 |
| CUBN | 2.0080068 | 1.71E-07 | GP280\|IFCR\|MGA1 | 80848 |
| KIF27 | -1.0214224 | 1.71E-07 | Krp5 | 246209 |
| SPATA5 | 0.7741053 | 1.71E-07 | | 361935 |
| MTHFD2 | 0.72602046 | 1.71E-07 | | 680308 |
| PLEKHG5 | 1.1178672 | 1.75E-07 | Tech | 310999 |
| FAM214A | -0.65628093 | 1.77E-07 | RGD1310552 | 300836 |
| GCK | -2.0055048 | 1.81E-07 | GLUKA\|RNGK2 | 24385 |
| KDSR | -0.7911181 | 1.81E-07 | Fvt1 | 360833 |
| WASH1 | -0.5980873 | 1.84E-07 | ORF19\|Wash\|Wash2 | 367328 |
| EXTL2 | -0.638926 | 1.94E-07 | | 310803 |
| NUDT6 | 1.2347888 | 1.98E-07 | Gfg | 207120 |
| THBD | 1.3691242 | 2.02E-07 | | 83580 |
| FRS3 | -1.0738213 | 2.04E-07 | | 316213 |
| PPAN | 0.7546475 | 2.14E-07 | | 298699 |

| | | | | |
|---|---|---|---|---|
| PSEN2 | -0.95383596 | 2.16E-07 | | 81751 |
| TAF12 | 0.7070982 | 2.20E-07 | | 682902 |
| PIEZO1 | 1.3669088 | 2.22E-07 | Fam38a\|Mib | 361430 |
| POLR1B | 0.85832816 | 2.26E-07 | RPA2\|Rpo1-2 | 83582 |
| JAK1 | 0.6872115 | 2.32E-07 | | 84598 |
| ST8SIA5 | 1.5325679 | 2.32E-07 | SIAT8E | 364901 |
| HEATR1 | 0.97933376 | 2.34E-07 | Heatr1 | 103689953 |
| NIPSNAP1 | -0.55160445 | 2.35E-07 | | 360971 |
| DIAPH1 | 0.7853345 | 2.44E-07 | Diap1 | 307483 |
| BID | -0.71934384 | 2.61E-07 | | 64625 |
| SHROOM4 | 1.8252568 | 2.68E-07 | RGD1563434 | 317391 |
| CAMKK1 | 0.8717963 | 2.94E-07 | | 60341 |
| GPR4 | 1.6370957 | 2.95E-07 | | 308408 |
| COL5A1 | 1.3160245 | 3.08E-07 | | 85490 |
| CISH | -1.1958967 | 3.11E-07 | Cis | 83681 |
| WISP1 | 1.8965559 | 3.13E-07 | | 65154 |
| SPIRE1 | 0.5689147 | 3.20E-07 | | 307348 |
| DCLK1 | 0.74735874 | 3.34E-07 | Ania4\|Cpg16\|Dcamkl1 | 83825 |
| KCNMB4 | -0.9178091 | 3.37E-07 | | 66016 |
| MCM5 | 0.86161107 | 3.38E-07 | | 291885 |
| FAM189B | -0.7885296 | 3.38E-07 | RGD1306107 | 310640 |
| PDE6D | -0.7544312 | 3.39E-07 | | 363272 |
| NFE2L3 | -1.3880478 | 3.41E-07 | | 312331 |
| RBM11 | -1.686274 | 3.45E-07 | | 288321 |
| GPR3 | 1.6430944 | 3.48E-07 | Gpcr3 | 266769 |
| CTPS2 | -0.5169076 | 3.54E-07 | | 619580 |
| SEMA4A | -1.0655426 | 3.60E-07 | | 310630 |
| TMEM182 | 1.8668323 | 3.73E-07 | RGD1563696 | 501129 |
| SH3BP5 | -0.78393334 | 3.89E-07 | SH3BP-5\|Sab\|Vesp18 | 117186 |
| ANO1 | 1.9408473 | 3.91E-07 | Tmem16a | 309135 |
| MAGEE2 | -1.1971086 | 3.91E-07 | RGD1564872 | 302392 |
| PHACTR1 | -0.4929978 | 3.94E-07 | | 306844 |
| CRY1 | 0.66173536 | 4.20E-07 | | 299691 |
| BBX | 1.1857326 | 4.25E-07 | | 303970 |
| EXOC5 | 0.6701291 | 4.31E-07 | Sec10l1 | 60627 |
| LIPG | 0.99999833 | 4.38E-07 | lipase | 291437 |
| TUBA8 | 1.4122529 | 4.45E-07 | | 500377 |
| RASIP1 | -0.8784184 | 4.49E-07 | | 292912 |
| PHACTR3 | -0.74370235 | 4.52E-07 | Scapin1\|scapinin | 362284 |
| MORN4 | -0.7344535 | 4.60E-07 | RGD1307336 | 293950 |
| GINS2 | 0.9516325 | 4.76E-07 | RGD1311055 | 292058 |
| TEC | 1.0602809 | 4.79E-07 | | 84492 |
| LIX1 | -0.71060413 | 4.83E-07 | | 292381 |

| | | | | |
|---|---|---|---|---|
| CRELD1 | -0.76312834 | 4.87E-07 | | 312638 |
| PCGF2 | -0.76668996 | 5.66E-07 | Rnf110 | 287662 |
| ARIH1 | 0.778141 | 5.85E-07 | | 300756 |
| ABCG1 | -0.99028426 | 5.87E-07 | Abc8 | 85264 |
| RXFP3 | 1.8421255 | 6.14E-07 | GPCR135\|RGD1311550\|Rln3r | 294807 |
| KCNJ14 | 0.88561434 | 6.25E-07 | Kir2.4 | 276720 |
| PAK1IP1 | 0.8685161 | 6.37E-07 | | 361232 |
| FGF3 | 1.916067 | 6.37E-07 | Int2 | 170633 |
| RPP38 | 0.7958621 | 6.46E-07 | | 291317 |
| MYADML2 | -1.8337954 | 6.52E-07 | RGD1311499 | 303744 |
| USP46 | -0.5572835 | 6.76E-07 | RGD1564808 | 289584 |
| NOG | 1.203545 | 6.77E-07 | | 25495 |
| FAM107B | 0.8400705 | 6.98E-07 | | 498796 |
| NETO2 | 0.7171609 | 7.04E-07 | | 307757 |
| CHST12 | -0.99370766 | 7.57E-07 | | 304322 |
| RAB33A | -1.1109589 | 7.82E-07 | RGD1563280 | 317580 |
| RNF182 | -1.174699 | 8.11E-07 | RGD1560399 | 498726 |
| SMTNL2 | -1.0423588 | 8.31E-07 | | 679629 |
| SFMBT1 | 0.7640687 | 8.31E-07 | Sfmbt | 58967 |
| DOT1L | 1.225937 | 8.76E-07 | | 362831 |
| DHX40 | -0.69222575 | 8.81E-07 | | 287595 |
| NOL10 | 0.62870246 | 9.12E-07 | | 313981 |
| PHLDA1 | 1.0072471 | 9.22E-07 | Tdag | 29380 |
| TYRO3 | 0.7946464 | 9.49E-07 | Brt | 25232 |
| EBF3 | 1.7820029 | 9.59E-07 | | 361668 |
| AGBL5 | -0.7093159 | 9.90E-07 | | 362710 |
| NEUROD1 | -1.4374832 | 9.93E-07 | | 29458 |
| MUM1 | -0.8432682 | 1.11E-06 | | 362838 |
| HEPHL1 | 2.001065 | 1.14E-06 | RGD1564835 | 500946 |
| RHBDD3 | -0.65523726 | 1.19E-06 | RGD1311827 | 289753 |
| PM20D2 | -0.5811111 | 1.20E-06 | Acy1l2 | 313130 |
| MFSD6 | -0.62734413 | 1.21E-06 | RGD1562317 | 301388 |
| TRIM23 | -0.6204789 | 1.21E-06 | Ard1\|Arfd1 | 81002 |
| DLEU7 | -1.3473 | 1.24E-06 | RGD1305071 | 290308 |
| PAK6 | 0.85644025 | 1.27E-06 | | 296078 |
| RHBDF2 | -1.5396582 | 1.28E-06 | Rhbdl6 | 303690 |
| DMTN | -0.46067888 | 1.31E-06 | Epb4.9\|Epb49 | 361069 |
| TMEM169 | -0.63025296 | 1.32E-06 | | 690294 |
| SV2B | -1.1675445 | 1.33E-06 | | 117556 |
| FURIN | 0.85177886 | 1.40E-06 | Pace\|Pcsk3 | 54281 |
| CACNB1 | 0.829659 | 1.48E-06 | | 50688 |
| ACKR3 | -0.89520156 | 1.50E-06 | Cmkor1\|Cxcr7\|Rdc1 | 84348 |
| GPR88 | 0.56655765 | 1.55E-06 | | 64443 |

| | | | | |
|---|---|---|---|---|
| ZDHHC14 | 1.0382525 | 1.59E-06 | Dhhc14 | 499014 |
| KMT2E | -0.5801543 | 1.67E-06 | Mll5 | 311968 |
| FDFT1 | -0.5545826 | 1.74E-06 | | 29580 |
| S100A10 | 0.73875827 | 1.77E-06 | p11 | 81778 |
| APOLD1 | 1.394856 | 1.77E-06 | Verge | 444983 |
| KCNH4 | -1.2192159 | 1.78E-06 | Bec2 | 114032 |
| LRSAM1 | -0.7647853 | 1.81E-06 | RGD1564403 | 311866 |
| KLHL23 | -0.50264704 | 1.82E-06 | RGD1307166 | 311114 |
| TAC1 | 1.7067176 | 1.82E-06 | PPTA3\|Ppt5fl\|RATPPTA3\|TA( | 24806 |
| SLC30A3 | 1.1141921 | 1.84E-06 | ZnT3 | 366568 |
| RIN1 | 1.6419429 | 1.84E-06 | | 207119 |
| ACTR3B | -0.59213006 | 1.90E-06 | RGD1565759 | 362298 |
| CTRB1 | 1.2300013 | 1.91E-06 | Ctrb | 24291 |
| OPRL1 | -0.82617456 | 1.97E-06 | KOR-3\|KOR3\|LC132\|MOR-C\| | 29256 |
| FYTTD1 | 0.55285794 | 1.99E-06 | Ac1176\|RGD1306899 | 360726 |
| JAK2 | 0.8380113 | 1.99E-06 | | 24514 |
| ANKS1B | 0.6663737 | 2.02E-06 | AIDA-1\|EB-1\|RGD1565556 | 314721 |
| HP1BP3 | -0.42026517 | 2.03E-06 | | 313647 |
| HSF1 | 0.59237087 | 2.05E-06 | | 79245 |
| WDR45B | 0.736058 | 2.07E-06 | RGD1305141\|WIPI-3\|Wdr45l | 360682 |
| RAB30 | -0.98914146 | 2.08E-06 | Rsb30 | 308821 |
| FLRT1 | 1.3457637 | 2.08E-06 | RGD1565152 | 499308 |
| MRTO4 | 0.5574501 | 2.12E-06 | RGD1311709 | 298586 |
| GRM1 | -1.2940531 | 2.16E-06 | Gprc1a | 24414 |
| PTGER4 | 1.7010512 | 2.17E-06 | EP4\|Ptger\|Ptgerep4 | 84023 |
| SH2B2 | -0.9922354 | 2.18E-06 | Aps | 114203 |
| NUAK1 | -1.0095615 | 2.19E-06 | RGD1309956 | 299694 |
| NOP56 | 0.6859545 | 2.21E-06 | Nol5a | 362214 |
| C1S | 0.96534324 | 2.28E-06 | r-gsp | 192262 |
| ANKRD34A | -0.8755667 | 2.31E-06 | Ankrd34\|RGD1308412 | 295283 |
| UBE2G2 | 0.7997212 | 2.34E-06 | | 294331 |
| ACVR1C | 1.5148302 | 2.40E-06 | Alk-7\|Alk7\|habrec1 | 245921 |
| P2RY14 | 1.9522917 | 2.41E-06 | Gpr105 | 171108 |
| EZH1 | -0.90584284 | 2.46E-06 | | 303547 |
| CD59 | -0.5351856 | 2.66E-06 | Cd59a\|Cd59b\|MAC-IP\|MACII | 25407 |
| RGD1305537 | 0.8518127 | 2.66E-06 | | 363528 |
| MXD4 | -0.6188515 | 2.75E-06 | | 360961 |
| RWDD2A | -1.2374972 | 2.75E-06 | Rwdd2 | 363110 |
| LEF1 | 1.3694166 | 2.80E-06 | | 161452 |
| MAP7D1 | 0.87675613 | 3.05E-06 | Mtap7d1 | 681287 |
| GPR22 | 0.5187866 | 3.07E-06 | | 298944 |
| MLLT3 | 0.5354938 | 3.07E-06 | Af-9\|Af9 | 114510 |
| GRM6 | -1.2690232 | 3.12E-06 | mGluR6 | 24419 |

| | | | | |
|---|---|---|---|---|
| MED15 | 0.88584226 | 3.17E-06 | Pcqap | 360743 |
| TRMT61A | 0.90787005 | 3.18E-06 | Gcd14\|RGD1359191\|Trm61 | 314462 |
| PNRC1 | -1.1859689 | 3.19E-06 | Prol2 | 286988 |
| TRIM24 | 0.5507146 | 3.20E-06 | | 500084 |
| RYK | 0.48857123 | 3.21E-06 | | 140585 |
| CCDC184 | -0.8354163 | 3.21E-06 | | 500925 |
| PNPLA3 | 0.5412728 | 3.25E-06 | Adpn | 362972 |
| DDX3X | 0.61162984 | 3.26E-06 | | 317335 |
| CBX8 | -0.77355605 | 3.33E-06 | | 303731 |
| AMOTL2 | -0.72246194 | 3.36E-06 | Lccp | 65157 |
| MUSTN1 | 1.5033047 | 3.40E-06 | Mustang | 290553 |
| SMR3B | 1.9007351 | 3.43E-06 | Arp\|P2-VA1\|RATSMR1A\|Smr | 24867 |
| PPP4R1 | 0.5003029 | 3.52E-06 | Pp4r1 | 140943 |
| WIPI2 | 0.59669757 | 3.53E-06 | | 288498 |
| CHRNA5 | -1.0467976 | 3.57E-06 | | 25102 |
| SEMA4G | -0.8667875 | 3.59E-06 | | 361764 |
| ANKRD6 | -0.76407593 | 3.61E-06 | | 500430 |
| STK40 | 0.6637745 | 3.61E-06 | Lyk4\|RGD1307310 | 360230 |
| PSD2 | -0.88123 | 3.78E-06 | | 307500 |
| PTK2 | 0.50821215 | 3.91E-06 | FAK\|FRNK\|p125FAK | 25614 |
| ADAMTS1 | 1.6356838 | 3.93E-06 | | 79252 |
| TMEM2 | 0.9436111 | 3.98E-06 | | 309400 |
| CAMK1G | 0.65445167 | 4.01E-06 | | 171358 |
| DUSP23 | -1.3225392 | 4.08E-06 | | 360881 |
| SLC6A15 | -0.55569607 | 4.13E-06 | Ntt73 | 282712 |
| EVA1A | 1.090327 | 4.15E-06 | Fam176a\|RGD1559797\|Tmem | 500221 |
| FLOT2 | -0.824491 | 4.20E-06 | | 83764 |
| UCK2 | 0.7105085 | 4.28E-06 | Umpk | 304944 |
| RCAN2 | 0.67361695 | 4.29E-06 | Dcip2\|Dscr1l1\|Zaki-4 | 140666 |
| SLC17A5 | -0.5625441 | 4.46E-06 | | 363103 |
| GLRB | -0.499963 | 4.53E-06 | | 25456 |
| POLG | 1.0007834 | 4.56E-06 | | 85472 |
| KY | -1.4094783 | 4.59E-06 | | 315962 |
| LRRC28 | -0.76801777 | 4.67E-06 | | 361588 |
| KLHL7 | -0.6506814 | 4.74E-06 | | 362303 |
| SNAP47 | -0.87896234 | 4.74E-06 | RGD735194\|SNAP-47 | 303183 |
| 42984 | -0.6011752 | 4.76E-06 | | 691335 |
| CHODL | -1.4929341 | 4.78E-06 | | 288289 |
| RGD1560108 | -0.43833774 | 4.79E-06 | | 499309 |
| EDRF1 | 0.54816943 | 4.90E-06 | RGD1306820 | 309069 |
| CNTFR | 0.72213113 | 4.95E-06 | | 313173 |
| LAMB3 | 1.7776339 | 5.01E-06 | | 305078 |
| HS6ST1 | 0.5523936 | 5.03E-06 | | 316325 |

| | | | | |
|---|---|---|---|---|
| PKD2 | 0.824576 | 5.14E-06 | RGD1559992\|Trpp2 | 498328 |
| CTBP2 | 0.5165653 | 5.20E-06 | | 81717 |
| POP1 | 0.5550531 | 5.20E-06 | | 315045 |
| ARHGAP17 | 0.8164248 | 5.39E-06 | Rich1 | 63994 |
| KLF2 | 1.3385966 | 5.52E-06 | Lklf | 306330 |
| SEMA4F | -0.8862101 | 5.93E-06 | | 29745 |
| CDT1 | 0.92074645 | 6.04E-06 | Ris2 | 292071 |
| CD200 | -0.64339024 | 6.06E-06 | Cspmo2\|MRCOX2\|Mox2 | 24560 |
| SIDT1 | -0.6057637 | 6.46E-06 | | 288109 |
| SIGMAR1 | 0.45852426 | 6.50E-06 | Oprs1 | 29336 |
| GPR165 | -1.2783878 | 6.53E-06 | RGD1564566 | 296866 |
| ADGRB3 | -0.7330781 | 6.67E-06 | Bai3 | 301309 |
| THAP3 | -0.98641676 | 6.70E-06 | RGD1308109 | 362667 |
| MBNL2 | -0.6090734 | 7.04E-06 | | 680445 |
| SERPINE1 | 1.582575 | 7.13E-06 | PAI1A\|Pai1\|Pai1aa\|Planh\|RA | 24617 |
| GBX1 | 1.7539651 | 7.19E-06 | | 246149 |
| ZFP746 | 0.50782764 | 7.19E-06 | RGD1306209\|Znf746 | 312303 |
| GABRA1 | -0.7911423 | 7.22E-06 | | 29705 |
| DIP2C | 1.0089853 | 7.51E-06 | RGD1560155 | 307067 |
| PPP2R2B | -0.44699788 | 7.90E-06 | Pppr2b2 | 60660 |
| ARHGAP20 | -1.062759 | 7.97E-06 | RARhoGAP | 367085 |
| GPR6 | -1.4827788 | 8.05E-06 | | 83683 |
| GMEB2 | 1.0013711 | 8.15E-06 | | 83635 |
| PINX1 | 0.8684557 | 8.17E-06 | RGD1566025 | 305963 |
| ROCK2 | 0.7401482 | 8.20E-06 | ROCK-II\|ROK | 25537 |
| HMGA2 | 1.7194638 | 8.42E-06 | Hmgic | 84017 |
| DDX20 | 0.65328026 | 8.54E-06 | Dp103 | 84473 |
| LOC682102 | -0.9888662 | 8.75E-06 | | 682102 |
| RND3 | 0.6508618 | 9.11E-06 | Arhe\|RHOE | 295588 |
| AIPL1 | 1.7215822 | 9.75E-06 | | 59110 |
| PRSS35 | 1.2670162 | 9.79E-06 | | 315866 |
| RXRB | -0.709745 | 1.02E-05 | RXR-beta | 361801 |
| IFT88 | -0.60537183 | 1.02E-05 | Ttc10 | 305918 |
| TOR1B | 0.5286337 | 1.02E-05 | | 311854 |
| RNF39 | 0.929221 | 1.02E-05 | Hzfw1\|Lirf | 171387 |
| LRRC17 | 0.96172076 | 1.03E-05 | | 502715 |
| TMEM143 | -0.8737467 | 1.03E-05 | RGD1305013 | 308593 |
| SH3GL1 | 0.6641487 | 1.05E-05 | SH3P8 | 81922 |
| GNAI1 | 0.4840802 | 1.07E-05 | BPGTPB | 25686 |
| AHI1 | 0.7709468 | 1.11E-05 | Ahi-1 | 308923 |
| CRLF3 | 0.7767652 | 1.12E-05 | Cytor4 | 54395 |
| REEP1 | -0.51963574 | 1.12E-05 | RGD1305230 | 362384 |
| LRRC56 | -0.7891569 | 1.13E-05 | RGD1311654 | 365389 |

| | | | | |
|---|---|---|---|---|
| CALCOCO1 | -0.6872678 | 1.13E-05 | | 246047 |
| ARHGEF9 | 0.41439357 | 1.13E-05 | | 66013 |
| GFOD1 | 1.3111984 | 1.13E-05 | | 306842 |
| DDIT3 | 1.1716156 | 1.13E-05 | CHOP\|CHOP-10\|Chop10\|Gac | 29467 |
| AEN | 0.58869857 | 1.14E-05 | Isg20l1\|RGD1305051 | 361594 |
| SUOX | -0.5694383 | 1.15E-05 | | 81805 |
| ETF1 | 0.7109775 | 1.15E-05 | | 307503 |
| SLAIN2 | 0.45768374 | 1.17E-05 | RGD1311593 | 305310 |
| GPC2 | -0.5883749 | 1.19E-05 | | 171517 |
| MNT | -0.8582651 | 1.21E-05 | | 287521 |
| KLHL36 | -0.8790037 | 1.22E-05 | | 498957 |
| GTPBP4 | 1.0413588 | 1.22E-05 | Crfg\|RGD1563564 | 114300 |
| PLK3 | 0.5974189 | 1.25E-05 | Cnk\|Fnk | 58936 |
| ACSL3 | 0.60574263 | 1.27E-05 | Acs3\|Facl3 | 114024 |
| CHST8 | 0.5964559 | 1.27E-05 | | 308511 |
| DISC1 | 1.3237586 | 1.28E-05 | | 307940 |
| FBN2 | 1.3433425 | 1.30E-05 | | 689008 |
| SKP2 | 0.5353479 | 1.30E-05 | RGD1562456 | 294790 |
| USP16 | 0.87004346 | 1.32E-05 | | 288306 |
| FAM65B | 0.9654181 | 1.32E-05 | Ab2-162\|RGD1306939 | 306934 |
| FZD1 | -0.8275878 | 1.33E-05 | | 58868 |
| UST | 1.0437173 | 1.34E-05 | | 361450 |
| AGFG1 | 0.48620692 | 1.34E-05 | | 363266 |
| ADRA1A | 1.5817711 | 1.34E-05 | Adra1c | 29412 |
| APC2 | -0.98123884 | 1.36E-05 | | 299611 |
| ZFP786 | -0.7909573 | 1.41E-05 | | 100158223 |
| RGS12 | -0.6019134 | 1.41E-05 | | 54292 |
| SRPK1 | 0.5421408 | 1.42E-05 | | 361811 |
| PNOC | 1.4730601 | 1.42E-05 | N23K\|Npnc1 | 25516 |
| JADE3 | 1.0835265 | 1.43E-05 | RGD1563945 | 299305 |
| GPR85 | -0.59075433 | 1.43E-05 | Srep2 | 64020 |
| SCN3B | -0.73834014 | 1.48E-05 | Scnb3 | 245956 |
| MYO9B | 0.7637376 | 1.48E-05 | | 25486 |
| SMPD3 | -0.6857319 | 1.49E-05 | cca1 | 94338 |
| GPR139 | -1.578461 | 1.50E-05 | | 293545 |
| DOCK8 | 1.2737018 | 1.55E-05 | | 499337 |
| KBTBD8 | 0.83205956 | 1.59E-05 | RGD1563166 | 500262 |
| FAM98A | 0.43262425 | 1.65E-05 | RGD1305486 | 313873 |
| QPCT | -0.9462288 | 1.66E-05 | Qpctl1\|RGD1562284 | 313837 |
| RAP1B | 0.7506237 | 1.66E-05 | | 171337 |
| RGD1359127 | 0.82780486 | 1.69E-05 | | 299612 |
| LOC314140 | 0.6524896 | 1.70E-05 | | 314140 |
| LGI1 | -1.0688856 | 1.70E-05 | | 252892 |

| | | | | |
|---|---|---|---|---|
| ADORA2A | -1.735874 | 1.75E-05 | A2ar\|ADENO\|Adora2l1 | 25369 |
| DES | 1.4596198 | 1.75E-05 | | 64362 |
| FAM185A | 0.63159645 | 1.78E-05 | RGD1562135 | 499979 |
| IQSEC3 | 0.91643864 | 1.78E-05 | sag | 404781 |
| GRPEL2 | 0.7165067 | 1.81E-05 | Afap1l1 | 688777 |
| CDR2 | 0.5496149 | 1.81E-05 | RGD1310578 | 308958 |
| TDRD7 | 0.48013663 | 1.84E-05 | Pctaire2bp | 85425 |
| QTRT2 | 0.6236451 | 1.88E-05 | Qtrtd1 | 288364 |
| SPTSSB | -1.2580304 | 1.89E-05 | | 100362555 |
| WBSCR22 | 0.64965624 | 1.89E-05 | | 368084 |
| STRADB | -0.4698745 | 1.90E-05 | Als2cr2\|RGD1559449 | 501146 |
| TAF1C | 0.68430364 | 1.90E-05 | | 361420 |
| FAM110D | -1.0217975 | 1.93E-05 | Grrp1\|RGD1564994 | 500563 |
| CEP70 | -0.81666386 | 1.93E-05 | | 367153 |
| ARHGEF28 | -0.7162797 | 1.96E-05 | Rgnef | 361882 |
| CLN8 | -0.578793 | 1.97E-05 | | 306619 |
| CDC14A | 1.0829693 | 1.97E-05 | | 310806 |
| TOX3 | -0.69983387 | 2.04E-05 | Tnrc9 | 291908 |
| UTP20 | 0.86299187 | 2.14E-05 | RGD1560606 | 314713 |
| BTBD10 | 0.4444746 | 2.15E-05 | RGD1306301 | 308890 |
| MB21D2 | -0.5345009 | 2.21E-05 | RGD1559643 | 498100 |
| RGD1306227 | -0.7342393 | 2.21E-05 | | 310377 |
| FYN | 0.42917925 | 2.24E-05 | | 25150 |
| SPIRE2 | -0.7114268 | 2.24E-05 | | 307925 |
| USP7 | 0.71191597 | 2.34E-05 | Hausp | 360471 |
| TRIQK | -0.8763315 | 2.35E-05 | | 500413 |
| GPR52 | -1.0711756 | 2.39E-05 | | 684623 |
| POLL | -0.72658974 | 2.41E-05 | | 361767 |
| EAF1 | 0.6044422 | 2.41E-05 | | 306261 |
| MAT2B | -0.71800137 | 2.59E-05 | | 683630 |
| APLP2 | 0.4543415 | 2.60E-05 | APLP-2\|Apph\|WALPLP2 | 64312 |
| HOMER2 | 0.74575526 | 2.72E-05 | Vesl-2 | 29547 |
| CCNF | 0.7937148 | 2.72E-05 | | 117524 |
| CCDC28A | -1.1651317 | 2.75E-05 | RGD1310326 | 361454 |
| CARF | -0.8988426 | 2.77E-05 | Als2cr8 | 301446 |
| CSNK2A2 | 0.50321984 | 2.78E-05 | | 307641 |
| ING4 | -0.56335753 | 2.79E-05 | | 297597 |
| RAB20 | 1.3043 | 2.80E-05 | | 689377 |
| CD63 | 0.48863277 | 2.87E-05 | | 29186 |
| ZFP90 | -0.8119344 | 2.88E-05 | | 498945 |
| KL | 1.7558788 | 2.90E-05 | | 83504 |
| NOC3L | 0.7142255 | 2.90E-05 | RGD1560656 | 361753 |
| SYTL5 | 1.532306 | 2.90E-05 | | 302538 |

| | | | | |
|---|---|---|---|---|
| USP36 | 0.6773198 | 2.91E-05 | | 303700 |
| EPHA3 | -1.1606913 | 2.95E-05 | | 29210 |
| LRFN5 | -0.6818887 | 2.95E-05 | | 314164 |
| TMEM25 | -0.8241418 | 2.97E-05 | | 689172 |
| ZFYVE9 | 0.61857194 | 2.99E-05 | | 313477 |
| GCLC | 0.66398966 | 3.03E-05 | Glclc | 25283 |
| PAK7 | -0.50225145 | 3.05E-05 | PAK-5\|PAK-7 | 311450 |
| AKIRIN1 | 0.4189056 | 3.07E-05 | | 595134 |
| DGKA | 0.7506357 | 3.07E-05 | Dagk1 | 140866 |
| ST3GAL3 | -0.68850714 | 3.15E-05 | Siat6 | 64445 |
| IGSF8 | -0.7526169 | 3.16E-05 | | 304979 |
| NR4A1 | 0.8427543 | 3.23E-05 | HMR\|Ngfi-b\|Nur77 | 79240 |
| PDE8A | 1.5737009 | 3.23E-05 | RNPDE8A | 308776 |
| PLAA | 0.5160917 | 3.25E-05 | PLA2P\|Plap | 116645 |
| GOLGB1 | -0.776303 | 3.28E-05 | | 192243 |
| FNDC3B | 1.1798513 | 3.28E-05 | RGD1311673 | 294925 |
| TRMT10C | 0.54266685 | 3.33E-05 | Rg9mtd1 | 304012 |
| LMO7 | -0.53625363 | 3.35E-05 | | 361084 |
| ZFP180 | 0.539561 | 3.35E-05 | rKr1 | 246279 |
| NSUN2 | 0.41097412 | 3.38E-05 | RGD1311954 | 361191 |
| CPNE4 | -0.5717217 | 3.45E-05 | | 367160 |
| PLEKHA3 | 0.5255029 | 3.53E-05 | | 295674 |
| SNTG2 | -1.1350543 | 3.58E-05 | | 298936 |
| PLPP6 | -0.62889516 | 3.58E-05 | Ppapdc2 | 619549 |
| SLC30A4 | 0.54896474 | 3.58E-05 | | 64469 |
| PUS3 | 0.74455327 | 3.64E-05 | RGD1310757 | 315554 |
| LOC100125362 | 0.5085785 | 3.65E-05 | | 100125362 |
| AUH | -0.6037643 | 3.67E-05 | | 361215 |
| DIO3 | 1.3018848 | 3.67E-05 | 5DIII\|DIOIII | 29475 |
| LPL | -0.6849811 | 3.69E-05 | | 24539 |
| ABI1 | 0.47359344 | 3.74E-05 | E3b1 | 79249 |
| LRRC20 | -0.63568723 | 3.79E-05 | RGD1566140 | 499430 |
| ZFP217 | 1.2785103 | 3.79E-05 | Znf217 | 311764 |
| SPTBN4 | 0.690293 | 3.86E-05 | Spnb4 | 308458 |
| PXDC1 | 0.7435586 | 3.88E-05 | RGD1311307 | 361238 |
| ZBTB2 | 0.50025356 | 3.90E-05 | | 308126 |
| LZIC | -0.4655248 | 3.93E-05 | | 366507 |
| ARRDC4 | -1.01276 | 3.93E-05 | Ab1-209\|LRRG00117 | 293019 |
| FHDC1 | 0.9313067 | 3.99E-05 | RGD1311955 | 295161 |
| MFAP3L | 1.1268398 | 4.04E-05 | | 306424 |
| DLG1 | 0.9289756 | 4.09E-05 | Dlgh1\|SAP97 | 25252 |
| GPRC5C | -1.4309026 | 4.09E-05 | | 287805 |
| STOML1 | -0.912747 | 4.12E-05 | RGD1559463 | 300748 |

| | | | | |
|---|---|---|---|---|
| CDC6 | 0.76084006 | 4.13E-05 | | 360621 |
| MAML3 | 1.4926767 | 4.14E-05 | Glrp1 | 310405 |
| MICU3 | 0.52570915 | 4.16E-05 | Efha2 | 364601 |
| CSNK1A1 | 0.44439343 | 4.18E-05 | | 113927 |
| PIANP | -0.7880445 | 4.18E-05 | PANP\|RGD1304952\|leda-1 | 312711 |
| TARS | 0.61816424 | 4.22E-05 | | 294810 |
| PAPPA | 1.6625159 | 4.26E-05 | | 313262 |
| SNX9 | 0.6418308 | 4.33E-05 | | 683687 |
| GMPS | 0.5733719 | 4.34E-05 | | 295088 |
| HSPA1B | 1.301831 | 4.35E-05 | HSP70.2\|Hsp70-1\|Hsp70-2\|H | 294254 |
| FFAR4 | 1.700831 | 4.36E-05 | Gpr120\|O3far1 | 294075 |
| PDE4C | 1.7066025 | 4.45E-05 | Dpde1 | 290646 |
| RYBP | 0.52183354 | 4.59E-05 | | 312603 |
| GPR63 | 1.2530905 | 4.71E-05 | | 297952 |
| GSPT1 | 0.6912279 | 4.78E-05 | | 24420 |
| MAPK15 | -1.170575 | 4.78E-05 | Erk7 | 286997 |
| CNTN2 | -1.07381 | 4.82E-05 | TAG-1\|TAG-564\|TAG1\|Tax | 25356 |
| NFX1 | 0.3948233 | 4.82E-05 | | 313166 |
| KCNJ12 | 0.87608784 | 4.89E-05 | IRK2\|Kir2.1\|Kir2.2 | 117052 |
| NOLC1 | 0.45543104 | 4.90E-05 | Nopp140 | 64896 |
| AFTPH | 0.57212704 | 4.90E-05 | RGD1311920 | 305544 |
| CUX2 | -0.9501184 | 4.99E-05 | Cutl2 | 288665 |
| CHST7 | 0.7684695 | 5.02E-05 | Gn6st-4 | 302302 |
| CDH17 | 1.6838155 | 5.18E-05 | | 117048 |
| TBC1D17 | -0.6419177 | 5.20E-05 | | 292886 |
| ITM2C | -0.7251217 | 5.24E-05 | | 301575 |
| GSPT2 | 0.48441496 | 5.29E-05 | RGD1563213 | 501582 |
| CDK13 | 0.6838744 | 5.43E-05 | Cdc2l5 | 306998 |
| SRD5A1 | -0.5419333 | 5.63E-05 | S5AR 1 | 24950 |
| LOC688459 | 1.670172 | 5.78E-05 | | 688459 |
| KPNA4 | 0.6832937 | 5.95E-05 | importin | 361959 |
| SLC35E3 | 0.3921884 | 5.99E-05 | RGD1564876 | 362883 |
| RABGEF1 | 0.44292256 | 6.03E-05 | | 360797 |
| TTC8 | -0.66576236 | 6.14E-05 | | 299246 |
| PGAM1 | 0.51403296 | 6.19E-05 | PGAM-B\|Pgmut | 24642 |
| LPCAT2 | 0.8675699 | 6.19E-05 | LPCAT-2\|RGD1563994 | 100359680 |
| LSS | 0.47092003 | 6.24E-05 | | 81681 |
| DNAJC12 | -0.8063608 | 6.25E-05 | Jdp1 | 619393 |
| MIOS | 0.52154845 | 6.26E-05 | RGD1308432 | 362324 |
| NUTM1 | 1.1681896 | 6.33E-05 | Nut\|RGD1564529 | 366153 |
| VWC2L | -1.4620417 | 6.33E-05 | RGD1565501 | 501160 |
| ADAMTS5 | 1.2069982 | 6.36E-05 | | 304135 |
| ZBTB5 | -0.68663055 | 6.41E-05 | | 298084 |

| | | | | |
|---|---|---|---|---|
| COBL | -0.50902236 | 6.44E-05 | | 305497 |
| EPHB6 | -0.86060816 | 6.53E-05 | | 312275 |
| MFSD9 | -0.6942437 | 6.59E-05 | RGD1562212 | 316356 |
| TTC9 | 0.6224589 | 6.69E-05 | RGD1565985 | 500689 |
| IL17RA | 0.61163193 | 6.82E-05 | Il17r | 312679 |
| MAFK | -0.61563605 | 7.17E-05 | | 246760 |
| RNF150 | 0.78642243 | 7.18E-05 | RGD1304572 | 364983 |
| ZFAND5 | 0.5366909 | 7.23E-05 | Zfp216 | 293960 |
| PDIK1L | -0.41942006 | 7.28E-05 | RGD1307476\|Stk35l2 | 313609 |
| HAS1 | 1.446656 | 7.31E-05 | | 282821 |
| FAM118A | -0.7019109 | 7.37E-05 | RGD1560783 | 300120 |
| CX3CL1 | 0.60973 | 7.42E-05 | Cx3c\|Scyd1 | 89808 |
| CUL9 | -0.67350787 | 7.45E-05 | Parc\|RGD1562008 | 316228 |
| EIF4A1 | 0.7466136 | 7.47E-05 | | 287436 |
| MMS22L | 0.6067737 | 7.50E-05 | RGD1304693 | 313108 |
| PNCK | -0.8705656 | 7.59E-05 | Camk1b | 29660 |
| CHRM2 | 1.338223 | 7.59E-05 | Acm2 | 81645 |
| ARRB2 | -0.75203335 | 7.63E-05 | BARRES | 25388 |
| UHRF1BP1L | 0.48176548 | 7.63E-05 | | 363009 |
| 42987 | 0.5782184 | 7.70E-05 | Eseptin\|Msf\|Slpa | 83788 |
| BAD | -0.535574 | 7.98E-05 | | 64639 |
| ACTN4 | 0.5552806 | 8.00E-05 | | 63836 |
| MTMR14 | -0.68917155 | 8.07E-05 | RGD1304842 | 312634 |
| TEF | -0.535716 | 8.07E-05 | | 29362 |
| SYK | 0.8088468 | 8.11E-05 | p72syk | 25155 |
| BBC3 | -0.8880315 | 8.11E-05 | Puma | 317673 |
| TMEM53 | -0.9829783 | 8.27E-05 | RGD1307066 | 313529 |
| TOR4A | 1.0359523 | 8.44E-05 | RGD1308019 | 311795 |
| CLIP2 | -0.5369201 | 8.51E-05 | CLIP-115\|Cyln2 | 29264 |
| XKR8 | 0.7255713 | 8.53E-05 | RGD1305649\|XRG8 | 313033 |
| EPDR1 | 0.46413752 | 8.55E-05 | Epdr2\|MERP2 | 291180 |
| DIRAS1 | -0.8976908 | 8.84E-05 | | 366826 |
| IFNGR2 | -0.4437649 | 8.88E-05 | | 360697 |
| PRKAR1A | 0.49870515 | 8.98E-05 | RIIA | 25725 |
| KLHL14 | -1.0380788 | 9.21E-05 | RGD1566178 | 364823 |
| DDX18 | 0.685768 | 9.26E-05 | MrDb | 308490 |
| OTUD3 | 0.9555002 | 9.28E-05 | RGD1560468 | 500572 |
| ARID3C | 1.5952957 | 9.28E-05 | RGD1560943 | 502946 |
| MAST1 | -0.60119736 | 9.42E-05 | Sast | 353118 |
| CIRBP | -0.72022355 | 9.57E-05 | | 81825 |
| ARF6 | 0.44755846 | 9.57E-05 | | 79121 |
| ZHX1 | -0.44386098 | 9.62E-05 | | 171159 |
| DDAH2 | -0.56704617 | 9.77E-05 | | 294239 |

| | | | | |
|---|---|---|---|---|
| ARG2 | 0.5525799 | 1.01E-04 | | 29215 |
| EPB41L4B | 1.0035497 | 1.02E-04 | RGD1562988 | 500464 |
| UNC119 | -0.6921164 | 1.03E-04 | RRG4\|Uncl19 | 29402 |
| AMIGO2 | -0.78318065 | 1.03E-04 | Ali1 | 300186 |
| CYS1 | -0.99969214 | 1.03E-04 | | 690489 |
| POU6F1 | 0.9946661 | 1.03E-04 | Brn5 | 116545 |
| VIPAS39 | -0.49750003 | 1.04E-04 | Spe39\|Vipar\|hSPE-39 | 681989 |
| GDNF | 1.6205387 | 1.07E-04 | gndf | 25453 |
| MPEG1 | -1.1206188 | 1.09E-04 | Mpg-1 | 64552 |
| PALMD | -0.99204695 | 1.10E-04 | | 310811 |
| CXCR4 | -0.614934 | 1.11E-04 | | 60628 |
| COQ10B | 0.5414451 | 1.11E-04 | RGD1359509 | 301416 |
| LDLRAD4 | 0.8536135 | 1.11E-04 | | 679578 |
| LUZP1 | 0.72387207 | 1.11E-04 | Luzp | 79428 |
| PPRC1 | 0.7030932 | 1.13E-04 | | 294007 |
| MSS51 | 0.8746994 | 1.14E-04 | Zmynd17 | 289904 |
| TPPP3 | -1.1469681 | 1.15E-04 | CGI-38\|RGD1305061 | 291966 |
| VPS9D1 | -0.5139628 | 1.16E-04 | RGD1565149 | 307923 |
| CTU2 | 0.70760155 | 1.16E-04 | Ncs2 | 292069 |
| LYPD1 | -0.7614784 | 1.17E-04 | Lypdc1 | 360838 |
| SPEM1 | 0.72259897 | 1.18E-04 | | 691981 |
| USP28 | 0.65976477 | 1.21E-04 | | 315639 |
| PRR15 | 1.2841624 | 1.21E-04 | RGD1311589 | 312358 |
| CEP131 | -0.7585036 | 1.23E-04 | Azi1 | 360672 |
| WFS1 | 0.7863216 | 1.23E-04 | | 83725 |
| PARP16 | -0.77560216 | 1.23E-04 | ARTD15\|LRRGT00109\|RGD13 | 315760 |
| COX19 | -0.611949 | 1.24E-04 | RGD1305631 | 304330 |
| SLC35F2 | -0.8826118 | 1.25E-04 | | 300713 |
| LRRC10B | 1.3151493 | 1.25E-04 | RGD1564983 | 309208 |
| STEAP1 | 1.3752855 | 1.25E-04 | Steap | 297738 |
| LMO2 | 1.2145776 | 1.26E-04 | | 362176 |
| BMP6 | 0.74068916 | 1.27E-04 | VGR | 25644 |
| GRIN1 | 0.56461716 | 1.36E-04 | GluN1\|NMDAR1\|NR1 | 24408 |
| PTCHD1 | 1.2826488 | 1.36E-04 | RGD1564527 | 317517 |
| ARPC5 | 0.51324344 | 1.36E-04 | | 360854 |
| FAM136A | -0.5223806 | 1.39E-04 | RGD1304825 | 297415 |
| CYP2S1 | -1.3139243 | 1.41E-04 | | 308445 |
| PNPO | -0.6106862 | 1.41E-04 | | 64533 |
| MVK | -0.59577304 | 1.42E-04 | Lrbp | 81727 |
| IMPDH2 | 0.5349753 | 1.44E-04 | IMPD 2\|IMPDH 2 | 301005 |
| SOGA3 | -0.4308755 | 1.45E-04 | Kiaa0408\|RGD1307525\|Rgd1 | 292199 |
| CHRNA3 | -1.1553017 | 1.46E-04 | | 25101 |
| TRPV2 | -0.7396124 | 1.49E-04 | Vrl1 | 29465 |

| | | | | |
|---|---|---|---|---|
| B4GALT7 | -0.6417235 | 1.52E-04 | | 364675 |
| ZFP467 | -0.6218755 | 1.52E-04 | Znf467 | 500110 |
| GPSM2 | 0.4762065 | 1.52E-04 | RGD1560967 | 362021 |
| TBKBP1 | 0.5302589 | 1.55E-04 | Prosapip2 | 266764 |
| ICK | -0.6124105 | 1.57E-04 | | 84411 |
| NAB1 | 0.438405 | 1.61E-04 | | 64824 |
| SLC18A2 | 0.5498699 | 1.62E-04 | MNAT\|VMAT-2\|VMAT2 | 25549 |
| PAPOLA | 0.44860807 | 1.63E-04 | | 314417 |
| RALYL | -0.802912 | 1.65E-04 | RGD1305844 | 294883 |
| ZFAT | 0.6622704 | 1.66E-04 | Zfat1\|Zfp406 | 362925 |
| TGFBR2 | 1.289802 | 1.66E-04 | TGF-beta 2\|Tgfbr2T | 81810 |
| POLR1C | 0.42630336 | 1.67E-04 | Ac2-127\|Rpo1-1 | 301246 |
| PTER | 1.0322663 | 1.69E-04 | Rpr-1\|Rpr1 | 63852 |
| CDYL2 | 0.95827675 | 1.69E-04 | | 292044 |
| AKAP7 | -0.7377203 | 1.69E-04 | AKAP-18\|AKAP18d\|Akap15\|A | 361458 |
| EFNB3 | -0.5920111 | 1.69E-04 | ELK-L3 | 360546 |
| GIPC3 | 1.6110032 | 1.69E-04 | RGD1563255 | 500789 |
| NOCT | 0.5487951 | 1.74E-04 | Ccr4\|Ccrn4l\|Ccrn4lb | 310395 |
| BCAS3 | -0.5178273 | 1.74E-04 | RGD1560788 | 363662 |
| DAZAP1 | 0.45886728 | 1.75E-04 | | 362836 |
| MXI1 | -0.47463524 | 1.76E-04 | MXI-WR | 25701 |
| HDAC5 | 0.48885363 | 1.77E-04 | | 84580 |
| MC5R | 1.6079197 | 1.77E-04 | | 25726 |
| POMK | -0.58920217 | 1.78E-04 | RGD1310810\|Sgk196 | 306549 |
| PODXL | -0.6923312 | 1.78E-04 | PC\|PCLP-1\|podocalyxin | 192181 |
| IRF6 | 1.2191257 | 1.82E-04 | | 364081 |
| PECAM1 | -1.3774275 | 1.82E-04 | CD31\|Pecam | 29583 |
| TRPV6 | -0.93850434 | 1.82E-04 | CaT1\|Ecac2\|Otrpc3 | 114246 |
| RGD1308117 | -0.9308683 | 1.82E-04 | | 361066 |
| ADAMTS2 | -1.2053839 | 1.84E-04 | RGD1565950 | 287899 |
| TNFRSF12A | 1.0359695 | 1.85E-04 | Fn14 | 302965 |
| ACOT3 | -0.62287575 | 1.85E-04 | RGD1564089 | 314304 |
| MMP17 | 0.8486851 | 1.88E-04 | | 288626 |
| TMEM68 | -0.5819448 | 1.88E-04 | RGD1309006 | 312946 |
| METTL13 | 0.46811473 | 1.89E-04 | RGD1311526 | 289159 |
| GALNT16 | -0.53217185 | 1.91E-04 | Galntl1 | 362760 |
| DAB1 | 0.71107423 | 1.92E-04 | | 266729 |
| IL1RAP | 0.9475349 | 2.00E-04 | IL-1RAcp\|Il1racpb | 25466 |
| ETV4 | 0.7502681 | 2.00E-04 | Pea3 | 360635 |
| PITX1 | 1.5863794 | 2.00E-04 | Bft | 113983 |
| CLASP2 | -0.45628664 | 2.00E-04 | | 114514 |
| TADA2B | 0.43645033 | 2.01E-04 | ADA2B\|RGD1561605 | 289717 |
| LTBP1 | 0.91750896 | 2.01E-04 | | 59107 |

| | | | | |
|---|---|---|---|---|
| DUSP26 | -0.6058669 | 2.02E-04 | RGD1310090 | 306527 |
| ADAMTS7 | -0.74705875 | 2.04E-04 | ADAMTS7B\|COMPase | 315879 |
| FAIM2 | -0.72629684 | 2.05E-04 | Lfg\|NMP35 | 246274 |
| CYB5R1 | -0.7076814 | 2.05E-04 | Nqo3a2 | 304805 |
| MEPE | 1.3387411 | 2.06E-04 | | 79110 |
| PTPN1 | 0.64497983 | 2.07E-04 | Ptp | 24697 |
| TPM4 | 0.82039404 | 2.07E-04 | Tpm4.2\|Tpm4.2cy | 24852 |
| RGS2 | 0.7381537 | 2.08E-04 | | 84583 |
| TEX10 | 0.5948173 | 2.11E-04 | | 298065 |
| KCNB1 | 0.86163217 | 2.11E-04 | DRK1PC\|Kcr1-1\|Kv2.1\|Shab | 25736 |
| CBFB | 0.6844125 | 2.12E-04 | Pebp2 | 361391 |
| TP73 | 1.3698492 | 2.13E-04 | P73\|Trp73 | 362675 |
| STAU1 | 0.3779561 | 2.14E-04 | Stau | 84496 |
| ARL14EP | 0.7410797 | 2.14E-04 | ARF7EP\|RGD1311463 | 311279 |
| IGF2 | 1.5672538 | 2.17E-04 | IGFII\|RNIGF2 | 24483 |
| CDK19 | -0.5549146 | 2.17E-04 | Cdc2l6\|Cdk11 | 309804 |
| LRRC59 | 0.4165463 | 2.20E-04 | | 287633 |
| RRS1 | 0.7456884 | 2.28E-04 | | 297784 |
| ANKRD12 | -0.6539705 | 2.30E-04 | | 316775 |
| LRRC4 | 0.68762976 | 2.31E-04 | | 641521 |
| RAB15 | 0.59578437 | 2.36E-04 | | 299156 |
| PPP2R3A | -0.8871097 | 2.40E-04 | | 363122 |
| OS9 | -0.4035507 | 2.40E-04 | Os-9 | 362891 |
| SEMA3E | 0.8192995 | 2.40E-04 | | 296789 |
| CMAS | -0.65439934 | 2.42E-04 | | 312826 |
| OPRD1 | 1.2956082 | 2.46E-04 | | 24613 |
| NLN | 0.36868855 | 2.46E-04 | | 117041 |
| HSD11B1 | 1.0060592 | 2.47E-04 | LRRGT00065 | 25116 |
| MRRF | -0.47376493 | 2.50E-04 | | 311903 |
| 42803 | -0.62812364 | 2.53E-04 | March-IX\|Rnf179 | 679272 |
| LRTM2 | -1.2352533 | 2.57E-04 | | 680883 |
| LHX2 | -0.7194863 | 2.57E-04 | LH2A\|Lh-2 | 296706 |
| GADD45A | 0.8438053 | 2.60E-04 | Ddit1\|Gadd45 | 25112 |
| PLCL2 | 0.49759674 | 2.67E-04 | | 301173 |
| UTP15 | 0.69488347 | 2.68E-04 | RGD1310992 | 310019 |
| PRUNE | -0.5479131 | 2.71E-04 | | 310664 |
| SMOC2 | -1.477256 | 2.71E-04 | | 292401 |
| YARS | 0.430864 | 2.71E-04 | | 313047 |
| RRP8 | 0.6214512 | 2.71E-04 | RGD1308302 | 308911 |
| ARHGAP31 | 1.1611425 | 2.71E-04 | Cdgap | 288093 |
| DESI1 | 0.42232907 | 2.72E-04 | DeSI-1\|Fam152b\|Pppde2\|RG | 315160 |
| RHEB | 0.5235065 | 2.74E-04 | | 26954 |
| ASGR1 | 1.3252411 | 2.80E-04 | ASGR\|RATRHL1\|RHL1 | 24210 |

| | | | | |
|---|---|---|---|---|
| PFKP | 0.4141759 | 2.84E-04 | ATP-PFK\|PFK-C\|PFK-P | 60416 |
| P4HA1 | 0.66392094 | 2.85E-04 | PHalphaI | 64475 |
| ARMC7 | -0.6029792 | 2.85E-04 | RGD1310043 | 287827 |
| SFMBT2 | 1.2059495 | 2.86E-04 | | 307106 |
| OSBPL6 | 0.9556358 | 2.87E-04 | | 311129 |
| USP20 | -0.5476291 | 2.93E-04 | | 311856 |
| ADAMTS15 | -0.94080544 | 2.93E-04 | | 300474 |
| CTDP1 | 0.4451366 | 2.93E-04 | | 291414 |
| ZW10 | 0.4267897 | 2.94E-04 | | 363059 |
| AP3S2 | -0.52299076 | 3.01E-04 | | 683402 |
| CDYL | 0.79401857 | 3.01E-04 | | 361237 |
| E2F4 | 0.48663586 | 3.02E-04 | | 100360427 |
| NDRG1 | 0.6972355 | 3.02E-04 | Ndr1 | 299923 |
| PPM1H | 0.7127094 | 3.03E-04 | | 314897 |
| NCOR2 | 0.8484824 | 3.11E-04 | | 360801 |
| NFATC1 | 0.7563985 | 3.12E-04 | | 100361818 |
| RGD1311164 | -0.46310118 | 3.15E-04 | | 297607 |
| SLC6A7 | -0.712864 | 3.28E-04 | Prot | 117100 |
| NFRKB | 0.6441671 | 3.31E-04 | | 315523 |
| ABHD6 | -0.4444501 | 3.32E-04 | | 305795 |
| KCND2 | -0.5871494 | 3.33E-04 | Kv4.2\|RK5\|Shal1 | 65180 |
| NRSN1 | 0.46664348 | 3.35E-04 | Vmp | 291129 |
| ADD3 | -0.5705513 | 3.37E-04 | | 25230 |
| CYB561 | -0.60420316 | 3.39E-04 | | 303601 |
| ELP2 | -0.41660187 | 3.39E-04 | Statip1 | 307545 |
| OSBPL1A | -0.36456922 | 3.50E-04 | | 259221 |
| RGD1312005 | -0.9231772 | 3.50E-04 | | 291580 |
| AMIGO3 | 0.73918504 | 3.52E-04 | | 316003 |
| TEX264 | -0.50394505 | 3.53E-04 | | 300988 |
| SIRT5 | -0.6803784 | 3.54E-04 | | 306840 |
| ABCD1 | -0.59828013 | 3.59E-04 | RGD1562128 | 363516 |
| ATP13A3 | 0.7735666 | 3.59E-04 | | 678704 |
| ARFGAP3 | -0.45546272 | 3.60E-04 | | 503165 |
| RALGDS | -0.51777 | 3.62E-04 | Rgds | 29622 |
| ADGRE5 | 1.0853325 | 3.64E-04 | Cd97 | 361383 |
| CEBPA | 0.7842719 | 3.67E-04 | DBPCEP | 24252 |
| ATP5SL | -0.4827523 | 3.72E-04 | | 361520 |
| IFT46 | -0.6445975 | 3.72E-04 | RGD1307682 | 300675 |
| YTHDF3 | 0.40102714 | 3.86E-04 | | 361920 |
| ESPNL | 1.5138139 | 3.86E-04 | RGD1562432 | 301606 |
| NIP7 | 0.4737033 | 3.90E-04 | CGI-37\|Nip7p\|pEachy | 192180 |
| LOC691141 | 1.0126132 | 3.96E-04 | | 691141 |
| BLOC1S6 | -0.54068834 | 4.01E-04 | Pldn | 317630 |

| | | | | |
|---|---|---|---|---|
| LRRC8B | 1.0917848 | 4.03E-04 | RGD1563429 | 305135 |
| ECH1 | -0.6108725 | 4.03E-04 | Pxel | 64526 |
| KDM1A | 0.36938342 | 4.03E-04 | Aof2\|Kdm1\|RGD1562975 | 500569 |
| DNASE2B | 1.5245335 | 4.18E-04 | Dlad\|UOX | 59296 |
| CYP11B2 | 1.5138268 | 4.25E-04 | Cp45as\|Cyp11b3\|RNCP45AS | 24294 |
| SERTM1 | 0.830482 | 4.26E-04 | | 690333 |
| ANAPC2 | -0.45406723 | 4.37E-04 | | 296558 |
| CAND1 | 0.57176256 | 4.39E-04 | Tip120\|Tip120A | 117152 |
| FAM20B | 0.37168434 | 4.45E-04 | RGD1311162 | 304885 |
| TCP11L2 | -0.7020828 | 4.48E-04 | RGD1307494 | 314683 |
| SRC | 0.7262243 | 4.55E-04 | | 83805 |
| NEBL | 0.81218123 | 4.55E-04 | Lasp-2 | 307189 |
| TMIE | -1.1572524 | 4.56E-04 | RGD1562523 | 501061 |
| QPCTL | -0.47004032 | 4.56E-04 | RGD1308128 | 292687 |
| SERPINB7 | 1.505848 | 4.56E-04 | Megsin | 117092 |
| NOC4L | 0.5610823 | 4.58E-04 | RGD1310661 | 360828 |
| HAGH | -0.6924093 | 4.59E-04 | Glo2\|RSP29 | 24439 |
| PCDHA2 | -0.5102854 | 4.59E-04 | rCNRv02 | 393086 |
| TSPAN2 | -0.4740401 | 4.59E-04 | Tspan-2 | 64521 |
| NRIP3 | 0.5417548 | 4.59E-04 | | 361625 |
| DSE | 0.5761879 | 4.59E-04 | Sart2 | 365583 |
| TRMT44 | 0.69999003 | 4.66E-04 | Mettl19\|RGD1308380 | 305443 |
| PCDHA12 | -0.5639758 | 4.67E-04 | rCNRv12 | 116779 |
| OXTR | 1.3693583 | 4.69E-04 | OT-R\|OTR\|OTR1 | 25342 |
| ZBTB22 | -0.48097795 | 4.72E-04 | Bing1\|Zfp297\|Znf297 | 309630 |
| REM2 | 0.7931316 | 4.73E-04 | | 64626 |
| PRRT1 | -0.61716336 | 4.87E-04 | DSPD1\|Ng5\|Orf31 | 406167 |
| GRHL2 | 1.4127645 | 4.90E-04 | RGD1561191 | 299979 |
| RBBP5 | 0.38272616 | 4.91E-04 | | 304794 |
| CLK3 | 0.43377548 | 4.91E-04 | | 171305 |
| AGPAT5 | 0.37164775 | 4.95E-04 | RGD1306405 | 306582 |
| PRKX | 0.47665304 | 4.96E-04 | | 501563 |
| BTAF1 | 0.86330533 | 5.02E-04 | RGD1564130 | 368042 |
| PCSK4 | -1.0208651 | 5.09E-04 | | 171085 |
| MPPED1 | -0.6468225 | 5.10E-04 | RGD1308244 | 362971 |
| RGD1310110 | -0.6975931 | 5.15E-04 | | 361032 |
| DNAJC25 | 0.5981939 | 5.15E-04 | | 362526 |
| H2AFY2 | -0.4295681 | 5.20E-04 | RGD1561371 | 361844 |
| POLRMT | 0.61828196 | 5.28E-04 | | 299604 |
| NACAD | -0.5167076 | 5.31E-04 | | 289786 |
| PCSK5 | 0.6855457 | 5.31E-04 | PC6\|Pc5 | 116548 |
| STK17B | 0.713867 | 5.33E-04 | Drak2 | 170904 |
| ZFP36L2 | 0.57546765 | 5.41E-04 | | 298765 |

| | | | | |
|---|---|---|---|---|
| RGD1305938 | -0.5517365 | 5.43E-04 | | 310362 |
| POLH | 0.64425135 | 5.48E-04 | | 316235 |
| RBM14 | 0.50478554 | 5.49E-04 | CoAA | 170900 |
| PLAGL1 | 0.6053042 | 5.49E-04 | Lot1\|Zac1 | 25157 |
| MARK3 | 0.3763424 | 5.61E-04 | | 170577 |
| PCMTD2 | -0.57118213 | 5.62E-04 | RGD1305684 | 311726 |
| ZFP362 | -0.6280068 | 5.62E-04 | RGD1306498 | 297879 |
| HCN1 | 0.86012423 | 5.63E-04 | | 84390 |
| GPR21 | -0.44384855 | 5.65E-04 | Gpr21 | 311911 |
| GPR45 | -0.5449812 | 5.67E-04 | | 301372 |
| DBP | -0.8268259 | 5.68E-04 | | 24309 |
| PCDHA6 | -0.50182587 | 5.68E-04 | rCNRv06 | 393088 |
| EMD | 0.42466643 | 5.68E-04 | | 25437 |
| GPR156 | -1.0159905 | 5.73E-04 | Gababl | 260430 |
| SLC48A1 | -0.3968802 | 5.78E-04 | HRG-1\|Hrg1 | 300191 |
| PRKCZ | -0.5678957 | 5.87E-04 | 14-3-3-zetaisoform\|Pkcz\|r14 | 25522 |
| FTSJ3 | 0.51962984 | 6.00E-04 | | 303608 |
| NMT2 | 0.43841127 | 6.07E-04 | | 291318 |
| CHRAC1 | -0.6103794 | 6.09E-04 | | 315058 |
| HSPA2 | 0.5357155 | 6.14E-04 | Hspt70\|Hst70 | 60460 |
| TMEM86A | -0.7264606 | 6.16E-04 | RGD1305749 | 308602 |
| SLC40A1 | 1.4047453 | 6.16E-04 | Fpn1\|Slc11a3\|Slc39a1 | 170840 |
| SLC37A4 | -0.5257178 | 6.18E-04 | G6pt1 | 29573 |
| LOC500877 | -1.3927882 | 6.18E-04 | | 500877 |
| LINGO1 | 0.6413327 | 6.21E-04 | Lrrn6a | 315691 |
| PRMT2 | -0.62597674 | 6.24E-04 | Hrmt1l1 | 499420 |
| SLC28A3 | -1.4889654 | 6.31E-04 | Cnt3 | 140944 |
| CAMK2N1 | 0.49420154 | 6.33E-04 | | 287005 |
| TMTC3 | 0.5425814 | 6.33E-04 | RGD1306351 | 314785 |
| TACR2 | 1.4816157 | 6.36E-04 | Tac2r | 25007 |
| CITED4 | -1.242663 | 6.37E-04 | MRG-2\|Mrg2 | 114491 |
| PRKRA | -0.56349343 | 6.37E-04 | RAX | 311130 |
| ABCB10 | -0.54307234 | 6.38E-04 | | 361439 |
| SEC24D | 0.83843863 | 6.48E-04 | | 310843 |
| LEPROT | -0.4193454 | 6.53E-04 | Ob-Rgrp\|Obrgrp | 56766 |
| PKLR | -0.9121864 | 6.56E-04 | PK1\|PKL\|Pklg | 24651 |
| NOC2L | 0.3833833 | 6.76E-04 | RGD1309387 | 313777 |
| TTLL1 | -0.56456447 | 6.79E-04 | | 362969 |
| ZNRF2 | -0.70722926 | 6.85E-04 | Znrf1 | 362367 |
| RIT2 | -0.5996122 | 6.87E-04 | Rin\|Rit1 | 291713 |
| CALHM2 | -1.169739 | 6.87E-04 | Fam26b\|RGD1308276 | 294019 |
| PNMAL2 | -0.50676274 | 6.95E-04 | RGD1560435 | 308393 |
| XPO4 | 0.72588956 | 6.95E-04 | | 290280 |

| | | | | |
|---|---|---|---|---|
| PACSIN3 | -0.63019615 | 6.97E-04 | SdpIII | 311187 |
| SYNGR3 | -0.4901226 | 6.97E-04 | | 302975 |
| JRK | 0.63231033 | 6.97E-04 | | 315073 |
| CALCRL | 0.9037969 | 6.98E-04 | CLR\|Crlr\|RATCRLR\|RNCLR | 25029 |
| CYHR1 | -0.8087888 | 6.99E-04 | RGD1561328 | 100362155 |
| KIF5B | 0.42993933 | 6.99E-04 | Khc | 117550 |
| NFKBIE | -0.7287623 | 7.15E-04 | Slc35b2 | 316241 |
| OSBPL2 | -0.37741908 | 7.17E-04 | | 296461 |
| PLEKHA5 | 0.5339661 | 7.17E-04 | Pepp2\|TRS1 | 246237 |
| PDE10A | 0.9542575 | 7.17E-04 | Pde10a3 | 63885 |
| STRADA | -0.49474013 | 7.18E-04 | Lyk5 | 303605 |
| RASAL2 | 0.9257782 | 7.19E-04 | | 304893 |
| BEND7 | 1.0420662 | 7.35E-04 | RGD1305898 | 361275 |
| PHC1 | 0.48105434 | 7.35E-04 | | 312690 |
| IGF2BP2 | 1.0324925 | 7.49E-04 | RGD1305614 | 303824 |
| TPK1 | -0.6448231 | 7.55E-04 | | 680668 |
| CHD6 | -0.6291748 | 7.64E-04 | CHD-6 | 311607 |
| CHKA | 0.42648286 | 7.71E-04 | CK-R\|Chk | 29194 |
| TMUB2 | -0.3868561 | 7.73E-04 | RGD1304758 | 303567 |
| DOC2A | -0.58669204 | 7.80E-04 | | 65031 |
| USP10 | 0.53595936 | 7.80E-04 | | 307905 |
| CTC1 | -0.45218167 | 7.83E-04 | RGD1563106 | 303238 |
| B3GNT2 | -0.6569873 | 7.84E-04 | B3gnt1 | 305571 |
| CCKAR | -1.4610728 | 7.84E-04 | Cck-ar | 24889 |
| OGDHL | -0.44704428 | 7.85E-04 | | 290566 |
| NKD1 | -0.644302 | 7.96E-04 | | 364952 |
| PDE7B | 0.7817121 | 8.00E-04 | | 140929 |
| PHF20 | -0.40930417 | 8.04E-04 | RGD1305020 | 311575 |
| ZC3H6 | -0.8585249 | 8.05E-04 | Zc3h6 | 103690012 |
| FSD1 | -0.5281462 | 8.13E-04 | | 301506 |
| P2RY1 | -0.8910657 | 8.19E-04 | P2y\|P2y1 | 25265 |
| PURB | 0.4570141 | 8.19E-04 | pur-beta | 498407 |
| TMCC2 | -0.6654097 | 8.26E-04 | RGD1311960 | 305095 |
| ILVBL | -0.6613915 | 8.34E-04 | | 362843 |
| CBFA2T3 | 0.8064325 | 8.46E-04 | | 361431 |
| TWIST1 | 1.1040758 | 8.48E-04 | Twist | 85489 |
| PHLPP1 | 0.5765138 | 8.52E-04 | Phlpp\|Plekhe1\|Scop | 59265 |
| PWP2 | 0.4914513 | 8.54E-04 | | 690297 |
| SHQ1 | 0.618061 | 8.56E-04 | RGD1310610 | 297483 |
| NCALD | -0.3158198 | 8.67E-04 | | 553106 |
| TIGAR | 0.42375776 | 8.67E-04 | | 502894 |
| BYSL | 0.44311565 | 8.71E-04 | | 359727 |
| MLF1 | 0.7294841 | 8.72E-04 | | 310483 |

| | | | | |
|---|---|---|---|---|
| BHLHE23 | 1.2740167 | 8.75E-04 | Bhlhb4\|RGD1559760 | 499952 |
| VAC14 | -0.38587335 | 8.80E-04 | | 307842 |
| KCTD6 | 0.4704236 | 8.86E-04 | | 305792 |
| MAD2L2 | -0.52181566 | 8.86E-04 | | 313702 |
| RNF4 | 0.43907994 | 8.86E-04 | | 29274 |
| TUSC5 | 1.3077588 | 8.86E-04 | DSPB1 | 360576 |
| D2HGDH | -0.7407282 | 9.03E-04 | RGD1307976 | 301624 |
| PPP1R21 | -0.44658285 | 9.11E-04 | RGD1565310 | 362697 |
| PIGW | 0.58660764 | 9.11E-04 | | 378774 |
| SH3BP1 | 0.7184306 | 9.11E-04 | | 300067 |
| FOPNL | -0.64248985 | 9.11E-04 | RGD1305823 | 360461 |
| RITA1 | -0.6617135 | 9.16E-04 | RGD1306772\|Rita | 288683 |
| DHX15 | 0.37572557 | 9.16E-04 | | 289693 |
| TXNRD1 | 0.43897274 | 9.16E-04 | Tr | 58819 |
| TRPM4 | -0.7388621 | 9.16E-04 | LTrpC-4\|Mls2s | 171143 |
| WNT7A | -0.44404274 | 9.17E-04 | | 114850 |
| SLC24A4 | 0.65773785 | 9.17E-04 | Nckx4 | 314396 |
| RBFA | -0.6280929 | 9.22E-04 | RGD1311910 | 307235 |
| LIPA | -0.48053014 | 9.24E-04 | Chole\|Chole2\|LAL\|Lip1 | 25055 |
| BACH1 | 0.8233174 | 9.38E-04 | | 304127 |
| APEX2 | 0.5453076 | 9.39E-04 | | 317628 |
| SYT9 | -0.6798635 | 9.45E-04 | Sytv | 60564 |
| PCSK7 | 0.42560545 | 9.45E-04 | PC7 | 29606 |
| DGCR8 | 0.3542616 | 9.58E-04 | | 287954 |
| ATP1A1 | 0.5371216 | 9.66E-04 | Nkaa1b | 24211 |
| KLHL21 | 0.52049124 | 9.70E-04 | RGD1305863 | 313743 |
| DMTF1 | 0.74144053 | 9.78E-04 | | 114485 |
| PDK1 | 0.49384907 | 9.83E-04 | | 116551 |
| VLDLR | 0.4947575 | 9.90E-04 | | 25696 |
| INPP5D | 0.9800654 | 9.92E-04 | | 54259 |
| COASY | -0.5244441 | 9.95E-04 | | 287711 |
| AATK | -0.7821023 | 9.97E-04 | | 690853 |
| TFRC | 0.6394503 | 0.001003031 | Trfr | 64678 |
| IVNS1ABP | 0.55952954 | 0.001011257 | | 289089 |
| FZD4 | 0.8261322 | 0.001014388 | | 64558 |
| MYOCD | 1.4390517 | 0.001021537 | Mycd | 246297 |
| P2RX2 | -1.0103608 | 0.001063429 | P2X2 | 114115 |
| WNT3 | -1.0721376 | 0.001075465 | INT4 | 24882 |
| NGLY1 | -0.5888386 | 0.001078654 | | 361014 |
| DCAF4 | -0.5240574 | 0.001085269 | Wdr21 | 362762 |
| SOX4 | -0.666895 | 0.001087743 | | 364712 |
| CDH15 | -1.0086491 | 0.00109044 | | 361432 |
| SHMT2 | 0.7363225 | 0.001096925 | | 299857 |

| | | | | |
|---|---|---|---|---|
| NOP9 | 0.46830082 | 0.001098523 | RGD1308396 | 290235 |
| DIEXF | 0.6520972 | 0.001115796 | Def | 305076 |
| XPO1 | 0.64450747 | 0.001121369 | | 85252 |
| CHD1 | 0.70372075 | 0.001121369 | | 308215 |
| OSBPL8 | 0.42345697 | 0.001121503 | RGD1561474 | 314824 |
| FRMD3 | -0.49199355 | 0.001123587 | | 298141 |
| IFT52 | -0.48374653 | 0.001123977 | RGD1311004 | 362265 |
| MMP16 | 0.7201553 | 0.001137568 | Mt3-mmp | 65205 |
| NTF3 | -0.91430616 | 0.001137797 | | 81737 |
| UGCG | -0.408513 | 0.001153735 | | 83626 |
| PPIF | 0.4912731 | 0.001161996 | CyP-D\|CypD\|PPIase | 282819 |
| ACAP1 | 0.87039363 | 0.001164913 | Centb1 | 287443 |
| LANCL2 | -0.40217626 | 0.001165692 | | 362375 |
| LOC100302465 | 1.327915 | 0.001170263 | | 100302465 |
| GPR176 | -0.6699465 | 0.001175242 | Gm1012\|Gpr\|RBU-15 | 117257 |
| BORCS5 | -0.97918296 | 0.00118038 | Loh12cr1 | 362452 |
| TEAD1 | 1.1423496 | 0.001182609 | TEF-1 | 361630 |
| TMCC3 | 0.6989743 | 0.001187061 | RGD1307241 | 314751 |
| CCDC82 | -0.55604905 | 0.001195598 | | 300359 |
| KCNH3 | -0.70602757 | 0.001198509 | Bec1\|Elk2 | 27150 |
| WSCD2 | -1.1164229 | 0.001199013 | Wscd2-ps1 | 360824 |
| EHD1 | -0.48566034 | 0.001199013 | RGD1306960 | 293692 |
| BNIP3 | -0.6796134 | 0.00120515 | | 84480 |
| STAMBP | -0.60814065 | 0.001211643 | Amsh | 171565 |
| PTPDC1 | 0.50092864 | 0.001211643 | | 291022 |
| ANXA7 | 0.4737687 | 0.001220179 | | 155423 |
| NPY2R | 1.174916 | 0.001229187 | | 66024 |
| TMEM106C | -0.50079596 | 0.001229815 | RGD1311532 | 315286 |
| BRINP3 | -0.8265199 | 0.001230632 | Fam5c | 286901 |
| PDCD6IP | 0.46702114 | 0.001230632 | AIP1\|Alix\|RGD1561176 | 501083 |
| NUDT18 | -0.6500521 | 0.001234377 | RGD1311802 | 361068 |
| SLC45A3 | -1.2641804 | 0.001236445 | RGD1309764 | 304785 |
| TRAPPC3 | -0.42669937 | 0.001236766 | | 362599 |
| HARS | 0.37955672 | 0.001241341 | Dnd1 | 307492 |
| SFPQ | 0.59208065 | 0.001242321 | | 252855 |
| SPRTN | 0.60773844 | 0.001254553 | RGD1559496 | 292101 |
| LRRC16B | -0.5078315 | 0.001264088 | Lrrc16b | 361041 |
| RAB9B | -0.56710654 | 0.001279833 | RGD1562951 | 367915 |
| OLFM4 | 0.9606124 | 0.001293636 | | 290409 |
| NUP93 | 0.45545802 | 0.001306387 | | 291874 |
| ING1 | 0.40621704 | 0.001317824 | p33ING1b\|p33ING1c | 306626 |
| KCNK3 | 0.85008407 | 0.001321135 | Task-1\|rTASK | 29553 |
| CYSRT1 | -1.1947086 | 0.001322035 | RGD1560880 | 499747 |

| | | | | |
|---|---|---|---|---|
| MMP15 | 0.5081275 | 0.00132987 | | 291848 |
| NXF1 | 0.5484511 | 0.001358358 | Mex67h\|Tap | 59087 |
| URI1 | -0.58499944 | 0.001370479 | RGD1310358 | 308537 |
| WDSUB1 | -0.56799155 | 0.001376847 | | 362137 |
| CEP295 | 0.6915539 | 0.001376847 | RGD1311723 | 363018 |
| PSTPIP2 | 0.85054135 | 0.001378867 | RGD1563090 | 307248 |
| FUCA2 | -0.77564716 | 0.001382232 | | 292485 |
| SULT2B1 | -0.75320095 | 0.001387261 | ST2B1 | 292915 |
| BCL2L13 | 0.55887043 | 0.001388408 | | 312682 |
| ASAP1 | 0.8193613 | 0.001390398 | Ddef1 | 314961 |
| EML5 | 0.90218073 | 0.001393962 | | 444982 |
| RNF6 | 0.40930188 | 0.00139603 | | 304271 |
| SERINC2 | 0.8296205 | 0.001399499 | Tde2l | 313057 |
| NAA25 | 0.5468156 | 0.001402575 | LRRGT00164\|Mdm20\|RGD13 | 360811 |
| AGK | -0.46472198 | 0.001407379 | RGD1562046 | 502749 |
| TOR1AIP1 | 0.45769244 | 0.001441091 | Lap1b\|Lap1c | 246314 |
| ANKRD13D | -0.5459929 | 0.001462133 | RGD1309156 | 361699 |
| DIRC2 | -0.5177993 | 0.001462133 | | 303902 |
| HPCAL4 | -0.65064806 | 0.001465978 | NVP-2\|Nvjp2 | 50872 |
| NPAP60 | 0.37747714 | 0.001465978 | Nup50\|Rtp60 | 25497 |
| TNIP1 | -0.5493757 | 0.001474619 | | 363599 |
| SPAST | -0.37951675 | 0.001488689 | Spg4 | 362700 |
| BCORL1 | -0.75904447 | 0.001489755 | RGD1566108 | 302810 |
| HMGCR | 0.49033245 | 0.001514448 | 3H3M | 25675 |
| MYLIP | 0.5107507 | 0.001516362 | MIR | 306825 |
| RNF112 | -0.5731631 | 0.001517352 | BFP\|Bfb\|Zfp179\|Znf179 | 24916 |
| PLA2G4A | 1.1080424 | 0.001519433 | Pla2c\|Pla2g4\|cPLA2 | 24653 |
| RRAGA | 0.5169129 | 0.00152748 | Raga | 117044 |
| 42796 | -0.45689023 | 0.00153112 | MARCH-II\|RGD1306395 | 362849 |
| FAM217B | -0.88913196 | 0.001552614 | RGD1311952 | 311692 |
| SYT3 | -0.7124081 | 0.00156439 | SIII | 25731 |
| ZFP672 | 0.51476556 | 0.00156439 | Znf672 | 303165 |
| IMPDH1 | -0.459075 | 0.001573219 | IMPD 1\|IMPDH 1 | 362329 |
| TVP23A | -0.81732684 | 0.001580614 | Fam18a\|RGD1566405 | 360467 |
| KIFAP3 | -0.34634274 | 0.001587511 | | 289168 |
| RAB27A | -0.86322165 | 0.001593067 | ram | 50645 |
| RBM18 | 0.33643258 | 0.001593067 | | 311902 |
| GLUL | 0.38647527 | 0.001595521 | Glns | 24957 |
| PXN | 0.6410701 | 0.001595521 | | 360820 |
| DEPDC1B | 0.5622565 | 0.001595577 | | 310074 |
| CYB561D2 | -0.57080746 | 0.001609138 | | 363137 |
| RGD1559896 | -0.34047088 | 0.001613119 | | 498967 |
| CD55 | -0.96448857 | 0.001613967 | Daf\|Daf1 | 64036 |

| | | | | |
|---|---|---|---|---|
| NDST3 | 1.1187375 | 0.001627725 | | 295430 |
| YPEL3 | -0.76707417 | 0.001634564 | RGD1564579 | 293491 |
| DIS3 | 0.6484738 | 0.001654481 | RGD1304646 | 306103 |
| PLXND1 | -0.66652536 | 0.001662025 | | 312652 |
| AHCYL2 | 0.5672737 | 0.001666295 | RGD1564895 | 312192 |
| LSM10 | -0.5304686 | 0.001670468 | | 366468 |
| ZFP523 | -0.43426538 | 0.001670468 | Znf76 | 361809 |
| BCL11A | -0.89537686 | 0.001678072 | | 305589 |
| CLK2 | 0.4075584 | 0.001678072 | | 365842 |
| GLI2 | 1.040282 | 0.001678072 | | 304729 |
| NCBP3 | 0.48621503 | 0.001684382 | RGD1308139 | 360563 |
| RAB3GAP2 | 0.56497294 | 0.001684382 | RGD1311518 | 289350 |
| ZFP295 | 0.62897 | 0.001685243 | Znf295 | 304056 |
| MAP3K7 | 0.41928765 | 0.001690861 | Tak1 | 313121 |
| NEUROD4 | -1.0769848 | 0.001694129 | | 288821 |
| SYT4 | 0.5085085 | 0.001725132 | | 64440 |
| FNDC3A | 0.80316854 | 0.001731631 | Fndc3 | 306022 |
| HTR7 | 0.8087953 | 0.001731631 | 5Ht7 | 65032 |
| TAF1A | 0.46075687 | 0.001742467 | | 360893 |
| ZMYND8 | -0.4993775 | 0.001744596 | Prkcbp1\|spikar | 296374 |
| RQCD1 | 0.35330376 | 0.001749681 | Rqcd1 | 301513 |
| LDAH | -0.34138876 | 0.001750941 | RGD1311648 | 313949 |
| CYP39A1 | -0.94485986 | 0.001756238 | | 301264 |
| ACPP | 1.3829361 | 0.001760973 | 5'-NT\|Acpp11\|FRAP\|Ppal\|RN | 56780 |
| AZIN1 | 0.5310784 | 0.001783718 | Oazi\|Oazin | 58961 |
| SEMA3C | 0.5066261 | 0.001806473 | | 296787 |
| CARNMT1 | 0.6264513 | 0.001806898 | RGD1311863 | 293871 |
| SLC27A3 | -0.43738768 | 0.001808057 | | 295219 |
| GOLGA7B | 0.6574997 | 0.001808057 | | 309378 |
| SIPA1L2 | 0.5657982 | 0.00181921 | Sersap2\|Spar2 | 361442 |
| APLF | -0.669483 | 0.001822101 | RGD1565557 | 500247 |
| NFIX | -0.61413676 | 0.001822101 | Nfix | 81524 |
| MRPL10 | -0.44270423 | 0.001824196 | L10mt\|MRP-L10\|MRP-L8 | 691075 |
| ZBTB24 | 0.49677444 | 0.001824461 | Bif1a\|ZNF450 | 365590 |
| SMARCD3 | -0.60978127 | 0.001829849 | | 296732 |
| GALK2 | -0.5638727 | 0.001834044 | | 296117 |
| SMARCA5 | 0.6474431 | 0.001834044 | | 307766 |
| TXNDC11 | 0.35675973 | 0.001842396 | RGD1307538 | 302899 |
| TBC1D9 | -0.5926496 | 0.001843215 | RGD1308221 | 304645 |
| TPH1 | 1.3470098 | 0.001868024 | Tph | 24848 |
| PNMA1 | -1.1105555 | 0.001874523 | MA1 | 170636 |
| JAKMIP1 | -0.7710171 | 0.001885091 | Marlin-1\|Marlin1 | 305434 |
| ARPP21 | 0.4453368 | 0.001891512 | Arpp-21\|Ppp1r1c\|Ppp1r1cl\|F | 363153 |

| | | | | |
|---|---|---|---|---|
| KCNMB2 | 0.95317215 | 0.001892463 | Kcmb2 | 294961 |
| MAP2K5 | -0.37010694 | 0.001892602 | Mek5 | 29568 |
| STC2 | 0.84162474 | 0.001892602 | | 63878 |
| NKIRAS2 | -0.44531527 | 0.00191021 | | 287707 |
| SLC20A1 | 0.6861232 | 0.00191021 | PiT-1 | 81826 |
| RNF128 | 0.62710315 | 0.001913067 | RGD1566282 | 315911 |
| PCDHA7 | -0.47689474 | 0.00191786 | rCNRv07 | 393089 |
| SPOP | -0.542691 | 0.00194976 | | 287643 |
| TRPC1 | -0.52421254 | 0.001959578 | Trrp1 | 89821 |
| MYO19 | 0.7242043 | 0.001980904 | RGD1559961 | 497974 |
| FBXO41 | -0.82166547 | 0.001982358 | RGD1566130 | 312504 |
| GALE | -0.53963566 | 0.001982854 | | 114860 |
| NAGA | -0.41248217 | 0.00198457 | | 315165 |
| KCND1 | -0.6402698 | 0.001986341 | Kv4.1 | 116695 |
| SERPINB8 | 1.3687408 | 0.002010378 | | 288937 |
| ZFP36 | 0.76764697 | 0.002015744 | Tis11 | 79426 |
| TBC1D8 | -0.4443501 | 0.00201678 | | 680133 |
| PDZD8 | 0.5937467 | 0.002042684 | Pdzk8 | 308000 |
| ALYREF | 0.62773097 | 0.002062709 | Thoc4 | 690585 |
| EXO1 | 0.76198125 | 0.002064636 | | 305000 |
| FADS6 | -0.6694298 | 0.002069547 | RGD1311950 | 303671 |
| PTP4A3 | -0.6268247 | 0.002074878 | | 362930 |
| PPEF1 | -0.81774217 | 0.002097266 | | 317498 |
| NEIL1 | -0.71396124 | 0.002099288 | | 367090 |
| PIK3R1 | -0.64042646 | 0.002099413 | PI3KA | 25513 |
| DARS2 | -0.48116833 | 0.002103762 | RGD1308286 | 304919 |
| MED9 | -0.56644887 | 0.002105041 | RGD1563669 | 497914 |
| BRIX1 | 0.48678407 | 0.002134174 | Bxdc2\|RGD1308508 | 294799 |
| ETNK2 | -0.7629403 | 0.002158593 | EKI 2 | 360843 |
| ADM | 0.98095083 | 0.002187922 | Ap\|H39316\|RATAP | 25026 |
| LOC688452 | 0.7606615 | 0.002189862 | Spaca6 | 688452 |
| SENP2 | 0.5625889 | 0.002194873 | Axam | 78973 |
| CHAF1B | 0.49415588 | 0.002200997 | | 288242 |
| API5 | 0.39791232 | 0.002205337 | | 362170 |
| RUNX1T1 | -0.83150077 | 0.002217144 | Cbfa2t1 | 362489 |
| CARS2 | -0.49256566 | 0.002218968 | RGD1311612 | 361184 |
| SIN3A | -0.43387488 | 0.002218968 | | 363067 |
| MBLAC2 | -0.6480122 | 0.002231585 | RGD1306703 | 365627 |
| CAT | -0.39946076 | 0.002231585 | CS1\|Cas1\|Cat01\|Catl\|Cs-1 | 24248 |
| ESR1 | 1.1407906 | 0.002235185 | ER-alpha\|Esr\|RNESTROR | 24890 |
| DDX51 | 0.43235165 | 0.002315611 | | 304570 |
| SFRP4 | 1.0897659 | 0.002315611 | | 89803 |
| NDRG4 | -0.48663014 | 0.002326731 | Bdm1\|Ndr4\|smap8 | 64457 |

| | | | | |
|---|---|---|---|---|
| INIP | 0.49384215 | 0.002329927 | RGD1307983 | 298032 |
| FSCN3 | 1.01676 | 0.002329927 | | 296947 |
| GRIP2 | -0.678192 | 0.002331911 | Abp | 171571 |
| IGF1 | 1.2108998 | 0.002331911 | | 24482 |
| XKR4 | -0.59722596 | 0.002337273 | XRG4 | 297801 |
| VAT1 | -0.4805402 | 0.00235633 | | 287721 |
| ST6GALNAC6 | -0.46399876 | 0.00235633 | Siat7F | 407765 |
| SLC44A2 | -0.44919485 | 0.00235633 | RGD1309680 | 363024 |
| SLC24A3 | 0.5326149 | 0.00235633 | Nckx3 | 85267 |
| ORC1 | 0.9550075 | 0.00235633 | Orc1l | 313479 |
| NDST4 | -1.3208786 | 0.00235929 | | 362035 |
| RNF219 | 0.35777017 | 0.002367857 | RGD1564706 | 361088 |
| MCM4 | 0.64696366 | 0.002370721 | Mcmd4 | 29728 |
| PCDHA4 | -0.38769412 | 0.002405588 | Cnr1 | 116741 |
| RIT1 | -0.36586645 | 0.00246091 | RGD1559874 | 499652 |
| RIIAD1 | 1.0830668 | 0.002471164 | | 100270669 |
| ZMYND10 | -0.7166203 | 0.002474723 | | 363139 |
| MYBBP1A | 0.6370758 | 0.002480002 | DBP | 60571 |
| GTF3C5 | -0.38180616 | 0.002491373 | | 362095 |
| ACTL6A | 0.40495035 | 0.002514606 | RGD1307747 | 361925 |
| NKX3-1 | 1.2755096 | 0.002515555 | | 305999 |
| DISP2 | -0.6134049 | 0.002556581 | | 311324 |
| ARFGAP2 | 0.383457 | 0.002566636 | Zfp289 | 362162 |
| NDEL1 | 0.4134144 | 0.002567926 | NUDE2\|Nudel | 170845 |
| SPRED1 | 0.8288227 | 0.002567926 | | 296072 |
| PELI3 | -0.749179 | 0.002570655 | RGD1305989 | 309157 |
| CDO1 | -0.601678 | 0.002577847 | | 81718 |
| SLC25A32 | 0.49625838 | 0.002578091 | RGD1565789 | 315023 |
| SLC6A8 | 0.570552 | 0.002578091 | CHOT1\|CHT1\|CRT\|CT1 | 50690 |
| ROR2 | 0.6379546 | 0.002579488 | | 306782 |
| NFXL1 | 0.7204357 | 0.002600309 | RGD1359201 | 289595 |
| NPLOC4 | 0.5224384 | 0.002606698 | Npl4 | 140639 |
| SLC25A42 | -0.4402367 | 0.002609801 | | 689414 |
| MFAP2 | -0.44396982 | 0.002639372 | | 313662 |
| TTC19 | 0.5083155 | 0.002639372 | RGD1311797 | 691506 |
| GPATCH2 | 0.6533226 | 0.002639372 | Gpatc2 | 289362 |
| LEMD3 | 0.40933073 | 0.002642271 | RGD1561001 | 680066 |
| RFXAP | 0.61888963 | 0.002650466 | | 499617 |
| MCOLN3 | 1.2846128 | 0.002662769 | | 308022 |
| PLPPR4 | 0.4323068 | 0.002678223 | Lppr4\|Prg-1\|Prg1 | 295401 |
| VPS37D | -0.67426366 | 0.002682377 | | 687208 |
| ASUN | 0.42108116 | 0.002704072 | | 690728 |
| PDK2 | -0.44153845 | 0.002713821 | | 81530 |

| | | | | |
|---|---|---|---|---|
| ZFP395 | -0.49914 | 0.002722246 | | 305972 |
| CLDN3 | -1.327775 | 0.002735947 | | 65130 |
| SV2C | 1.3235396 | 0.002735947 | | 29643 |
| MAFF | 0.6750922 | 0.002736387 | | 366960 |
| FKBP1A | 0.31115192 | 0.002739557 | FKBP12\|Fkbp2 | 25639 |
| ITPR3 | 1.1131233 | 0.002747432 | IP3R3\|IP3R3X | 25679 |
| CYP11B3 | 1.2890716 | 0.002758251 | CYPXIB3 | 353498 |
| ZBTB16 | -1.1543972 | 0.002766361 | Lx\|Plzf\|Zfp145\|Znf145 | 353227 |
| BPHL | -0.57253987 | 0.002774489 | | 361239 |
| HNRNPF | 0.6241387 | 0.002774495 | Hnrpf | 64200 |
| ZFP18 | 0.42917475 | 0.002779537 | Zfp535\|Zkscan6\|Znf18 | 303226 |
| HPN | 1.3208531 | 0.002788 | | 29135 |
| B4GALNT4 | -0.4399508 | 0.002813124 | RGD1310052 | 309105 |
| HERC3 | -0.5670623 | 0.002856638 | | 362377 |
| SLC30A10 | 0.7927553 | 0.002856638 | RGD1305098 | 289353 |
| SLC35B4 | -0.37633476 | 0.002863239 | | 296969 |
| ATAD3A | 0.32929096 | 0.002863239 | Atad3 | 298682 |
| TBCK | -0.3988576 | 0.002872885 | RGD1307816\|Tbckl | 295446 |
| NOL3 | -0.4864958 | 0.002873274 | Arc | 85383 |
| NIFK | 0.5546444 | 0.002931919 | Mki67ip | 246042 |
| OPRK1 | -1.2661084 | 0.002945226 | | 29335 |
| LRRC75B | -0.89601713 | 0.002945226 | Fam211b | 100365744 |
| TBC1D25 | -0.5088058 | 0.002945226 | Oatl1\|RGD1559711 | 302552 |
| EEA1 | 0.5295778 | 0.002949305 | | 314764 |
| BRINP1 | 0.58867437 | 0.002951721 | Dbc1\|Dbccr1 | 140610 |
| SPTLC1 | 0.39000693 | 0.00295267 | Lcb1\|RGD1306617\|Spt1 | 361213 |
| ARL4D | 0.7306127 | 0.00295267 | Arf4l | 303559 |
| HGS | 0.36550793 | 0.002959091 | Hrs | 56084 |
| FAM26F | 1.3035073 | 0.002999965 | RGD1304835 | 294430 |
| ANO6 | 0.74228007 | 0.003006419 | Tmem16f | 315272 |
| LOC361346 | 0.6765565 | 0.003009195 | Las2 | 361346 |
| RIMS4 | 0.9382637 | 0.003045795 | Rim4gamma | 266976 |
| PSMB11 | 1.047436 | 0.003048026 | RGD1308773 | 290206 |
| SEC24B | 0.36643368 | 0.003058924 | | 295461 |
| NMBR | 1.0635699 | 0.00308967 | NMB-R | 25264 |
| HNRNPU | 0.44230294 | 0.003123493 | Hnrpu\|SN1 | 117280 |
| RBPJ | 0.64910966 | 0.003143098 | | 679028 |
| WDR6 | -0.50963014 | 0.003184359 | | 301007 |
| SLC12A6 | -0.7204348 | 0.00319381 | Kcc3 | 691209 |
| CIR1 | -0.8313745 | 0.003194235 | 1700023b02rik\|Cir\|RGD1309 | 362149 |
| ATRAID | -0.5178097 | 0.003194644 | RGD1311605 | 298841 |
| DCAKD | -0.30305466 | 0.003211073 | | 360639 |
| DND1 | 1.0508786 | 0.003215087 | | 679841 |

| | | | | |
|---|---|---|---|---|
| LCK | -0.7098316 | 0.003231213 | Lck1\|Lcktkr | 313050 |
| NR1D1 | -0.4220045 | 0.003231213 | REV-ERBAALPHA | 252917 |
| RSRP1 | -0.39317167 | 0.003245802 | RGD1359529 | 362626 |
| FBL | 0.6051908 | 0.003247209 | | 292747 |
| BBS2 | -0.5848547 | 0.003267711 | | 113948 |
| ZFP639 | 0.4032127 | 0.003267711 | Znf639 | 683504 |
| MRPS2 | 0.35226396 | 0.003272506 | | 362094 |
| CDK18 | -1.0914624 | 0.003274151 | PCTAIRE3\|Pctk3 | 289019 |
| FAM222B | 0.5397364 | 0.003284344 | RGD1566149 | 497960 |
| DYNC1I1 | -0.30571437 | 0.003284492 | Dnci1\|Dncic1 | 29564 |
| ENGASE | -0.6188538 | 0.003294754 | RGD1310868 | 303702 |
| PCBP4 | -0.31813145 | 0.003294754 | | 363133 |
| CHSY1 | 0.3295991 | 0.003300143 | | 292999 |
| SCN4A | -1.1988186 | 0.003331173 | NCHVS\|Nav1.4\|microI | 25722 |
| ZFP800 | 0.69300103 | 0.003331173 | RGD1560157 | 500057 |
| TMEM87B | 0.45328468 | 0.003332206 | RGD1307907 | 362212 |
| PPP1R10 | 0.43675312 | 0.003333753 | Fb19\|Pnuts | 65045 |
| ZFYVE19 | -0.47516018 | 0.003340885 | | 499871 |
| FN3K | -1.18426 | 0.003346953 | Fnsk | 498034 |
| MYCN | -0.46250296 | 0.003346953 | N-myc\|Nmuc1\|Nmyc | 298894 |
| MON2 | 0.66772544 | 0.003356609 | RGD1308808 | 314894 |
| EMILIN3 | 1.0215374 | 0.003366859 | | 362262 |
| SLC30A2 | -1.2455347 | 0.003374921 | ZnT-2\|Znt2 | 25362 |
| PURG | 0.5889565 | 0.003383831 | | 361162 |
| PTGIR | 1.2722929 | 0.003413173 | | 292661 |
| FSTL5 | -0.35508668 | 0.003419268 | | 365823 |
| FCHO1 | -0.44358185 | 0.003452883 | | 290639 |
| CHMP1A | -0.41559383 | 0.003452883 | Pcoln3 | 365024 |
| EMP3 | 1.2313905 | 0.003452986 | | 81505 |
| RPS6KA6 | 0.47431427 | 0.00346415 | RGD1560817 | 317203 |
| NPAS2 | 0.5455637 | 0.00346415 | | 316351 |
| SHD | -0.56852055 | 0.00346793 | | 316507 |
| ELOVL4 | 0.3928217 | 0.003474993 | | 315851 |
| HMGCL | -0.47724962 | 0.003477131 | | 79238 |
| JUP | -0.6522347 | 0.003483449 | | 81679 |
| MFN1 | 0.42748147 | 0.003500277 | Fzo1b | 192647 |
| SLC2A8 | -0.49465877 | 0.003501701 | Glut8 | 85256 |
| ZDHHC20 | 0.5142524 | 0.003510928 | RGD1305755 | 305923 |
| CCDC32 | -0.6085224 | 0.003520041 | Gm631 | 296081 |
| ZPBP2 | -1.0296645 | 0.003535947 | | 363676 |
| FADD | 0.8071368 | 0.003547048 | Mort1 | 266610 |
| FBXO30 | -0.5166577 | 0.003554638 | | 308283 |
| MTPAP | 0.4852335 | 0.003593015 | Papd1 | 307050 |

| | | | | |
|---|---|---|---|---|
| FOXP2 | -0.6970624 | 0.003596837 | RGD1559697 | 500037 |
| LOC100125367 | 0.70325047 | 0.003596837 | C1orf74 | 100125367 |
| OSCAR | 1.1101156 | 0.003609305 | RGD1559897 | 292537 |
| KCNA4 | 0.64054894 | 0.003625348 | KCHAN\|Kv1.4\|Kv4\|RHK1\|RK3 | 25469 |
| FAM129C | -0.9176881 | 0.00363949 | | 498604 |
| NAT14 | -0.5717621 | 0.00363949 | RGD1565991\|Zfp628 | 361500 |
| SMIM17 | -0.68858564 | 0.003641049 | RGD1559613 | 499067 |
| AKAP9 | -0.52254903 | 0.003653498 | CG-NAP\|Gisp\|Yotiao | 246150 |
| ORC2 | 0.6520922 | 0.003659118 | Orc2l | 301430 |
| BCKDHA | -0.4117469 | 0.003673959 | BCKDA\|E1a | 25244 |
| RNF139 | 0.37509334 | 0.003691356 | TRC8 | 315000 |
| DNAAF5 | 0.6442604 | 0.003704938 | Heatr2 | 304332 |
| APBB3 | -0.45330966 | 0.003707689 | Fe65l2 | 117026 |
| TYMS | 0.58960056 | 0.003712491 | | 29261 |
| PXYLP1 | -0.49903175 | 0.003724332 | Acpl2 | 315939 |
| DTX2 | 0.49222773 | 0.003763813 | | 304591 |
| ZFAND6 | -0.3773252 | 0.003771226 | Za20d3\|zgc:101121 | 293067 |
| REEP2 | -0.35044065 | 0.003773972 | | 682105 |
| N4BP3 | -0.7732296 | 0.003786323 | C330016o10rik\|RGD1308356 | 303112 |
| CCDC28B | -0.6469193 | 0.003786323 | | 682445 |
| RPS6KB2 | 0.38501665 | 0.003795693 | | 361696 |
| RGS14 | -0.69307476 | 0.003797186 | | 114705 |
| PUS7 | 0.67190504 | 0.003797186 | RGD1307054 | 296751 |
| RANGAP1 | 0.41167435 | 0.003804426 | | 362965 |
| LRCH1 | 0.58920455 | 0.003804426 | | 502020 |
| TENM3 | -0.6950083 | 0.003808736 | Odz3 | 306451 |
| TFAP4 | -0.67507505 | 0.003827042 | Tcfap4 | 360482 |
| TBK1 | 0.40808332 | 0.003830434 | | 299827 |
| TMEM223 | -0.5691301 | 0.003839845 | | 690285 |
| IFT81 | -0.6005608 | 0.003870222 | Cdv-1\|Cdv1 | 373066 |
| ITSN1 | 0.6095012 | 0.003875809 | Itsn | 29491 |
| FOXO3 | -0.89672095 | 0.003884692 | Fkhrl1\|Foxo3a | 294515 |
| PPP4R3A | 0.38124272 | 0.003885117 | RGD1309059\|Smek1 | 314388 |
| CSRP2 | -0.6035509 | 0.003912169 | SmLIM | 29317 |
| RGD1308147 | 0.37867263 | 0.003940911 | | 307008 |
| WNT9A | -0.85681874 | 0.003952992 | | 287357 |
| POLR3E | 0.34231988 | 0.003956293 | RGD1308086 | 361640 |
| TBC1D14 | -0.4229025 | 0.003971899 | SRF-2\|UR-NR#2 | 360956 |
| NCS1 | 0.49619532 | 0.003984428 | Freq | 65153 |
| KTI12 | 0.5730082 | 0.003994137 | | 685656 |
| NRSN2 | -0.8531885 | 0.003997315 | | 689978 |
| CISD2 | 0.42032087 | 0.004040704 | RGD1566242 | 295457 |
| RHOQ | 0.6162784 | 0.004042533 | Tc10 | 85428 |

| | | | | |
|---|---|---|---|---|
| SDE2 | 0.3299553 | 0.004044133 | RGD1305572 | 289315 |
| PHOSPHO2 | 0.3698227 | 0.004055211 | | 295663 |
| BNIP3L | -0.42774191 | 0.004081985 | Nix | 140923 |
| RBBP7 | 0.4944776 | 0.004090612 | | 83712 |
| GRIN3B | 0.5486226 | 0.004097416 | GluN3B\|NMDAR3B\|NR3B | 170796 |
| GEMIN5 | 0.62930375 | 0.004097416 | | 691231 |
| CDC16 | 0.28580996 | 0.004109758 | | 290875 |
| GPATCH11 | 0.39006427 | 0.004110932 | Ccdc75\|RGD1311424 | 362685 |
| SLC5A4 | -1.2244954 | 0.00411264 | Slc5a4a | 294341 |
| RGD1564804 | -0.6460024 | 0.004115207 | | 313551 |
| FN3KRP | -0.46802858 | 0.004123667 | RGD1304570 | 303755 |
| UBE2F | 0.41229182 | 0.004123667 | RGD1307608 | 363284 |
| AMZ2 | -0.40870693 | 0.004130729 | RGD1304846 | 360650 |
| SLC9B2 | -0.7989277 | 0.004135514 | Nhedc2 | 679958 |
| SUN1 | 0.59840167 | 0.004141931 | Unc84a | 360773 |
| SLC25A39 | -0.4051778 | 0.004168978 | RGD1306193 | 360636 |
| ATF1 | 0.51113796 | 0.004168978 | Atf-1 | 315305 |
| SPATA5L1 | 0.5796718 | 0.004168978 | | 691729 |
| IDH1 | -0.4401083 | 0.004169217 | | 24479 |
| UBE2D3 | 0.3849438 | 0.004222871 | | 81920 |
| EML1 | -0.66444993 | 0.004228776 | | 362783 |
| PRPF18 | -0.37852702 | 0.004251674 | KCRF\|Pprf18 | 171552 |
| ACSL1 | 0.34920642 | 0.004251674 | ACS\|Acas\|COAA\|Facl2 | 25288 |
| LAMC1 | 0.7354476 | 0.004274136 | B2e | 117036 |
| NAPEPLD | 0.77145904 | 0.004277008 | NAPE-PLD | 296757 |
| DDX52 | 0.5531235 | 0.004299835 | Rok1 | 85432 |
| ULK2 | -0.33573863 | 0.004315719 | | 303206 |
| SH3GLB1 | 0.4935133 | 0.004339478 | | 292156 |
| CEBPG | 0.5350657 | 0.004339478 | C/EBP\|CEBPRNA | 25301 |
| ABHD8 | -0.7104634 | 0.004342752 | | 306338 |
| SYT6 | 0.79413813 | 0.004346998 | | 60565 |
| RECK | 0.7344724 | 0.004355526 | | 313488 |
| MPLKIP | 0.4674159 | 0.004360512 | | 684996 |
| PCDHA10 | -0.61110485 | 0.004377054 | rCNRv10 | 116778 |
| ELK3 | 0.551439 | 0.004377054 | | 362871 |
| NEPRO | 0.6178104 | 0.004377054 | RGD1311458 | 303948 |
| SLC5A3 | 1.0699618 | 0.004377054 | Smit | 114507 |
| TUBB4B | 0.28440395 | 0.004377083 | Tubb2\|Tubb2c | 296554 |
| PLEKHA4 | -0.593785 | 0.004413514 | | 308584 |
| MCMBP | 0.49211955 | 0.004427683 | MCM-BP\|RGD1306730 | 309009 |
| TJP3 | -0.85026395 | 0.00444724 | | 314640 |
| UXS1 | 0.37348494 | 0.004454902 | UGD | 246232 |
| VWC2 | 0.865909 | 0.004454902 | | 501231 |

| | | | | |
|---|---|---|---|---|
| MEPCE | -0.39203325 | 0.004465452 | | 304361 |
| CSTF2 | 0.29538757 | 0.004491416 | | 683927 |
| FUNDC1 | -0.5906039 | 0.004509994 | | 363442 |
| CUL3 | 0.36666733 | 0.004511394 | | 301555 |
| ALKBH1 | 0.41555202 | 0.004555387 | Alkbh | 362766 |
| CEP85L | 0.7712387 | 0.004556595 | | 100365935 |
| THBS2 | 1.1592374 | 0.004579172 | TSP-2 | 292406 |
| CCDC71 | -0.5524986 | 0.0045851 | | 498678 |
| PRTN3 | -0.91662306 | 0.004593573 | | 314615 |
| TUBA4A | 0.42560843 | 0.004598877 | Tuba4 | 316531 |
| TSPAN17 | -0.51210874 | 0.004600765 | Fbxo23\|GHB-R | 306771 |
| TMEM202 | 1.0555767 | 0.004626515 | | 691306 |
| RHOA | 0.49570227 | 0.004637632 | Arha\|Arha2 | 117273 |
| GNE | 0.51726013 | 0.004663155 | Uae1 | 114711 |
| ALKBH2 | -0.8614533 | 0.004668592 | RGD1306377 | 304578 |
| KDM4A | -0.49764067 | 0.004688337 | Jmjd2a | 313539 |
| HNRNPH1 | 0.46152624 | 0.004688713 | Hnrph\|Hnrph1 | 140931 |
| PAX1 | 1.1155308 | 0.00469122 | Pax-1 | 311505 |
| TUBG1 | -0.47636124 | 0.004773877 | | 252921 |
| NFYA | 0.48423356 | 0.004798219 | CBF-A | 29508 |
| PRKAA1 | 0.3957278 | 0.004844106 | AMPKalpha1 | 65248 |
| POLD1 | 0.49632335 | 0.004845094 | | 59294 |
| CNNM2 | -0.65561235 | 0.004852129 | | 294014 |
| AKAP5 | -0.8019565 | 0.00488042 | AKAP150\|Akap79\|P150 | 171026 |
| RNF145 | -0.44215265 | 0.004890411 | RGD1309561 | 287212 |
| COMMD7 | -0.49026376 | 0.004907491 | | 296285 |
| GAR1 | 0.56295294 | 0.004907491 | Nola1 | 499709 |
| NTRK3 | 0.7086573 | 0.004907491 | trkC | 29613 |
| SMYD5 | -0.3803803 | 0.004916756 | | 312503 |
| PCDHA5 | -0.5710929 | 0.004936068 | rCNRv05 | 393087 |
| FIG4 | -0.4024622 | 0.004944686 | Kiaa0274\|RGD1311375\|Sac3 | 309855 |
| 42802 | -0.4099952 | 0.00495365 | Mir | 312656 |
| MZF1 | -0.408548 | 0.004959129 | Zfp98 | 361508 |
| PPP1R1A | 0.6349888 | 0.004966796 | | 58977 |
| PEAK1 | 1.2385699 | 0.00496826 | RGD1312026 | 315686 |
| RNF167 | -0.35393977 | 0.00497668 | RGD1305972 | 360554 |
| PPP3CC | -0.38419008 | 0.004986884 | | 171378 |
| TRPC4 | -0.5223144 | 0.005011922 | Trp4\|Trrp4 | 84494 |
| NAP1L3 | -0.72873604 | 0.005018402 | | 170914 |
| CHST10 | 0.52101976 | 0.005029481 | Hnk-1st | 140568 |
| RPAP2 | -0.4148779 | 0.005081292 | RGD1309034 | 305120 |
| F2R | 0.36301935 | 0.005097748 | Par1\|TRGPC | 25439 |
| TTC14 | 0.639218 | 0.005097748 | | 310314 |

| | | | | |
|---|---|---|---|---|
| GAS7 | -0.5019012 | 0.00510623 | | 85246 |
| SDK2 | 1.196949 | 0.005116664 | | 360652 |
| SPDYA | -0.76542056 | 0.005124109 | Gs4\|Lm23\|Spdy1\|Spy1 | 192209 |
| BOK | -0.701874 | 0.005136154 | Bok-BH3 | 29884 |
| EMC9 | -0.8998065 | 0.005136166 | Fam158a\|RGD1308113 | 290224 |
| TMPO | 0.32697394 | 0.005138164 | LAP2 | 25359 |
| KCNK1 | 0.36667544 | 0.005139695 | Twik | 59324 |
| TERF1 | 0.5834361 | 0.005147464 | | 297758 |
| TGIF1 | 0.6937128 | 0.005147464 | Tfig\|Tgif | 316742 |
| HPGD | 1.1811209 | 0.005147464 | | 79242 |
| FJX1 | 0.5376544 | 0.005156112 | | 366140 |
| CDC25B | -0.48590094 | 0.005168988 | | 171103 |
| SVOP | -0.60297906 | 0.005182482 | | 171442 |
| HEMK1 | -0.5954756 | 0.005182482 | RGD1308293 | 300989 |
| ZDHHC5 | 0.6428169 | 0.005216695 | DHHC-5 | 362156 |
| RCOR1 | 0.44634447 | 0.005250085 | RGD1305743 | 102554884 |
| IL1RL2 | 1.1775132 | 0.005268157 | | 171106 |
| NASP | 0.58709365 | 0.005275937 | | 298441 |
| FBXL6 | -0.5333863 | 0.005339098 | | 362941 |
| MED8 | 0.45743755 | 0.005354017 | | 362575 |
| SOBP | -0.631317 | 0.005394171 | RGD1560479\|Sobpl\|jcx1 | 309860 |
| LOC361635 | -0.35061666 | 0.005394371 | | 361635 |
| CDCA4 | 0.38979167 | 0.005422936 | | 500727 |
| FZR1 | -0.509077 | 0.005477336 | | 314642 |
| MELTF | -1.0656476 | 0.005504429 | Magea1\|Mfi2 | 288038 |
| FAM220A | -0.41932482 | 0.005504429 | Sipar | 498145 |
| CCDC93 | 0.5447762 | 0.005507648 | | 304743 |
| TFDP1 | 0.34993988 | 0.005546859 | | 361178 |
| GLRX | -0.48387155 | 0.005596032 | Glrx1\|Grx | 64045 |
| KIF3B | -0.31642193 | 0.005596032 | | 296284 |
| ASCC3 | 0.49893677 | 0.005650464 | Helic1 | 309887 |
| PPFIA4 | -0.76504517 | 0.005665115 | | 140592 |
| ISG20L2 | 0.7826963 | 0.005676423 | | 361977 |
| TNRC6B | -0.6648719 | 0.005709778 | Cbl27 | 192178 |
| TNFRSF25 | 1.0643183 | 0.005733798 | | 500592 |
| TMEM141 | -0.6813307 | 0.005735843 | RGD1561492 | 499755 |
| WNT4 | 0.8826062 | 0.005740115 | | 84426 |
| CLN5 | 0.60114914 | 0.005746855 | | 306128 |
| WDR3 | 0.3375209 | 0.005779792 | | 310720 |
| ABTB1 | -0.6536297 | 0.005859064 | | 297432 |
| ERI2 | 0.51821464 | 0.005869466 | Exod1 | 691484 |
| PIGM | 0.56453097 | 0.005895754 | GPI-MT-I | 79112 |
| CORO1C | 0.5309538 | 0.005907807 | RGD1564490 | 501841 |

| | | | | |
|---|---|---|---|---|
| PKDCC | 0.49999848 | 0.005909331 | RGD1311939\|Sgk493 | 313860 |
| RGD1307315 | -0.4464015 | 0.00591617 | | 362793 |
| MSANTD3 | 0.42342773 | 0.005918904 | RGD1308165 | 362516 |
| EPHA2 | 1.0286369 | 0.005919148 | | 366492 |
| FBXO31 | -0.6204082 | 0.005923453 | RGD1561069 | 498959 |
| MAGED2 | -0.35522595 | 0.005938329 | | 113947 |
| AKAP12 | 0.4540927 | 0.005938329 | AKAP12A\|AKAP12B\|AKAP12( | 83425 |
| ACVR1B | -0.52200156 | 0.005945652 | Skr2 | 29381 |
| CNTN3 | -0.86453986 | 0.005964303 | BIG-1\|Pang | 54279 |
| YPEL2 | -0.9454961 | 0.00598012 | RGD1311564 | 360590 |
| MSL3L2 | 0.7059998 | 0.005988806 | RGD1308699 | 309790 |
| PRR13 | -0.40907007 | 0.006000093 | RGD1307129 | 363004 |
| FAM129B | 0.54152995 | 0.006012125 | RGD1307018 | 362115 |
| NR2F6 | 0.46301594 | 0.006019252 | | 245980 |
| SLC35F5 | -0.36386016 | 0.006063604 | | 288993 |
| NOV | -0.5798898 | 0.006083168 | | 81526 |
| ASIC1 | -0.42842707 | 0.006100501 | Accn2 | 79123 |
| SVBP | -0.8070299 | 0.006107451 | Ccdc23\|RGD1311232 | 362578 |
| NABP1 | 0.7346499 | 0.006111566 | Obfc2a\|RGD1306658 | 363227 |
| TMEM205 | -0.6299742 | 0.006129767 | RGD1563250 | 300441 |
| NAT10 | 0.37204736 | 0.006133545 | RGD1306717 | 311257 |
| CANT1 | -0.5045032 | 0.006144266 | srapy | 246272 |
| RNF180 | -0.37024853 | 0.006163667 | | 685384 |
| CISD3 | -0.6918637 | 0.006166389 | Mel13\|RGD1559720 | 287661 |
| IPO7 | 0.39745855 | 0.006166389 | | 308939 |
| DFNB31 | 0.5800239 | 0.006166389 | Cip98\|Whrn | 313255 |
| PACSIN2 | 0.63012505 | 0.006166389 | SdpII | 124461 |
| EPRS | 0.5938935 | 0.006239232 | | 289352 |
| AP4B1 | -0.3710514 | 0.006244707 | | 310746 |
| TRANK1 | 0.74175256 | 0.006244707 | RGD1563130 | 316022 |
| AGFG2 | -0.5693147 | 0.006261388 | Hrbl | 304375 |
| SAMHD1 | 0.49654856 | 0.006312627 | | 311580 |
| SAMD5 | 0.8477639 | 0.006338316 | RGD1311012 | 365038 |
| LIG4 | 0.66022676 | 0.006354985 | | 290907 |
| LRRFIP1 | 0.3652048 | 0.006383461 | | 367314 |
| RAB2B | -0.3584162 | 0.006421351 | | 305853 |
| TAF2 | 0.30742243 | 0.006444161 | TAFIIB | 170844 |
| SYT12 | 0.75857925 | 0.006454528 | Srg1 | 191595 |
| ATG4C | -0.42357612 | 0.006478442 | | 313391 |
| VGLL4 | 0.35816586 | 0.006478442 | | 297523 |
| AQP11 | -0.61925507 | 0.006514886 | | 286758 |
| NARS | 0.44664752 | 0.006528283 | LRRGT00113 | 291556 |
| IGFBPL1 | -0.6693971 | 0.006588765 | | 366366 |

| | | | | |
|---|---|---|---|---|
| INO80C | -0.36147293 | 0.006594931 | RGD1310199 | 291737 |
| TACR1 | -1.0150286 | 0.006642334 | Tac1r | 24807 |
| LENG8 | 0.5379686 | 0.006642334 | | 361506 |
| FRAS1 | 1.1956491 | 0.006642334 | | 289486 |
| NRAP | -0.94512296 | 0.006742248 | | 307982 |
| RAI2 | -0.4963332 | 0.006761554 | RGD1560139 | 501555 |
| ATP5S | -0.8233238 | 0.006810316 | | 362749 |
| CNR1 | -0.59843177 | 0.006857776 | SKR6R | 25248 |
| FGF8 | 1.2183199 | 0.00687262 | | 29349 |
| PEX14 | -0.43377188 | 0.006914046 | | 64460 |
| CDC42EP2 | -0.8827879 | 0.006927021 | | 309175 |
| KCTD15 | -0.39976555 | 0.006962657 | | 499129 |
| BAMBI | -0.64160264 | 0.006993972 | | 83837 |
| TIMP4 | 0.61468256 | 0.006993972 | | 680130 |
| TARBP2 | -0.3733423 | 0.007031335 | | 363006 |
| HMG20A | -0.3441888 | 0.007074592 | RGD1564760 | 315689 |
| TMEM114 | 1.1656243 | 0.007099612 | RGD1560593 | 501675 |
| KCNE2 | -0.7082734 | 0.007102726 | Mirp1 | 171138 |
| BEND5 | -0.6119894 | 0.007102726 | | 362564 |
| MMS19 | -0.3799196 | 0.007102726 | Mms19l | 171124 |
| PARD6B | 0.4408882 | 0.007102726 | | 362279 |
| RGD1310769 | -0.47779393 | 0.007161525 | | 299207 |
| RBAK | 0.40330344 | 0.007161525 | RGD1562875 | 288489 |
| WDR4 | 0.66925246 | 0.007161525 | | 690032 |
| NECAB3 | -0.8259294 | 0.007169013 | Apba2bp | 311562 |
| MRPL34 | -0.7356181 | 0.007183372 | | 290632 |
| DTX3 | -0.41688272 | 0.00719039 | RGD1566181 | 500847 |
| FRY | -0.8267952 | 0.007203703 | RGD1307034 | 304244 |
| LDB1 | -0.3872721 | 0.007323738 | | 309447 |
| NR0B1 | 1.0694661 | 0.007323738 | Ahch\|DAX-1 | 58850 |
| DEAF1 | -0.55294883 | 0.007333743 | NUDR | 83632 |
| TARDBP | 0.3015849 | 0.007337708 | Tdp-43 | 298648 |
| ATG10 | -0.7369567 | 0.007339263 | | 688555 |
| ARL4A | 0.5981951 | 0.007339263 | Arl4 | 29308 |
| TDRD6 | -1.1833401 | 0.007401314 | | 316254 |
| TMEM106B | -0.30082932 | 0.007401314 | LRRGT00101 | 312132 |
| PCDH7 | 0.76379704 | 0.007401314 | | 360942 |
| CDK5RAP1 | -0.53716445 | 0.007436976 | | 252827 |
| SNRPA | 0.3253762 | 0.007436976 | | 292729 |
| SLC43A1 | 0.850067 | 0.007438363 | | 311168 |
| TMEM178A | 0.5657237 | 0.007500956 | Tmem178 | 362691 |
| TES | 0.5907334 | 0.007521784 | | 500040 |
| TRAPPC4 | -0.64285696 | 0.007616847 | Sbdn | 367073 |

| | | | | |
|---|---|---|---|---|
| TOP1MT | -0.59850335 | 0.007637879 | | 300029 |
| CBLN1 | -0.61230785 | 0.007648109 | | 498922 |
| HIBCH | -0.42652848 | 0.007669291 | | 301384 |
| CRY2 | 0.55243105 | 0.00776325 | | 170917 |
| CBLN4 | -0.76772314 | 0.007824593 | | 499947 |
| FRA10AC1 | -0.6739591 | 0.007824593 | RGD1309482 | 365458 |
| LRRC7 | 0.74293244 | 0.007852638 | | 117284 |
| NHLH2 | -0.99175644 | 0.007871 | | 295327 |
| PHYHD1 | 0.6154232 | 0.007871 | | 296621 |
| ATXN2L | 0.67690605 | 0.007871434 | RGD1565868 | 361649 |
| WDR5 | 0.3201417 | 0.007872803 | | 362093 |
| VARS2 | -0.37360045 | 0.007873848 | Vars2l\|Varsl | 309596 |
| HRH1 | 0.7724292 | 0.007873848 | Hisr | 24448 |
| PLXNA2 | 0.79055667 | 0.007896097 | | 289392 |
| RNGTT | -0.30786905 | 0.007917888 | | 313131 |
| ATG13 | -0.5128015 | 0.007935271 | Harbi1\|Harbi1l\|RGD1310685 | 362164 |
| DEF6 | -1.148882 | 0.007943125 | | 309642 |
| SLC9A5 | 0.49069187 | 0.007967993 | Nhe5 | 192215 |
| TMEM55B | -0.39198777 | 0.008072576 | RGD1307475 | 364298 |
| SARS | 0.39547387 | 0.008072576 | Sars1 | 266975 |
| OLIG2 | 0.36800122 | 0.008075546 | | 304103 |
| BCOR | 0.44199145 | 0.008079262 | RGD1562735 | 317346 |
| MFSD3 | -0.58512545 | 0.008091276 | | 500899 |
| IWS1 | 0.46569702 | 0.008118515 | RGD1304762 | 291705 |
| ST8SIA4 | -0.7798197 | 0.008123681 | Siat8d | 116696 |
| RBM46 | -0.7862903 | 0.008132804 | RGD1560205 | 310548 |
| RGD1311188 | 0.5532214 | 0.008146983 | | 315088 |
| CYR61 | 0.95583934 | 0.008146983 | | 83476 |
| SGMS1 | 0.44758293 | 0.008148881 | Mob\|Tmem23 | 353229 |
| SLCO1C1 | 0.6127385 | 0.008177418 | Bsat1\|Oatp14\|Slc21a14 | 84511 |
| DPF1 | -0.58634126 | 0.008184612 | Neud4 | 50545 |
| RIOK3 | 0.3015605 | 0.008250138 | | 361293 |
| PTPRH | -0.9853315 | 0.008254619 | Bem2 | 171125 |
| DUOX1 | 1.0812548 | 0.008254619 | | 266807 |
| RGS9 | -0.7473781 | 0.008289885 | | 29481 |
| COQ10A | -0.4284515 | 0.008343717 | RGD1562447 | 362810 |
| ANKEF1 | 1.2020665 | 0.008350425 | Ankrd5 | 296184 |
| PJA2 | -0.35182375 | 0.008373673 | Neurodap1\|praja2 | 192256 |
| ST3GAL2 | -0.54038936 | 0.008385118 | Siat4b\|Siat5 | 64442 |
| C1QL1 | 0.43491355 | 0.008396974 | | 363686 |
| SMOX | 0.34669903 | 0.008435047 | | 308652 |
| PIH1D1 | -0.47758472 | 0.008503411 | RGD1309809 | 292898 |
| WLS | 0.37792286 | 0.008532126 | Gpr177 | 362065 |

| | | | | |
|---|---|---|---|---|
| TMF1 | 0.63926774 | 0.008532126 | | 114206 |
| TTC33 | -0.42918146 | 0.008553218 | RGD1564042 | 294774 |
| KEAP1 | -0.48125336 | 0.008556129 | Inrf2 | 117519 |
| RASGRF1 | -0.44856116 | 0.008556129 | | 192213 |
| KPNB1 | 0.56883645 | 0.008556129 | Impnb | 24917 |
| CUL4A | -0.3229395 | 0.008583327 | RGD1563853 | 361181 |
| GALNT3 | -0.77247727 | 0.008656305 | | 366061 |
| PRPSAP2 | -0.40071207 | 0.008660885 | Pap41 | 117272 |
| CCDC40 | -0.65257496 | 0.008794177 | | 287867 |
| CFAP97 | -0.57325876 | 0.008794177 | RGD1307325 | 306469 |
| NFKB1 | 0.59376323 | 0.008794177 | EBP-1\|NF-kB | 81736 |
| ZFP189 | 0.6014152 | 0.008794177 | | 313219 |
| EFNA5 | 0.8253024 | 0.008794177 | Lerk7 | 116683 |
| IL11 | -0.8465593 | 0.008835731 | Il-11 | 171040 |
| BANP | -0.35008252 | 0.008862481 | | 292064 |
| FSCN1 | 0.5502662 | 0.008885613 | Fascin | 683788 |
| ELAC2 | 0.37857094 | 0.00893575 | | 282826 |
| LMBR1L | -0.5199525 | 0.009033621 | RGD1309631 | 300215 |
| HERPUD2 | -0.53365934 | 0.009096939 | RGD1307343 | 300463 |
| MAPK1IP1 | -0.50829536 | 0.009096939 | RGD1565431 | 499280 |
| GNL3 | 0.5454993 | 0.009097274 | Ns | 290556 |
| KCNK13 | 1.0015669 | 0.009097274 | THIK-1\|prdx1 | 64120 |
| KLF13 | -0.5957496 | 0.009102651 | RGD1565099 | 499171 |
| LZTFL1 | -0.57112455 | 0.009116273 | | 316102 |
| XKR7 | 1.15487 | 0.009132666 | | 311549 |
| BTG4 | -1.0542605 | 0.009133044 | SCIR-27 | 315650 |
| FLOT1 | -0.57301366 | 0.009232132 | Rareg | 64665 |
| WDR77 | 0.35151115 | 0.009239938 | Ac2-269\|MEP-50\|RGD131047 | 310769 |
| NPY | 0.94925565 | 0.009372319 | NPY02\|RATNPY\|RATNPY02 | 24604 |
| PPP1R1B | 0.43043384 | 0.009427398 | Darpp-32\|Darpp32 | 360616 |
| ELF3 | -1.1831183 | 0.009427593 | | 304815 |
| KATNB1 | -0.4570654 | 0.009427593 | | 291852 |
| SGCA | 0.9726473 | 0.009429377 | | 303468 |
| ZC3H11A | 0.73905 | 0.009568936 | RGD1308290 | 360845 |
| KCNK5 | 0.898048 | 0.009568936 | TASK2 | 364241 |
| CDK8 | 0.34470317 | 0.009589051 | RGD1560888 | 498140 |
| ASCC1 | -0.4827921 | 0.009596734 | Asc-1\|CGI-18 | 294512 |
| CFB | 0.99190015 | 0.009596734 | Bf\|Da1-24 | 294257 |
| CDK2AP2 | -0.44704607 | 0.009633761 | | 688405 |
| USP13 | 0.3351111 | 0.009750959 | | 310306 |
| BZW1 | 0.4769006 | 0.009750959 | | 363232 |
| RASA1 | 0.46877527 | 0.009786127 | GAPX\|Rasa | 25676 |
| MAK16 | 0.60697174 | 0.0098085 | RGD1311297\|Rbm13 | 306526 |

| | | | | |
|---|---|---|---|---|
| INTS12 | 0.40871048 | 0.009844609 | Phf22 | 295448 |
| RAB27B | 0.7839486 | 0.009873074 | | 84590 |
| MSRB1 | -0.37079212 | 0.00987861 | Sepx1\|selX | 685059 |
| MAP3K12 | -0.36958146 | 0.009950466 | DLK\|MUK\|PK\|Zpk | 25579 |
| PDE3B | 0.55044544 | 0.009950466 | | 29516 |
| VAV3 | 0.66234994 | 0.009977018 | RGD1565941 | 295378 |
| TMPRSS6 | 1.0566071 | 0.009998485 | | 315388 |
| HSCB | -0.53802866 | 0.010015708 | RGD1311005 | 360826 |
| TTC39C | 0.5109129 | 0.010026858 | | 686179 |
| NOTCH4 | -0.5388284 | 0.010029635 | | 406162 |
| RPS6KA3 | 0.7044436 | 0.010052121 | RGD1563860 | 501560 |
| GNA11 | 0.5249587 | 0.010080943 | | 81662 |
| PVRL1 | 0.6996072 | 0.010245699 | HveC\|Pvrl1\|nectin-1 | 192183 |
| PBX1 | -0.7391616 | 0.010258959 | | 304947 |
| TBP | 0.43738207 | 0.010327858 | TFIID | 117526 |
| TMEM229B | -0.5321619 | 0.010432375 | RGD1562622 | 503035 |
| HINT3 | -0.47867116 | 0.010432375 | HINT-3\|HINT-4\|Hint4 | 246769 |
| MET | 0.7387908 | 0.01045371 | Hgfr | 24553 |
| HBP1 | -0.462579 | 0.010578232 | Hmgb1 | 27080 |
| BMS1 | 0.4483293 | 0.01058447 | Bms1l | 362426 |
| POLDIP2 | -0.2770717 | 0.01068762 | | 287544 |
| CDK12 | 0.72352934 | 0.01068762 | Crk7\|Crkrs\|Pksc | 192350 |
| GLDN | 0.7637601 | 0.010722259 | Colm\|Crgl2 | 315675 |
| SDHC | -0.27536422 | 0.010724688 | | 289217 |
| AFAP1L1 | 0.7564462 | 0.010724688 | RGD1311580 | 291565 |
| BFAR | -0.32794145 | 0.010738542 | | 304709 |
| KMT5A | 0.734331 | 0.010748866 | Pr-set7\|RGD1305893\|Setd8 | 689820 |
| ERBB3 | 0.7934279 | 0.010816959 | nuc-ErbB3 | 29496 |
| RAB43 | 0.8154377 | 0.010816959 | | 500249 |
| MYBL1 | 0.7658934 | 0.010897748 | | 297783 |
| JAKMIP3 | -0.56649476 | 0.010920779 | Necc2\|RGD1307177 | 365380 |
| PHRF1 | 0.45779696 | 0.010951469 | | 245925 |
| MRVI1 | 1.1131667 | 0.010966362 | | 308899 |
| COMTD1 | -0.38202608 | 0.010984688 | | 305685 |
| RIOK2 | 0.51462376 | 0.010996142 | | 308201 |
| PCGF5 | 0.7161094 | 0.011064585 | | 681178 |
| PRRT2 | -0.5121309 | 0.011066943 | DSPB3\|RGD1564195 | 361651 |
| HDAC9 | -0.72362775 | 0.011070985 | RGD1310748\|RGD1563092 | 687001 |
| XRCC5 | -0.4033135 | 0.011080731 | Ku80\|Kup80 | 363247 |
| HPCAL1 | -0.5157591 | 0.011088899 | NVP-3\|Nvp3\|VILIP-3 | 50871 |
| BMF | -0.83336586 | 0.011220546 | | 246142 |
| DDX39A | 0.40229562 | 0.011228486 | Ddx39\|Ddxl | 89827 |
| LMO4 | -0.76822114 | 0.011260265 | | 362051 |

| | | | | |
|---|---|---|---|---|
| DDX56 | 0.33297938 | 0.011277332 | Ddx21\|RH-II/Gu | 289780 |
| PTBP1 | 0.5902062 | 0.01129836 | PTB1\|Ptb\|Pybp\|Pybp1\|Pybp[illegible] | 29497 |
| METTL18 | 0.68845814 | 0.011364142 | RGD1306783 | 304928 |
| GOLPH3L | -0.35808986 | 0.011370461 | | 310669 |
| SRPRA | 0.31458172 | 0.011376195 | Srpr | 315548 |
| TRPC3 | -0.5177324 | 0.011378234 | TrpC3c\|Trrp3 | 60395 |
| ANKRD34C | 1.1576787 | 0.011426149 | RGD1305403 | 300889 |
| CCDC148 | -0.6401252 | 0.011467871 | RGD1561169 | 311051 |
| G3BP1 | 0.29962832 | 0.011478704 | G3bp\|RGD1310666 | 171092 |
| HSPB11 | -0.645847 | 0.011492399 | | 685284 |
| SLC19A2 | 0.44552222 | 0.011543294 | | 289175 |
| TRMT12 | 0.47362167 | 0.011628785 | RGD1307127 | 314999 |
| GRIP1 | 0.5774581 | 0.011628785 | | 84016 |
| FNDC1 | 0.82207674 | 0.011628785 | | 308099 |
| SLC39A6 | 0.297998 | 0.011633094 | | 291733 |
| DCAF10 | 0.5334191 | 0.011633094 | RGD1304730\|Wdr32 | 313242 |
| DDI2 | 0.62694544 | 0.011633094 | RGD1311815 | 313668 |
| BANK1 | -1.1293828 | 0.01166602 | LRRGT00104 | 365948 |
| PNKD | -0.36677125 | 0.011666717 | Mr-1 | 100188944 |
| SNX16 | 0.35181302 | 0.011668363 | | 64088 |
| LOC363337 | 0.9969346 | 0.011669235 | | 363337 |
| ANAPC7 | -0.29959714 | 0.011677975 | | 304490 |
| CDV3 | 0.5922145 | 0.011718756 | | 315970 |
| ARHGAP25 | 0.9579346 | 0.011726267 | RGD1562105 | 500246 |
| PLOD3 | 0.3947694 | 0.011803632 | | 288583 |
| TEKT2 | -0.93230605 | 0.011809778 | | 298532 |
| SRRM3 | -0.4555189 | 0.011809778 | RGD1307391 | 685890 |
| MOSPD3 | -0.51183414 | 0.011891845 | | 288557 |
| RAD51 | 0.44940814 | 0.01191205 | RGD1563603 | 499870 |
| PARD6A | -0.63022274 | 0.011914798 | Par-6a\|Par6a | 307799 |
| SMIM20 | -0.5709944 | 0.011914798 | RGD1565192 | 501923 |
| GTF2IRD1 | -0.46814457 | 0.011914798 | Gtf3 | 246770 |
| ARHGEF7 | 0.43864736 | 0.011914798 | P85spr\|Pak3bp | 114559 |
| LFNG | 0.44454327 | 0.011976814 | | 170905 |
| RGD1309028 | -0.6242827 | 0.012039031 | | 299265 |
| CASP9 | -0.4441282 | 0.012155866 | Apaf3\|Casp-9-CTD\|Casp9_v1 | 58918 |
| ADAP1 | -0.4007203 | 0.012155866 | Centa1\|p42IP4 | 171097 |
| PI16 | -0.7531638 | 0.012169292 | | 294312 |
| TCEAL1 | -0.6499034 | 0.012176254 | | 302593 |
| OGFOD1 | 0.5635196 | 0.012176254 | RGD1308848 | 307657 |
| SEMA6B | -0.58345705 | 0.012255387 | | 84609 |
| ZFP9 | -0.5530713 | 0.012271681 | | 100158232 |
| PAFAH1B3 | -0.60000294 | 0.012316116 | | 114113 |

| | | | | |
|---|---|---|---|---|
| PTP4A2 | 0.5444053 | 0.012316116 | | 85237 |
| CLCN5 | 0.99063647 | 0.012317729 | CLC5 | 25749 |
| CRAT | -0.43266946 | 0.012332224 | | 311849 |
| NUDT16 | -0.59488744 | 0.012413353 | RGD1311387 | 363129 |
| CELF6 | -0.35775325 | 0.012423826 | Brunol6 | 300758 |
| CDK2 | 0.42351413 | 0.012472671 | | 362817 |
| FAM227A | -0.5646193 | 0.012541516 | RGD1305939 | 300074 |
| RAD18 | 0.48092374 | 0.012632669 | | 362412 |
| HSPA14 | 0.34657094 | 0.012634094 | | 307133 |
| PPP1R12A | 0.4900083 | 0.012656978 | M110\|MBSP\|Mypt1 | 116670 |
| ABTB2 | 0.52522504 | 0.012657382 | Cca3 | 171440 |
| E2F3 | 0.6646341 | 0.012657382 | RGD1561600 | 291105 |
| U2AF1 | 0.47243112 | 0.012676724 | | 687575 |
| SLC7A5 | 0.64414984 | 0.01268377 | E16\|TA1 | 50719 |
| ALKBH4 | 0.4706152 | 0.012723504 | RGD1308608 | 288587 |
| B3GALT5 | -0.60520667 | 0.012782929 | | 288161 |
| FEM1A | 0.43934175 | 0.012786375 | | 316131 |
| IRGQ | -0.4388204 | 0.012798666 | RGD1309409 | 292708 |
| HPS4 | -0.36125737 | 0.012807782 | | 304555 |
| PIK3CB | 0.6236102 | 0.012816889 | | 85243 |
| ASCC2 | 0.56986994 | 0.012851208 | RGD1561422 | 498402 |
| WDR45 | -0.4638789 | 0.012886619 | | 302559 |
| PRKCQ | -0.4221288 | 0.01289057 | Pkcq | 85420 |
| ZFP575 | -0.6800261 | 0.012934943 | Znf575 | 308430 |
| OSBPL9 | -0.42365465 | 0.012934943 | | 298369 |
| FADS3 | -0.3418549 | 0.012934943 | | 286922 |
| CDKN2AIPNL | -0.51119804 | 0.012948143 | RGD1308696 | 287278 |
| EME1 | 0.70008147 | 0.01298084 | | 287634 |
| GRK5 | 0.5016199 | 0.012997693 | Gprk5 | 59075 |
| LCMT2 | 0.6007042 | 0.013089576 | | 296098 |
| FAM134A | -0.4706435 | 0.013092566 | RGD1306844 | 363252 |
| ARF1 | 0.2535262 | 0.013098046 | | 64310 |
| SEC13 | 0.2940295 | 0.013098046 | Sec13l1 | 297522 |
| KIFC3 | 0.33618397 | 0.013098096 | KRP4 | 307644 |
| UROD | -0.39947385 | 0.013158204 | porphyrinogen carboxy-lyase | 29421 |
| RCAN1 | 0.3098915 | 0.013201669 | Dscr1\|Mcip1 | 266766 |
| PSENEN | -0.3733969 | 0.013263653 | RGD1312037 | 292788 |
| KIF18B | 0.54401064 | 0.013345053 | RGD1310360 | 303575 |
| MEOX2 | 1.1383067 | 0.013349704 | | 29279 |
| CMTR2 | 0.40433457 | 0.013375053 | Ftsjd1\|RGD1309394 | 292016 |
| TMA16 | 0.66073817 | 0.013386441 | RGD1305222 | 290686 |
| KCNJ5 | 1.139306 | 0.01343561 | | 29713 |
| PDXP | -0.627077 | 0.013486668 | Rbp1 | 727679 |

| | | | | |
|---|---|---|---|---|
| GABARAPL1 | -0.3361119 | 0.013505544 | Gec1 | 689161 |
| FNTB | -0.32460427 | 0.013505544 | | 64511 |
| CRIPT | -0.5646176 | 0.013523571 | | 56725 |
| FUT7 | 0.94434845 | 0.013527854 | | 296564 |
| SAPCD2 | 0.5823559 | 0.013559467 | | 680531 |
| PLK4 | 0.45452306 | 0.013645521 | | 310344 |
| WHSC1 | 0.56271935 | 0.013645521 | RGD1565590 | 680537 |
| RAPGEF3 | 0.41137132 | 0.013665983 | Epac | 59326 |
| ITGA11 | -1.0088171 | 0.013686501 | | 315744 |
| NUDT22 | -0.7193156 | 0.013696576 | | 293703 |
| E2F6 | 0.47217578 | 0.013747766 | | 313978 |
| CCDC175 | -0.68771666 | 0.013760201 | | 500668 |
| TRPT1 | -0.62095404 | 0.013778904 | Tpt1h | 293704 |
| MAD2L1BP | 0.46765026 | 0.013778904 | | 316237 |
| CYTH1 | -0.42518106 | 0.013819262 | Pscd1\|Sec7 | 116691 |
| MAPK3 | -0.44389716 | 0.013960122 | ERK1\|ERT2\|Erk-1\|Esrk1\|MAP | 50689 |
| SRP68 | 0.2553731 | 0.013960122 | | 363707 |
| PNKP | 0.41672245 | 0.013967045 | | 308576 |
| APBA2 | 0.44258118 | 0.013969976 | Mint2 | 83610 |
| NCOA4 | 0.2536262 | 0.01399342 | | 619385 |
| DVL1 | 0.41346362 | 0.014030753 | dvl-1 | 83721 |
| CEP104 | 0.44551 | 0.014032714 | GlyBP | 246295 |
| ZFP35 | 0.46781194 | 0.014046456 | Zfp239 | 307547 |
| TAS1R3 | 1.1216946 | 0.014046456 | T1r3 | 170634 |
| CNNM1 | 0.64462084 | 0.0140548 | | 309387 |
| CROT | 0.31769052 | 0.014072188 | | 83842 |
| CLDN9 | -1.0229357 | 0.014098884 | | 287099 |
| VKORC1 | -0.584093 | 0.014123564 | | 309004 |
| ZFAND2B | -0.54397416 | 0.014211214 | RGD1306260 | 363253 |
| MGAT1 | -0.33278877 | 0.014421033 | | 81519 |
| POPDC3 | 0.64929414 | 0.014451542 | | 641520 |
| TPD52L3 | -1.1269943 | 0.014482369 | RGD1309391\|Trpd52l3 | 293894 |
| SLC9A3R1 | 0.2791779 | 0.01448429 | | 59114 |
| ZFP358 | -0.40169713 | 0.01453809 | | 360754 |
| ELOVL5 | 0.6144908 | 0.014565824 | rELO1 | 171400 |
| SLC22A23 | 0.546605 | 0.014620495 | Nritp | 64559 |
| WBSCR17 | 0.59746975 | 0.014620495 | | 288611 |
| GRIN2A | 1.1237993 | 0.014670977 | GluN2A\|NMDAR2A\|NR2A | 24409 |
| EGLN3 | -0.36376014 | 0.014675131 | PHD-3\|PHD3\|SM-20 | 54702 |
| RGD1307443 | -0.6906255 | 0.014710051 | | 361244 |
| SLC35A5 | -0.42048854 | 0.014710051 | RGD1564361 | 498081 |
| RSPO3 | -0.7410595 | 0.014804967 | RGD1563246 | 498997 |
| USP1 | 0.47711465 | 0.01482081 | | 313387 |

| | | | | |
|---|---|---|---|---|
| CELF1 | 0.6000399 | 0.014868277 | Cugbp1 | 362160 |
| HSPA12A | -0.4305887 | 0.014875834 | | 307997 |
| RTFDC1 | -0.34047946 | 0.014875834 | RGD1311072 | 296410 |
| GZF1 | 0.31087232 | 0.014882887 | RGD1562321\|Zfp336 | 311508 |
| ITGA7 | 0.5440397 | 0.014901846 | | 81008 |
| ZFP414 | -0.53686273 | 0.01492048 | RGD1308123\|Znf414 | 299647 |
| EIF1B | -0.46571234 | 0.014938674 | RGD1309241 | 301068 |
| POLR1A | 0.60301477 | 0.014951176 | Rpa1\|Rpo1-4 | 83581 |
| ENTPD6 | -0.42570895 | 0.014967268 | | 85260 |
| RGD1565498 | 0.47882587 | 0.015042431 | | 500843 |
| KBTBD2 | 0.3364676 | 0.015217288 | | 312372 |
| LRRC42 | -0.41950768 | 0.015220032 | RGD1308919 | 298309 |
| MGAT5B | -0.5500894 | 0.015225536 | | 303693 |
| LRRC55 | -0.77066547 | 0.015227393 | | 311171 |
| PAPD7 | 0.287108 | 0.015227393 | Pols | 306672 |
| COL26A1 | -0.7256622 | 0.015306952 | | 685612 |
| MCCC1 | -0.42266458 | 0.015306952 | | 294972 |
| ST6GAL1 | 0.30966005 | 0.015314262 | Siat1 | 25197 |
| FSTL4 | 0.6717412 | 0.015316743 | | 303130 |
| TMEM45A | 1.1240699 | 0.015316743 | | 680866 |
| HIBADH | -0.38129097 | 0.01535139 | | 63938 |
| RGD1304694 | 0.4424138 | 0.01535139 | | 362974 |
| ZFP865 | -0.4434425 | 0.015435516 | RGD1565545\|Znf865 | 308337 |
| C2CD5 | -0.28698078 | 0.015568304 | RGD1304592 | 362461 |
| KAT8 | -0.3544251 | 0.015594526 | Myst1 | 310194 |
| INO80 | 0.46175486 | 0.015594526 | Inoc1\|RGD1310969 | 296084 |
| CTTN | 0.32078534 | 0.015613903 | Cttnb | 60465 |
| UBAP2 | 0.53915226 | 0.015616591 | | 313169 |
| PRKRIP1 | 0.42274854 | 0.015644388 | | 498171 |
| TMEM42 | -0.65324134 | 0.015668258 | RGD1307118 | 363171 |
| LOC688765 | 0.57121044 | 0.015714934 | | 688765 |
| FEZ1 | -0.38018176 | 0.015769731 | | 81730 |
| MRPL14 | -0.63503325 | 0.015814753 | L14mt\|L32mt\|MRP-L14\|MRF | 301250 |
| PEO1 | -0.37689152 | 0.015904427 | | 309441 |
| FAM127B | -0.40364155 | 0.015928643 | Cxx1a | 679038 |
| SLC27A2 | -0.83193815 | 0.015976228 | | 65192 |
| GRB10 | -0.3867585 | 0.016015625 | RGD1566234 | 498416 |
| SLC44A1 | 0.42136672 | 0.016015625 | Cdw92\|Ctl1 | 85254 |
| GRP | -0.75931096 | 0.016039979 | | 171101 |
| LRRC6 | -0.97423357 | 0.016069578 | Lrtp\|Tslrp | 299920 |
| BRE | -0.40150228 | 0.016130334 | | 362704 |
| SLC22A1 | -1.0284437 | 0.016288156 | Oct1\|Orct1\|Roct1 | 24904 |
| ISPD | -0.5942794 | 0.016308328 | Nip | 493574 |

| | | | | |
|---|---|---|---|---|
| TSPYL1 | -0.30987325 | 0.016322302 | Tspyl | 29544 |
| PITPNB | 0.24328303 | 0.016322302 | | 114561 |
| EPHA5 | 0.42673656 | 0.016352832 | EHK-1\|Els1\|Els1. | 79208 |
| NOP58 | 0.49382052 | 0.016380163 | Nap65\|Nol5 | 60373 |
| SPON1 | 0.5561997 | 0.016466217 | Sponf | 64456 |
| ATPAF2 | -0.41099223 | 0.016475318 | | 303190 |
| POLE4 | -0.47191355 | 0.016498253 | | 362385 |
| ZC3H14 | 0.34251946 | 0.016498253 | Npuk68 | 192359 |
| DIS3L2 | -0.39656162 | 0.01651732 | RGD1560168 | 367307 |
| TMEM39A | 0.46733102 | 0.016577259 | RGD1306421 | 288092 |
| RNF125 | -0.96201056 | 0.016589224 | | 361296 |
| ATP6V0A2 | 0.57361895 | 0.016614597 | Cc1-3\|J6b7\|Tj6 | 116455 |
| HIGD2A | -0.31916255 | 0.01662287 | RGD1309691 | 290999 |
| TMEM184B | -0.44063416 | 0.016624494 | RGD1306591 | 362959 |
| AMN | -0.93529457 | 0.016706992 | | 314459 |
| DOC2B | 0.7099712 | 0.016706992 | | 81820 |
| SATB1 | -0.48735073 | 0.016751138 | | 316164 |
| CIAPIN1 | 0.42386988 | 0.01675911 | | 307649 |
| USP15 | 0.44951677 | 0.016768282 | Ubp109 | 171329 |
| ZFP93 | -0.88612896 | 0.01677114 | | 296399 |
| TBX22 | -1.0988843 | 0.016847761 | Tbx 22 | 302369 |
| PCDHA9 | -0.42499796 | 0.016847761 | rCNRv09 | 393090 |
| SLC3A2 | 0.46668822 | 0.016847761 | Mdu1 | 50567 |
| BTRC | 0.47048524 | 0.016847761 | beta-TrCP1 | 361765 |
| HSPA9 | 0.6106112 | 0.016955905 | Crp40\|GRP-75\|Hspa9a\|PBP7 | 291671 |
| LINGO3 | 0.6689029 | 0.01696763 | | 690755 |
| GRN | -0.40805992 | 0.017058002 | PGRN | 29143 |
| MFSD13A | -0.43792814 | 0.017107178 | RGD1309313\|Tmem180 | 309454 |
| TAL2 | 0.94439465 | 0.017111102 | Tail2 | 685229 |
| ADARB1 | 0.49465048 | 0.017120646 | Adar2\|Red1 | 25367 |
| USP47 | 0.29100683 | 0.017180504 | | 308896 |
| OSCP1 | -0.37596297 | 0.017206836 | RGD1306596 | 362595 |
| RGD1565059 | -0.7956027 | 0.017226197 | | 499630 |
| GLYCTK | -0.54001683 | 0.017226197 | | 684314 |
| LOC100911367 | 0.5705844 | 0.017255178 | | 100911367 |
| ADAT1 | 0.4908129 | 0.017425861 | | 690810 |
| PTPRE | 0.55275923 | 0.017460024 | PTPepsilon | 114767 |
| CSTF1 | 0.33187506 | 0.017541908 | | 311670 |
| PEX13 | 0.3088094 | 0.01754587 | | 305581 |
| KIF19 | -0.71489835 | 0.017548606 | Kif19a\|RGD1559936 | 303659 |
| STRAP | 0.30872402 | 0.017723577 | | 297699 |
| AHRR | -1.0712992 | 0.01773828 | | 498999 |
| STK4 | 0.39856547 | 0.017745363 | | 311622 |

| | | | | |
|---|---|---|---|---|
| CACNB3 | -0.37787104 | 0.017808218 | CACH3B | 25297 |
| RHBDL1 | -0.7174056 | 0.017820828 | Rhbdl | 117025 |
| GPR160 | -1.0051844 | 0.017873589 | | 499588 |
| LIAS | -0.52962303 | 0.017964575 | | 305348 |
| TMEM107 | -0.5602709 | 0.017983124 | | 691750 |
| RNF19B | -0.3713631 | 0.018190706 | Ibrdc3 | 313806 |
| MTERF4 | 0.3561925 | 0.018190706 | Mterfd2 | 363289 |
| STRN3 | 0.31938872 | 0.018196741 | Gs2na\|Sg2na | 114520 |
| VOPP1 | -0.31295136 | 0.01823828 | Ecop\|RGD1306494 | 362374 |
| SEC22A | -0.5121607 | 0.018246332 | Sec22l2 | 117513 |
| PTRH2 | 0.3744117 | 0.018246332 | RGD1306819 | 287593 |
| ZFP259 | 0.2686922 | 0.018263027 | RGD1562173 | 500989 |
| MYH7 | -0.7258306 | 0.018271018 | Bmyo\|Myhcb\|myHC-beta\|m | 29557 |
| TSC22D2 | 0.5230815 | 0.018339287 | | 499624 |
| WDR55 | 0.50927746 | 0.018367928 | LRRG00133\|RGD1305640 | 307494 |
| ANKMY2 | -0.5344849 | 0.018450173 | Gna14 | 314046 |
| SCARF1 | 1.0565767 | 0.018450173 | | 303313 |
| ZBTB20 | -0.70726675 | 0.018489016 | RGD1560387 | 288105 |
| NT5C1A | -0.8567769 | 0.018526126 | | 313574 |
| KCNJ9 | -0.45441687 | 0.018526126 | Girk3\|Kir3.3 | 116560 |
| RAB11FIP2 | -0.3372077 | 0.018526126 | RGD1308538 | 308003 |
| PBX4 | -0.6258097 | 0.01855699 | Edg4 | 361131 |
| ABHD14A | -0.7208101 | 0.0185698 | Dorz1\|RGD1309721 | 300982 |
| FARS2 | -0.53872687 | 0.018574912 | Fars1 | 306879 |
| GPR61 | -0.5982247 | 0.01859059 | | 310780 |
| BROX | -0.3832211 | 0.018613735 | RGD1307161 | 305031 |
| ZCCHC8 | 0.4474043 | 0.01883635 | | 288661 |
| HRK | -0.4555785 | 0.018928798 | Bid3\|Dp5 | 117271 |
| USP11 | -0.30351844 | 0.018964652 | Uhx1 | 408217 |
| PRSS36 | -0.6603448 | 0.019085756 | | 497040 |
| KLHDC1 | -0.65808123 | 0.019107081 | | 314190 |
| AAK1 | 0.53069514 | 0.019107081 | RGD1563580 | 500244 |
| PDGFC | 0.74214494 | 0.019112902 | SCDGF | 79429 |
| SURF2 | 0.75827086 | 0.019112902 | | 619345 |
| STIM1 | -0.3825069 | 0.019153882 | | 361618 |
| ZCCHC24 | 0.30016264 | 0.019153882 | RGD1306164 | 361104 |
| PPP1R15B | 0.42969123 | 0.01926163 | | 304799 |
| CBX7 | -0.637661 | 0.019262733 | | 362962 |
| CCM2 | 0.3330669 | 0.019262733 | | 305505 |
| ASB8 | -0.46658117 | 0.019325513 | | 315287 |
| SEC23B | 0.46436992 | 0.019342592 | | 362226 |
| FGF22 | 0.56972426 | 0.019345097 | | 170579 |
| CAMSAP3 | -0.42870077 | 0.019371033 | RGD1307246 | 689074 |

| | | | | |
|---|---|---|---|---|
| ATRIP | 0.3553003 | 0.019394368 | | 301014 |
| CPSF7 | 0.39920464 | 0.019409502 | RGD1305441 | 365407 |
| METRNL | 0.83298993 | 0.019446347 | | 316842 |
| VEZT | 0.27029166 | 0.019518334 | | 299738 |
| SFTPA1 | 1.0221486 | 0.019572165 | PSAP\|PSP-A\|SP-A\|Sftp1\|Sftp | 24773 |
| TRIM3 | -0.3747128 | 0.019587457 | Berp\|Rnf22 | 83616 |
| PGAM5 | 0.47645056 | 0.019648103 | RGD1312028 | 288731 |
| RNF34 | -0.35055918 | 0.01968367 | Momo | 282845 |
| BMP7 | 0.5152237 | 0.01968367 | BMP-7 | 85272 |
| IDI1 | 0.5555658 | 0.01968367 | | 89784 |
| SPTLC2 | 0.5762459 | 0.01968367 | Pomt2 | 366697 |
| SCNN1G | 1.0740247 | 0.01968367 | ENaC | 24768 |
| LARS | 0.5295304 | 0.0197218 | | 291624 |
| EBF1 | -1.0893587 | 0.019734237 | Ebf\|Olf1 | 116543 |
| ZFP503 | 0.55792856 | 0.019744331 | Nolz-1\|Nolz1\|Znf503 | 305687 |
| NTNG2 | 0.64086926 | 0.01978038 | | 311836 |
| CCNL2 | 0.6622199 | 0.01978038 | | 298686 |
| PKNOX2 | -0.5545568 | 0.019788744 | | 680549 |
| FBXO3 | -0.28655007 | 0.019788744 | | 690634 |
| PSME4 | 0.48247126 | 0.01982289 | | 498433 |
| GNB5 | -0.48535332 | 0.019877074 | | 83579 |
| BRAP | 0.31765878 | 0.01990787 | | 687346 |
| RAB36 | -0.407907 | 0.019967712 | | 690407 |
| FAM163B | 0.5645095 | 0.019967712 | | 685169 |
| F8 | 0.5986479 | 0.019967712 | | 302470 |
| PWWP2B | 0.70017093 | 0.019967712 | Pwwp2 | 361671 |
| GPR155 | -0.38577795 | 0.020069456 | | 311730 |
| TNIP2 | 0.6872239 | 0.020082528 | | 305451 |
| APBB1 | -0.36937773 | 0.02017821 | FE65 | 29722 |
| ALG9 | -0.32408738 | 0.020257102 | Dibd1 | 367083 |
| KDELR3 | 0.44304436 | 0.020277565 | | 315131 |
| MESDC1 | 0.28909972 | 0.020294696 | | 308795 |
| SEPHS1 | 0.33469778 | 0.02044995 | | 291314 |
| AIG1 | -0.42043447 | 0.020462526 | RGD1562920 | 292486 |
| CCDC91 | -0.60597724 | 0.0204778 | Ccdc91 | 103690009 |
| CALY | -0.7370509 | 0.02062924 | Calcyon\|Drd1ip | 192349 |
| FIP1L1 | 0.42561775 | 0.02062924 | | 289582 |
| GJB2 | 0.87944615 | 0.02066671 | CXN-26\|Cx26 | 394266 |
| SNRNP40 | 0.26904786 | 0.020728698 | RGD1309198 | 313056 |
| MPDU1 | -0.49394405 | 0.020758046 | | 303244 |
| ZMPSTE24 | 0.4096436 | 0.020809803 | | 313564 |
| LYRM7 | -0.7201525 | 0.020845149 | | 686506 |
| DHX33 | 0.6332344 | 0.021010872 | | 287464 |

| | | | | |
|---|---|---|---|---|
| MAP2K1 | 0.46555167 | 0.021087652 | Mek1 | 170851 |
| TRAK1 | -0.44992325 | 0.021176055 | RGD1307844 | 316085 |
| HCN3 | -0.58350736 | 0.021250414 | | 114245 |
| COX11 | 0.3311598 | 0.021282801 | | 690300 |
| DECR2 | -0.7600805 | 0.021355916 | Dcrakl\|putativeperoxisomal2 | 64461 |
| NEMP2 | 0.3647779 | 0.021356642 | RGD1560421\|Tmem194b | 503257 |
| MPHOSPH10 | 0.49548197 | 0.021419972 | | 293828 |
| FAM212A | -0.98108214 | 0.021421747 | RGD1560778 | 316001 |
| CDC45 | 0.44240254 | 0.021473555 | Cdc45l | 287961 |
| TRIM36 | 0.64333475 | 0.02148699 | | 291597 |
| ATF4 | 0.45487285 | 0.021494819 | | 79255 |
| NVL | 0.39071107 | 0.021549927 | | 289323 |
| SHTN1 | -0.3805176 | 0.021569615 | RGD1311558\|Shootin1 | 292139 |
| MCCC2 | -0.4543736 | 0.021715138 | | 361884 |
| TWIST2 | -1.073762 | 0.021729017 | Dermo1 | 59327 |
| WDR34 | -0.35201335 | 0.021729643 | | 296618 |
| UBE2D1 | -0.5558267 | 0.02176936 | | 361831 |
| ANGPT2 | 0.9950371 | 0.02176936 | Agpt2\|Ang-2 | 89805 |
| MSTN | -0.9719133 | 0.021809116 | Gdf8 | 29152 |
| SPIN2A | -0.66989213 | 0.021809116 | RGD1560898\|Spin2 | 317395 |
| CDH7 | -0.7649024 | 0.02182333 | Cad7 | 29162 |
| NOP14 | 0.37772366 | 0.021906657 | Nol14\|RGD1305605 | 289724 |
| AZIN2 | -0.74265516 | 0.02193875 | Adc\|Azl2\|ODC-p\|RGD156477 | 366473 |
| TUBGCP3 | 0.33934143 | 0.021972759 | | 306599 |
| VPS37C | 0.3543599 | 0.022023605 | RGD1309258 | 308178 |
| COG3 | 0.46740416 | 0.022023605 | | 361073 |
| SCLY | -0.41821226 | 0.02206824 | | 363285 |
| MIDN | 0.48622456 | 0.02206824 | | 314623 |
| SLC5A10 | 1.0390279 | 0.02206824 | | 303205 |
| MAP7D2 | 0.6383688 | 0.022087066 | Brelin\|Mtap7d2\|RGD156485 | 317508 |
| DDX31 | 0.40275928 | 0.0221322 | | 311835 |
| ALDH3A2 | 0.41460213 | 0.0221322 | Aldh4\|FALDH | 65183 |
| EHD3 | 0.45017213 | 0.02218002 | Ehd2 | 192249 |
| CCDC116 | 0.72348905 | 0.022228336 | Sdf2l1 | 287936 |
| SCHIP1 | 0.44048518 | 0.02227399 | | 295105 |
| NEUROD2 | -0.57589155 | 0.022393469 | | 54276 |
| ZFYVE1 | -0.39420262 | 0.022393469 | | 299188 |
| ST18 | 0.9323301 | 0.022393469 | Nzf3\|r-MyT3 | 266680 |
| SFI1 | -0.3593983 | 0.02240066 | RGD1560636 | 305467 |
| IARS | 0.5432607 | 0.02241786 | Iarsl | 306804 |
| THNSL1 | -0.41821316 | 0.02246256 | | 498805 |
| KRT16 | 1.0515325 | 0.022464631 | Ka16\|Krt14\|Krt14l | 303530 |
| SLC39A10 | 0.41648242 | 0.022551391 | | 363229 |

| | | | | |
|---|---|---|---|---|
| SSR1 | 0.23996204 | 0.022580769 | Ac2-238 | 361233 |
| TMEM95 | 0.99867725 | 0.02259108 | | 691982 |
| MRPS12 | -0.37447104 | 0.022634368 | | 292758 |
| TMEM19 | -0.45933667 | 0.022701034 | | 299800 |
| RBM26 | 0.35263962 | 0.022743149 | RGD1308297 | 306137 |
| DPYD | -0.40603387 | 0.022779727 | | 81656 |
| DPM2 | -0.34453657 | 0.022779727 | | 29640 |
| ICAM5 | 0.6350212 | 0.023105703 | | 313785 |
| NR1H4 | 1.0299447 | 0.02314844 | Fxr | 60351 |
| AMPD2 | -0.35778847 | 0.02316695 | Ampd | 362015 |
| SF3A1 | 0.30246204 | 0.023192532 | | 305479 |
| RTF1 | -0.37656203 | 0.023204755 | Gtl7 | 366169 |
| BNIP1 | -0.47995958 | 0.02324584 | Bnip1l1 | 140932 |
| C2 | 0.65125006 | 0.023256697 | | 24231 |
| EIF5A | 0.3161303 | 0.02325939 | | 287444 |
| PODXL2 | -0.46132386 | 0.02325958 | RGD1305815 | 297433 |
| GPRC5D | 1.0628122 | 0.023274288 | RGD1563887 | 500349 |
| EBPL | -0.81103384 | 0.023533585 | | 361054 |
| RNFT2 | -0.47578886 | 0.023533585 | RGD1560195\|Tmem118 | 304521 |
| SDR39U1 | -0.47096366 | 0.023644267 | RGD1309307 | 361044 |
| SYNJ2BP | 0.44070727 | 0.02364881 | Omp25 | 64531 |
| HDGFRP3 | 0.23701152 | 0.0237519 | | 252941 |
| POFUT2 | 0.2858751 | 0.02376734 | | 309686 |
| AGPAT4 | 0.3172423 | 0.023829859 | | 170919 |
| ARHGAP26 | 0.6762538 | 0.023879444 | | 307459 |
| PTH2 | 1.0623065 | 0.0239008 | RGD1559447\|Tifp39\|Tip39 | 499149 |
| HNMT | -0.7935456 | 0.023969265 | | 81676 |
| EXOC6B | 0.51423776 | 0.024024148 | RGD1560638 | 500233 |
| ENHO | -0.77281755 | 0.024061421 | RGD1565232 | 100912292 |
| ZFP598 | 0.38935694 | 0.024068875 | Ntrap\|Znf598 | 287119 |
| HEATR5B | -0.47741523 | 0.024102585 | | 362683 |
| RAB26 | -0.6244838 | 0.024215654 | | 171111 |
| ZMAT3 | 0.60328597 | 0.024220372 | PAG608\|Wig1 | 64394 |
| FBXW17 | -0.52784884 | 0.024243416 | RGD1566133 | 361219 |
| TRIM45 | -0.6569792 | 0.024276678 | | 295323 |
| SMIM3 | -0.43222162 | 0.02429222 | Nid67 | 286910 |
| PCDHA3 | -0.4330155 | 0.024306206 | | 116780 |
| HIPK2 | 0.72276527 | 0.024310857 | | 362342 |
| DSG3 | 1.039333 | 0.024361389 | | 291752 |
| DUSP8 | -0.4727663 | 0.024405938 | Hb5\|M3/6 | 361679 |
| NR1I3 | -0.47867706 | 0.024651837 | CAR | 65035 |
| MAP3K5 | -0.5257784 | 0.024662388 | Ask1\|RGD1306565 | 365057 |
| RNMTL1 | 0.5180982 | 0.0246857 | RGD1309077\|Rnmtl1 | 360569 |

| | | | | |
|---|---|---|---|---|
| NXNL2 | 0.8030359 | 0.0246857 | Rdcvf2 | 689232 |
| CLTC | 0.4065264 | 0.02470702 | | 54241 |
| DCTD | 0.5415651 | 0.02474614 | | 290741 |
| GMPPB | 0.35616145 | 0.024762154 | RGD1560458 | 363145 |
| SHISA2 | 0.67983407 | 0.024807438 | RGD1561256\|Tmem46 | 498528 |
| N4BP2L2 | 0.4002115 | 0.024890715 | | 288416 |
| RGS7 | 0.4333365 | 0.024890715 | | 54296 |
| PLPPR3 | -0.36046594 | 0.025061086 | Lppr3\|Prg-2 | 314614 |
| RGD1309870 | -0.9370675 | 0.025096979 | | 289778 |
| SMARCA2 | -0.3563186 | 0.025176045 | | 361745 |
| ATMIN | 0.3480729 | 0.025186088 | Asciz\|RGD1305781 | 315037 |
| CCBL2 | -0.4399762 | 0.025223214 | Kat3 | 541589 |
| CIDEA | 1.0524814 | 0.025334332 | | 291541 |
| CRIP2 | -0.36622167 | 0.025359625 | CRP2 | 338401 |
| HIRA | 0.5345021 | 0.025456967 | | 363849 |
| TWISTNB | 0.56608593 | 0.025475383 | | 362728 |
| ENTPD3 | -0.6030954 | 0.025481807 | NTPDase3 | 316077 |
| PMP22 | 0.30523092 | 0.025500532 | Gas-3 | 24660 |
| TRPC7 | -0.9399961 | 0.025614291 | | 282822 |
| KATNBL1 | 0.4713918 | 0.025625682 | | 691543 |
| USP24 | 0.584131 | 0.025641214 | | 313427 |
| RAVER1 | 0.44098482 | 0.025709024 | Raver1h | 298705 |
| FNDC5 | -0.41621748 | 0.025931872 | | 260327 |
| PMAIP1 | 0.9662358 | 0.025931872 | Noxa | 492821 |
| NR2F1 | -0.6552084 | 0.02600139 | COUP-TFI | 81808 |
| PDIA5 | 0.75548553 | 0.02600139 | | 360722 |
| HPDL | 0.6105229 | 0.026015805 | Gloxd1\|RGD1310014 | 313521 |
| INHBB | 0.5896933 | 0.02605992 | | 25196 |
| CAPN1 | -0.8966967 | 0.026103247 | | 29153 |
| EHF | 1.021225 | 0.026108855 | | 295965 |
| ARFIP1 | 0.44239298 | 0.026112726 | | 60382 |
| MCL1 | 0.3782775 | 0.026118701 | | 60430 |
| SIKE1 | 0.24370816 | 0.026174108 | RGD1311316\|Sike | 362007 |
| SRPK2 | 0.29034224 | 0.026197786 | | 296753 |
| ZBTB9 | 0.33083826 | 0.026280228 | 3930402F13Rik | 294289 |
| ZFP213 | -0.49818873 | 0.026280472 | Znf213 | 287094 |
| NTM | -0.28882337 | 0.026316976 | Hnt\|RNU16845 | 50864 |
| AIFM1 | -0.3125007 | 0.026340885 | Aif\|Pdcd8 | 83533 |
| CASP8AP2 | 0.4594791 | 0.026352275 | | 313128 |
| LDHA | 0.43479967 | 0.026459916 | Ldh1 | 24533 |
| VPS25 | -0.42980328 | 0.026719892 | | 681059 |
| SLC25A20 | -0.3190037 | 0.026738843 | | 117035 |
| LOC100910945 | -0.46788025 | 0.026746746 | | 100910945 |

| | | | | |
|---|---|---|---|---|
| SNCA | -0.6050173 | 0.026817571 | | 29219 |
| PLEKHA1 | 0.25035676 | 0.026821863 | RGD1564153 | 361659 |
| AKT2 | 0.5054009 | 0.026897125 | | 25233 |
| KLF4 | 0.90540993 | 0.02692236 | GKLF | 114505 |
| DEPDC5 | -0.40845168 | 0.026927 | | 305464 |
| NGB | -0.6928232 | 0.026994903 | | 85382 |
| IVD | -0.39578953 | 0.027016677 | | 24513 |
| PSMG1 | -0.47655776 | 0.027284902 | Dscr2 | 288236 |
| GORASP2 | 0.2791223 | 0.027284902 | GRASP55\|Grs2 | 113961 |
| ATP10A | 0.7995832 | 0.02731377 | | 365266 |
| TBL3 | 0.31145382 | 0.027333464 | | 287120 |
| QDPR | -0.28917953 | 0.02734116 | | 64192 |
| SPSB4 | -0.6106562 | 0.027451115 | RGD1562307 | 300950 |
| TYR | -0.9649715 | 0.02745966 | C | 308800 |
| RNASEH2A | -0.5348562 | 0.02767752 | | 364974 |
| SDPR | -0.81919664 | 0.027761383 | | 316384 |
| CACNG8 | 0.47115186 | 0.027788473 | | 140729 |
| IFNG | 0.96064746 | 0.027807288 | IFNG2 | 25712 |
| RBL1 | 0.63687265 | 0.027824886 | | 680111 |
| RANBP6 | 0.26083702 | 0.027894719 | | 309326 |
| GPM6A | -0.5804157 | 0.028009865 | M6a | 306439 |
| TIMELESS | 0.5192112 | 0.02803853 | Tim | 83508 |
| SLITRK3 | -0.5794967 | 0.028080568 | | 310519 |
| CEP120 | -0.25485238 | 0.028080568 | Ccdc100\|RGD1565619 | 307302 |
| MIPEP | -0.30178705 | 0.02809246 | | 81684 |
| CDC42SE1 | 0.23582925 | 0.028155314 | | 499672 |
| SLC41A3 | -0.53476036 | 0.028180331 | | 641603 |
| TMEM8A | -0.41689226 | 0.02822031 | M83\|Tmem8 | 303004 |
| ACTN2 | -0.4354072 | 0.028269438 | | 291245 |
| VPS53 | -0.28410313 | 0.028292362 | RGD1311391 | 287535 |
| COMMD1 | -0.6125608 | 0.028329818 | | 289831 |
| CDH22 | 0.44884676 | 0.028351601 | | 29182 |
| HIST3H2A | -0.5952332 | 0.028359205 | H2a | 64646 |
| EFNA1 | -0.98023325 | 0.028365927 | B61 | 94268 |
| RNF111 | 0.2780099 | 0.028372575 | | 300813 |
| WDHD1 | 0.58513665 | 0.028372575 | | 305827 |
| LASP1 | -0.43409932 | 0.028413778 | | 29278 |
| CDK2AP1 | 0.51430607 | 0.028428685 | | 360804 |
| SDC2 | 0.40368718 | 0.02844198 | HSPG | 25615 |
| DGUOK | -0.7135703 | 0.028450714 | | 297389 |
| TNFRSF11A | -0.5661064 | 0.0285034 | RANK\|RGD1563614 | 498206 |
| SPATA2L | 0.43527207 | 0.028505994 | RGD1560840 | 498963 |
| TANGO2 | -0.4104154 | 0.028557584 | RGD1310348 | 360738 |

| | | | | |
|---|---|---|---|---|
| LYAR | 0.6589188 | 0.028607719 | | 289707 |
| TAF5L | 0.6951462 | 0.028670723 | | 307927 |
| FIBP | -0.52428794 | 0.028681373 | | 282837 |
| N4BP2L1 | -0.5847066 | 0.028689358 | | 498131 |
| PCBP2 | 0.33674398 | 0.028723367 | | 363005 |
| RGD1309104 | 0.38246995 | 0.028741185 | | 289084 |
| EIF2AK4 | -0.41502848 | 0.028792525 | | 114859 |
| DPYSL2 | -0.5521578 | 0.02881728 | Crmp2\|TOAD-64 | 25416 |
| NAAA | -0.9980757 | 0.02890205 | Asahl | 497009 |
| CHMP6 | -0.37391767 | 0.028960612 | RGD1565325 | 287873 |
| ACKR2 | -0.9902222 | 0.02896535 | Ccbp2\|D6 | 140473 |
| TMBIM6 | 0.24474776 | 0.02903055 | Ab1-011\|Ac1-149\|Cc1-27\|Te | 24822 |
| SAPCD1 | -0.5870442 | 0.029040692 | Ng23 | 406170 |
| GPAT4 | 0.29193175 | 0.029048502 | Agpat6\|RGD1310520 | 290843 |
| GPR26 | 0.79486054 | 0.02906515 | | 192153 |
| DMC1 | -0.954366 | 0.029074695 | Dmc1h | 362960 |
| ACAA1 | -0.3464555 | 0.029277159 | Acaa\|Acaa1a\|Pktaa | 24157 |
| MKRN1 | -0.32218188 | 0.029280579 | | 296988 |
| GDAP2 | 0.2811545 | 0.02932361 | | 362004 |
| GPANK1 | -0.55446744 | 0.029328948 | Bat4 | 415064 |
| BLOC1S5 | -0.52388054 | 0.029396914 | Muted | 306868 |
| MAPRE1 | 0.30936545 | 0.029399548 | Eb1 | 114764 |
| ZIC2 | 0.5563749 | 0.029415943 | | 361096 |
| SLC2A13 | -0.44457495 | 0.029434267 | Hmit | 171147 |
| TIMP3 | 0.66125244 | 0.029497175 | | 25358 |
| SLC2A1 | 0.4447203 | 0.029523157 | GLUTB\|GTG1\|Glut1\|Gtg3\|RA | 24778 |
| GFRA2 | 0.7065282 | 0.029685542 | Retl2 | 25136 |
| RARS | 0.41149497 | 0.029705409 | | 287191 |
| FAF1 | 0.34779784 | 0.029716888 | | 140657 |
| TPM2 | 0.8883627 | 0.029716888 | | 500450 |
| SPCS3 | 0.35186204 | 0.029738758 | | 680782 |
| NUDT14 | -0.6949743 | 0.029768933 | | 299346 |
| TRAPPC2B | -0.56372106 | 0.02983081 | | 100910318 |
| SESN2 | 0.63824654 | 0.029846989 | RGD1566319 | 502988 |
| EXOC3L1 | -0.6660272 | 0.029977003 | Exoc3l\|RGD1311132 | 291961 |
| MYPOP | -0.6344545 | 0.03004929 | P42pop\|RGD1565160 | 499090 |
| SYT13 | -0.48842904 | 0.030050488 | | 80977 |
| LOC689986 | -0.37586313 | 0.030110314 | | 689986 |
| IFT122 | -0.36788243 | 0.030110314 | Wdr10 | 312651 |
| SWSAP1 | -0.48390603 | 0.03014336 | RGD1308026 | 363029 |
| LRRC34 | -0.8412541 | 0.030171528 | RGD1565052 | 499589 |
| ZHX2 | 0.55784607 | 0.030193167 | | 314988 |
| FBXL14 | 0.4120996 | 0.030285114 | | 312675 |

| | | | | |
|---|---|---|---|---|
| SLC4A5 | -0.92623943 | 0.030432098 | NBC4 | 297386 |
| SRCIN1 | -0.5569027 | 0.030438466 | P140\|Snip | 56029 |
| IRX6 | 1.0180786 | 0.030557713 | RGD1564830 | 307715 |
| TRERF1 | -0.6184687 | 0.03057336 | | 316219 |
| WNT16 | -1.0102184 | 0.030616408 | | 500047 |
| ZFP697 | 0.8302689 | 0.030626684 | RGD1308711\|Znf697 | 295310 |
| CACNB4 | 0.641858 | 0.030679084 | | 58942 |
| THSD7A | -0.6736665 | 0.030782651 | RGD1566201 | 500032 |
| PCDHGA2 | -0.493664 | 0.030789359 | | 498846 |
| LARS2 | -0.39411244 | 0.030789359 | | 363172 |
| CUL2 | 0.24325298 | 0.030789359 | | 361258 |
| CLINT1 | 0.54087186 | 0.030789359 | Clint\|Enth\|Epn4 | 360515 |
| NIT1 | -0.2994909 | 0.030802669 | | 289222 |
| AARS | 0.37740648 | 0.030811546 | | 292023 |
| LHFPL5 | -0.72176534 | 0.030828876 | Tmhs | 294303 |
| BCL2L11 | -0.5232826 | 0.030841896 | Bim\|BimL | 64547 |
| HDX | 0.7631826 | 0.030876638 | RGD1563666 | 317617 |
| APCS | 0.9416904 | 0.030882161 | Sap | 29339 |
| SLC4A10 | -0.45854294 | 0.03095376 | NCBE | 295645 |
| CCDC65 | -0.37604862 | 0.03095376 | | 362994 |
| RHOT1 | 0.26838666 | 0.031272113 | | 303351 |
| STAT2 | -0.37317953 | 0.031295735 | | 288774 |
| SCN1A | -0.72736907 | 0.031329434 | | 81574 |
| STEAP2 | 0.40834293 | 0.03136245 | | 312052 |
| RAMP2 | -0.6928447 | 0.031428874 | | 58966 |
| CPT1C | -0.35671914 | 0.031512763 | CPT IC\|CPT1-B\|CPTI-B | 308579 |
| FBXL20 | -0.6405506 | 0.031550296 | Fbl2\|Fbxl2 | 64039 |
| NRG1 | 0.55282503 | 0.0316391 | | 112400 |
| S100PBP | -0.5235443 | 0.031642627 | RGD1564943 | 500551 |
| NAA40 | 0.51848483 | 0.031666256 | Nat11\|RGD1565838 | 361718 |
| IL25 | -1.0208933 | 0.031695936 | RGD1561632 | 501996 |
| DCPS | -0.34323475 | 0.031695936 | Hint-5 | 266605 |
| CORO2A | -0.488291 | 0.031715386 | | 313235 |
| RNF5 | -0.45976725 | 0.031731933 | | 407784 |
| HK2 | 0.47732583 | 0.03179401 | | 25059 |
| GEMIN8 | -0.41399086 | 0.0318039 | Fam51a1 | 363462 |
| POU1F1 | 1.0202218 | 0.0318039 | GHF1\|GHF1A\|PIT1Z\|Pit1 | 25517 |
| DCAF7 | -0.3669658 | 0.031858068 | RGD1305140\|Wdr68 | 303602 |
| ATP6V1B2 | 0.42750973 | 0.031886395 | Atp6b1b2\|Atp6b2\|Vatb | 117596 |
| SMAP2 | 0.3646476 | 0.03202195 | RGD1308418\|Smap1l | 298500 |
| ATP11A | 0.49652663 | 0.03202961 | Ua20 | 306600 |
| DDB2 | -0.5134682 | 0.03221289 | | 100362121 |
| RGD1308601 | 0.568617 | 0.0323027 | | 307249 |

| | | | | |
|---|---|---|---|---|
| TRIP12 | 0.48558667 | 0.03232823 | Gtl6\|TRIP-12 | 316575 |
| RAD21 | -0.26925308 | 0.032404546 | | 314949 |
| RGS5 | 0.9978223 | 0.03242148 | | 54294 |
| POGLUT1 | 0.39570394 | 0.032437116 | Ktelc1\|RGD1306248 | 288091 |
| PATL1 | 0.5256846 | 0.032437116 | Pat1b\|RGD1305514 | 361736 |
| SLC1A4 | 0.54220474 | 0.032437116 | ASCT1 | 305540 |
| XRCC4 | -0.39937168 | 0.032484468 | | 309995 |
| LRRC41 | 0.385255 | 0.032493487 | RGD1311221 | 362566 |
| LOC680039 | 0.46632928 | 0.032493487 | | 680039 |
| TLK2 | 0.31502602 | 0.032496337 | | 303592 |
| NCSTN | -0.26698932 | 0.032625727 | | 289231 |
| SPATS2L | -0.40924394 | 0.032649647 | RGD1309930 | 316426 |
| MPP4 | 0.7599157 | 0.032875896 | Dlg6 | 58808 |
| SLC5A8 | 1.0145775 | 0.033064038 | RGD1564146 | 500820 |
| BARHL2 | 0.9757056 | 0.033074524 | Barh\|MBH1 | 65050 |
| ERF | 0.47308475 | 0.033135038 | | 292721 |
| 42982 | -0.764245 | 0.033291806 | EG3-1RVC\|EG3RVC\|Pnutl2 | 287606 |
| B4GALT5 | 0.33854157 | 0.033291806 | | 362275 |
| MTERF2 | -0.63347954 | 0.033466984 | Mterfd3\|RGD1311836 | 366856 |
| USF1 | -0.32495648 | 0.033466984 | | 83586 |
| PKNOX1 | 0.6513941 | 0.03349886 | | 294322 |
| LOXHD1 | 1.0126847 | 0.033543635 | | 291427 |
| PANK4 | 0.3367602 | 0.03362458 | Fang1 | 171053 |
| DHRS3 | 0.47700343 | 0.033664837 | | 313689 |
| PHF1 | -0.4050112 | 0.033805452 | Tctex3 | 294287 |
| CDC14B | 0.7017648 | 0.033805452 | | 361195 |
| ZFYVE16 | -0.5173794 | 0.033865884 | RGD1564784 | 499508 |
| AKR1E2 | -0.483803 | 0.033865884 | Akr1cl2\|Akr1e1 | 307091 |
| CHURC1 | -0.601889 | 0.033934236 | | 299154 |
| USP49 | -0.6277844 | 0.034041964 | RGD1310513 | 316211 |
| CRYM | -0.53354144 | 0.03418983 | CDK108 | 117024 |
| SRSF9 | 0.3987796 | 0.034256175 | Sfrs9 | 288701 |
| CDPF1 | -0.5317033 | 0.034327634 | RGD1306001 | 362975 |
| RPH3A | -0.6514723 | 0.034347903 | | 171039 |
| CCDC53 | -0.4767593 | 0.034459036 | RGD1563761 | 299707 |
| PHF14 | -0.41190183 | 0.034459036 | RGD1563764 | 500030 |
| MINK1 | 0.39588234 | 0.034459036 | MEKKK 6 | 303259 |
| NET1 | -0.32405576 | 0.034461897 | | 307098 |
| TLE3 | 0.3218495 | 0.03449508 | Esp3 | 84424 |
| PAPD4 | 0.43107226 | 0.03457216 | | 361878 |
| MESP1 | -0.97030115 | 0.034614347 | | 308766 |
| GABRQ | 0.72086304 | 0.034620695 | | 65187 |
| RAB14 | 0.27399725 | 0.03472653 | | 94197 |

| | | | | |
|---|---|---|---|---|
| MFSD14B | 0.24501477 | 0.034771945 | Hiatl1\|RGD1308377 | 306687 |
| MCM10 | 0.53480804 | 0.034814026 | | 307126 |
| TNFAIP2 | -0.80376416 | 0.034970935 | | 299339 |
| RASSF6 | -1.0063156 | 0.03497475 | | 305251 |
| VRK3 | -0.3422794 | 0.035095934 | | 361565 |
| SEC61A2 | 0.2270908 | 0.03526537 | | 361273 |
| PEX5 | 0.5473612 | 0.03528934 | PTS1-BP\|PTS1R | 312703 |
| NME7 | -0.45875064 | 0.03532765 | Nm23-r7 | 171566 |
| DDX19B | 0.31900993 | 0.035466228 | Zd10a | 690693 |
| DNMT1 | 0.34042725 | 0.035562135 | | 84350 |
| GPRASP1 | -0.24657959 | 0.035594195 | pips | 171407 |
| CCNO | -0.83561397 | 0.03564552 | RGD1565217 | 499528 |
| CACUL1 | 0.45192146 | 0.03565909 | RGD1308127 | 365493 |
| PLAT | 0.5291216 | 0.03566246 | PATISS\|tPA | 25692 |
| MRGBP | 0.6698152 | 0.035740416 | RGD1308612 | 311713 |
| PDE12 | 0.5205767 | 0.03574115 | LRRGT00074\|RGD1310975 | 306231 |
| B3GNT3 | -0.95138377 | 0.03587172 | | 290638 |
| TGFB1 | 0.6852653 | 0.035985652 | Tgfb | 59086 |
| CHML | 0.8247513 | 0.036103465 | | 689102 |
| CDK5RAP3 | -0.41523105 | 0.03611087 | C53 | 80278 |
| TSPAN15 | -0.38312688 | 0.0361306 | | 679462 |
| PPP2R5A | 0.3153681 | 0.036137 | | 312754 |
| PDZD7 | -0.5707858 | 0.036299657 | Pdzk7 | 293996 |
| JPH3 | -0.42839488 | 0.036348023 | | 307916 |
| SLK | 0.5161005 | 0.036348023 | Stk2 | 54308 |
| MMP11 | -0.59045744 | 0.03639457 | ST3 | 25481 |
| ALAS1 | 0.27712983 | 0.0364049 | ALAS | 65155 |
| SLC35D3 | 0.7441862 | 0.03642329 | | 308717 |
| NOP2 | 0.30664676 | 0.036445048 | Nol1 | 314969 |
| ZFP39 | -0.43446797 | 0.036447357 | | 303173 |
| ZFP397 | 0.6151469 | 0.036568984 | Znf397 | 100909408 |
| ITPRIPL2 | 0.9190477 | 0.03661702 | RGD1564255 | 499253 |
| ALDH1L2 | -0.44637772 | 0.0366413 | RGD1309458 | 299699 |
| MCEE | -0.5760194 | 0.036644287 | | 293829 |
| ELP5 | 0.28309202 | 0.036664568 | DERP6\|Rai12 | 287446 |
| SNN | -0.3975091 | 0.03668417 | | 29140 |
| TUBG2 | -0.46170938 | 0.03676427 | | 680991 |
| CDK5 | -0.45083693 | 0.03676427 | | 140908 |
| TUBA3B | 0.98023945 | 0.036987737 | RGD1565155 | 500363 |
| PPWD1 | 0.34223482 | 0.037053958 | RGD1310204 | 294711 |
| ZC3H8 | -0.52011245 | 0.037083894 | Zc3hdc8 | 311414 |
| SC5D | 0.3328679 | 0.037091367 | Sc5dl | 114100 |
| SLC35D1 | 0.5396388 | 0.037110966 | | 298280 |

| | | | | |
|---|---|---|---|---|
| RGD1309079 | 0.42236546 | 0.037203502 | Ab2-095 | 315891 |
| SZRD1 | 0.2651899 | 0.03728946 | C1orf144\|RGD1560286 | 500575 |
| PGGT1B | 0.33448288 | 0.03728946 | | 81746 |
| CROCC | -0.508079 | 0.037360642 | | 313663 |
| ZGPAT | 0.29030257 | 0.03737169 | RGD1310801 | 296478 |
| ADPGK | 0.31411034 | 0.03744561 | RGD1306103 | 315722 |
| HTR5A | -0.67148125 | 0.037459828 | 5HT5A\|MR22 | 25689 |
| PERP | 0.590914 | 0.037459828 | | 292949 |
| RPS6KB1 | 0.37649137 | 0.03754645 | | 83840 |
| CREG1 | -0.26421645 | 0.03756935 | Creg | 289185 |
| NMRK1 | -0.60891306 | 0.037603244 | Nrk1 | 499330 |
| ARRDC3 | -0.42791298 | 0.037853606 | LRRGT00048 | 309945 |
| ARHGEF2 | -0.50566804 | 0.038085505 | | 310635 |
| ARHGEF25 | -0.41118413 | 0.0381102 | Geft | 314904 |
| COIL | 0.32240334 | 0.038111575 | | 50998 |
| NKAP | -0.59048504 | 0.038151268 | 2610020o08rik | 298342 |
| ETHE1 | -0.36724612 | 0.038245164 | | 292710 |
| NBR1 | -0.32224736 | 0.038245164 | Ca125\|RGD1311421 | 303554 |
| RPF1 | 0.58397985 | 0.038253207 | Bxdc5 | 499725 |
| SLC37A1 | 0.5571413 | 0.03828822 | | 294321 |
| SRSF2 | 0.53308135 | 0.038305525 | Sfrs2 | 494445 |
| ALG10 | 0.7551815 | 0.038305525 | Alg10b\|KCR1 | 245960 |
| SLC16A8 | 0.9730974 | 0.038305525 | Mct3 | 65200 |
| TRIM54 | -0.9230427 | 0.038337693 | Rnf30 | 362708 |
| PKM | 0.5263086 | 0.038356286 | PKM12\|Pk3\|Pkm2 | 25630 |
| KIT | -0.5006792 | 0.038370207 | | 64030 |
| FDXR | -0.5313651 | 0.0385795 | AR | 79122 |
| SNX7 | 0.398886 | 0.038629517 | | 310815 |
| ENPP1 | 0.72993696 | 0.038642246 | Npps\|Pc1 | 85496 |
| PAWR | 0.5601135 | 0.03864341 | Par-4\|Par4 | 64513 |
| SIRT7 | 0.42087793 | 0.038659964 | | 303745 |
| CDC42EP3 | -0.44628868 | 0.03866876 | | 313838 |
| LSG1 | 0.32500356 | 0.03869334 | RGD1309089 | 288029 |
| ZFP46 | -0.440568 | 0.03882037 | | 298558 |
| TMEM41A | 0.5112608 | 0.03897502 | | 681708 |
| ADCYAP1 | 0.6414165 | 0.03897502 | Pacap | 24166 |
| CCSAP | 0.43907428 | 0.03899341 | RGD1563235 | 307926 |
| ZFP513 | 0.26082954 | 0.039262705 | Znf513 | 313913 |
| CPTP | -0.57773244 | 0.03937554 | Gltpd1 | 313771 |
| FBXO44 | -0.54700243 | 0.03943288 | RGD1562463 | 500587 |
| HSPA4 | 0.30015755 | 0.039481673 | Hsp110\|Hsp70\|irp94 | 266759 |
| ANKRD52 | 0.68396103 | 0.039582606 | RGD1307124 | 362811 |
| NOSIP | -0.43927485 | 0.03969875 | | 292894 |

| | | | | |
|---|---|---|---|---|
| RNF141 | -0.36530054 | 0.039720736 | | 308900 |
| TMEM179 | -0.5009015 | 0.039815802 | RGD1310269 | 314472 |
| RAB11B | -0.41509232 | 0.039815802 | | 79434 |
| CHEK1 | 0.50829446 | 0.039869428 | | 140583 |
| RAB3A | -0.39905143 | 0.040116232 | RAB3 | 25531 |
| ERAL1 | -0.3119239 | 0.040128108 | | 363646 |
| TACC1 | 0.471592 | 0.0402329 | Tacc1a | 306562 |
| FXYD6 | -0.32307225 | 0.040235706 | Php | 63847 |
| PHF21B | 0.6334849 | 0.040290035 | RGD1308739 | 300117 |
| GORAB | 0.44201726 | 0.0403276 | Scyl1bp1 | 304923 |
| SIL1 | -0.56205726 | 0.040359624 | | 291673 |
| AK2 | 0.5197275 | 0.04037918 | | 24184 |
| UBN1 | 0.28962803 | 0.04038311 | | 302935 |
| PIAS3 | -0.45282632 | 0.04042212 | | 83614 |
| JPH4 | -0.51739067 | 0.04046165 | | 445271 |
| PGD | -0.33300146 | 0.040510073 | Cc2-27 | 100360180 |
| ACADS | -0.35135093 | 0.040542137 | Scad | 64304 |
| CAP2 | 0.34800616 | 0.04064034 | | 116653 |
| ZBTB8B | -0.50454503 | 0.040657874 | RGD1562327\|Zbtb8\|Zbtb8l | 500553 |
| PADI2 | -0.50108236 | 0.040773638 | Pdi2 | 29511 |
| ERLEC1 | -0.46808836 | 0.040773638 | RGD1306508 | 289874 |
| TCTN1 | -0.3580323 | 0.040773638 | RGD1566266 | 304486 |
| ST8SIA2 | -0.69829434 | 0.040880658 | Siat8b | 117523 |
| VWA5B2 | -0.4674993 | 0.040880658 | RGD1564491 | 303812 |
| HNRNPAB | 0.33188716 | 0.040880658 | A1F-C1\|Hnrpab | 83498 |
| FGF7 | 0.96359503 | 0.040880658 | Kgf | 29348 |
| LHX4 | 0.79457676 | 0.040988516 | | 360858 |
| MED4 | -0.36088556 | 0.041012213 | Vdrip | 306030 |
| YTHDF1 | 0.23782311 | 0.041012213 | | 296467 |
| CDCA7 | 0.40414777 | 0.041012213 | | 311742 |
| CBLB | -0.6886468 | 0.041081518 | | 171136 |
| SAP30BP | 0.32180488 | 0.041081518 | RGD1565537 | 360662 |
| METTL6 | 0.3548717 | 0.04120367 | | 290564 |
| MYLK | -0.6736588 | 0.04122695 | | 288057 |
| ITPR1 | -0.43509513 | 0.041246727 | I145TR\|IP3R1\|InsP3R\|InsP3R | 25262 |
| CABLES1 | -0.31079614 | 0.041444834 | | 307585 |
| SIRT3 | -0.3815202 | 0.041483484 | | 293615 |
| OBFC1 | -0.48785168 | 0.041653346 | | 294025 |
| LGMN | -0.4387056 | 0.04174131 | Prsc1 | 63865 |
| SOAT1 | 0.6703569 | 0.04174131 | Acat-1 | 81782 |
| CDH9 | -0.8313626 | 0.041754942 | | 29163 |
| ICA1 | -0.4694552 | 0.04176689 | Ica69 | 81024 |
| KDM1B | -0.31811586 | 0.04183083 | Aof1 | 306819 |

| | | | | |
|---|---|---|---|---|
| CEP57L1 | -0.5037394 | 0.041924905 | cep57R | 294519 |
| CAMK1 | -0.48706043 | 0.042015456 | Camki\|Gaip | 171503 |
| SRFBP1 | 0.40503505 | 0.042069778 | p49/STRAP | 291469 |
| TRIM2 | -0.30773902 | 0.0421367 | | 361970 |
| TSPAN31 | 0.2815228 | 0.042165656 | Sas | 362890 |
| TNRC18 | -0.57466465 | 0.042166073 | Zfp469 | 304302 |
| GPR101 | 0.9520749 | 0.042299286 | RGD1564196 | 317608 |
| GPX3 | -0.553081 | 0.042353693 | GPx-3\|GPx-P\|GSHPx-3\|GSHP | 64317 |
| ZFP521 | 0.37923446 | 0.042425748 | Znf521 | 307579 |
| PRR5 | -0.6435585 | 0.0425764 | Protor1 | 315189 |
| RASGRP1 | 0.4747592 | 0.042591874 | Rasgrp | 29434 |
| GGACT | -0.52912664 | 0.04270827 | A2ld1 | 290500 |
| GTF2F1 | 0.2910501 | 0.04270827 | | 316123 |
| POLR1E | 0.46758574 | 0.042786576 | Praf1\|RGD1565773 | 313245 |
| MYRIP | 0.47532845 | 0.043024935 | Slac2c | 360034 |
| MGC95210 | -0.425667 | 0.043049075 | | 287798 |
| ANK1 | 0.63647246 | 0.043114673 | | 306570 |
| SOCS6 | 0.3937871 | 0.043192185 | | 307200 |
| LOC688390 | -0.661496 | 0.04323364 | | 688390 |
| CSPG5 | -0.47791874 | 0.043257702 | Caleb\|Ngc | 50568 |
| TIA1 | -0.39518464 | 0.043443162 | | 312510 |
| GM2A | -0.31562626 | 0.043772697 | | 282838 |
| FBP2 | -0.8136375 | 0.043803196 | FBPase | 114508 |
| ZFP367 | 0.37631714 | 0.043803196 | Znf367 | 306695 |
| CRK | 0.55220175 | 0.043810494 | Crko | 54245 |
| LYPLA2 | 0.41494977 | 0.043842703 | | 83510 |
| RGD1565033 | -0.5849035 | 0.04386195 | | 498014 |
| EIF2B5 | 0.26243556 | 0.04387254 | | 192234 |
| FAM192A | -0.44237024 | 0.044030946 | Nip30\|RGD1307433 | 307652 |
| EPN2 | 0.2372426 | 0.044030946 | | 60443 |
| SV2A | -0.44024414 | 0.044125818 | Sv2 | 117559 |
| OVCA2 | -0.4248868 | 0.04432062 | RGD1564623 | 497954 |
| CD226 | -0.8940893 | 0.044405293 | | 307199 |
| GLRA2 | -0.4702876 | 0.04443878 | RNNEOGLY | 24397 |
| NUP62 | 0.32010227 | 0.044462617 | Np62 | 65274 |
| TSC22D1 | -0.30597922 | 0.04449759 | TS22A\|TSC-22\|Tgfb1i4 | 498545 |
| MYBL2 | 0.502166 | 0.04449759 | | 296344 |
| TOLLIP | 0.3704476 | 0.044588216 | | 361677 |
| FPGS | 0.30807278 | 0.04468685 | | 687266 |
| PPM1J | -0.68527406 | 0.044715635 | | 295341 |
| ADSS | -0.2692896 | 0.044726953 | Adss2 | 289276 |
| FOSL2 | 0.8236007 | 0.044848252 | Fra2 | 25446 |
| NEK1 | -0.42037255 | 0.044919748 | | 290705 |

| | | | | |
|---|---|---|---|---|
| FUT11 | 0.37389797 | 0.044919748 | | 286971 |
| LDLR | 0.5501626 | 0.044919748 | LDLRA | 300438 |
| PA2G4 | 0.2990845 | 0.04504876 | Ebp1 | 288778 |
| PCYT1A | 0.5348623 | 0.045138597 | CCT-alpha\|CTP | 140544 |
| PPP1CA | 0.24071914 | 0.045298655 | PP1alpha | 24668 |
| TBC1D10A | 0.41351005 | 0.045365326 | Tbc1d10 | 360968 |
| RHBDF1 | 0.40119374 | 0.045441378 | | 303008 |
| HES1 | 0.45293653 | 0.045441378 | | 29577 |
| ADAMTS10 | 0.37756658 | 0.045457304 | | 314655 |
| MYH9L1 | 0.5848633 | 0.045488283 | | 25745 |
| BTBD6 | -0.35536626 | 0.045565758 | | 690367 |
| DYNC2LI1 | -0.5108052 | 0.045751467 | RGD1310286 | 298767 |
| SLCO4A1 | -0.40388513 | 0.045751467 | OATP-E\|Slc21a12 | 171144 |
| SHISA5 | -0.38090327 | 0.045767896 | | 301013 |
| FANCE | -0.50799674 | 0.045877352 | RGD1561045 | 309643 |
| CNP | -0.27311778 | 0.046021413 | CNPF\|CNPI\|CNPII\|Cnp1 | 25275 |
| C2CD2L | -0.36675042 | 0.046023235 | Tmem24 | 300666 |
| MPV17L2 | -0.42423078 | 0.046042807 | RGD1308064 | 290645 |
| ABHD17B | 0.38789335 | 0.04624426 | Cgi67l\|Fam108b1\|RGD13052 | 309399 |
| SYNGR4 | -0.89858645 | 0.046292707 | | 292916 |
| DDIT4 | -0.54277253 | 0.04639547 | Rtp801 | 140942 |
| SLC1A6 | -0.644496 | 0.04661186 | EAAT4 | 84012 |
| ZFP709 | 0.4258446 | 0.046780914 | Hit40\|Znf14 | 266773 |
| FGF13 | -0.47861964 | 0.04680864 | FGF-13 | 84488 |
| EEF2K | -0.41597348 | 0.046873935 | SMEF2K | 25435 |
| USP31 | 0.6709861 | 0.046946503 | | 308959 |
| SYNGR1 | -0.5001973 | 0.04698841 | | 29205 |
| FAM160A2 | 0.4618909 | 0.047051255 | FHIP | 293343 |
| TTC30B | -0.37847698 | 0.047068115 | | 499814 |
| ERO1A | 0.30140406 | 0.04717244 | Ero1l | 171562 |
| PMVK | -0.48543307 | 0.04733122 | | 310645 |
| RALGAPA1 | 0.45262334 | 0.047342747 | GRIPE\|Garnl1\|Tulip1\|p240 | 56785 |
| MAN1A2 | 0.58379173 | 0.04750291 | Man1b | 295319 |
| GPC1 | 0.44138172 | 0.047604747 | HSPG M12\|HSPGM12 | 58920 |
| NTN1 | 0.6117326 | 0.047619216 | netrin-1 | 114523 |
| ZBED4 | 0.6373196 | 0.04774105 | | 315211 |
| RBM33 | 0.62802976 | 0.047770053 | RGD1310651 | 362297 |
| CH25H | 0.8603463 | 0.047844984 | | 309527 |
| RHPN2 | -0.6892688 | 0.047902454 | | 308516 |
| FAT4 | -0.68607205 | 0.04809833 | RGD1564291 | 310341 |
| GABRD | -0.7051563 | 0.048128575 | GABAA-RD | 29689 |
| HACD3 | 0.26696298 | 0.04816285 | Ptplad1\|RGD1565496 | 300783 |
| DEAR | 0.41171983 | 0.04816285 | | 446170 |

| | | | | |
|---|---|---|---|---|
| HOMEZ | -0.4243091 | 0.048186395 | | 260325 |
| DPY19L1 | 0.24113286 | 0.048194468 | RGD1305822 | 315496 |
| MED11 | -0.46381506 | 0.04836857 | RGD1563202 | 287456 |
| HID1 | -0.34535822 | 0.04836857 | RGD1311422 | 287822 |
| RASSF5 | 0.42041242 | 0.04836857 | Maxp1\|Nore1 | 54355 |
| MEST | 0.52117264 | 0.04836857 | | 58827 |
| SPRN | -0.7253101 | 0.048448265 | | 541462 |
| WDR46 | 0.33697125 | 0.04854942 | Bing4 | 309628 |
| GMPPA | -0.34430286 | 0.048574977 | | 501167 |
| WDR26 | 0.35423997 | 0.048592504 | RGD1565589 | 498301 |
| SENP7 | -0.26622754 | 0.04878576 | | 288167 |
| ZFP583 | 0.3633975 | 0.04878576 | RGD1562044 | 499068 |
| SLC25A14 | -0.30390215 | 0.048834655 | Bmcp1 | 85263 |
| ARHGEF11 | -0.52531695 | 0.048962355 | Gtrap48 | 78966 |
| SQLE | 0.28197297 | 0.048962355 | | 29230 |
| LMO1 | -0.5265091 | 0.049009074 | Dat1\|Lmo3 | 245979 |
| SLC17A6 | 0.6108406 | 0.049009074 | Dnpi\|Vglut2 | 84487 |
| PACSIN1 | 0.6023192 | 0.04935037 | | 29704 |
| HEXDC | -0.33909637 | 0.049379133 | | 100216475 |
| SCYL3 | 0.4040984 | 0.049412906 | RGD1308992 | 360866 |
| UMPS | 0.3245573 | 0.049528614 | | 288051 |
| HSDL1 | -0.39937967 | 0.04956813 | RGD1308433 | 361418 |
| ISOC1 | -0.25592902 | 0.049739197 | DR-NR#1\|RGD1307632 | 364879 |
| CDC25A | 0.31931496 | 0.049744014 | | 171102 |
| YARS2 | 0.38417423 | 0.049744014 | CGI-04\|RGD1311696 | 287924 |
| NR4A3 | 0.46416077 | 0.049744014 | NOR-2 | 58853 |
| SHISA8 | 0.7559386 | 0.049744014 | RGD1563996 | 315163 |
| RBPMS | 0.8294909 | 0.049915213 | RGD1561067 | 498642 |
| KLHL25 | -0.38356707 | 0.04996569 | RGD1310815 | 293023 |
| ENPP5 | -0.21971719 | 0.04996569 | E-NPP 5\|Npp5 | 316249 |
| ZFP777 | 0.28266618 | 0.04996569 | RGD1566056\|Znf777 | 502764 |
| TNFAIP1 | -0.2973561 | 0.049998462 | Edp1 | 287543 |
| PABPN1 | 0.28493616 | 0.049998462 | | 116697 |

**W+B and RX2+B v V+B LIST**

| Gene Symbol | Gene ID | log2fc8-1 | log2FC_db_wp ( WP and DB v DW ) |
|---|---|---|---|
| MARCH2 | 362849 | 1.0184354 | -1.0184354 |
| MARCH8 | 312656 | 0.5621597 | -0.5621597 |
| MARCH9 | 679272 | 1.2818643 | -1.2818643 |
| MARCH11 | 499558 | -0.93179196 | 0.93179196 |
| SEP6 | 691335 | 1.0019512 | -1.0019512 |
| AATK | 690853 | 1.1776892 | -1.1776892 |
| ABCB10 | 361439 | 0.7567853 | -0.7567853 |
| ABCG1 | 85264 | 0.9672269 | -0.9672269 |
| ABHD14A | 300982 | 0.98221266 | -0.98221266 |
| ABHD6 | 305795 | 0.49674532 | -0.49674532 |
| ABHD8 | 306338 | 1.4025161 | -1.4025161 |
| ABI1 | 79249 | -0.3251728 | 0.3251728 |
| ABTB1 | 297432 | 1.1204531 | -1.1204531 |
| ACAA1 | 24157 | 1.1053482 | -1.1053482 |
| ACAN | 58968 | -1.5901684 | 1.5901684 |
| ACPP | 56780 | -1.9549279 | 1.9549279 |
| ACSL1 | 25288 | -0.25970042 | 0.25970042 |
| ACSL4 | 113976 | -0.5156731 | 0.5156731 |
| ACTN1 | 81634 | -1.6871865 | 1.6871865 |
| ACTN2 | 291245 | 0.9964308 | -0.9964308 |
| ACTR3B | 362298 | 1.1455386 | -1.1455386 |
| ACVR1 | 79558 | -0.89764786 | 0.89764786 |
| ACVR1C | 245921 | -2.020098 | 2.020098 |
| ADAMTS1 | 79252 | -1.3923155 | 1.3923155 |
| ADAMTS15 | 300474 | 1.4631126 | -1.4631126 |
| ADAMTS2 | 287899 | 1.6056658 | -1.6056658 |
| ADAMTS5 | 304135 | -2.0638483 | 2.0638483 |
| ADAMTS7 | 315879 | 1.2275528 | -1.2275528 |
| ADAP1 | 171097 | 1.135771 | -1.135771 |
| ADCK3 | 360887 | 1.6959999 | -1.6959999 |
| ADCY7 | 84420 | -1.5113342 | 1.5113342 |
| ADCY8 | 29241 | -1.4688996 | 1.4688996 |
| ADGRB3 | 301309 | 0.9077113 | -0.9077113 |
| ADGRE5 | 361383 | -1.1002142 | 1.1002142 |
| ADM | 25026 | -0.8431437 | 0.8431437 |
| ADORA1 | 29290 | -1.5732242 | 1.5732242 |
| ADORA2A | 25369 | 2.0874836 | -2.0874836 |
| ADRA1A | 29412 | -3.6117969 | 3.6117969 |
| ADRA1B | 24173 | -1.7472442 | 1.7472442 |
| ADRA1D | 29413 | -2.5322816 | 2.5322816 |
| ADSS | 289276 | 0.4811795 | -0.4811795 |

| | | | |
|---|---|---|---|
| AEN | 361594 | -0.62921613 | 0.62921613 |
| AFAP1L1 | 291565 | -0.63077354 | 0.63077354 |
| AFTPH | 305544 | -0.60534877 | 0.60534877 |
| AGBL5 | 362710 | 0.69131386 | -0.69131386 |
| AGFG1 | 363266 | -0.32273915 | 0.32273915 |
| AGFG2 | 304375 | 0.84993124 | -0.84993124 |
| AGPAT4 | 170919 | 0.29892495 | -0.29892495 |
| AGTR1A | 24180 | -2.2749994 | 2.2749994 |
| AHCYL2 | 312192 | -0.7150247 | 0.7150247 |
| AHI1 | 308923 | -0.50456995 | 0.50456995 |
| AIFM1 | 83533 | 0.48263925 | -0.48263925 |
| AIG1 | 292486 | 0.9889844 | -0.9889844 |
| AIPL1 | 59110 | -1.7069412 | 1.7069412 |
| AJAP1 | 687031 | -0.8164796 | 0.8164796 |
| AK1 | 24183 | 1.2302434 | -1.2302434 |
| AK4 | 29223 | -1.5553226 | 1.5553226 |
| AKAP7 | 361458 | 0.8108324 | -0.8108324 |
| AKT2 | 25233 | -0.7264136 | 0.7264136 |
| ALDH1B1 | 298079 | 1.5108378 | -1.5108378 |
| ALG9 | 367083 | 0.6862229 | -0.6862229 |
| ALKBH4 | 288587 | -0.537583 | 0.537583 |
| ALOXE3 | 287424 | -1.671209 | 1.671209 |
| AMN | 314459 | 3.0723894 | -3.0723894 |
| AMOTL2 | 65157 | 0.9606952 | -0.9606952 |
| AMPD2 | 362015 | 0.99161434 | -0.99161434 |
| AMZ2 | 360650 | 0.91489744 | -0.91489744 |
| ANAPC2 | 296558 | 1.145698 | -1.145698 |
| ANAPC7 | 304490 | 0.78994304 | -0.78994304 |
| ANGPT2 | 89805 | -1.492057 | 1.492057 |
| ANK1 | 306570 | -0.7242867 | 0.7242867 |
| ANKEF1 | 296184 | -1.1365056 | 1.1365056 |
| ANKMY2 | 314046 | 0.89643574 | -0.89643574 |
| ANKRD12 | 316775 | 0.84804845 | -0.84804845 |
| ANKRD13D | 361699 | 1.2342212 | -1.2342212 |
| ANKRD33B | 310200 | -3.6498084 | 3.6498084 |
| ANKRD34A | 295283 | 1.2890335 | -1.2890335 |
| ANKRD34C | 300889 | -2.4182699 | 2.4182699 |
| ANKRD52 | 362811 | -1.0459763 | 1.0459763 |
| ANKRD6 | 500430 | 1.0373784 | -1.0373784 |
| ANKS1B | 314721 | -0.758562 | 0.758562 |
| ANO1 | 309135 | -2.3566136 | 2.3566136 |
| ANO4 | 299714 | -1.6268474 | 1.6268474 |
| ANO6 | 315272 | -0.7700067 | 0.7700067 |

| | | | |
|---|---|---|---|
| ANXA11 | 290527 | -1.4498796 | 1.4498796 |
| ANXA7 | 155423 | -0.34464207 | 0.34464207 |
| AP1S3 | 367304 | -1.3531628 | 1.3531628 |
| AP3S2 | 683402 | 1.3042969 | -1.3042969 |
| APBA2 | 83610 | -0.557084 | 0.557084 |
| APBB1 | 29722 | 1.1575422 | -1.1575422 |
| APC2 | 299611 | 1.1566436 | -1.1566436 |
| APCS | 29339 | -2.2406497 | 2.2406497 |
| APOLD1 | 444983 | -1.2710346 | 1.2710346 |
| AQP11 | 286758 | 0.8479076 | -0.8479076 |
| ARC | 54323 | -3.9109151 | 3.9109151 |
| AREG | 29183 | -3.612455 | 3.612455 |
| ARFGAP3 | 503165 | 0.5375412 | -0.5375412 |
| ARFIP1 | 60382 | -1.0706038 | 1.0706038 |
| ARHGAP10 | 688429 | -0.851316 | 0.851316 |
| ARHGAP17 | 63994 | -0.656478 | 0.656478 |
| ARHGAP20 | 367085 | 0.9537013 | -0.9537013 |
| ARHGAP25 | 500246 | -1.0276208 | 1.0276208 |
| ARHGEF25 | 314904 | 1.3536028 | -1.3536028 |
| ARHGEF28 | 361882 | 1.0131143 | -1.0131143 |
| ARHGEF3 | 290541 | -1.7099833 | 1.7099833 |
| ARID5A | 316327 | -1.8440592 | 1.8440592 |
| ARIH1 | 300756 | -0.68741125 | 0.68741125 |
| ARL14EP | 311279 | -0.94357884 | 0.94357884 |
| ARMC7 | 287827 | 0.30938548 | -0.30938548 |
| ARMCX6 | 363496 | 1.8307469 | -1.8307469 |
| ARPP21 | 363153 | -0.39693156 | 0.39693156 |
| ARRDC4 | 293019 | 0.89794666 | -0.89794666 |
| ARSJ | 311013 | -2.640662 | 2.640662 |
| ASAP1 | 314961 | -1.2466823 | 1.2466823 |
| ASCC1 | 294512 | 1.1025828 | -1.1025828 |
| ASCC2 | 498402 | -0.7461903 | 0.7461903 |
| ASCC3 | 309887 | -0.6759984 | 0.6759984 |
| ASGR1 | 24210 | -2.110998 | 2.110998 |
| ASUN | 690728 | -0.6417649 | 0.6417649 |
| ATF1 | 315305 | -0.5402889 | 0.5402889 |
| ATG10 | 688555 | 1.2703956 | -1.2703956 |
| ATG16L1 | 363278 | -0.6789589 | 0.6789589 |
| ATMIN | 315037 | -0.43551126 | 0.43551126 |
| ATP11A | 306600 | -1.0544498 | 1.0544498 |
| ATP13A3 | 678704 | -1.1816852 | 1.1816852 |
| ATP1B1 | 25650 | -0.36451632 | 0.36451632 |
| ATP2B1 | 29598 | -0.34199724 | 0.34199724 |

| | | | |
|---|---|---|---|
| ATP5SL | 361520 | 0.81120443 | -0.81120443 |
| ATP6V0A2 | 116455 | -1.0298127 | 1.0298127 |
| ATPAF2 | 303190 | 1.1358901 | -1.1358901 |
| ATRAID | 298841 | 1.178951 | -1.178951 |
| AUH | 361215 | 1.0459806 | -1.0459806 |
| B3GNT2 | 305571 | 0.6440816 | -0.6440816 |
| B3GNT7 | 316583 | 1.7183048 | -1.7183048 |
| B4GALT7 | 364675 | 1.2259122 | -1.2259122 |
| BAALC | 140720 | -0.8724806 | 0.8724806 |
| BACH1 | 304127 | -1.7171786 | 1.7171786 |
| BAD | 64639 | 1.2983373 | -1.2983373 |
| BAG3 | 293524 | -2.8785598 | 2.8785598 |
| BAI1 | 362931 | -0.7593267 | 0.7593267 |
| BAIAP2 | 117542 | -1.7085128 | 1.7085128 |
| BANP | 292064 | 0.29416624 | -0.29416624 |
| BAZ1A | 314126 | -1.0645523 | 1.0645523 |
| BBC3 | 317673 | 1.5297897 | -1.5297897 |
| BBS2 | 113948 | 0.7999286 | -0.7999286 |
| BBX | 303970 | -2.2494485 | 2.2494485 |
| BCAS3 | 363662 | 1.1168277 | -1.1168277 |
| BCKDHA | 25244 | 0.8446007 | -0.8446007 |
| BCL11A | 305589 | 0.9717859 | -0.9717859 |
| BCOR | 317346 | -0.53614026 | 0.53614026 |
| BCORL1 | 302810 | 0.8902297 | -0.8902297 |
| BDNF | 24225 | -1.5365307 | 1.5365307 |
| BEND5 | 362564 | 1.4132191 | -1.4132191 |
| BEND7 | 361275 | -1.3499432 | 1.3499432 |
| BHLHE22 | 365748 | 2.9591675 | -2.9591675 |
| BHLHE23 | 499952 | -1.4622635 | 1.4622635 |
| BHLHE40 | 79431 | -2.205643 | 2.205643 |
| BID | 64625 | 0.8444431 | -0.8444431 |
| BLES03 | 266609 | 1.273569 | -1.273569 |
| BLOC1S5 | 306868 | 0.92540556 | -0.92540556 |
| BLOC1S6 | 317630 | 0.49731073 | -0.49731073 |
| BMPER | 300455 | -1.1944522 | 1.1944522 |
| BNIP3 | 84480 | 1.0849622 | -1.0849622 |
| BNIP3L | 140923 | 1.0528262 | -1.0528262 |
| BOK | 29884 | 1.6824747 | -1.6824747 |
| BPHL | 361239 | 1.2076544 | -1.2076544 |
| BRE | 362704 | 1.037709 | -1.037709 |
| BRINP3 | 286901 | 1.3785894 | -1.3785894 |
| BROX | 305031 | 0.6399443 | -0.6399443 |
| BSDC1 | 297890 | 0.70258904 | -0.70258904 |

| | | | |
|---|---|---|---|
| BTBD6 | 690367 | 0.77760196 | -0.77760196 |
| BTG4 | 315650 | 1.8256004 | -1.8256004 |
| BTRC | 361765 | -0.5605399 | 0.5605399 |
| BVES | 365603 | -1.4005638 | 1.4005638 |
| C1QL2 | 288979 | -3.3434925 | 3.3434925 |
| C1QL3 | 680404 | -1.6164814 | 1.6164814 |
| C1S | 192262 | -1.7718594 | 1.7718594 |
| C2CD2L | 300666 | 0.6377958 | -0.6377958 |
| CABLES1 | 307585 | 0.7776117 | -0.7776117 |
| CACNB1 | 50688 | -0.37143254 | 0.37143254 |
| CACNB3 | 25297 | 1.0428525 | -1.0428525 |
| CACNB4 | 58942 | -1.2364125 | 1.2364125 |
| CACNG3 | 140724 | -1.3195261 | 1.3195261 |
| CALCA | 24241 | -2.2412775 | 2.2412775 |
| CALCOCO1 | 246047 | 0.6982477 | -0.6982477 |
| CALCRL | 25029 | -1.2262877 | 1.2262877 |
| CALHM2 | 294019 | 0.9394895 | -0.9394895 |
| CALY | 192349 | 1.3429091 | -1.3429091 |
| CAMK1G | 171358 | -0.5225028 | 0.5225028 |
| CAMKK1 | 60341 | -0.6206306 | 0.6206306 |
| CAPN2 | 29154 | -0.51820725 | 0.51820725 |
| CAPN5 | 171495 | 1.1776319 | -1.1776319 |
| CARF | 301446 | 0.7784706 | -0.7784706 |
| CARS | 293638 | -0.6864734 | 0.6864734 |
| CAT | 24248 | 0.48430485 | -0.48430485 |
| CAV1 | 25404 | -1.3746773 | 1.3746773 |
| CBFA2T3 | 361431 | -1.1455648 | 1.1455648 |
| CBFB | 361391 | -0.62609076 | 0.62609076 |
| CBLN1 | 498922 | 1.3264753 | -1.3264753 |
| CBLN2 | 291388 | -0.9794108 | 0.9794108 |
| CBX8 | 303731 | 0.7328807 | -0.7328807 |
| CCBL2 | 541589 | 1.2280177 | -1.2280177 |
| CCDC129 | 500139 | -1.8191351 | 1.8191351 |
| CCDC148 | 311051 | 1.48715 | -1.48715 |
| CCDC175 | 500668 | 1.6173832 | -1.6173832 |
| CCDC184 | 500925 | 1.306872 | -1.306872 |
| CCDC28A | 361454 | 2.0075464 | -2.0075464 |
| CCDC28B | 682445 | 1.71312 | -1.71312 |
| CCDC32 | 296081 | 0.83603305 | -0.83603305 |
| CCDC40 | 287867 | 1.0490103 | -1.0490103 |
| CCDC53 | 299707 | 1.4643389 | -1.4643389 |
| CCDC86 | 293738 | -1.2498329 | 1.2498329 |
| CCDC92 | 100036765 | -0.5050791 | 0.5050791 |

| | | | |
|---|---|---|---|
| CCDC93 | 304743 | -0.6232316 | 0.6232316 |
| CCKAR | 24889 | 2.4626114 | -2.4626114 |
| CCKBR | 25706 | -2.2215497 | 2.2215497 |
| CCND1 | 58919 | -1.5370443 | 1.5370443 |
| CCNG2 | 29157 | 1.8686941 | -1.8686941 |
| CCNL2 | 298686 | -0.5288063 | 0.5288063 |
| CCPG1 | 363098 | 1.1234567 | -1.1234567 |
| CD200 | 24560 | 1.1759683 | -1.1759683 |
| CD226 | 307199 | 1.8789576 | -1.8789576 |
| CD55 | 64036 | 0.7351284 | -0.7351284 |
| CD59 | 25407 | 1.0753154 | -1.0753154 |
| CDC14A | 310806 | -1.7185223 | 1.7185223 |
| CDC25B | 171103 | 0.9233059 | -0.9233059 |
| CDC37L1 | 293886 | -0.5121295 | 0.5121295 |
| CDC42EP3 | 313838 | 0.95548755 | -0.95548755 |
| CDC42EP4 | 303653 | 1.3727965 | -1.3727965 |
| CDCA4 | 500727 | -0.63640547 | 0.63640547 |
| CDCA7L | 619566 | -0.6370459 | 0.6370459 |
| CDH15 | 361432 | 1.6700618 | -1.6700618 |
| CDH7 | 29162 | 1.4516338 | -1.4516338 |
| CDK19 | 309804 | 0.8173471 | -0.8173471 |
| CDK5 | 140908 | 1.1217455 | -1.1217455 |
| CDK5RAP1 | 252827 | 1.0830865 | -1.0830865 |
| CDK5RAP3 | 80278 | 1.1755166 | -1.1755166 |
| CDK8 | 498140 | -0.36754665 | 0.36754665 |
| CDKN2AIPNL | 287278 | 0.944174 | -0.944174 |
| CDO1 | 81718 | 1.284292 | -1.284292 |
| CDYL | 361237 | -1.3846971 | 1.3846971 |
| CDYL2 | 292044 | -1.3537384 | 1.3537384 |
| CELF6 | 300758 | 0.9918951 | -0.9918951 |
| CEP57L1 | 294519 | 0.5916561 | -0.5916561 |
| CFB | 294257 | -1.421618 | 1.421618 |
| CFLAR | 117279 | -1.0501424 | 1.0501424 |
| CH25H | 309527 | -1.1801852 | 1.1801852 |
| CHD1 | 308215 | -1.0756745 | 1.0756745 |
| CHD6 | 311607 | 0.6874034 | -0.6874034 |
| CHML | 689102 | -2.2738986 | 2.2738986 |
| CHMP1A | 365024 | 0.81924915 | -0.81924915 |
| CHMP6 | 287873 | 1.1243142 | -1.1243142 |
| CHN2 | 84031 | 1.0361619 | -1.0361619 |
| CHODL | 288289 | 2.231705 | -2.231705 |
| CHRAC1 | 315058 | 0.6425566 | -0.6425566 |
| CHRM2 | 81645 | -1.6941352 | 1.6941352 |

| | | | |
|---|---|---|---|
| CHRNA5 | 25102 | 1.1824613 | -1.1824613 |
| CHST10 | 140568 | -0.43606564 | 0.43606564 |
| CHST12 | 304322 | 1.2780493 | -1.2780493 |
| CHST7 | 302302 | -0.5012247 | 0.5012247 |
| CHSY1 | 292999 | -0.34165925 | 0.34165925 |
| CHURC1 | 299154 | 1.9833574 | -1.9833574 |
| CIR1 | 362149 | 1.5603967 | -1.5603967 |
| CISD3 | 287661 | 1.9258878 | -1.9258878 |
| CISH | 83681 | 1.444542 | -1.444542 |
| CITED2 | 114490 | -1.593563 | 1.593563 |
| CLASP2 | 114514 | 0.70147526 | -0.70147526 |
| CLDN9 | 287099 | 1.4729203 | -1.4729203 |
| CLDND1 | 288182 | -0.4175044 | 0.4175044 |
| CLIP2 | 29264 | 1.1156156 | -1.1156156 |
| CLMP | 286939 | 1.8699203 | -1.8699203 |
| CLN5 | 306128 | -0.8822822 | 0.8822822 |
| CLN8 | 306619 | 0.57413274 | -0.57413274 |
| CMAS | 312826 | 1.2177485 | -1.2177485 |
| CMPK2 | 314004 | 0.8633549 | -0.8633549 |
| CMTR2 | 292016 | -1.0163047 | 1.0163047 |
| CNGA4 | 85258 | -3.4251666 | 3.4251666 |
| CNNM1 | 309387 | -0.75169325 | 0.75169325 |
| CNNM2 | 294014 | 0.48656547 | -0.48656547 |
| CNP | 25275 | 0.96667814 | -0.96667814 |
| CNR1 | 25248 | 0.5381837 | -0.5381837 |
| CNTFR | 313173 | -0.86567485 | 0.86567485 |
| CNTN2 | 25356 | 1.4346097 | -1.4346097 |
| CNTN3 | 54279 | 1.2686888 | -1.2686888 |
| COASY | 287711 | 0.91764677 | -0.91764677 |
| COBL | 305497 | 0.6472813 | -0.6472813 |
| COG3 | 361073 | -0.77866966 | 0.77866966 |
| COL26A1 | 685612 | 1.0989481 | -1.0989481 |
| COL4A1 | 290905 | -0.5921858 | 0.5921858 |
| COL5A1 | 85490 | -1.4546968 | 1.4546968 |
| COMMD1 | 289831 | 1.6570169 | -1.6570169 |
| COMTD1 | 305685 | 1.2443414 | -1.2443414 |
| COX11 | 690300 | -0.71707284 | 0.71707284 |
| COX19 | 304330 | 1.2216917 | -1.2216917 |
| CPNE4 | 367160 | 1.0142878 | -1.0142878 |
| CPNE8 | 362988 | 1.2241803 | -1.2241803 |
| CRAT | 311849 | 0.6730089 | -0.6730089 |
| CREBL2 | 362453 | 0.47197583 | -0.47197583 |
| CREM | 25620 | -1.4372749 | 1.4372749 |

| | | | |
|---|---|---|---|
| CRHBP | 29625 | -3.949865 | 3.949865 |
| CRIP2 | 338401 | 1.1930181 | -1.1930181 |
| CRIPT | 56725 | 1.1421247 | -1.1421247 |
| CRLF3 | 54395 | -0.7444929 | 0.7444929 |
| CROT | 83842 | -0.85425067 | 0.85425067 |
| CRY1 | 299691 | -0.6343994 | 0.6343994 |
| CRYM | 117024 | 1.5057176 | -1.5057176 |
| CSPG5 | 50568 | 0.853645 | -0.853645 |
| CSRNP1 | 363165 | -1.3948185 | 1.3948185 |
| CSRP2 | 29317 | 1.7276167 | -1.7276167 |
| CTDP1 | 291414 | -0.30255198 | 0.30255198 |
| CTPS2 | 619580 | 0.95587426 | -0.95587426 |
| CTSF | 361704 | 1.2331759 | -1.2331759 |
| CUBN | 80848 | -2.3821952 | 2.3821952 |
| CUL9 | 316228 | 1.498927 | -1.498927 |
| CUX2 | 288665 | 0.9874247 | -0.9874247 |
| CX3CL1 | 89808 | -0.6751452 | 0.6751452 |
| CXCR4 | 60628 | 0.8277072 | -0.8277072 |
| CXXC5 | 291670 | -0.8889998 | 0.8889998 |
| CYB561 | 303601 | 1.338949 | -1.338949 |
| CYB561D2 | 363137 | 1.6831125 | -1.6831125 |
| CYB5R1 | 304805 | 1.3150365 | -1.3150365 |
| CYGB | 170520 | 1.8016915 | -1.8016915 |
| CYHR1 | 100362155.chr | 1.5363264 | -1.5363264 |
| CYLD | 312937 | 0.6384103 | -0.6384103 |
| CYP11B2 | 24294 | -1.7526295 | 1.7526295 |
| CYP11B3 | 353498 | -2.2118537 | 2.2118537 |
| CYP26B1 | 312495 | -3.2631261 | 3.2631261 |
| CYP27B1 | 114700 | -2.6242201 | 2.6242201 |
| CYR61 | 83476 | 2.1667793 | -2.1667793 |
| CYS1 | 690489 | 1.237013 | -1.237013 |
| CYTH1 | 116691 | 0.5620195 | -0.5620195 |
| D2HGDH | 301624 | 1.048223 | -1.048223 |
| DAAM1 | 314212 | 1.1973464 | -1.1973464 |
| DARS2 | 304919 | 0.78181463 | -0.78181463 |
| DBNDD1 | 361437 | 2.8623035 | -2.8623035 |
| DBP | 24309 | 1.0060977 | -1.0060977 |
| DCAF10 | 313242 | -0.7000545 | 0.7000545 |
| DCAF4 | 362762 | 1.1153011 | -1.1153011 |
| DCAKD | 360639 | 0.59539396 | -0.59539396 |
| DCLK1 | 83825 | -0.5291546 | 0.5291546 |
| DCLK3 | 316023 | 3.1585078 | -3.1585078 |
| DCPS | 266605 | 0.740661 | -0.740661 |

| | | | |
|---|---|---|---|
| DDAH2 | 294239 | 1.7449005 | -1.7449005 |
| DDHD1 | 305816 | -0.571644 | 0.571644 |
| DDIT4 | 140942 | 0.8063096 | -0.8063096 |
| DDX19B | 690693 | -0.27219203 | 0.27219203 |
| DDX20 | 84473 | -0.47132653 | 0.47132653 |
| DDX21 | 317399 | -1.0781173 | 1.0781173 |
| DDX3X | 317335 | -0.42627296 | 0.42627296 |
| DEAF1 | 83632 | 0.9790304 | -0.9790304 |
| DEF6 | 309642 | 1.9787971 | -1.9787971 |
| DEPDC5 | 305464 | 0.60728824 | -0.60728824 |
| DES | 64362 | -1.3894205 | 1.3894205 |
| DESI1 | 315160 | -0.3931853 | 0.3931853 |
| DGCR8 | 287954 | -0.29063565 | 0.29063565 |
| DGUOK | 297389 | 0.75320053 | -0.75320053 |
| DHRS9 | 170635 | -4.409934 | 4.409934 |
| DHX40 | 287595 | 0.8696716 | -0.8696716 |
| DHX57 | 366532 | 0.9110711 | -0.9110711 |
| DIAPH1 | 307483 | -0.6938537 | 0.6938537 |
| DIEXF | 305076 | -0.45094988 | 0.45094988 |
| DIP2C | 307067 | -1.248641 | 1.248641 |
| DIRAS1 | 366826 | 1.1724926 | -1.1724926 |
| DIRC2 | 303902 | 0.7612408 | -0.7612408 |
| DIS3 | 306103 | -0.55803543 | 0.55803543 |
| DISC1 | 307940 | -1.470493 | 1.470493 |
| DISP2 | 311324 | 0.9088202 | -0.9088202 |
| DKK2 | 295445 | -4.432964 | 4.432964 |
| DLC1 | 58834 | -2.8727405 | 2.8727405 |
| DLEU7 | 290308 | 1.9654785 | -1.9654785 |
| DLG1 | 25252 | -1.559255 | 1.559255 |
| DLGAP2 | 116681 | -3.2651875 | 3.2651875 |
| DLK1 | 114587 | -1.0597112 | 1.0597112 |
| DMP1 | 25312 | -3.0781653 | 3.0781653 |
| DNAJB4 | 295549 | 0.95473754 | -0.95473754 |
| DNAJB5 | 313811 | -0.76502573 | 0.76502573 |
| DNAJC25 | 362526 | -0.40392032 | 0.40392032 |
| DNASE2B | 59296 | -3.7200184 | 3.7200184 |
| DOC2A | 65031 | 1.2574142 | -1.2574142 |
| DOCK8 | 499337 | -1.8718106 | 1.8718106 |
| DOK4 | 361364 | 1.2097769 | -1.2097769 |
| DOK5 | 502694 | -1.1789211 | 1.1789211 |
| DOT1L | 362831 | -1.4931031 | 1.4931031 |
| DPF1 | 50545 | 1.4777873 | -1.4777873 |
| DPF3 | 299186 | -1.2533493 | 1.2533493 |

| | | | |
|---|---|---|---|
| DPM2 | 29640 | 1.2595621 | -1.2595621 |
| DPY19L1 | 315496 | -0.47533405 | 0.47533405 |
| DPYD | 81656 | 0.5710019 | -0.5710019 |
| DPYSL2 | 25416 | 1.0006807 | -1.0006807 |
| DRD5 | 25195 | -5.998091 | 5.998091 |
| DSE | 365583 | -0.54123944 | 0.54123944 |
| DSG3 | 291752 | -3.3121614 | 3.3121614 |
| DTX3 | 500847 | 0.92782456 | -0.92782456 |
| DUOX1 | 266807 | -1.1840279 | 1.1840279 |
| DUSP14 | 360580 | -1.9785913 | 1.9785913 |
| DUSP23 | 360881 | 1.6762038 | -1.6762038 |
| DUSP26 | 306527 | 1.5668025 | -1.5668025 |
| DUSP4 | 60587 | -2.832405 | 2.832405 |
| DUSP5 | 171109 | -2.4208353 | 2.4208353 |
| DUSP6 | 116663 | -1.7100483 | 1.7100483 |
| DYNC1I1 | 29564 | 0.8546949 | -0.8546949 |
| EBF1 | 116543 | 1.8131684 | -1.8131684 |
| EBF3 | 361668 | -1.4570917 | 1.4570917 |
| EBPL | 361054 | 1.8555979 | -1.8555979 |
| ECEL1 | 60417 | -2.5625746 | 2.5625746 |
| ECH1 | 64526 | 0.8102297 | -0.8102297 |
| EDNRA | 24326 | -1.5468034 | 1.5468034 |
| EEA1 | 314764 | -1.3007274 | 1.3007274 |
| EEF2K | 25435 | 0.9492289 | -0.9492289 |
| EFNA4 | 310643 | 1.9769019 | -1.9769019 |
| EFNB3 | 360546 | 1.1232499 | -1.1232499 |
| EFR3A | 362923 | -1.2757814 | 1.2757814 |
| EGR2 | 114090 | -1.3127321 | 1.3127321 |
| EGR3 | 25148 | -2.6038425 | 2.6038425 |
| EHBP1 | 305556 | 0.5023931 | -0.5023931 |
| EHBP1L1 | 309169 | 2.03378 | -2.03378 |
| EHD1 | 293692 | 1.2024988 | -1.2024988 |
| EHD4 | 192204 | -1.3532994 | 1.3532994 |
| EHF | 295965 | -2.370194 | 2.370194 |
| EIF1B | 301068 | 1.0278952 | -1.0278952 |
| EIF2B5 | 192234 | 0.2963045 | -0.2963045 |
| EIF5A | 287444 | 0.4837144 | -0.4837144 |
| ELAVL4 | 432358 | 1.4632306 | -1.4632306 |
| ELF3 | 304815 | 2.784469 | -2.784469 |
| ELMOD1 | 315670 | 1.4567773 | -1.4567773 |
| ELOVL7 | 361895 | -2.4834602 | 2.4834602 |
| ELP2 | 307545 | 0.6777394 | -0.6777394 |
| EMC9 | 290224 | 2.015691 | -2.015691 |

| | | | |
|---|---|---|---|
| EMILIN3 | 362262 | -1.2687697 | 1.2687697 |
| EML1 | 362783 | 0.8281443 | -0.8281443 |
| EML4 | 313861 | -0.95312595 | 0.95312595 |
| EML5 | 444982 | -1.1021036 | 1.1021036 |
| EMP1 | 25314 | -4.5941815 | 4.5941815 |
| EMP3 | 81505 | -1.6464361 | 1.6464361 |
| ENTPD6 | 85260 | 0.9032556 | -0.9032556 |
| EPB41L4B | 500464 | -0.9424875 | 0.9424875 |
| EPDR1 | 291180 | -0.7201802 | 0.7201802 |
| EPHA3 | 29210 | 1.0821818 | -1.0821818 |
| EPHA4 | 316539 | 0.9911312 | -0.9911312 |
| EPHB6 | 312275 | 1.5996736 | -1.5996736 |
| ERAL1 | 363646 | 0.8207574 | -0.8207574 |
| ERCC1 | 292673 | -0.49968156 | 0.49968156 |
| ERRFI1 | 313729 | -1.0401924 | 1.0401924 |
| ESR1 | 24890 | -1.1356808 | 1.1356808 |
| ETHE1 | 292710 | 1.3022593 | -1.3022593 |
| ETV4 | 360635 | -0.70073 | 0.70073 |
| ETV5 | 303828 | -2.085397 | 2.085397 |
| EVA1A | 500221 | -0.9362583 | 0.9362583 |
| EXO1 | 305000 | -0.83079153 | 0.83079153 |
| EXOC3L1 | 291961 | 1.5982518 | -1.5982518 |
| EXOC5 | 60627 | -0.7554035 | 0.7554035 |
| EXTL2 | 310803 | 0.46723768 | -0.46723768 |
| EZR | 54319 | -1.5817354 | 1.5817354 |
| F2R | 25439 | -0.67870164 | 0.67870164 |
| FADS3 | 286922 | 0.8076848 | -0.8076848 |
| FAIM2 | 246274 | 1.2188671 | -1.2188671 |
| FAM102B | 365903 | -0.8933741 | 0.8933741 |
| FAM107B | 498796 | -0.6215519 | 0.6215519 |
| FAM110C | 500638 | -4.870246 | 4.870246 |
| FAM110D | 500563 | 2.551216 | -2.551216 |
| FAM118A | 300120 | 1.0343237 | -1.0343237 |
| FAM127B | 679038 | 0.98202944 | -0.98202944 |
| FAM129B | 362115 | -0.3663069 | 0.3663069 |
| FAM129C | 498604 | 1.7675214 | -1.7675214 |
| FAM134A | 363252 | 0.820052 | -0.820052 |
| FAM136A | 297415 | 1.1968178 | -1.1968178 |
| FAM150B | 679566 | -4.36002 | 4.36002 |
| FAM163B | 685169 | -0.50279844 | 0.50279844 |
| FAM184B | 289671 | 2.1667516 | -2.1667516 |
| FAM192A | 307652 | 0.84725934 | -0.84725934 |
| FAM196A | 100233213 | -1.6162378 | 1.6162378 |

| | | | |
|---|---|---|---|
| FAM212B | 310764 | -2.0356653 | 2.0356653 |
| FAM214A | 300836 | 1.1667423 | -1.1667423 |
| FAM220A | 498145 | 0.8113638 | -0.8113638 |
| FAM222B | 497960 | -0.9767736 | 0.9767736 |
| FAM26F | 294430 | -2.4521663 | 2.4521663 |
| FAM43A | 288031 | -1.4174006 | 1.4174006 |
| FAM65B | 306934 | -0.95911074 | 0.95911074 |
| FAM84A | 313969 | -1.5026602 | 1.5026602 |
| FAR1 | 293173 | -1.3329574 | 1.3329574 |
| FARS2 | 306879 | 1.2801592 | -1.2801592 |
| FBN2 | 689008 | -0.93448967 | 0.93448967 |
| FBXL14 | 312675 | -0.8897135 | 0.8897135 |
| FBXL20 | 64039 | 0.5463844 | -0.5463844 |
| FBXL6 | 362941 | 1.4751589 | -1.4751589 |
| FBXO16 | 305970 | 1.7636634 | -1.7636634 |
| FBXO31 | 498959 | 1.4363247 | -1.4363247 |
| FBXO41 | 312504 | 1.2192515 | -1.2192515 |
| FBXO44 | 500587 | 1.0990988 | -1.0990988 |
| FBXW17 | 361219 | 1.3822906 | -1.3822906 |
| FCHO1 | 290639 | 1.3593366 | -1.3593366 |
| FDFT1 | 29580 | 1.176812 | -1.176812 |
| FDXR | 79122 | 1.5488063 | -1.5488063 |
| FEZ1 | 81730 | 1.0974953 | -1.0974953 |
| FFAR4 | 294075 | -2.5157187 | 2.5157187 |
| FGF13 | 84488 | 1.0222241 | -1.0222241 |
| FGF2 | 54250 | -4.242713 | 4.242713 |
| FGF3 | 170633 | -2.5571735 | 2.5571735 |
| FGF8 | 29349 | -2.0306342 | 2.0306342 |
| FGF9 | 25444 | -1.3539819 | 1.3539819 |
| FGFR1 | 79114 | -2.3136618 | 2.3136618 |
| FHL2 | 63839 | -0.5822633 | 0.5822633 |
| FIBP | 282837 | 1.2658006 | -1.2658006 |
| FIG4 | 309855 | 0.70847756 | -0.70847756 |
| FILIP1 | 246776 | 2.7372005 | -2.7372005 |
| FJX1 | 366140 | -0.46838948 | 0.46838948 |
| FKBP5 | 361810 | -0.6876135 | 0.6876135 |
| FLOT1 | 64665 | 1.4665278 | -1.4665278 |
| FLOT2 | 83764 | 1.4728571 | -1.4728571 |
| FLRT1 | 499308 | -1.623491 | 1.623491 |
| FLT1 | 54251 | -2.632133 | 2.632133 |
| FMNL1 | 287746 | -0.7881743 | 0.7881743 |
| FN3KRP | 303755 | 0.49876475 | -0.49876475 |
| FNDC3A | 306022 | -1.3759376 | 1.3759376 |

| | | | |
|---|---|---|---|
| FNDC3B | 294925 | -1.8846078 | 1.8846078 |
| FOXP2 | 500037 | 1.0561723 | -1.0561723 |
| FRA10AC1 | 365458 | 1.5293473 | -1.5293473 |
| FRAS1 | 289486 | -1.3616799 | 1.3616799 |
| FRMD3 | 298141 | 0.5577625 | -0.5577625 |
| FRMD6 | 257646 | -1.4485581 | 1.4485581 |
| FRS3 | 316213 | 1.4993623 | -1.4993623 |
| FST | 24373 | 3.1059115 | -3.1059115 |
| FSTL5 | 365823 | 0.41745064 | -0.41745064 |
| FTSJ3 | 303608 | -0.36455974 | 0.36455974 |
| FUCA2 | 292485 | 1.1626443 | -1.1626443 |
| FUNDC1 | 363442 | 0.85060465 | -0.85060465 |
| FURIN | 54281 | -0.75249976 | 0.75249976 |
| FUT7 | 296564 | -1.479171 | 1.479171 |
| FXYD6 | 63847 | 1.0193477 | -1.0193477 |
| FYTTD1 | 360726 | -0.31878003 | 0.31878003 |
| FZD1 | 58868 | 0.79249483 | -0.79249483 |
| FZD4 | 64558 | -1.4714973 | 1.4714973 |
| FZR1 | 314642 | 1.0390644 | -1.0390644 |
| GABARAPL1 | 689161 | 0.7775922 | -0.7775922 |
| GABRD | 29689 | 1.7964844 | -1.7964844 |
| GABRG2 | 29709 | 1.161652 | -1.161652 |
| GABRQ | 65187 | -0.915536 | 0.915536 |
| GADD45B | 299626 | -1.4834048 | 1.4834048 |
| GAL | 29141 | -2.946706 | 2.946706 |
| GALE | 114860 | 1.5647572 | -1.5647572 |
| GALK2 | 296117 | 1.3194399 | -1.3194399 |
| GALNT3 | 366061 | 0.69954914 | -0.69954914 |
| GALNT7 | 29750 | -1.491607 | 1.491607 |
| GALNT9 | 304571 | -1.1128696 | 1.1128696 |
| GALR1 | 50577 | -6.854205 | 6.854205 |
| GAS7 | 85246 | 0.69908094 | -0.69908094 |
| GBX1 | 246149 | -1.8644722 | 1.8644722 |
| GCGR | 24953 | -1.8537114 | 1.8537114 |
| GCK | 24385 | 4.057229 | -4.057229 |
| GCLC | 25283 | -1.5143988 | 1.5143988 |
| GCNT1 | 64043 | -3.3693295 | 3.3693295 |
| GDNF | 25453 | -1.8685989 | 1.8685989 |
| GEM | 297902 | -1.464599 | 1.464599 |
| GEMIN8 | 363462 | 1.0965458 | -1.0965458 |
| GFOD1 | 306842 | -1.8363347 | 1.8363347 |
| GFPT2 | 360518 | 1.2477304 | -1.2477304 |
| GFRA1 | 25454 | -2.5710554 | 2.5710554 |

| | | | |
|---|---|---|---|
| GINS2 | 292058 | -0.7468228 | 0.7468228 |
| GJB2 | 394266 | -1.0402259 | 1.0402259 |
| GLDN | 315675 | -0.80972207 | 0.80972207 |
| GLI2 | 304729 | -1.8104296 | 1.8104296 |
| GLRA2 | 24397 | 0.9108136 | -0.9108136 |
| GLRB | 25456 | 0.6576773 | -0.6576773 |
| GLRX | 64045 | 0.938423 | -0.938423 |
| GMPPA | 501167 | 1.1785016 | -1.1785016 |
| GOLGB1 | 192243 | 0.6012115 | -0.6012115 |
| GOLPH3L | 310669 | 0.26587063 | -0.26587063 |
| GPANK1 | 415064 | 1.29448 | -1.29448 |
| GPAT3 | 305166 | -1.7013336 | 1.7013336 |
| GPATCH2 | 289362 | -1.0378615 | 1.0378615 |
| GPC2 | 171517 | 1.5365086 | -1.5365086 |
| GPM6A | 306439 | 0.86119384 | -0.86119384 |
| GPR101 | 317608 | -2.4965632 | 2.4965632 |
| GPR12 | 80840 | 2.111301 | -2.111301 |
| GPR139 | 293545 | 1.1281028 | -1.1281028 |
| GPR156 | 260430 | 0.9838025 | -0.9838025 |
| GPR158 | 291352 | -3.9495723 | 3.9495723 |
| GPR21 | 311911 | 0.32613963 | -0.32613963 |
| GPR26 | 192153 | -1.0960113 | 1.0960113 |
| GPR3 | 266769 | -1.151957 | 1.151957 |
| GPR37 | 117549 | 1.8543845 | -1.8543845 |
| GPR4 | 308408 | -0.7456858 | 0.7456858 |
| GPR45 | 301372 | 0.54334664 | -0.54334664 |
| GPR50 | 117097 | -2.2457151 | 2.2457151 |
| GPR52 | 684623 | 1.0648395 | -1.0648395 |
| GPR61 | 310780 | 1.0142778 | -1.0142778 |
| GPR63 | 297952 | -2.409319 | 2.409319 |
| GPR68 | 314386 | -2.2744858 | 2.2744858 |
| GPR83 | 140595 | -2.6879358 | 2.6879358 |
| GPR85 | 64020 | 0.8875574 | -0.8875574 |
| GPR88 | 64443 | -0.5728938 | 0.5728938 |
| GPRASP1 | 171407 | 0.37424657 | -0.37424657 |
| GPRC5A | 312790 | -5.1008306 | 5.1008306 |
| GPRC5C | 287805 | 1.4053327 | -1.4053327 |
| GPX3 | 64317 | 1.0541209 | -1.0541209 |
| GRASP | 192254 | -2.925807 | 2.925807 |
| GREM1 | 50566 | -1.078983 | 1.078983 |
| GRHL2 | 299979 | -1.4455799 | 1.4455799 |
| GRIN2A | 24409 | -3.1377962 | 3.1377962 |
| GRIP1 | 84016 | -0.62972784 | 0.62972784 |

| | | | |
|---|---|---|---|
| GRK5 | 59075 | -0.7295949 | 0.7295949 |
| GRM1 | 24414 | 1.7649225 | -1.7649225 |
| GRM6 | 24419 | 1.5778718 | -1.5778718 |
| GRN | 29143 | 0.6929331 | -0.6929331 |
| GRP | 171101 | 1.7887181 | -1.7887181 |
| GRWD1 | 308592 | -0.5673416 | 0.5673416 |
| GSPT2 | 501582 | -0.37514183 | 0.37514183 |
| GTF2IRD1 | 246770 | 0.80117756 | -0.80117756 |
| GTF3C5 | 362095 | 0.5371806 | -0.5371806 |
| GTPBP4 | 114300 | -1.0845741 | 1.0845741 |
| GUCY1A3 | 497757 | 1.2316636 | -1.2316636 |
| GUCY1B3 | 25202 | 1.2535505 | -1.2535505 |
| GYPC | 364837 | -3.377353 | 3.377353 |
| GZF1 | 311508 | -0.41640782 | 0.41640782 |
| H2AFY2 | 361844 | 1.0627307 | -1.0627307 |
| HACE1 | 361866 | -1.2985991 | 1.2985991 |
| HAS1 | 282821 | -2.8278217 | 2.8278217 |
| HAUS8 | 290626 | -2.3493516 | 2.3493516 |
| HBEGF | 25433 | -0.8350807 | 0.8350807 |
| HCN1 | 84390 | -0.9801577 | 0.9801577 |
| HCN3 | 114245 | 1.0103825 | -1.0103825 |
| HDAC11 | 297453 | 1.309217 | -1.309217 |
| HDX | 317617 | -1.6189289 | 1.6189289 |
| HEATR5B | 362683 | 0.72567636 | -0.72567636 |
| HECA | 308624 | -0.65091056 | 0.65091056 |
| HECTD2 | 309514 | -2.1175406 | 2.1175406 |
| HEPHL1 | 500946 | -1.8817065 | 1.8817065 |
| HERC3 | 362377 | 0.5495569 | -0.5495569 |
| HERPUD2 | 300463 | 0.3756691 | -0.3756691 |
| HES6 | 316626 | 1.6718974 | -1.6718974 |
| HIC1 | 303310 | -1.6527004 | 1.6527004 |
| HIGD2A | 290999 | 1.5173984 | -1.5173984 |
| HIST3H2A | 64646 | 1.7466613 | -1.7466613 |
| HMG20A | 315689 | 0.5274166 | -0.5274166 |
| HMGA1 | 117062 | -2.1395297 | 2.1395297 |
| HMGA2 | 84017 | -2.8688316 | 2.8688316 |
| HMGCLL1 | 367112 | 1.3452446 | -1.3452446 |
| HOMER1 | 29546 | -1.7807477 | 1.7807477 |
| HOMER2 | 29547 | -0.7872916 | 0.7872916 |
| HP1BP3 | 313647 | 0.68039024 | -0.68039024 |
| HPCAL1 | 50871 | 1.0417874 | -1.0417874 |
| HPN | 29135 | -1.9175618 | 1.9175618 |
| HRH1 | 24448 | -1.7427148 | 1.7427148 |

| | | | |
|---|---|---|---|
| HRH3 | 85268 | 2.7061095 | -2.7061095 |
| HRK | 117271 | 0.6865796 | -0.6865796 |
| HS3ST5 | 294449 | -0.7514898 | 0.7514898 |
| HSD11B1 | 25116 | -0.91425896 | 0.91425896 |
| HSDL1 | 361418 | 0.47044277 | -0.47044277 |
| HSF2BP | 499413 | -3.7208776 | 3.7208776 |
| HSH2D | 100360518 | -2.35337 | 2.35337 |
| HSPA12A | 307997 | 0.68739927 | -0.68739927 |
| HSPA1B | 294254 | 1.3913642 | -1.3913642 |
| HSPA4L | 294993 | -1.1560411 | 1.1560411 |
| HSPB11 | 685284 | 1.5743185 | -1.5743185 |
| HSPH1 | 288444 | -0.590423 | 0.590423 |
| HTR1B | 25075 | -3.1142626 | 3.1142626 |
| HTR1F | 60448 | -3.0416794 | 3.0416794 |
| HTR7 | 65032 | -0.6663122 | 0.6663122 |
| ICA1 | 81024 | 0.96159065 | -0.96159065 |
| ICK | 84411 | 0.7886756 | -0.7886756 |
| IDH1 | 24479 | 0.74111575 | -0.74111575 |
| IER5 | 498256 | -1.1308113 | 1.1308113 |
| IER5L | 499772 | -0.9994531 | 0.9994531 |
| IFFO2 | 641315 | -1.5317856 | 1.5317856 |
| IFNG | 25712 | -2.047485 | 2.047485 |
| IFNGR2 | 360697 | 0.80174 | -0.80174 |
| IFRD1 | 29596 | -0.88495636 | 0.88495636 |
| IFT122 | 312651 | 0.8522669 | -0.8522669 |
| IFT52 | 362265 | 0.8561634 | -0.8561634 |
| IGF1 | 24482 | -1.4307919 | 1.4307919 |
| IGF2BP2 | 303824 | -1.0935881 | 1.0935881 |
| IGSF3 | 295325 | 1.2240644 | -1.2240644 |
| IGSF8 | 304979 | 1.394032 | -1.394032 |
| IGSF9 | 304982 | 2.1925702 | -2.1925702 |
| IL1RAP | 25466 | -1.0855343 | 1.0855343 |
| IL1RL2 | 171106 | -2.4500093 | 2.4500093 |
| IL25 | 501996 | 1.6658764 | -1.6658764 |
| IL34 | 498951 | 2.079161 | -2.079161 |
| IL6R | 24499 | -2.765288 | 2.765288 |
| IMPDH1 | 362329 | 1.1701523 | -1.1701523 |
| ING4 | 297597 | 1.3283275 | -1.3283275 |
| INHBA | 29200 | -2.3986232 | 2.3986232 |
| INIP | 298032 | -0.40450808 | 0.40450808 |
| INO80C | 291737 | 0.64258486 | -0.64258486 |
| INPP5D | 54259 | -1.4566199 | 1.4566199 |
| INSC | 293166 | -3.3949947 | 3.3949947 |

| | | | |
|---|---|---|---|
| IPPK | 306808 | -1.4250371 | 1.4250371 |
| IQSEC3 | 404781 | -0.8191139 | 0.8191139 |
| IRF6 | 364081 | -1.33301 | 1.33301 |
| IRGQ | 292708 | 0.5293753 | -0.5293753 |
| IRS2 | 29376 | -1.7454215 | 1.7454215 |
| IRX6 | 307715 | -1.503687 | 1.503687 |
| ISG20L2 | 361977 | -0.82030123 | 0.82030123 |
| ISM1 | 311760 | -3.1640146 | 3.1640146 |
| ISOC1 | 364879 | 0.45082656 | -0.45082656 |
| ITGA11 | 315744 | 1.8450263 | -1.8450263 |
| ITGA6 | 114517 | -2.3379338 | 2.3379338 |
| ITM2C | 301575 | 1.0857115 | -1.0857115 |
| ITPR3 | 25679 | -1.2132822 | 1.2132822 |
| ITSN1 | 29491 | -0.89033645 | 0.89033645 |
| IVD | 24513 | 0.46442395 | -0.46442395 |
| IWS1 | 291705 | -0.4862273 | 0.4862273 |
| JADE3 | 299305 | -1.4989351 | 1.4989351 |
| JAK1 | 84598 | -0.6616703 | 0.6616703 |
| JAK2 | 24514 | -1.0800953 | 1.0800953 |
| JAKMIP1 | 305434 | 1.5035542 | -1.5035542 |
| JAKMIP3 | 365380 | 0.5178917 | -0.5178917 |
| JPH3 | 307916 | 0.8579227 | -0.8579227 |
| JPH4 | 445271 | 0.83087945 | -0.83087945 |
| JUNB | 24517 | 1.1961887 | -1.1961887 |
| KATNBL1 | 691543 | -0.8469377 | 0.8469377 |
| KBTBD2 | 312372 | -0.7062145 | 0.7062145 |
| KCNA1 | 24520 | -2.9324117 | 2.9324117 |
| KCNA4 | 25469 | -1.1384053 | 1.1384053 |
| KCNB1 | 25736 | -0.9069853 | 0.9069853 |
| KCNC4 | 684516 | -1.2006938 | 1.2006938 |
| KCND2 | 65180 | 0.85068125 | -0.85068125 |
| KCNE2 | 171138 | 0.578719 | -0.578719 |
| KCNF1 | 298908 | -1.5706642 | 1.5706642 |
| KCNH3 | 27150 | 1.2644694 | -1.2644694 |
| KCNH4 | 114032 | 2.7187452 | -2.7187452 |
| KCNJ12 | 117052 | -0.89960426 | 0.89960426 |
| KCNJ4 | 116649 | -0.8559248 | 0.8559248 |
| KCNJ5 | 29713 | -1.992862 | 1.992862 |
| KCNK1 | 59324 | -0.40121496 | 0.40121496 |
| KCNK2 | 170899 | 2.3072448 | -2.3072448 |
| KCNK3 | 29553 | -1.1693228 | 1.1693228 |
| KCNMB2 | 294961 | -1.3542088 | 1.3542088 |
| KCNMB4 | 66016 | 1.8512713 | -1.8512713 |

| | | | |
|---|---|---|---|
| KCNN1 | 54261 | 1.5103672 | -1.5103672 |
| KCNN3 | 54263 | -1.9845036 | 1.9845036 |
| KCNV1 | 60326 | -1.0496781 | 1.0496781 |
| KCTD15 | 499129 | 0.79915696 | -0.79915696 |
| KCTD6 | 305792 | -0.46940184 | 0.46940184 |
| KDM1A | 500569 | -0.24557911 | 0.24557911 |
| KDM4A | 313539 | 0.65337425 | -0.65337425 |
| KDSR | 360833 | 0.5790338 | -0.5790338 |
| KIF27 | 246209 | 0.9834024 | -0.9834024 |
| KIFAP3 | 289168 | 0.73230666 | -0.73230666 |
| KITLG | 60427 | -1.4576013 | 1.4576013 |
| KL | 83504 | -3.6032693 | 3.6032693 |
| KLC4 | 316226 | 1.3974475 | -1.3974475 |
| KLF10 | 81813 | -2.4324164 | 2.4324164 |
| KLF11 | 313994 | 2.2614818 | -2.2614818 |
| KLF13 | 499171 | 0.9078538 | -0.9078538 |
| KLF14 | 312203 | -2.951865 | 2.951865 |
| KLF2 | 306330 | -1.3621902 | 1.3621902 |
| KLF5 | 84410 | -0.84676117 | 0.84676117 |
| KLF7 | 363243 | 1.1907438 | -1.1907438 |
| KLF9 | 117560 | -1.7787198 | 1.7787198 |
| KLHDC8A | 305096 | 3.1098526 | -3.1098526 |
| KLHDC8B | 306589 | 1.5225165 | -1.5225165 |
| KLHL14 | 364823 | 0.937737 | -0.937737 |
| KLHL23 | 311114 | 0.75821215 | -0.75821215 |
| KLHL24 | 303803 | 0.91934127 | -0.91934127 |
| KLHL25 | 293023 | 0.5308118 | -0.5308118 |
| KLHL29 | 298867 | -0.89261734 | 0.89261734 |
| KLHL36 | 498957 | 1.0984741 | -1.0984741 |
| KLHL5 | 305351 | 0.54883945 | -0.54883945 |
| KLHL7 | 362303 | 0.7690857 | -0.7690857 |
| KMT2E | 311968 | 0.7637932 | -0.7637932 |
| KRAS | 24525 | -0.9396105 | 0.9396105 |
| LAMB3 | 305078 | -1.7006404 | 1.7006404 |
| LAMP5 | 362220 | -2.8400226 | 2.8400226 |
| LANCL2 | 362375 | 0.60344684 | -0.60344684 |
| LASP1 | 29278 | 0.83576113 | -0.83576113 |
| LBH | 683626 | -1.3358486 | 1.3358486 |
| LCK | 313050 | 1.7219892 | -1.7219892 |
| LDAH | 313949 | 0.5557382 | -0.5557382 |
| LDB1 | 309447 | 0.9988345 | -0.9988345 |
| LDLRAD4 | 679578 | -0.878956 | 0.878956 |
| LEF1 | 161452 | -2.2935052 | 2.2935052 |

| | | | |
|---|---|---|---|
| LENG8 | 361506 | -1.1925628 | 1.1925628 |
| LEXM | 500516 | -4.3647046 | 4.3647046 |
| LGI1 | 252892 | 1.1382487 | -1.1382487 |
| LHFPL2 | 294643 | -1.3977135 | 1.3977135 |
| LHX2 | 296706 | 1.1847873 | -1.1847873 |
| LHX4 | 360858 | -0.7624696 | 0.7624696 |
| LIMK2 | 29524 | 1.1858875 | -1.1858875 |
| LINGO3 | 690755 | -0.95841247 | 0.95841247 |
| LIPA | 25055 | 0.5183348 | -0.5183348 |
| LIPG | 291437 | -1.3595645 | 1.3595645 |
| LIX1 | 292381 | 1.0929558 | -1.0929558 |
| LMNA | 60374 | -1.3383297 | 1.3383297 |
| LMO1 | 245979 | 1.6234221 | -1.6234221 |
| LMO2 | 362176 | -1.4608102 | 1.4608102 |
| LMO4 | 362051 | 1.6763406 | -1.6763406 |
| LOC100125362 | 100125362 | -0.4107643 | 0.4107643 |
| LOC100125367 | 100125367 | -0.791909 | 0.791909 |
| LOC100302465 | 100302465 | -2.2428775 | 2.2428775 |
| LOC100910945 | 100910945 | 1.6606487 | -1.6606487 |
| LOC314140 | 314140 | -0.47866273 | 0.47866273 |
| LOC500877 | 500877 | 1.3496305 | -1.3496305 |
| LOC680039 | 680039 | -0.6987722 | 0.6987722 |
| LOC680663 | 680663 | -1.9476616 | 1.9476616 |
| LOC682102 | 682102 | 1.5073361 | -1.5073361 |
| LOC688390 | 688390 | 1.0159336 | -1.0159336 |
| LOC688452 | 688452 | -1.316452 | 1.316452 |
| LOC688459 | 688459 | -3.1551375 | 3.1551375 |
| LOC688765 | 688765 | -0.62610525 | 0.62610525 |
| LPAR1 | 116744 | -3.2895625 | 3.2895625 |
| LPCAT2 | 100359680 | -0.8129258 | 0.8129258 |
| LPL | 24539 | 0.85419005 | -0.85419005 |
| LRCH1 | 502020 | -0.8170938 | 0.8170938 |
| LRFN5 | 314164 | 1.121322 | -1.121322 |
| LRRC10B | 309208 | -1.2030287 | 1.2030287 |
| LRRC16B | 361041 | 1.1732049 | -1.1732049 |
| LRRC17 | 502715 | -0.84104425 | 0.84104425 |
| LRRC20 | 499430 | 1.2454054 | -1.2454054 |
| LRRC26 | 311803 | -2.5454128 | 2.5454128 |
| LRRC28 | 361588 | 0.92711407 | -0.92711407 |
| LRRC34 | 499589 | 1.666489 | -1.666489 |
| LRRC4 | 641521 | -0.7702187 | 0.7702187 |
| LRRC42 | 298309 | 0.95530796 | -0.95530796 |
| LRRC55 | 311171 | 0.8532531 | -0.8532531 |

| | | | |
|---|---|---|---|
| LRRC56 | 365389 | 1.6173358 | -1.6173358 |
| LRRC6 | 299920 | 0.894351 | -0.894351 |
| LRRC61 | 500111 | 1.6301228 | -1.6301228 |
| LRRC71 | 310689 | 1.3037688 | -1.3037688 |
| LRRC8B | 305135 | -1.5113169 | 1.5113169 |
| LRRN1 | 500280 | 0.79205996 | -0.79205996 |
| LRRN3 | 81514 | 1.1927134 | -1.1927134 |
| LRRTM3 | 294380 | -0.68070257 | 0.68070257 |
| LRTM2 | 680883 | 1.338333 | -1.338333 |
| LTBP1 | 59107 | -0.5354814 | 0.5354814 |
| LUZP1 | 79428 | -0.898674 | 0.898674 |
| LYPD1 | 360838 | 0.981903 | -0.981903 |
| LYPLA2 | 83510 | 0.44875744 | -0.44875744 |
| LYRM7 | 686506 | 1.144265 | -1.144265 |
| LYRM9 | 497962 | 1.8536752 | -1.8536752 |
| LYVE1 | 293186 | -1.9220386 | 1.9220386 |
| LZIC | 366507 | 0.5858828 | -0.5858828 |
| LZTFL1 | 316102 | 0.6239684 | -0.6239684 |
| LZTS1 | 266711 | 1.459883 | -1.459883 |
| M6PR | 312689 | -0.55362326 | 0.55362326 |
| MAF1 | 315093 | 1.2809612 | -1.2809612 |
| MAFK | 246760 | 0.609026 | -0.609026 |
| MAGED2 | 113947 | 1.1529831 | -1.1529831 |
| MAGEE2 | 302392 | 1.4546404 | -1.4546404 |
| MAL2 | 362911 | 1.1839454 | -1.1839454 |
| MAML3 | 310405 | -2.5122805 | 2.5122805 |
| MAN1A1 | 294410 | -2.6563742 | 2.6563742 |
| MAN1A2 | 295319 | -1.0048935 | 1.0048935 |
| MAN1C1 | 362625 | -1.4619563 | 1.4619563 |
| MAN2A1 | 25478 | -1.7447019 | 1.7447019 |
| MAP2K3 | 303200 | -1.8593556 | 1.8593556 |
| MAP2K4 | 287398 | -0.89620936 | 0.89620936 |
| MAP2K5 | 29568 | 0.8882246 | -0.8882246 |
| MAP2K6 | 114495 | 1.7449335 | -1.7449335 |
| MAP3K12 | 25579 | 1.0383974 | -1.0383974 |
| MAP3K7 | 313121 | -0.4686874 | 0.4686874 |
| MAP7D1 | 681287 | -0.6060024 | 0.6060024 |
| MAPK15 | 286997 | 2.5421202 | -2.5421202 |
| MAPK1IP1 | 499280 | 0.7967403 | -0.7967403 |
| MAPK3 | 50689 | 1.2152724 | -1.2152724 |
| MAS1 | 25153 | -1.5918338 | 1.5918338 |
| MAT2B | 683630 | 0.6810566 | -0.6810566 |
| MB21D2 | 498100 | 0.7683288 | -0.7683288 |

| | | | |
|---|---|---|---|
| MBLAC2 | 365627 | 0.65403515 | -0.65403515 |
| MC5R | 25726 | -1.2832022 | 1.2832022 |
| MCCC2 | 361884 | 0.7789034 | -0.7789034 |
| MCEE | 293829 | 1.3414524 | -1.3414524 |
| MCHR1 | 83567 | -1.8251122 | 1.8251122 |
| MCL1 | 60430 | -0.5577267 | 0.5577267 |
| MCM5 | 291885 | -0.33948404 | 0.33948404 |
| MCOLN3 | 308022 | -1.3112947 | 1.3112947 |
| ME1 | 24552 | -0.7285156 | 0.7285156 |
| MED15 | 360743 | -0.7941409 | 0.7941409 |
| MED4 | 306030 | 0.919518 | -0.919518 |
| MED9 | 497914 | 1.1482514 | -1.1482514 |
| MEIS2 | 311311 | -1.1404159 | 1.1404159 |
| MELTF | 288038 | 1.8393661 | -1.8393661 |
| MEPCE | 304361 | 0.7829449 | -0.7829449 |
| MEST | 58827 | -0.84213364 | 0.84213364 |
| METRNL | 316842 | -1.0411627 | 1.0411627 |
| METTL18 | 304928 | -0.6321552 | 0.6321552 |
| MFAP2 | 313662 | 0.541674 | -0.541674 |
| MFAP3L | 306424 | -2.319295 | 2.319295 |
| MFN1 | 192647 | -0.5755169 | 0.5755169 |
| MFSD13A | 309454 | 0.64404565 | -0.64404565 |
| MFSD14B | 306687 | -0.8376799 | 0.8376799 |
| MFSD6 | 301388 | 0.665314 | -0.665314 |
| MFSD9 | 316356 | 0.5732246 | -0.5732246 |
| MGAT1 | 81519 | 0.5889136 | -0.5889136 |
| MGAT5B | 303693 | 1.1863223 | -1.1863223 |
| MICAL2 | 365352 | -0.9250694 | 0.9250694 |
| MICU3 | 364601 | -0.73718745 | 0.73718745 |
| MINK1 | 303259 | -0.5068592 | 0.5068592 |
| MKRN1 | 296988 | 0.7501028 | -0.7501028 |
| MLLT3 | 114510 | -0.9280919 | 0.9280919 |
| MMP10 | 117061 | -4.506247 | 4.506247 |
| MMP13 | 171052 | -5.4765487 | 5.4765487 |
| MMP17 | 288626.chr12.2 | -0.74054533 | 0.74054533 |
| MMP3 | 171045 | -3.3737705 | 3.3737705 |
| MMS19 | 171124 | 0.6874268 | -0.6874268 |
| MNT | 287521 | 0.9935537 | -0.9935537 |
| MON2 | 314894 | -0.6843731 | 0.6843731 |
| MORN4 | 293950 | 1.3041744 | -1.3041744 |
| MOSPD3 | 288557 | 0.751187 | -0.751187 |
| MPDU1 | 303244 | 1.1387473 | -1.1387473 |
| MPEG1 | 64552 | 0.9923391 | -0.9923391 |

| | | | |
|---|---|---|---|
| MPPED1 | 362971 | 1.5517403 | -1.5517403 |
| MPV17L2 | 290645 | 1.1072813 | -1.1072813 |
| MRM1 | 363661 | -0.40253547 | 0.40253547 |
| MRPL10 | 691075 | 0.88254374 | -0.88254374 |
| MRPL14 | 301250 | 1.6324209 | -1.6324209 |
| MRPL34 | 290632 | 1.77923 | -1.77923 |
| MRPS12 | 292758 | 1.1417923 | -1.1417923 |
| MRRF | 311903 | 0.69935894 | -0.69935894 |
| MSL3L2 | 309790 | -1.0766865 | 1.0766865 |
| MSRB1 | 685059 | 1.255364 | -1.255364 |
| MSTN | 29152 | 1.3471953 | -1.3471953 |
| MTERF2 | 366856 | 1.3015225 | -1.3015225 |
| MTHFD2 | 680308 | -0.6058509 | 0.6058509 |
| MTMR12 | 310155 | -2.155846 | 2.155846 |
| MTPAP | 307050 | -0.4685971 | 0.4685971 |
| MTSS1 | 362918 | 1.1206813 | -1.1206813 |
| MTUS2 | 498136 | 2.035879 | -2.035879 |
| MUSK | 81725 | -5.454042 | 5.454042 |
| MUSTN1 | 290553 | -0.9738472 | 0.9738472 |
| MVK | 81727 | 1.5795974 | -1.5795974 |
| MXD4 | 360961 | 0.5827962 | -0.5827962 |
| MXI1 | 25701 | 0.50135267 | -0.50135267 |
| MYADML2 | 303744 | 2.5961335 | -2.5961335 |
| MYBPH | 83708 | -0.956229 | 0.956229 |
| MYC | 24577 | -0.7247556 | 0.7247556 |
| MYCL | 298506 | 2.5626357 | -2.5626357 |
| MYH1 | 287408 | -2.2530081 | 2.2530081 |
| MYH4 | 360543 | -3.5680916 | 3.5680916 |
| MYH7 | 29557 | 1.8198395 | -1.8198395 |
| MYO1E | 25484 | -2.8696394 | 2.8696394 |
| MYO9B | 25486 | -0.92271817 | 0.92271817 |
| MYRIP | 360034 | -1.1458832 | 1.1458832 |
| MYT1L | 116668 | 1.4418831 | -1.4418831 |
| N4BP2L1 | 498131 | 0.57142353 | -0.57142353 |
| NAA40 | 361718 | -0.3950268 | 0.3950268 |
| NAAA | 497009 | 1.1311554 | -1.1311554 |
| NAB2 | 314910 | -2.149525 | 2.149525 |
| NAGA | 315165 | 0.841882 | -0.841882 |
| NAMPT | 297508 | -1.0371566 | 1.0371566 |
| NAP1L3 | 170914 | 0.95103115 | -0.95103115 |
| NAPEPLD | 296757 | -1.5130607 | 1.5130607 |
| NARF | 360681 | 1.2755058 | -1.2755058 |
| NAT14 | 361500 | 1.6346843 | -1.6346843 |

| | | | |
|---|---|---|---|
| NBR1 | 303554 | 0.380894 | -0.380894 |
| NCALD | 553106 | 0.59072584 | -0.59072584 |
| NCBP1 | 298075 | -0.4643829 | 0.4643829 |
| NCBP3 | 360563 | -0.32995352 | 0.32995352 |
| NCOR2 | 360801 | -0.64690495 | 0.64690495 |
| NCSTN | 289231 | 0.35748908 | -0.35748908 |
| NDRG4 | 64457 | 0.98793954 | -0.98793954 |
| NDST3 | 295430 | -1.1042091 | 1.1042091 |
| NDST4 | 362035 | 1.3141888 | -1.3141888 |
| NDUFAF4 | 362495 | -0.99508625 | 0.99508625 |
| NEDD9 | 291044 | -0.98134995 | 0.98134995 |
| NEFL | 83613 | -1.707625 | 1.707625 |
| NET1 | 307098 | 0.47926953 | -0.47926953 |
| NETO1 | 307206 | -1.0864983 | 1.0864983 |
| NEUROD1 | 29458 | 1.0511507 | -1.0511507 |
| NEUROD2 | 54276 | 0.49624836 | -0.49624836 |
| NEUROD4 | 288821 | 0.92741853 | -0.92741853 |
| NFATC1 | 100361818 | -1.4382105 | 1.4382105 |
| NFIB | 29227 | 0.49739763 | -0.49739763 |
| NFKBIE | 316241 | 0.68254673 | -0.68254673 |
| NFRKB | 315523 | -0.55807513 | 0.55807513 |
| NFX1 | 313166 | -0.36331108 | 0.36331108 |
| NGB | 85382 | 1.2327213 | -1.2327213 |
| NGF | 310738 | -1.8514184 | 1.8514184 |
| NHLH2 | 295327 | 1.1201491 | -1.1201491 |
| NIPSNAP1 | 360971 | 1.297048 | -1.297048 |
| NIPSNAP3B | 313211 | 1.4569368 | -1.4569368 |
| NIT1 | 289222 | 0.8626085 | -0.8626085 |
| NKAP | 298342 | 1.2555282 | -1.2555282 |
| NKD1 | 364952 | 1.1703247 | -1.1703247 |
| NKIRAS2 | 287707 | 0.8874032 | -0.8874032 |
| NMBR | 25264 | -0.97207475 | 0.97207475 |
| NME7 | 171566 | 0.61801004 | -0.61801004 |
| NOC3L | 361753 | -0.6718607 | 0.6718607 |
| NOCT | 310395 | -0.38256145 | 0.38256145 |
| NOG | 25495 | -0.9140755 | 0.9140755 |
| NOL10 | 313981 | -0.3505987 | 0.3505987 |
| NOL3 | 85383 | 1.4650979 | -1.4650979 |
| NOP2 | 314969 | 0.28176972 | -0.28176972 |
| NOSIP | 292894 | 1.5772415 | -1.5772415 |
| NOTCH4 | 406162 | 1.0245112 | -1.0245112 |
| NOV | 81526 | 1.4443339 | -1.4443339 |
| NPAP60 | 25497 | -0.4197655 | 0.4197655 |

| | | | |
|---|---|---|---|
| NPL | 304860 | -2.4390168 | 2.4390168 |
| NPLOC4 | 140639 | -0.4270526 | 0.4270526 |
| NPTX1 | 266777 | -2.6545897 | 2.6545897 |
| NPTX2 | 288475 | -3.609085 | 3.609085 |
| NPY1R | 29358 | 1.4898225 | -1.4898225 |
| NPY2R | 66024 | -1.3297244 | 1.3297244 |
| NR1D1 | 252917 | 0.50560164 | -0.50560164 |
| NR1D2 | 259241 | 0.41467774 | -0.41467774 |
| NR1I3 | 65035 | 0.6986578 | -0.6986578 |
| NR2F1 | 81808 | 1.0604424 | -1.0604424 |
| NRF1 | 312195 | -0.95736855 | 0.95736855 |
| NRIP1 | 304157 | 1.0291498 | -1.0291498 |
| NRIP3 | 361625 | -0.4573317 | 0.4573317 |
| NRN1 | 83834 | -2.2758472 | 2.2758472 |
| NRSN2 | 689978 | 1.8389419 | -1.8389419 |
| NTF3 | 81737 | 1.4229482 | -1.4229482 |
| NTM | 50864 | 1.2541375 | -1.2541375 |
| NTRK3 | 29613 | -0.6188653 | 0.6188653 |
| NUAK1 | 299694 | 0.9856098 | -0.9856098 |
| NUB1 | 296731 | 0.8141831 | -0.8141831 |
| NUDT14 | 299346 | 1.9057776 | -1.9057776 |
| NUDT16 | 363129 | 1.4501662 | -1.4501662 |
| NUDT18 | 361068 | 1.2897798 | -1.2897798 |
| NUDT6 | 207120 | -1.108387 | 1.108387 |
| NUTM1 | 366153 | -1.0077661 | 1.0077661 |
| NVL | 289323 | -0.4274949 | 0.4274949 |
| NYAP1 | 304376 | 2.2479792 | -2.2479792 |
| OGDHL | 290566 | 0.8859162 | -0.8859162 |
| OGFOD1 | 307657 | -1.2811823 | 1.2811823 |
| OLIG2 | 304103 | -0.64515185 | 0.64515185 |
| OPRL1 | 29256 | 1.2142957 | -1.2142957 |
| ORAI2 | 304592 | 1.5564501 | -1.5564501 |
| OS9 | 362891 | 0.8949342 | -0.8949342 |
| OSBP2 | 305475 | -1.1393825 | 1.1393825 |
| OSBPL1A | 259221 | 0.4714191 | -0.4714191 |
| OSBPL2 | 296461 | 0.5098632 | -0.5098632 |
| OSBPL6 | 311129 | -1.4557711 | 1.4557711 |
| OSCAR | 292537 | -1.0710164 | 1.0710164 |
| OSCP1 | 362595 | 0.8735821 | -0.8735821 |
| OTUD3 | 500572 | -1.2368755 | 1.2368755 |
| OVCA2 | 497954 | 0.8804624 | -0.8804624 |
| OXTR | 25342 | -1.5087687 | 1.5087687 |
| P2RX2 | 114115 | 1.5780302 | -1.5780302 |

| | | | |
|---|---|---|---|
| P2RY1 | 25265 | 0.70958847 | -0.70958847 |
| P2RY14 | 171108 | -3.1473017 | 3.1473017 |
| P4HA1 | 64475 | -0.59373695 | 0.59373695 |
| PACSIN1 | 29704 | -0.7012393 | 0.7012393 |
| PACSIN2 | 124461 | -0.6784627 | 0.6784627 |
| PAFAH1B3 | 114113 | 1.4949573 | -1.4949573 |
| PAK6 | 296078 | -0.74153435 | 0.74153435 |
| PAK7 | 311450 | 0.50949806 | -0.50949806 |
| PALMD | 310811 | 1.181603 | -1.181603 |
| PAM | 25508 | -1.4187125 | 1.4187125 |
| PANX1 | 315435 | -1.0286161 | 1.0286161 |
| PAPOLA | 314417 | -0.50829095 | 0.50829095 |
| PAPPA | 313262 | -2.3755753 | 2.3755753 |
| PARD6A | 307799 | 1.2980179 | -1.2980179 |
| PATL1 | 361736 | -0.7010471 | 0.7010471 |
| PAX1 | 311505 | -1.1285974 | 1.1285974 |
| PBX1 | 304947 | 0.7519762 | -0.7519762 |
| PCBP4 | 363133 | 0.9265226 | -0.9265226 |
| PCDH19 | 317183 | 1.8977797 | -1.8977797 |
| PCDH8 | 64865 | -1.3071531 | 1.3071531 |
| PCDHA10 | 116778 | 0.8411842 | -0.8411842 |
| PCDHA12 | 116779 | 0.84128857 | -0.84128857 |
| PCDHA2 | 393086 | 0.8769241 | -0.8769241 |
| PCDHA3 | 116780 | 0.68591297 | -0.68591297 |
| PCDHA4 | 116741 | 0.67511904 | -0.67511904 |
| PCDHA5 | 393087 | 0.6871557 | -0.6871557 |
| PCDHA6 | 393088 | 0.8123535 | -0.8123535 |
| PCDHA7 | 393089 | 0.75973856 | -0.75973856 |
| PCDHA9 | 393090 | 0.5936863 | -0.5936863 |
| PCDHGA2 | 498846 | 0.62522614 | -0.62522614 |
| PCGF2 | 287662 | 1.1421169 | -1.1421169 |
| PCGF5 | 681178 | -0.94097286 | 0.94097286 |
| PCSK1 | 25204 | -1.1705414 | 1.1705414 |
| PCSK4 | 171085 | 1.9182475 | -1.9182475 |
| PDCD6IP | 501083 | -0.48611596 | 0.48611596 |
| PDE10A | 63885 | -1.4955032 | 1.4955032 |
| PDE12 | 306231 | -0.8079652 | 0.8079652 |
| PDE6D | 363272 | 1.5448804 | -1.5448804 |
| PDE8A | 308776 | -3.1037586 | 3.1037586 |
| PDE9A | 191569 | -0.9505414 | 0.9505414 |
| PDGFB | 24628 | -2.7598 | 2.7598 |
| PDK1 | 116551 | -0.314967 | 0.314967 |
| PDP1 | 54705 | -1.9958472 | 1.9958472 |

| | | | |
|---|---|---|---|
| PDSS2 | 365592 | 1.6004679 | -1.6004679 |
| PDXK | 83578 | -2.4952135 | 2.4952135 |
| PDXP | 727679 | 1.1974871 | -1.1974871 |
| PDZD8 | 308000 | -0.95551646 | 0.95551646 |
| PEAK1 | 315686 | -2.5138175 | 2.5138175 |
| PECAM1 | 29583 | 1.6324714 | -1.6324714 |
| PENK | 29237 | -2.9786656 | 2.9786656 |
| PEO1 | 309441 | 1.0950004 | -1.0950004 |
| PEX14 | 64460 | 1.0501068 | -1.0501068 |
| PFKFB3 | 117276 | -2.1468453 | 2.1468453 |
| PGD | 100360180 | 0.7458782 | -0.7458782 |
| PHACTR1 | 306844 | 0.7627509 | -0.7627509 |
| PHACTR3 | 362284 | 0.9712113 | -0.9712113 |
| PHEX | 25512 | -3.0974095 | 3.0974095 |
| PHF14 | 500030 | 0.48963156 | -0.48963156 |
| PHF20 | 311575 | 0.6254803 | -0.6254803 |
| PHLDA1 | 29380 | -1.0356759 | 1.0356759 |
| PHOSPHO2 | 295663 | -0.24552187 | 0.24552187 |
| PI16 | 294312 | 1.547693 | -1.547693 |
| PIANP | 312711 | 1.1342764 | -1.1342764 |
| PIEZO1 | 361430 | -1.1661532 | 1.1661532 |
| PIGM | 79112 | -1.5399154 | 1.5399154 |
| PIH1D1 | 292898 | 1.2580909 | -1.2580909 |
| PIK3C3 | 65052 | 1.0280429 | -1.0280429 |
| PIK3CB | 85243 | -0.8965338 | 0.8965338 |
| PIK3CD | 366508 | 2.0016377 | -2.0016377 |
| PIK3IP1 | 305472 | 1.0467699 | -1.0467699 |
| PITX1 | 113983 | -3.433762 | 3.433762 |
| PJA2 | 192256 | 0.38071728 | -0.38071728 |
| PKD2 | 498328 | -1.0872481 | 1.0872481 |
| PKLR | 24651 | 2.0960937 | -2.0960937 |
| PKNOX1 | 294322 | -0.9983487 | 0.9983487 |
| PKNOX2 | 680549 | 0.7530365 | -0.7530365 |
| PLA2G4A | 24653 | -1.7562926 | 1.7562926 |
| PLAGL1 | 25157 | -0.4995203 | 0.4995203 |
| PLAUR | 50692 | -4.892946 | 4.892946 |
| PLCL1 | 84587 | -2.184532 | 2.184532 |
| PLCXD2 | 363781 | 1.5329875 | -1.5329875 |
| PLEKHG5 | 310999 | -1.0774326 | 1.0774326 |
| PLEKHH2 | 313866 | -2.63918 | 2.63918 |
| PLK2 | 83722 | -0.8687869 | 0.8687869 |
| PLK4 | 310344 | -0.31353673 | 0.31353673 |
| PLPPR3 | 314614 | 1.3086072 | -1.3086072 |

| | | | |
|---|---|---|---|
| PLPPR4 | 295401 | -0.35939395 | 0.35939395 |
| PLXNA2 | 289392 | -1.0976771 | 1.0976771 |
| PLXND1 | 312652 | 0.890478 | -0.890478 |
| PMP22 | 24660 | -0.6559772 | 0.6559772 |
| PMVK | 310645 | 1.8580743 | -1.8580743 |
| PNCK | 29660 | 1.7745042 | -1.7745042 |
| PNMA2 | 305977 | 1.2323488 | -1.2323488 |
| PNMAL1 | 361515 | 1.4421582 | -1.4421582 |
| PNMAL2 | 308393 | 0.617953 | -0.617953 |
| PNPO | 64533 | 0.6252789 | -0.6252789 |
| PNRC1 | 286988 | 1.235424 | -1.235424 |
| PODXL | 192181 | 0.6308847 | -0.6308847 |
| PODXL2 | 297433 | 1.2327952 | -1.2327952 |
| POLD1 | 59294 | 0.31041476 | -0.31041476 |
| POLE4 | 362385 | 1.2117931 | -1.2117931 |
| POLG | 85472 | -1.0260669 | 1.0260669 |
| POLL | 361767 | 1.2602353 | -1.2602353 |
| POLR1A | 83581 | -0.618907 | 0.618907 |
| POLR3E | 361640 | -0.6538515 | 0.6538515 |
| POMK | 306549 | 0.8079559 | -0.8079559 |
| POP1 | 315045 | -0.6307121 | 0.6307121 |
| POPDC2 | 360718 | 1.4212865 | -1.4212865 |
| POPDC3 | 641520 | -0.6938103 | 0.6938103 |
| POU4F2 | 171355 | -4.1835046 | 4.1835046 |
| POU4F3 | 364855 | -3.2644193 | 3.2644193 |
| POU6F1 | 116545 | -1.1412504 | 1.1412504 |
| PPEF1 | 317498 | 1.2263446 | -1.2263446 |
| PPFIA4 | 140592 | 1.3249093 | -1.3249093 |
| PPHLN1 | 366975 | -0.89656925 | 0.89656925 |
| PPIF | 282819 | -0.40926883 | 0.40926883 |
| PPM1H | 314897 | -0.7404914 | 0.7404914 |
| PPP1CA | 24668 | 0.3484464 | -0.3484464 |
| PPP1R12A | 116670 | -0.4603236 | 0.4603236 |
| PPP1R18 | 361790 | 1.5800347 | -1.5800347 |
| PPP2R2B | 60660 | 1.0847508 | -1.0847508 |
| PPP2R3A | 363122 | 0.93942696 | -0.93942696 |
| PPP2R5A | 312754 | -0.5296165 | 0.5296165 |
| PPP3CC | 171378 | 0.7928479 | -0.7928479 |
| PPP4R3A | 314388 | -0.26442242 | 0.26442242 |
| PPRC1 | 294007 | -0.74539727 | 0.74539727 |
| PRAGMIN | 306506 | -1.5715748 | 1.5715748 |
| PRICKLE1 | 315259 | -1.5160234 | 1.5160234 |
| PRIMA1 | 690195 | -4.5349665 | 4.5349665 |

| | | | |
|---|---|---|---|
| PRKAA1 | 65248 | -0.53383154 | 0.53383154 |
| PRKAR2A | 29699 | -1.7279683 | 1.7279683 |
| PRKCQ | 85420 | 0.6951319 | -0.6951319 |
| PRKCZ | 25522 | 0.98278683 | -0.98278683 |
| PRKG2 | 25523 | -1.4253035 | 1.4253035 |
| PRKRA | 311130 | 1.2576514 | -1.2576514 |
| PRMT2 | 499420 | 1.3451191 | -1.3451191 |
| PRR13 | 363004 | 0.84493697 | -0.84493697 |
| PRR15 | 312358 | -1.0145926 | 1.0145926 |
| PRRT1 | 406167 | 1.2082696 | -1.2082696 |
| PRRT2 | 361651 | 0.9984324 | -0.9984324 |
| PRRT4 | 500059 | 1.8222485 | -1.8222485 |
| PRSS12 | 85266 | 1.6767384 | -1.6767384 |
| PRSS23 | 308807 | -2.5837831 | 2.5837831 |
| PRSS35 | 315866 | -2.0219622 | 2.0219622 |
| PRUNE | 310664 | 0.6687089 | -0.6687089 |
| PSD2 | 307500 | 1.2645555 | -1.2645555 |
| PSENEN | 292788 | 1.3393663 | -1.3393663 |
| PSME3 | 287716 | -0.2551777 | 0.2551777 |
| PSME4 | 498433 | -0.70672363 | 0.70672363 |
| PSMG1 | 288236 | 1.2615643 | -1.2615643 |
| PSTPIP2 | 307248 | -1.2847602 | 1.2847602 |
| PTAFR | 58949 | -4.0518913 | 4.0518913 |
| PTBP1 | 29497 | -0.45423856 | 0.45423856 |
| PTCHD1 | 317517 | -1.8570472 | 1.8570472 |
| PTGER4 | 84023 | -2.125272 | 2.125272 |
| PTGS2 | 29527 | -2.3201041 | 2.3201041 |
| PTH2R | 81753 | -2.445413 | 2.445413 |
| PTPDC1 | 291022 | -0.88963217 | 0.88963217 |
| PTPN1 | 24697 | -0.3797934 | 0.3797934 |
| PTPRE | 114767 | -0.6064726 | 0.6064726 |
| PTPRH | 171125 | 1.9810252 | -1.9810252 |
| PTPRN | 116660 | -0.87433666 | 0.87433666 |
| PTPRR | 94202.chr7.2 | 2.134171 | -2.134171 |
| PURG | 361162 | -0.59971136 | 0.59971136 |
| PUS3 | 315554 | -0.87038594 | 0.87038594 |
| PUS7 | 296751 | -0.7606243 | 0.7606243 |
| PVR | 25066 | -1.5210682 | 1.5210682 |
| PVRL1 | 192183 | -0.618076 | 0.618076 |
| PWWP2B | 361671 | -0.5893987 | 0.5893987 |
| PXN | 360820 | -0.61022323 | 0.61022323 |
| QDPR | 64192 | 1.2607766 | -1.2607766 |
| QPCT | 313837 | 1.2014691 | -1.2014691 |

| | | | |
|---|---|---|---|
| R3HDM1 | 304763 | -0.93854046 | 0.93854046 |
| RAB11B | 79434 | 0.882444 | -0.882444 |
| RAB20 | 689377 | -1.370957 | 1.370957 |
| RAB27B | 84590 | -1.699384 | 1.699384 |
| RAB2B | 305853 | 0.8936181 | -0.8936181 |
| RAB30 | 308821 | 1.1979102 | -1.1979102 |
| RAB33A | 317580 | 2.0812652 | -2.0812652 |
| RAB36 | 690407 | 1.0716302 | -1.0716302 |
| RAB3A | 25531 | 1.0002227 | -1.0002227 |
| RAB3D | 140665 | 1.7104299 | -1.7104299 |
| RAB43 | 500249 | -1.0554436 | 1.0554436 |
| RAB9B | 367915 | 0.6534948 | -0.6534948 |
| RAD21 | 314949 | 0.46561742 | -0.46561742 |
| RAI14 | 294804 | -1.4602149 | 1.4602149 |
| RALGAPA1 | 56785 | -0.4897604 | 0.4897604 |
| RALGDS | 29622 | 0.93801826 | -0.93801826 |
| RALYL | 294883 | 1.3761951 | -1.3761951 |
| RAMP2 | 58966 | 1.5750382 | -1.5750382 |
| RAMP3 | 56820 | 2.9852746 | -2.9852746 |
| RANBP6 | 309326 | -0.20998384 | 0.20998384 |
| RAP1B | 171337 | -0.7228956 | 0.7228956 |
| RASA1 | 25676 | -0.7775865 | 0.7775865 |
| RASAL2 | 304893 | -1.3808692 | 1.3808692 |
| RASGEF1C | 360519 | 1.6070697 | -1.6070697 |
| RASGRF1 | 192213 | 0.5586639 | -0.5586639 |
| RASGRP2 | 361714 | 1.6968195 | -1.6968195 |
| RASIP1 | 292912 | 1.7952964 | -1.7952964 |
| RASL11A | 304268 | -3.7108407 | 3.7108407 |
| RASSF8 | 312846 | -3.7016206 | 3.7016206 |
| RAVER1 | 298705 | -0.3390794 | 0.3390794 |
| RBAK | 288489 | -0.7255992 | 0.7255992 |
| RBFA | 307235 | 1.1691409 | -1.1691409 |
| RBM11 | 288321 | 2.2333245 | -2.2333245 |
| RBM12 | 652928 | -0.8207188 | 0.8207188 |
| RBM24 | 690139 | 1.451968 | -1.451968 |
| RBM46 | 310548 | 1.0298349 | -1.0298349 |
| RBMS1 | 362138 | -1.9032842 | 1.9032842 |
| RBPMS | 498642 | -0.87195694 | 0.87195694 |
| RCAN2 | 140666 | -0.7658639 | 0.7658639 |
| RDH10 | 353252 | -1.5777113 | 1.5777113 |
| REEP1 | 362384 | 0.88295263 | -0.88295263 |
| REEP2 | 682105 | 0.91989726 | -0.91989726 |
| REM2 | 64626 | -0.6479169 | 0.6479169 |

| | | | |
|---|---|---|---|
| RET | 24716 | -1.3632734 | 1.3632734 |
| RFXAP | 499617 | -0.5034862 | 0.5034862 |
| RGD1304810 | 306504 | -1.4057148 | 1.4057148 |
| RGD1305014 | 309029 | -0.93383926 | 0.93383926 |
| RGD1305587 | 294499 | 2.1736956 | -2.1736956 |
| RGD1306227 | 310377 | 0.99300516 | -0.99300516 |
| RGD1307443 | 361244 | 0.8686419 | -0.8686419 |
| RGD1307461 | 300990 | 1.7910181 | -1.7910181 |
| RGD1308147 | 307008 | -0.3366559 | 0.3366559 |
| RGD1309028 | 299265 | 1.2496943 | -1.2496943 |
| RGD1309079 | 315891 | -0.5708009 | 0.5708009 |
| RGD1310110 | 361032 | 1.0895427 | -1.0895427 |
| RGD1310769 | 299207 | 1.0653373 | -1.0653373 |
| RGD1312005 | 291580 | 1.0583966 | -1.0583966 |
| RGD1560108 | 499309 | 0.961564 | -0.961564 |
| RGD1561849 | 500393 | -1.1715535 | 1.1715535 |
| RGD1561931 | 302396 | 0.8947085 | -0.8947085 |
| RGD1563072 | 313595 | 1.8446188 | -1.8446188 |
| RGD1564036 | 497895 | 1.2878602 | -1.2878602 |
| RGD1564804 | 313551 | 1.4052379 | -1.4052379 |
| RGS12 | 54292 | 1.0016395 | -1.0016395 |
| RGS14 | 114705 | 1.3399516 | -1.3399516 |
| RGS16 | 360857 | -2.010148 | 2.010148 |
| RGS17 | 308118 | -0.698156 | 0.698156 |
| RGS2 | 84583 | -0.40462452 | 0.40462452 |
| RGS20 | 362477 | -1.5197148 | 1.5197148 |
| RGS4 | 29480 | -1.4679604 | 1.4679604 |
| RGS5 | 54294 | -2.4916008 | 2.4916008 |
| RHBDF2 | 303690 | 2.3448224 | -2.3448224 |
| RHBDL1 | 117025 | 1.6193181 | -1.6193181 |
| RHOQ | 85428 | -1.2808673 | 1.2808673 |
| RHOV | 171581 | 2.48307 | -2.48307 |
| RHPN2 | 308516 | 0.9445657 | -0.9445657 |
| RIN1 | 207119 | -2.4847422 | 2.4847422 |
| RIT1 | 499652 | 0.43847728 | -0.43847728 |
| RIT2 | 291713 | 1.3842981 | -1.3842981 |
| RNASEH2A | 364974 | 1.6685215 | -1.6685215 |
| RND1 | 362993 | 1.7105467 | -1.7105467 |
| RNF111 | 300813 | -0.30827323 | 0.30827323 |
| RNF145 | 287212 | 0.68509424 | -0.68509424 |
| RNF150 | 364983 | -0.74297404 | 0.74297404 |
| RNF152 | 293561 | 1.9067645 | -1.9067645 |
| RNF167 | 360554 | 0.67522365 | -0.67522365 |

| | | | |
|---|---|---|---|
| RNF180 | 685384 | 0.40405306 | -0.40405306 |
| RNF182 | 498726 | 1.5751994 | -1.5751994 |
| RNF19B | 313806 | 0.6673586 | -0.6673586 |
| RNF217 | 292188 | -0.64998645 | 0.64998645 |
| RNF34 | 282845 | 0.41530994 | -0.41530994 |
| RNF39 | 171387 | -0.73396945 | 0.73396945 |
| RNF5 | 407784 | 1.0183594 | -1.0183594 |
| RNFT2 | 304521 | 0.75005347 | -0.75005347 |
| RNGTT | 313131 | 0.5777789 | -0.5777789 |
| ROCK2 | 25537 | -0.97574705 | 0.97574705 |
| RPH3A | 171039 | 0.8633184 | -0.8633184 |
| RPS6KA2 | 117269 | 1.3743302 | -1.3743302 |
| RPS6KA3 | 501560 | -0.86574256 | 0.86574256 |
| RPS6KA5 | 314384 | 0.6330049 | -0.6330049 |
| RPS6KA6 | 317203 | -0.65257776 | 0.65257776 |
| RPS6KB1 | 83840 | -0.6176701 | 0.6176701 |
| RRAD | 83521 | -3.126623 | 3.126623 |
| RRAGD | 297960 | 0.9920717 | -0.9920717 |
| RRP8 | 308911 | -0.91041267 | 0.91041267 |
| RSPO3 | 498997 | 1.8153454 | -1.8153454 |
| RSRP1 | 362626 | -0.6119179 | 0.6119179 |
| RT1-S3 | 294228 | -2.6764774 | 2.6764774 |
| RTF1 | 366169 | 0.8068774 | -0.8068774 |
| RTFDC1 | 296410 | 0.5922399 | -0.5922399 |
| RTN4RL1 | 303311 | -1.1240036 | 1.1240036 |
| RTN4RL2 | 311169 | -1.19438 | 1.19438 |
| RUNX1 | 50662 | -2.4223049 | 2.4223049 |
| RUNX1T1 | 362489 | 0.62385863 | -0.62385863 |
| RUNX2 | 367218 | -1.8867472 | 1.8867472 |
| RXFP3 | 294807 | -2.7440076 | 2.7440076 |
| RXRB | 361801 | 1.4743892 | -1.4743892 |
| RXRG | 83574 | 2.587212 | -2.587212 |
| RYBP | 312603 | -0.2905089 | 0.2905089 |
| RYK | 140585 | -0.49521574 | 0.49521574 |
| SAMD4A | 305826 | -2.2564552 | 2.2564552 |
| SAMD5 | 365038 | -1.5033222 | 1.5033222 |
| SAMHD1 | 311580 | -0.70444477 | 0.70444477 |
| SAP30BP | 360662 | -0.2611519 | 0.2611519 |
| SATB1 | 316164 | 0.87971145 | -0.87971145 |
| SC5D | 114100 | -0.57787454 | 0.57787454 |
| SCG2 | 24765 | -2.7256134 | 2.7256134 |
| SCLY | 363285 | 0.79577345 | -0.79577345 |
| SCN1A | 81574 | 1.082227 | -1.082227 |

| | | | |
|---|---|---|---|
| SCN1B | 29686 | -2.0425715 | 2.0425715 |
| SCN2A | 24766 | 1.2908763 | -1.2908763 |
| SCN3B | 245956 | 0.8881229 | -0.8881229 |
| SCN4A | 25722 | 2.258376 | -2.258376 |
| SCNN1G | 24768 | -1.5599666 | 1.5599666 |
| SCRT2 | 366229 | -2.0606408 | 2.0606408 |
| SDC1 | 25216 | -1.3037148 | 1.3037148 |
| SDCBP | 83841 | -0.3909704 | 0.3909704 |
| SDCCAG8 | 305002 | 1.5519366 | -1.5519366 |
| SDHC | 289217 | 0.9934685 | -0.9934685 |
| SDK2 | 360652 | -1.9475592 | 1.9475592 |
| SDR39U1 | 361044 | 1.1415017 | -1.1415017 |
| SEC13 | 297522 | 0.3394574 | -0.3394574 |
| SEC16B | 89868 | -2.1565294 | 2.1565294 |
| SEC24B | 295461 | -0.5457721 | 0.5457721 |
| SEC24D | 310843 | -1.615148 | 1.615148 |
| SELE | 25544 | -1.9188021 | 1.9188021 |
| SEMA3A | 29751 | -2.3533933 | 2.3533933 |
| SEMA3D | 246262 | -1.2178336 | 1.2178336 |
| SEMA3E | 296789 | -0.8667002 | 0.8667002 |
| SEMA4F | 29745 | 1.4172438 | -1.4172438 |
| SEMA4G | 361764 | 1.1764392 | -1.1764392 |
| SEMA6B | 84609 | 1.1183448 | -1.1183448 |
| SEMA6C | 29744 | 2.2092333 | -2.2092333 |
| SEPHS1 | 291314 | -0.42142496 | 0.42142496 |
| SERPINB2 | 60325 | -4.882518 | 4.882518 |
| SERPINB8 | 288937 | -2.8327084 | 2.8327084 |
| SERTAD1 | 361526 | -1.5891567 | 1.5891567 |
| SERTM1 | 690333 | -0.82669574 | 0.82669574 |
| SESN1 | 294518 | 1.064322 | -1.064322 |
| SFI1 | 305467 | 0.84867126 | -0.84867126 |
| SFMBT1 | 58967 | -0.62853 | 0.62853 |
| SFMBT2 | 307106 | -1.643639 | 1.643639 |
| SFRP4 | 89803 | -1.0778743 | 1.0778743 |
| SGCA | 303468 | -1.9697973 | 1.9697973 |
| SGMS1 | 353229 | -0.8720232 | 0.8720232 |
| SH2B2 | 114203 | 1.1817917 | -1.1817917 |
| SH2D3C | 362111 | 3.8986006 | -3.8986006 |
| SH3BP1 | 300067 | -0.7575102 | 0.7575102 |
| SH3BP5 | 117186 | 1.111334 | -1.111334 |
| SH3GLB1 | 292156 | -0.39560103 | 0.39560103 |
| SHC4 | 679845 | -2.4207256 | 2.4207256 |
| SHISA5 | 301013 | 0.8572069 | -0.8572069 |

| | | | |
|---|---|---|---|
| SHQ1 | 297483.chr4.1 | -0.90175235 | 0.90175235 |
| SHROOM2 | 317435 | 0.94060034 | -0.94060034 |
| SHROOM4 | 317391 | -3.285885 | 3.285885 |
| SHTN1 | 292139 | 0.7538315 | -0.7538315 |
| SIAH3 | 692004 | -2.7351675 | 2.7351675 |
| SIDT1 | 288109 | 0.9055039 | -0.9055039 |
| SIK1 | 59329 | -1.636797 | 1.636797 |
| SIK3 | 684112 | -1.1976771 | 1.1976771 |
| SIL1 | 291673 | 1.5942905 | -1.5942905 |
| SIM1 | 309888 | -2.920881 | 2.920881 |
| SIN3A | 363067 | 0.41163677 | -0.41163677 |
| SIPA1L2 | 361442 | -0.8146569 | 0.8146569 |
| SIRT3 | 293615 | 1.1229125 | -1.1229125 |
| SIRT5 | 306840 | 1.8049828 | -1.8049828 |
| SKP2 | 294790 | -1.1556857 | 1.1556857 |
| SLAIN2 | 305310 | -0.46348232 | 0.46348232 |
| SLC16A14 | 316578 | -2.186105 | 2.186105 |
| SLC16A8 | 65200 | -1.2989899 | 1.2989899 |
| SLC17A5 | 363103 | 0.6097403 | -0.6097403 |
| SLC17A6 | 84487 | -0.9674586 | 0.9674586 |
| SLC18A2 | 25549 | -0.6991742 | 0.6991742 |
| SLC1A6 | 84012 | 1.3026911 | -1.3026911 |
| SLC22A1 | 24904 | 4.297857 | -4.297857 |
| SLC22A3 | 29504 | 2.5468235 | -2.5468235 |
| SLC24A4 | 314396 | -0.37487805 | 0.37487805 |
| SLC25A14 | 85263 | 0.5005022 | -0.5005022 |
| SLC25A20 | 117035 | 0.8498117 | -0.8498117 |
| SLC25A37 | 306000 | -1.8937051 | 1.8937051 |
| SLC25A39 | 360636 | 1.3247854 | -1.3247854 |
| SLC27A2 | 65192 | 1.1857318 | -1.1857318 |
| SLC28A3 | 140944 | 1.9427401 | -1.9427401 |
| SLC2A3 | 25551 | -1.5491256 | 1.5491256 |
| SLC30A10 | 289353 | -1.2617816 | 1.2617816 |
| SLC30A3 | 366568 | -0.72683805 | 0.72683805 |
| SLC30A4 | 64469 | -0.8386116 | 0.8386116 |
| SLC35B4 | 296969 | 0.3722925 | -0.3722925 |
| SLC35D1 | 298280 | -1.6176689 | 1.6176689 |
| SLC35D3 | 308717 | -1.6887423 | 1.6887423 |
| SLC35E3 | 362883 | -0.41176802 | 0.41176802 |
| SLC35F5 | 288993 | 0.37466466 | -0.37466466 |
| SLC37A4 | 29573 | 1.1339908 | -1.1339908 |
| SLC40A1 | 170840 | -2.2319286 | 2.2319286 |
| SLC41A2 | 362861 | -1.2773671 | 1.2773671 |

| | | | |
|---|---|---|---|
| SLC4A11 | 311423 | -1.7083405 | 1.7083405 |
| SLC4A7 | 117955 | -2.0753016 | 2.0753016 |
| SLC5A10 | 303205 | -1.8730152 | 1.8730152 |
| SLC5A3 | 114507 | -1.519374 | 1.519374 |
| SLC5A8 | 500820 | -1.7296529 | 1.7296529 |
| SLC6A15 | 282712 | 0.711413 | -0.711413 |
| SLC6A17 | 613226 | -1.3041549 | 1.3041549 |
| SLC6A7 | 117100 | 1.1860822 | -1.1860822 |
| SLC6A8 | 50690 | -0.67422664 | 0.67422664 |
| SLC9A2 | 24783 | -1.3528452 | 1.3528452 |
| SLC9B2 | 679958 | 1.9674516 | -1.9674516 |
| SLCO2A1 | 24546 | -4.5949497 | 4.5949497 |
| SLCO3A1 | 140915 | -1.3849361 | 1.3849361 |
| SLCO4A1 | 171144 | 1.2394837 | -1.2394837 |
| SLITRK4 | 302473 | 0.9764185 | -0.9764185 |
| SMAD3 | 25631 | -2.0338812 | 2.0338812 |
| SMAGP | 300236 | -2.016632 | 2.016632 |
| SMARCA2 | 361745 | 0.8919621 | -0.8919621 |
| SMARCD3 | 296732 | 1.4601693 | -1.4601693 |
| SMIM17 | 499067 | 1.050431 | -1.050431 |
| SMIM20 | 501923 | 1.1410125 | -1.1410125 |
| SMIM3 | 286910 | 0.81856567 | -0.81856567 |
| SMOC2 | 292401 | 2.2719147 | -2.2719147 |
| SMPD3 | 94338 | 1.0461674 | -1.0461674 |
| SMPDL3B | 362619 | -2.7852929 | 2.7852929 |
| SMR3B | 24867 | -1.3570703 | 1.3570703 |
| SMTN | 289734 | -1.9741095 | 1.9741095 |
| SMYD5 | 312503 | 0.71387887 | -0.71387887 |
| SNCA | 29219 | 1.2924238 | -1.2924238 |
| SNN | 29140 | 0.44895124 | -0.44895124 |
| SNRNP40 | 313056 | 0.3734343 | -0.3734343 |
| SNX9 | 683687 | -1.3578744 | 1.3578744 |
| SOAT1 | 81782 | -1.5339559 | 1.5339559 |
| SOCS3 | 89829 | -1.716651 | 1.716651 |
| SOGA3 | 292199 | 0.7784561 | -0.7784561 |
| SORBS2 | 114901 | -1.709716 | 1.709716 |
| SOX4 | 364712 | 1.2455767 | -1.2455767 |
| SP7 | 300260 | -4.5945225 | 4.5945225 |
| SPAST | 362700 | 0.5576073 | -0.5576073 |
| SPATA2L | 498963 | -0.4327329 | 0.4327329 |
| SPCS3 | 680782 | -0.78490406 | 0.78490406 |
| SPHK1 | 170897 | -3.063561 | 3.063561 |
| SPHKAP | 316561 | 1.4794879 | -1.4794879 |

| | | | |
|---|---|---|---|
| SPIRE1 | 307348 | -0.45116532 | 0.45116532 |
| SPIRE2 | 307925 | 1.3488461 | -1.3488461 |
| SPNS2 | 100270678 | -0.66034466 | 0.66034466 |
| SPOP | 287643 | 1.0444767 | -1.0444767 |
| SPRED3 | 308478 | -1.8490422 | 1.8490422 |
| SPRR1A | 499660 | -5.059616 | 5.059616 |
| SPRTN | 292101 | -1.1680199 | 1.1680199 |
| SPRY2 | 306141 | -1.6108091 | 1.6108091 |
| SPRY4 | 291610 | -2.687808 | 2.687808 |
| SPSB1 | 313722 | -2.0201378 | 2.0201378 |
| SQLE | 29230 | 0.30901468 | -0.30901468 |
| SRD5A1 | 24950 | 0.84864455 | -0.84864455 |
| SRPK1 | 361811 | -0.41135523 | 0.41135523 |
| SRPRA | 315548 | -0.29907146 | 0.29907146 |
| SRRM3 | 685890 | 0.9302858 | -0.9302858 |
| SRSF9 | 288701 | 0.4320581 | -0.4320581 |
| SRXN1 | 296271 | -4.2581544 | 4.2581544 |
| SS18 | 361295 | -0.85127825 | 0.85127825 |
| SSR1 | 361233 | -0.19632502 | 0.19632502 |
| SSTR2 | 54305 | -0.9175802 | 0.9175802 |
| SSTR3 | 171044 | -3.1048226 | 3.1048226 |
| SSTR4 | 25555 | -1.9505814 | 1.9505814 |
| ST18 | 266680 | -1.01843 | 1.01843 |
| ST3GAL2 | 64442 | 0.8283027 | -0.8283027 |
| ST3GAL3 | 64445 | 0.82038593 | -0.82038593 |
| ST6GAL1 | 25197 | -0.36678654 | 0.36678654 |
| ST6GALNAC6 | 407765 | 0.9609258 | -0.9609258 |
| ST8SIA5 | 364901 | -2.471564 | 2.471564 |
| STAC2 | 363674 | -0.914047 | 0.914047 |
| STAMBP | 171565 | 0.9105669 | -0.9105669 |
| STC2 | 63878 | -1.6325964 | 1.6325964 |
| STEAP1 | 297738 | -1.2303724 | 1.2303724 |
| STEAP2 | 312052 | -0.30343154 | 0.30343154 |
| STIM1 | 361618 | 0.4927793 | -0.4927793 |
| STK26 | 317589 | -3.2381546 | 3.2381546 |
| STOML1 | 300748 | 1.3919854 | -1.3919854 |
| STRADA | 303605 | 0.83367527 | -0.83367527 |
| STT3B | 363160 | -0.8150197 | 0.8150197 |
| STX11 | 292483 | -3.5040588 | 3.5040588 |
| SULT2B1 | 292915 | 1.1133765 | -1.1133765 |
| SUN1 | 360773 | -0.8405576 | 0.8405576 |
| SUOX | 81805 | 0.60807633 | -0.60807633 |
| SV2A | 117559 | 1.0175225 | -1.0175225 |

| | | | |
|---|---|---|---|
| SV2B | 117556 | 1.2700803 | -1.2700803 |
| SV2C | 29643 | -3.5015628 | 3.5015628 |
| SVBP | 362578 | 1.7390351 | -1.7390351 |
| SVOP | 171442 | 1.1907222 | -1.1907222 |
| SYK | 25155 | -0.5262172 | 0.5262172 |
| SYNJ2 | 84018 | -1.7209162 | 1.7209162 |
| SYNPO | 60324 | -1.3905915 | 1.3905915 |
| SYPL2 | 362018 | -2.3790407 | 2.3790407 |
| SYT10 | 60567 | -3.5787656 | 3.5787656 |
| SYT12 | 191595 | -0.9574822 | 0.9574822 |
| SYT13 | 80977 | 0.60884726 | -0.60884726 |
| SYT3 | 25731 | 1.6051745 | -1.6051745 |
| SYT6 | 60565 | -0.7595564 | 0.7595564 |
| SYT9 | 60564 | 0.75644463 | -0.75644463 |
| SYTL5 | 302538 | -1.7463497 | 1.7463497 |
| TAC1 | 24806 | -1.1776572 | 1.1776572 |
| TACC1 | 306562 | -1.1586126 | 1.1586126 |
| TACR2 | 25007 | -2.0917666 | 2.0917666 |
| TACR3 | 24808 | -2.047815 | 2.047815 |
| TADA2B | 289717 | -0.44116637 | 0.44116637 |
| TAF12 | 682902 | -0.5072301 | 0.5072301 |
| TAGAP | 308097 | -1.8094736 | 1.8094736 |
| TARS | 294810 | -0.57766813 | 0.57766813 |
| TAS1R3 | 170634 | -1.8621123 | 1.8621123 |
| TBC1D14 | 360956 | 0.71468055 | -0.71468055 |
| TBC1D17 | 292886 | 1.2522173 | -1.2522173 |
| TBC1D25 | 302552 | 0.55837303 | -0.55837303 |
| TBC1D9 | 304645 | 0.7860574 | -0.7860574 |
| TBCK | 295446 | 0.8337724 | -0.8337724 |
| TBX21 | 303496 | -1.7847887 | 1.7847887 |
| TCEAL1 | 302593 | 0.9830134 | -0.9830134 |
| TCP11L2 | 314683 | 0.6612161 | -0.6612161 |
| TCTN1 | 304486 | 0.77660185 | -0.77660185 |
| TDRD6 | 316254 | 2.2980094 | -2.2980094 |
| TEAD1 | 361630 | -2.510715 | 2.510715 |
| TEF | 29362 | 0.608276 | -0.608276 |
| TEKT2 | 298532 | 1.7960329 | -1.7960329 |
| TENM3 | 306451 | 0.9096696 | -0.9096696 |
| TEX264 | 300988 | 1.285548 | -1.285548 |
| TFAP4 | 360482 | 0.829667 | -0.829667 |
| TGFB1 | 59086 | -0.9455964 | 0.9455964 |
| TGFBR2 | 81810 | -2.4523053 | 2.4523053 |
| TGIF1 | 316742 | -1.1005571 | 1.1005571 |

| | | | |
|---|---|---|---|
| THBD | 83580 | -1.7839245 | 1.7839245 |
| THNSL1 | 498805 | 0.4096687 | -0.4096687 |
| THRB | 24831 | -1.8771453 | 1.8771453 |
| THSD7A | 500032 | 0.6431325 | -0.6431325 |
| TIGAR | 502894 | -0.43789047 | 0.43789047 |
| TIMP1 | 116510 | -1.594023 | 1.594023 |
| TLK2 | 303592 | -0.35235876 | 0.35235876 |
| TM4SF1 | 295061 | -1.8866996 | 1.8866996 |
| TMCC2 | 305095 | 1.1049625 | -1.1049625 |
| TMCC3 | 314751 | -0.9580081 | 0.9580081 |
| TMEM106B | 312132 | 0.31767172 | -0.31767172 |
| TMEM106C | 315286 | 1.0677006 | -1.0677006 |
| TMEM107 | 691750 | 1.5588582 | -1.5588582 |
| TMEM141 | 499755 | 1.8384227 | -1.8384227 |
| TMEM150C | 360916 | 1.5865021 | -1.5865021 |
| TMEM163 | 360839 | -0.7065779 | 0.7065779 |
| TMEM179 | 314472 | 1.2138616 | -1.2138616 |
| TMEM182 | 501129 | -2.9876928 | 2.9876928 |
| TMEM184B | 362959 | 0.3989972 | -0.3989972 |
| TMEM19 | 299800 | 0.858972 | -0.858972 |
| TMEM200C | 501201 | 1.0343357 | -1.0343357 |
| TMEM202 | 691306 | -1.2168857 | 1.2168857 |
| TMEM218 | 300516 | 1.4755671 | -1.4755671 |
| TMEM229B | 503035 | 0.8901293 | -0.8901293 |
| TMEM246 | 362518 | 1.4409344 | -1.4409344 |
| TMEM25 | 689172 | 1.2348101 | -1.2348101 |
| TMEM39A | 288092 | 0.632154 | -0.632154 |
| TMEM41A | 681708 | -0.71854585 | 0.71854585 |
| TMEM42 | 363171 | 1.2895105 | -1.2895105 |
| TMEM45A | 680866 | -2.076463 | 2.076463 |
| TMEM55B | 364298 | 0.9689839 | -0.9689839 |
| TMEM65 | 500874 | -0.7801371 | 0.7801371 |
| TMEM68 | 312946 | 0.97808397 | -0.97808397 |
| TMEM8A | 303004 | 0.98735017 | -0.98735017 |
| TMIE | 501061 | 2.1490898 | -2.1490898 |
| TMOD1 | 25566 | -0.79898345 | 0.79898345 |
| TMPRSS6 | 315388 | -2.1276395 | 2.1276395 |
| TMTC4 | 290501 | 1.451589 | -1.451589 |
| TNFAIP1 | 287543 | 0.38224366 | -0.38224366 |
| TNFRSF11A | 498206 | 0.8378635 | -0.8378635 |
| TNFRSF12A | 302965 | -1.5811874 | 1.5811874 |
| TNFRSF25 | 500592 | -2.0133827 | 2.0133827 |
| TNIP1 | 363599 | 0.5466149 | -0.5466149 |

| | | | |
|---|---|---|---|
| TNIP2 | 305451 | -0.9859405 | 0.9859405 |
| TNRC18 | 304302 | 0.62138903 | -0.62138903 |
| TOR1AIP1 | 246314 | -0.38566485 | 0.38566485 |
| TOX3 | 291908 | 1.0046704 | -1.0046704 |
| TP73 | 362675 | -1.8165276 | 1.8165276 |
| TPBG | 83684 | -1.2230638 | 1.2230638 |
| TPD52L3 | 293894 | 2.1558177 | -2.1558177 |
| TPH1 | 24848 | -1.759721 | 1.759721 |
| TPK1 | 680668 | 1.1043934 | -1.1043934 |
| TPM4 | 24852 | -0.78833544 | 0.78833544 |
| TPPP3 | 291966 | 2.493314 | -2.493314 |
| TRAPPC2B | 100910318 | 1.4842247 | -1.4842247 |
| TRAPPC3 | 362599 | 1.1807364 | -1.1807364 |
| TRERF1 | 316219 | 0.6702241 | -0.6702241 |
| TRIB1 | 78969 | -2.3380017 | 2.3380017 |
| TRIB2 | 313974 | -0.85800225 | 0.85800225 |
| TRIM23 | 81002 | 0.6327717 | -0.6327717 |
| TRIM45 | 295323 | 1.0136446 | -1.0136446 |
| TRIM9 | 155812 | -1.8680766 | 1.8680766 |
| TRIQK | 500413 | 1.4419904 | -1.4419904 |
| TRMT10C | 304012 | -0.3914864 | 0.3914864 |
| TRMT6 | 311441 | -0.88919955 | 0.88919955 |
| TRPC1 | 89821 | 0.63926923 | -0.63926923 |
| TRPC4 | 84494 | 1.104643 | -1.104643 |
| TRPC6 | 89823 | -3.409288 | 3.409288 |
| TRPC7 | 282822 | 1.1666611 | -1.1666611 |
| TRPM4 | 171143 | 1.397249 | -1.397249 |
| TRPT1 | 293704 | 1.089093 | -1.089093 |
| TRPV6 | 114246 | 1.3203087 | -1.3203087 |
| TSC22D1 | 498545 | 0.61734474 | -0.61734474 |
| TSPAN17 | 306771 | 1.5900029 | -1.5900029 |
| TSPAN2 | 64521 | 0.8337795 | -0.8337795 |
| TSPAN5 | 362048 | -0.75329566 | 0.75329566 |
| TSPYL1 | 29544 | 0.55921113 | -0.55921113 |
| TTC19 | 691506 | -0.5289188 | 0.5289188 |
| TTC9 | 500689 | -0.55293614 | 0.55293614 |
| TTLL1 | 362969 | 1.3057308 | -1.3057308 |
| TTPAL | 296349 | -1.0306127 | 1.0306127 |
| TUBA8 | 500377 | -1.2659577 | 1.2659577 |
| TUBB4B | 296554 | 0.42896006 | -0.42896006 |
| TUBG2 | 680991 | 1.3486153 | -1.3486153 |
| TUBGCP3 | 306599 | -0.40801 | 0.40801 |
| TUSC5 | 360576 | -2.4632652 | 2.4632652 |

| | | | |
|---|---|---|---|
| TWIST1 | 85489 | -0.961156 | 0.961156 |
| TXNIP | 117514 | 1.4114915 | -1.4114915 |
| TXNRD1 | 58819 | -0.74920535 | 0.74920535 |
| TYMS | 29261 | -0.8746929 | 0.8746929 |
| TYR | 308800 | 2.1056445 | -2.1056445 |
| TYRO3 | 25232 | -0.3947916 | 0.3947916 |
| UAP1 | 498272 | -0.38090712 | 0.38090712 |
| UBASH3B | 315579 | -2.500164 | 2.500164 |
| UBE2D1 | 361831 | 1.0117896 | -1.0117896 |
| UHRF1BP1L | 363009 | -0.48036596 | 0.48036596 |
| ULK2 | 303206 | 0.4279266 | -0.4279266 |
| UNC119 | 29402 | 1.4818367 | -1.4818367 |
| UNC5B | 60630 | -2.809945 | 2.809945 |
| UOX | 114768 | -2.7210045 | 2.7210045 |
| USF1 | 83586 | 0.7900194 | -0.7900194 |
| USO1 | 56042 | -0.34504065 | 0.34504065 |
| USP11 | 408217 | 0.580222 | -0.580222 |
| USP20 | 311856 | 1.0782478 | -1.0782478 |
| USP28 | 315639 | -0.79407537 | 0.79407537 |
| USP46 | 289584 | 0.6767605 | -0.6767605 |
| USP49 | 316211 | 0.6756008 | -0.6756008 |
| UST | 361450 | -1.3118315 | 1.3118315 |
| UTP15 | 310019 | -0.9198079 | 0.9198079 |
| UTP20 | 314713 | -0.92448187 | 0.92448187 |
| VAC14 | 307842 | 0.91569674 | -0.91569674 |
| VASP | 361517 | -1.2618965 | 1.2618965 |
| VCL | 305679 | -1.6796203 | 1.6796203 |
| VEGFA | 83785 | -1.0660167 | 1.0660167 |
| VGF | 29461 | -3.8885574 | 3.8885574 |
| VGLL4 | 297523 | -0.45178235 | 0.45178235 |
| VIPR1 | 24875 | -2.1832316 | 2.1832316 |
| VKORC1 | 309004 | 1.4050825 | -1.4050825 |
| VOPP1 | 362374 | 0.8389173 | -0.8389173 |
| VPS25 | 681059 | 0.9774818 | -0.9774818 |
| VPS53 | 287535 | 0.69485587 | -0.69485587 |
| VSTM2B | 361560 | -0.8955742 | 0.8955742 |
| VWA1 | 298683 | 2.2382662 | -2.2382662 |
| VWA5B2 | 303812 | 1.0999229 | -1.0999229 |
| VWC2 | 501231 | -0.7515774 | 0.7515774 |
| VWC2L | 501160 | 2.1847382 | -2.1847382 |
| WASH1 | 367328 | 1.4061364 | -1.4061364 |
| WDHD1 | 305827 | -0.54014874 | 0.54014874 |
| WDR1 | 360950 | -0.49472558 | 0.49472558 |

| | | | |
|---|---|---|---|
| WDR34 | 296618 | 1.4017438 | -1.4017438 |
| WDR43 | 362703 | -0.6477642 | 0.6477642 |
| WDR45 | 302559 | 1.1314515 | -1.1314515 |
| WDR45B | 360682 | -0.67207795 | 0.67207795 |
| WDR6 | 301007 | 0.90195644 | -0.90195644 |
| WFS1 | 83725 | -0.6478758 | 0.6478758 |
| WIPI2 | 288498 | -0.4373512 | 0.4373512 |
| WISP1 | 65154 | -2.8569026 | 2.8569026 |
| WNT6 | 316526 | -2.9822986 | 2.9822986 |
| WNT7A | 114850 | 0.54971087 | -0.54971087 |
| WSCD1 | 287466 | -0.6364453 | 0.6364453 |
| WSCD2 | 360824 | 1.4729046 | -1.4729046 |
| XKR4 | 297801 | 0.6514431 | -0.6514431 |
| XKR7 | 311549 | -1.9481895 | 1.9481895 |
| XKR8 | 313033 | -1.112087 | 1.112087 |
| XPO4 | 290280 | -0.98097914 | 0.98097914 |
| XRCC4 | 309995 | 1.1296892 | -1.1296892 |
| YPEL3 | 293491 | 1.6609019 | -1.6609019 |
| YPEL5 | 298792 | 0.9776608 | -0.9776608 |
| ZBTB16 | 353227 | 1.1991673 | -1.1991673 |
| ZBTB22 | 309630 | 0.99966913 | -0.99966913 |
| ZBTB46 | 311718 | -1.4119025 | 1.4119025 |
| ZBTB5 | 298084 | 0.5326453 | -0.5326453 |
| ZBTB9 | 294289 | -0.53306437 | 0.53306437 |
| ZC2HC1A | 310244 | 1.0619466 | -1.0619466 |
| ZC3H18 | 292067 | -0.47602633 | 0.47602633 |
| ZC3H8 | 311414 | 0.7709499 | -0.7709499 |
| ZC4H2 | 367838 | 1.3166186 | -1.3166186 |
| ZDHHC14 | 499014 | -0.5482368 | 0.5482368 |
| ZDHHC20 | 305923 | -0.734937 | 0.734937 |
| ZDHHC5 | 362156 | -0.77307737 | 0.77307737 |
| ZFAND2B | 363253 | 1.1858829 | -1.1858829 |
| ZFAND6 | 293067 | 1.0159383 | -1.0159383 |
| ZFAT | 362925 | -0.368832 | 0.368832 |
| ZFP217 | 311764 | -2.161255 | 2.161255 |
| ZFP238 | 64619 | 1.1051159 | -1.1051159 |
| ZFP358 | 360754 | 0.9876498 | -0.9876498 |
| ZFP362 | 297879 | 0.492345 | -0.492345 |
| ZFP395 | 305972 | 0.5507338 | -0.5507338 |
| ZFP414 | 299647 | 1.2685622 | -1.2685622 |
| ZFP467 | 500110 | 0.87794447 | -0.87794447 |
| ZFP513 | 313913 | 0.50401187 | -0.50401187 |
| ZFP575 | 308430 | 1.4536451 | -1.4536451 |

| | | | |
|---|---|---|---|
| ZFP672 | 303165 | -0.67905396 | 0.67905396 |
| ZFP697 | 295310 | -1.3314832 | 1.3314832 |
| ZFP709 | 266773 | -0.7732385 | 0.7732385 |
| ZFP775 | 312309 | 1.252074 | -1.252074 |
| ZFP786 | 100158223 | 0.84110814 | -0.84110814 |
| ZFP90 | 498945 | 1.1531198 | -1.1531198 |
| ZFPM1 | 691504 | -1.1701488 | 1.1701488 |
| ZFYVE1 | 299188 | 0.5481313 | -0.5481313 |
| ZFYVE19 | 499871 | 1.4545959 | -1.4545959 |
| ZFYVE9 | 313477 | -0.6973551 | 0.6973551 |
| ZGPAT | 296478 | 0.540302 | -0.540302 |
| ZMIZ1 | 361103 | -1.3804868 | 1.3804868 |
| ZMYND8 | 296374 | 0.6084991 | -0.6084991 |
| ZNRF2 | 362367 | 0.7356772 | -0.7356772 |
| ZPBP2 | 363676 | 0.8807924 | -0.8807924 |
| ZW10 | 363059 | -0.37213176 | 0.37213176 |
| ZYX | 114636 | -0.7388414 | 0.7388414 |

**Epilepsy-linked V+B v CTRL**

GENE SYMBOL
GALR1
ARC
PLAUR
EGR2
FOSB
GAL
NPTX2
LAMP5
MMP9
EGR4
PRIMA1
EGR1
GRASP
SCG2
ADRA1D
GADD45B
DUSP5
KLF10
KCNA1
SCN1B
DUSP6
HTR1B
EGR3
PTGS2
GFRA1
HTR1F
INHBA
FGF2
NGF
RAG1
SERTAD1
BAG3
RGS4
NRN1
CITED2
THRB
TRIB1
PLCL1
PDP1
FOS
ADRA1B
LAMB3
RGS16

TAC1
BDNF
BAIAP2
HOMER1
GDNF
PRICKLE1
IFRD1
CREM
ADRA1A
CDKN1A
BHLHE40
SOCS3
PNOC
HAS1
KCNV1
TRIM9
IER2
SV2C
SPRY2
HSPA1A/HSPA1B
OPRD1
SLC2A3
PLK2
NPY2R
GADD45G
SPNS2
ERRFI1
JUNB
GRIN2A
PTPRN
DNAJB5
DUSP2
POLG
ADM
FGF7
CYR61
NPY
SSTR2
CALCRL
JUN
CAMKK1
KCNB1
CH25H
HCN1
GADD45A

NR4A1
METRNL
HTR7
TYRO3
DIAPH1
SPATA5
ZFP36
RGS2
MYC
GABRQ
NAMPT
GFRA2
SH3GL1
CACNB4
ADCYAP1
LINGO1
KCNA4
STAM
PACSIN1
PLK3
MCM6
FGF22
GRIN1
NRG1
LDLR
GRIN3B
PLAT
GNA11
ADARB1
NR4A3
SIGMAR1
HES1
SLC2A1
HNRNPU
ARHGEF7
PPP1R1B
FYN
PNKP
ARHGEF9
RAPGEF3
MCL1
AARS
KCNK1
WDR26
FKBP1A

PMP22
USF1
TRIM3
TWNK
DEPDC5
SCLY
ARRDC3
ITPR1
SV2A
ABHD6
KCNJ9
SLC4A10
STRADB
FGF13
STRADA
PRRT2
BCL2L11
SLC37A4
CRYM
BAD
TEF
DEAF1
SLITRK3
NGLY1
CNR1
MBNL2
PNPO
BEND5
CXCR4
NR2F1
CNNM2
ST3GAL3
GABRD
KCNH3
BID
SCN1A
SCN3B
ANKRD6
ST8SIA4
GABRA1
AKAP5
DBP
BBC3
NTF3
KCNMB4

TRPC7
PSEN2
PCDH19
ELAVL4
SCN2A
CHRNA5
LGI1
CNTN2
SV2B
SCN4A
GABRG2
OPRK1
IGSF9
GRM1
NEUROD1
KCNK2

**Epilepsy-linked WP+B and RX2+B reversal gene list**

| Gene Syml | ID |
|---|---|
| KCNK2 | 170899 |
| SCN4A | 25722 |
| IGSF9 | 304982 |
| CYR61 | 83476 |
| PCDH19 | 317183 |
| KCNMB4 | 66016 |
| GABRD | 29689 |
| GRM1 | 24414 |
| BBC3 | 317673 |
| CRYM | 117024 |
| ELAVL4 | 432358 |
| CNTN2 | 25356 |
| NTF3 | 81737 |
| BEND5 | 362564 |
| HSPA1A/H | 294254 |
| BAD | 64639 |
| SCN2A | 24766 |
| SV2B | 117556 |
| KCNH3 | 27150 |
| JUNB | 24517 |
| CHRNA5 | 25102 |
| TRPC7 | 282822 |
| GABRG2 | 29709 |
| LGI1 | 252892 |
| SLC37A4 | 29573 |
| TWNK | 309441 |
| SCN1A | 81574 |
| NR2F1 | 81808 |
| NEUROD1 | 29458 |
| ANKRD6 | 500430 |
| FGF13 | 84488 |
| SV2A | 117559 |
| DBP | 24309 |
| PRRT2 | 361651 |
| SCN3B | 245956 |
| BID | 64625 |
| STRADA | 303605 |
| CXCR4 | 60628 |
| ST3GAL3 | 64445 |
| USF1 | 83586 |
| PNPO | 64533 |
| TEF | 29362 |
| DEPDC5 | 305464 |

| | |
|---|---|
| CNR1 | 25248 |
| ABHD6 | 305795 |
| CNNM2 | 294014 |
| TYRO3 | 25232 |
| KCNK1 | 59324 |
| RGS2 | 84583 |
| MCL1 | 60430 |
| CAMKK1 | 60341 |
| PMP22 | 24660 |
| SPNS2 | 1E+08 |
| HTR7 | 65032 |
| DIAPH1 | 307483 |
| PACSIN1 | 29704 |
| MYC | 24577 |
| DNAJB5 | 313811 |
| ADM | 25026 |
| PLK2 | 83722 |
| PTPRN | 116660 |
| IFRD1 | 29596 |
| KCNB1 | 25736 |
| GABRQ | 65187 |
| SSTR2 | 54305 |
| HCN1 | 84390 |
| POLG | 85472 |
| NAMPT | 297508 |
| ERRFI1 | 313729 |
| METRNL | 316842 |
| KCNV1 | 60326 |
| KCNA4 | 25469 |
| TAC1 | 24806 |
| CH25H | 309527 |
| CALCRL | 25029 |
| CACNB4 | 58942 |
| EGR2 | 114090 |
| NPY2R | 66024 |
| CREM | 25620 |
| RGS4 | 29480 |
| GADD45B | 299626 |
| PRICKLE1 | 315259 |
| BDNF | 24225 |
| SLC2A3 | 25551 |
| SERTAD1 | 361526 |
| CITED2 | 114490 |
| SPRY2 | 306141 |
| LAMB3 | 305078 |

| | |
|---|---|
| BAIAP2 | 117542 |
| DUSP6 | 116663 |
| SOCS3 | 89829 |
| ADRA1B | 24173 |
| HOMER1 | 29546 |
| NGF | 310738 |
| TRIM9 | 155812 |
| GDNF | 25453 |
| THRB | 24831 |
| PDP1 | 54705 |
| RGS16 | 360857 |
| SCN1B | 29686 |
| PLCL1 | 84587 |
| BHLHE40 | 79431 |
| NRN1 | 83834 |
| PTGS2 | 29527 |
| TRIB1 | 78969 |
| INHBA | 29200 |
| DUSP5 | 171109 |
| KLF10 | 81813 |
| ADRA1D | 29413 |
| GFRA1 | 25454 |
| EGR3 | 25148 |
| SCG2 | 24765 |
| HAS1 | 282821 |
| LAMP5 | 362220 |
| BAG3 | 293524 |
| GRASP | 192254 |
| KCNA1 | 24520 |
| GAL | 29141 |
| HTR1F | 60448 |
| HTR1B | 25075 |
| GRIN2A | 24409 |
| SV2C | 29643 |
| NPTX2 | 288475 |
| ADRA1A | 29412 |
| ARC | 54323 |
| FGF2 | 54250 |
| PLAUR | 50692 |
| GALR1 | 50577 |

**Top WP+B and RX2+B in network by category**

| Synaptic Plasticity Group | Entrez ID |
|---|---|
| ADCY8 | 29241 |
| ADORA1 | 29290 |
| ADORA2A | 25369 |
| ARC | 54323 |
| BAIAP2 | 117542 |
| BDNF | 24225 |
| CACNB4 | 58942 |
| CDK5 | 140908 |
| CNR1 | 25248 |
| CNTN2 | 25356 |
| CREM | 25620 |
| DRD5 | 25195 |
| EFNA4 | 310643 |
| EFNB3 | 360546 |
| EGR2 | 114090 |
| EGR3 | 25148 |
| ERCC1 | 292673 |
| GRIN2A | 24409 |
| GRIP1 | 84016 |
| GRM1 | 24414 |
| HOMER1 | 29546 |
| HRH1 | 24448 |
| IGF1 | 24482 |
| INHBA | 29200 |
| JPH3 | 307916 |
| JPH4 | 445271 |
| JUNB | 24517 |
| KCNH3 | 27150 |
| KLF9 | 117560 |
| KLF10 | 81813 |
| KRAS | 24525 |
| LZTS1 | 266711 |
| NGF | 310738 |
| NPTX2 | 288475 |
| NTF3 | 81737 |
| NTRK3 | 29613 |
| PCDH8 | 64865 |
| PLK2 | 83722 |
| PPP1CA | 24668 |
| PRKAR2A | 29699 |

| | |
|---|---|
| RAB3A | 25531 |
| RASGRF1 | 192213 |
| RGS2 | 84583 |
| SNCA | 29219 |
| SYNPO | 60324 |
| TIMP1 | 116510 |
| VGF | 29461 |

| **Receptors and ion channels group** | **Entrez ID** |
|---|---|
| ADGRB1 | 362931 |
| ADGRB3 | 301309 |
| ADGRE5 | 361383 |
| ADORA1 | 29290 |
| ADORA2A | 25369 |
| ADRA1A | 29412 |
| ADRA1B | 24173 |
| ADRA1D | 29413 |
| AGTR1 | 24180 |
| ANO1 | 309135 |
| CACNB3 | 25297 |
| CALCRL | 25029 |
| CCKAR | 24889 |
| CCKBR | 25706 |
| CHRNA5 | 25102 |
| CNR1 | 25248 |
| DRD5 | 25195 |
| EDNRA | 24326 |
| F2R | 25439 |
| FFAR4 | 294075 |
| FZD1 | 58868 |
| FZD4 | 64558 |
| GABARAPL1 | 689161 |
| GABRD | 29689 |
| GABRG2 | 29709 |
| GABRQ | 65187 |
| GALR1 | 50577 |
| GCGR | 24953 |
| GPR3 | 266769 |
| GPR4 | 308408 |
| GPR12 | 80840 |
| GPR26 | 192153 |
| GPR37 | 117549 |
| GPR45 | 301372 |

| | |
|---|---|
| GPR50 | 117097 |
| GPR61 | 310780 |
| GPR63 | 297952 |
| GPR68 | 314386 |
| GPR83 | 140595 |
| GPR85 | 64020 |
| GPR88 | 64443 |
| GPR101 | 317608 |
| GPR139 | 293545 |
| GPR156 | 260430 |
| GPR158 | 291352 |
| GPRC5A | 312790 |
| GPRC5C | 287805 |
| GRIN2A | 24409 |
| GRIP1 | 84016 |
| GRM1 | 24414 |
| GRM6 | 24419 |
| HCN1 | 84390 |
| HCN3 | 114245 |
| HRH1 | 24448 |
| HRH3 | 85268 |
| HTR7 | 65032 |
| HTR1B | 25075 |
| HTR1F | 60448 |
| KCNA1 | 24520 |
| KCNA4 | 25469 |
| KCNB1 | 25736 |
| KCNC4 | 684516 |
| KCND2 | 65180 |
| KCNE2 | 171138 |
| KCNF1 | 298908 |
| KCNH3 | 27150 |
| KCNH4 | 114032 |
| KCNJ4 | 116649 |
| KCNJ5 | 29713 |
| KCNJ12 | 117052 |
| KCNK1 | 59324 |
| KCNK2 | 170899 |
| KCNK3 | 29553 |
| KCNMB2 | 294961 |
| KCNMB4 | 66016 |
| KCNN1 | 54261 |
| KCNN3 | 54263 |

| | |
|---|---|
| KCNV1 | 60326 |
| KCTD6 | 305792 |
| KCTD15 | 499129 |
| LPAR1 | 116744 |
| MAS1 | 25153 |
| MC5R | 25726 |
| MCHR1 | 83567 |
| NMBR | 25264 |
| NPY1R | 29358 |
| NPY2R | 66024 |
| OPRL1 | 29256 |
| OXTR | 25342 |
| P2RY1 | 25265 |
| P2RY14 | 171108 |
| PTAFR | 58949 |
| PTGER4 | 84023 |
| RXFP3 | 294807 |
| SCN1A | 81574 |
| SCN1B | 29686 |
| SCN2A | 24766 |
| SCN3B | 245956 |
| SCN4A | 25722 |
| SCNN1G | 24768 |
| SSTR2 | 54305 |
| SSTR3 | 171044 |
| SSTR4 | 25555 |
| TACR2 | 25007 |
| TACR3 | 24808 |
| TAS1R3 | 170634 |
| VIPR1 | 24875 |

| **Proliferation Group** | **Entrez ID** |
|---|---|
| AATK | 690853 |
| ADCY7 | 84420 |
| ADCY8 | 29241 |
| ADORA2A | 25369 |
| AKT2 | 25233 |
| ANKRD6 | 500430 |
| APBA2 | 83610 |
| APBB1 | 29722 |
| ARHGAP17 | 63994 |
| ARHGEF25 | 314904 |
| BAIAP2 | 117542 |

| | |
|---|---|
| BDNF | 24225 |
| CABLES1 | 307585 |
| CACNB3 | 25297 |
| CACNB4 | 58942 |
| CAMKK1 | 60341 |
| CAT | 24248 |
| CAV1 | 25404 |
| CCND1 | 58919 |
| CDH7 | 29162 |
| CDH15 | 361432 |
| CDK5 | 140908 |
| CFLAR | 117279 |
| CLASP2 | 114514 |
| CNP | 25275 |
| CNR1 | 25248 |
| CNTN2 | 25356 |
| COL4A1 | 290905 |
| COL5A1 | 85490 |
| CSPG5 | 50568 |
| CX3CL1 | 89808 |
| CXCR4 | 60628 |
| DISC1 | 307940 |
| DLC1 | 58834 |
| DOK4 | 361364 |
| DOK5 | 502694 |
| DPYSL2 | 25416 |
| EEA1 | 314764 |
| EFNB3 | 360546 |
| EGR3 | 25148 |
| EIF5A | 287444 |
| ELAVL4 | 432358 |
| EML1 | 362783 |
| ESR1 | 24890 |
| ETV4 | 360635 |
| ETV5 | 303828 |
| EXOC5 | 60627 |
| EZR | 54319 |
| FEZ1 | 81730 |
| FGF2 | 54250 |
| FGF3 | 170633 |
| FGF8 | 29349 |
| FGF9 | 25444 |
| FGF13 | 84488 |

| | |
|---|---|
| FGFR1 | 79114 |
| FKBP5 | 361810 |
| FZR1 | 314642 |
| GAL | 29141 |
| GAS7 | 85246 |
| GDNF | 25453 |
| GFRA1 | 25454 |
| GPM6A | 306439 |
| GPR3 | 266769 |
| GPR12 | 80840 |
| GRASP | 192254 |
| GRK5 | 59075 |
| GRN | 29143 |
| IFNG | 25712 |
| IGF1 | 24482 |
| IGSF9 | 304982 |
| IL6R | 24499 |
| IRS2 | 29376 |
| IRX6 | 307715 |
| ITGA6 | 114517 |
| JAK1 | 84598 |
| JAK2 | 24514 |
| KCNA1 | 24520 |
| KITLG | 60427 |
| KL | 83504 |
| KRAS | 24525 |
| LGI1 | 252892 |
| LPAR1 | 116744 |
| MAFK | 246760 |
| MAML3 | 310405 |
| MAP2K3 | 303200 |
| MAP2K4 | 287398 |
| MAP2K5 | 29568 |
| MAP2K6 | 114495 |
| MAPK3 | 50689 |
| MAPK15 | 286997 |
| MAR8 | 312656 |
| MFN1 | 192647 |
| MUSK | 81725 |
| NEUROD4 | 288821 |
| NFATC1 | 100361818 |
| NGB | 85382 |
| NGF | 310738 |

| | |
|---|---|
| NOG | 25495 |
| NPTX1 | 266777 |
| NR1D1 | 252917 |
| NRF1 | 312195 |
| NRN1 | 83834 |
| NTF3 | 81737 |
| NTRK3 | 29613 |
| PAK5 | 311450 |
| PAK6 | 296078 |
| PDGFB | 24628 |
| PDP1 | 54705 |
| PIK3C3 | 65052 |
| PIK3CB | 85243 |
| PIK3CD | 366508 |
| PJA2 | 192256 |
| PLAGL1 | 25157 |
| PLCL1 | 84587 |
| PLPPR4 | 295401 |
| POLR3E | 361640 |
| POU4F2 | 171355 |
| POU4F3 | 364855 |
| PRAG1 | 306506 |
| PRKAA1 | 65248 |
| PRKAR2A | 29699 |
| PRKCQ | 85420 |
| PRKCZ | 25522 |
| PTPDC1 | 291022 |
| PXN | 360820 |
| RAB33A | 317580 |
| RALGDS | 29622 |
| RAP1B | 171337 |
| RASA1 | 25676 |
| RASGRF1 | 192213 |
| RET | 24716 |
| RHOQ | 85428 |
| RHOV | 171581 |
| RIT1 | 499652 |
| RIT2 | 291713 |
| RND1 | 362993 |
| ROCK2 | 25537 |
| RTN4RL2 | 311169 |
| RXRB | 361801 |
| RXRG | 83574 |

| | |
|---|---|
| SCN1B | 29686 |
| SDC1 | 25216 |
| SEMA3A | 29751 |
| SEMA3D | 246262 |
| SH2D3C | 362111 |
| SMARCD3 | 296732 |
| SOCS3 | 89829 |
| SPAST | 362700 |
| SPRR1A | 499660 |
| SPRY2 | 306141 |
| SS18 | 361295 |
| SSTR3 | 171044 |
| SYK | 25155 |
| TGFB1 | 59086 |
| TGFBR2 | 81810 |
| TNFRSF12A | 302965 |
| TP73 | 362675 |
| Tpm4 | 24852 |
| TRPC1 | 89821 |
| TWIST1 | 85489 |
| VCL | 305679 |
| VEGFA | 83785 |
| VGF | 29461 |
| ZBTB18 | 64619 |

| **Neurogenesis group** | **Entrez ID** |
|---|---|
| AATK | 690853 |
| ABI1 | 79249 |
| ACSL4 | 113976 |
| ACVR1 | 79558 |
| ADAP1 | 171097 |
| ADCY7 | 84420 |
| ADCY8 | 29241 |
| ADGRB1 | 362931 |
| ADGRB3 | 301309 |
| ADGRE5 | 361383 |
| ADM | 25026 |
| ADORA2A | 25369 |
| AIPL1 | 59110 |
| AKT2 | 25233 |
| ANAPC2 | 296558 |
| ANKRD6 | 500430 |
| APBA2 | 83610 |

| | |
|---|---|
| APBB1 | 29722 |
| ARC | 54323 |
| AREG | 29183 |
| ARHGAP17 | 63994 |
| ARHGEF25 | 314904 |
| ARHGEF28 | 361882 |
| ARPP21 | 363153 |
| BAIAP2 | 117542 |
| BDNF | 24225 |
| BHLHE22 | 365748 |
| BHLHE23 | 499952 |
| BID | 64625 |
| BLOC1S6 | 317630 |
| BRINP3 | 286901 |
| CABLES1 | 307585 |
| CACNB3 | 25297 |
| CACNB4 | 58942 |
| CAMK1G | 171358 |
| CAMKK1 | 60341 |
| CAT | 24248 |
| CAV1 | 25404 |
| CBLN1 | 498922 |
| CBLN2 | 291388 |
| CCKAR | 24889 |
| CCND1 | 58919 |
| CDH7 | 29162 |
| CDH15 | 361432 |
| CDK5 | 140908 |
| CDK5RAP1 | 252827 |
| CDK5RAP3 | 80278 |
| CFLAR | 117279 |
| Chrm2 | 81645 |
| CLASP2 | 114514 |
| CLN8 | 306619 |
| CNP | 25275 |
| CNR1 | 25248 |
| CNTFR | 313173 |
| CNTN2 | 25356 |
| COL4A1 | 290905 |
| COL5A1 | 85490 |
| CREM | 25620 |
| CSPG5 | 50568 |
| CTSF | 361704 |

| | |
|---|---|
| CUX2 | 288665 |
| CX3CL1 | 89808 |
| CXCR4 | 60628 |
| CYP26B1 | 312495 |
| CYR61 | 83476 |
| DDIT4 | 140942 |
| DIAPH1 | 307483 |
| DISC1 | 307940 |
| DLC1 | 58834 |
| DLG1 | 25252 |
| DLK1 | 114587 |
| DOK4 | 361364 |
| DOK5 | 502694 |
| DPYSL2 | 25416 |
| EBF1 | 116543 |
| EBF3 | 361668 |
| ECEL1 | 60417 |
| EDNRA | 24326 |
| EEA1 | 314764 |
| EEF2K | 25435 |
| EFNA4 | 310643 |
| EFNB3 | 360546 |
| EGR2 | 114090 |
| EGR3 | 25148 |
| EHD1 | 293692 |
| EIF5A | 287444 |
| ELAVL4 | 432358 |
| EML1 | 362783 |
| EPHA3 | 29210 |
| EPHA4 | 316539 |
| EPHB6 | 312275 |
| ESR1 | 24890 |
| ETV4 | 360635 |
| ETV5 | 303828 |
| EXOC5 | 60627 |
| EZR | 54319 |
| F2R | 25439 |
| FAIM2 | 246274 |
| FBXO31 | 498959 |
| FEZ1 | 81730 |
| FGF2 | 54250 |
| FGF3 | 170633 |
| FGF8 | 29349 |

| | |
|---|---|
| FGF9 | 25444 |
| FGF13 | 84488 |
| FGFR1 | 79114 |
| FIG4 | 309855 |
| FKBP5 | 361810 |
| FLOT1 | 64665 |
| FLRT1 | 499308 |
| FOXP2 | 500037 |
| FST | 24373 |
| FZR1 | 314642 |
| GABRG2 | 29709 |
| GAL | 29141 |
| GAS7 | 85246 |
| GBX1 | 246149 |
| GDNF | 25453 |
| GEM | 297902 |
| GFRA1 | 25454 |
| GLDN | 315675 |
| GLI2 | 304729 |
| GPC2 | 171517 |
| GPM6A | 306439 |
| GPR3 | 266769 |
| GPR12 | 80840 |
| GRASP | 192254 |
| GRIN2A | 24409 |
| GRIP1 | 84016 |
| GRK5 | 59075 |
| GRM1 | 24414 |
| GRM6 | 24419 |
| GRN | 29143 |
| GUCY1A3 | 497757 |
| HCN1 | 84390 |
| HMG20A | 315689 |
| HOMER1 | 29546 |
| HOMER2 | 29547 |
| HSPA1A/HSPA1B | 294254 |
| HTR7 | 65032 |
| HTR1B | 25075 |
| IFNG | 25712 |
| IFT122 | 312651 |
| IGF1 | 24482 |
| IGSF9 | 304982 |
| IL1RAP | 25466 |

| | |
|---|---|
| IL6R | 24499 |
| INHBA | 29200 |
| IRS2 | 29376 |
| IRX6 | 307715 |
| ITGA6 | 114517 |
| ITM2C | 301575 |
| ITSN1 | 29491 |
| JAK1 | 84598 |
| JAK2 | 24514 |
| JUNB | 24517 |
| KCNA1 | 24520 |
| KCND2 | 65180 |
| KDM1A | 500569 |
| KDM4A | 313539 |
| KITLG | 60427 |
| KL | 83504 |
| KLF9 | 117560 |
| KRAS | 24525 |
| LCK | 313050 |
| LDB1 | 309447 |
| LEF1 | 161452 |
| LGI1 | 252892 |
| LHX2 | 296706 |
| LHX4 | 360858 |
| LIMK2 | 29524 |
| LMNA | 60374 |
| LMO4 | 362051 |
| LPAR1 | 116744 |
| LRRC4 | 641521 |
| LRRN1 | 500280 |
| LRRN3 | 81514 |
| LRRTM3 | 294380 |
| LRTM2 | 680883 |
| LZTS1 | 266711 |
| MAFK | 246760 |
| MAML3 | 310405 |
| MAN2A1 | 25478 |
| MAP2K3 | 303200 |
| MAP2K4 | 287398 |
| MAP2K5 | 29568 |
| MAP2K6 | 114495 |
| MAP3K12 | 25579 |
| MAPK3 | 50689 |

| | |
|---|---|
| MAPK15 | 286997 |
| MAR8 | 312656 |
| MCOLN3 | 308022 |
| MFN1 | 192647 |
| MINK1 | 303259 |
| MMP3 | 171045 |
| MUSK | 81725 |
| MYC | 24577 |
| MYT1L | 116668 |
| NAMPT | 297508 |
| NAPEPLD | 296757 |
| NCOR2 | 360801 |
| NCSTN | 289231 |
| NECTIN1 | 192183 |
| NEFL | 83613 |
| NEUROD1 | 29458 |
| NEUROD2 | 54276 |
| NEUROD4 | 288821 |
| NFATC1 | 100361818 |
| NFIB | 29227 |
| NGB | 85382 |
| NGF | 310738 |
| NOG | 25495 |
| NOTCH4 | 406162 |
| NPTX1 | 266777 |
| NPY1R | 29358 |
| NPY2R | 66024 |
| NR1D1 | 252917 |
| NR2F1 | 81808 |
| NRF1 | 312195 |
| NRN1 | 83834 |
| NTF3 | 81737 |
| NTRK3 | 29613 |
| NUAK1 | 299694 |
| NYAP1 | 304376 |
| OLIG2 | 304103 |
| OXTR | 25342 |
| P2RX2 | 114115 |
| PACSIN1 | 29704 |
| PAK5 | 311450 |
| PAK6 | 296078 |
| PANX1 | 315435 |
| PARD6A | 307799 |

| | |
|---|---|
| PBX1 | 304947 |
| PCDH8 | 64865 |
| PDGFB | 24628 |
| PDP1 | 54705 |
| PECAM1 | 29583 |
| PHACTR3 | 362284 |
| PIEZO1 | 361430 |
| PIK3C3 | 65052 |
| PIK3CB | 85243 |
| PIK3CD | 366508 |
| PJA2 | 192256 |
| PLAGL1 | 25157 |
| PLAUR | 50692 |
| PLCL1 | 84587 |
| PLPPR4 | 295401 |
| PLXNA2 | 289392 |
| PLXND1 | 312652 |
| PMP22 | 24660 |
| POLR3E | 361640 |
| POU4F2 | 171355 |
| POU4F3 | 364855 |
| PPP3CC | 171378 |
| PRAG1 | 306506 |
| PRKAA1 | 65248 |
| PRKAR2A | 29699 |
| PRKCQ | 85420 |
| PRKCZ | 25522 |
| PRSS12 | 85266 |
| PRUNE1 | 310664 |
| PSENEN | 292788 |
| PTGS2 | 29527 |
| PTPDC1 | 291022 |
| PTPRE | 114767 |
| PXN | 360820 |
| RAB11B | 79434 |
| RAB33A | 317580 |
| RAB3A | 25531 |
| RALGDS | 29622 |
| RAP1B | 171337 |
| RASA1 | 25676 |
| RASGRF1 | 192213 |
| REEP1 | 362384 |
| REM2 | 64626 |

| | |
|---|---|
| RET | 24716 |
| RGS2 | 84583 |
| RGS14 | 114705 |
| RHOQ | 85428 |
| RHOV | 171581 |
| RIT1 | 499652 |
| RIT2 | 291713 |
| RND1 | 362993 |
| ROCK2 | 25537 |
| RPS6KA3 | 501560 |
| RPS6KB1 | 83840 |
| RTN4RL1 | 303311 |
| RTN4RL2 | 311169 |
| RUNX1 | 50662 |
| RXRB | 361801 |
| RXRG | 83574 |
| RYK | 140585 |
| SCN1A | 81574 |
| SCN1B | 29686 |
| SDC1 | 25216 |
| SDCBP | 83841 |
| SDK2 | 360652 |
| SEMA3A | 29751 |
| SEMA3D | 246262 |
| SEMA3E | 296789 |
| SH2D3C | 362111 |
| SHTN1 | 292139 |
| SLC17A6 | 84487 |
| SLC18A2 | 25549 |
| SLC5A3 | 114507 |
| SLC6A15 | 282712 |
| SLITRK4 | 302473 |
| SMAD3 | 25631 |
| SMARCD3 | 296732 |
| SNCA | 29219 |
| SOCS3 | 89829 |
| SORBS2 | 114901 |
| SOX4 | 364712 |
| SPAST | 362700 |
| SPHK1 | 170897 |
| SPRR1A | 499660 |
| SPRY2 | 306141 |
| SS18 | 361295 |

| | |
|---|---|
| SSTR3 | 171044 |
| SUN1 | 360773 |
| SV2A | 117559 |
| SYK | 25155 |
| TAC1 | 24806 |
| TACC1 | 306562 |
| TCTN1 | 304486 |
| TGFB1 | 59086 |
| TGFBR2 | 81810 |
| TGIF1 | 316742 |
| THRB | 24831 |
| TMEM106B | 312132 |
| TNFRSF12A | 302965 |
| TP73 | 362675 |
| TPBG | 83684 |
| Tpm4 | 24852 |
| TRIM9 | 155812 |
| TRPC1 | 89821 |
| TRPC6 | 89823 |
| TWIST1 | 85489 |
| TYRO3 | 25232 |
| ULK2 | 303206 |
| UST | 361450 |
| VASP | 361517 |
| VCL | 305679 |
| VEGFA | 83785 |
| VGF | 29461 |
| VWC2 | 501231 |
| WNT6 | 316526 |
| WNT7A | 114850 |
| XRCC4 | 309995 |
| ZBTB18 | 64619 |

| **Transcription factors group** | **Entrez ID** |
|---|---|
| ABTB1 | 297432 |
| ANAPC2 | 296558 |
| ANKRD5 | 296184 |
| ANKRD6 | 500430 |
| APBB1 | 29722 |
| ARC | 54323 |
| ARHGEF25 | 314904 |
| ARID5A | 316327 |
| ASCC1 | 294512 |

| | |
|---|---|
| ASCC2 | 498402 |
| ASCC3 | 309887 |
| ATF1 | 315305 |
| ATMIN | 315037 |
| BACH1 | 304127 |
| BANP | 292064 |
| BAZ1A | 314126 |
| BBX | 303970 |
| BCL11A | 305589 |
| BCOR | 317346 |
| BHLHE22 | 365748 |
| BHLHE23 | 499952 |
| BHLHE40 | 79431 |
| BTBD6 | 690367 |
| CALCOCO1 | 246047 |
| CARF | 301446 |
| CBFA2T3 | 361431 |
| CBFB | 361391 |
| CBX8 | 303731 |
| CCRN4L | 310395 |
| CDK5 | 140908 |
| CHD1 | 308215 |
| CHD6 | 311607 |
| CHRAC1 | 315058 |
| CHURC1 | 299154 |
| CIR1 | 362149 |
| CITED2 | 114490 |
| CREBL2 | 362453 |
| CREM | 25620 |
| CRIP2 | 338401 |
| CRY1 | 299691 |
| CSRNP1 | 363165 |
| CSRP2 | 29317 |
| CTDP1 | 291414 |
| CUX2 | 288665 |
| DBP | 24309 |
| DDX20 | 84473 |
| DDX3X | 317335 |
| DEAF1 | 83632 |
| DEPDC5 | 305464 |
| DHX57 | 366532 |
| DIP2C | 307067 |
| DOT1L | 362831 |

| | |
|---|---|
| DPF1 | 50545 |
| DPF3 | 299186 |
| EBF1 | 116543 |
| EBF3 | 361668 |
| EEA1 | 314764 |
| EGR2 | 114090 |
| EGR3 | 25148 |
| EHF | 295965 |
| ELF3 | 304815 |
| ELP2 | 307545 |
| ESR1 | 24890 |
| ETV4 | 360635 |
| ETV5 | 303828 |
| FBXO41 | 312504 |
| FHL2 | 63839 |
| FOXP2 | 500037 |
| GAS7 | 85246 |
| GBX1 | 246149 |
| GLI2 | 304729 |
| GRHL2 | 299979 |
| GRM6 | 24419 |
| GTF2IRD1 | 246770 |
| GTF3C5 | 362095 |
| GZF1 | 311508 |
| HDAC11 | 297453 |
| HES6 | 316626 |
| HIC1 | 303310 |
| HMG20A | 315689 |
| HMGA1 | 117062 |
| HMGA2 | 84017 |
| HP1BP3 | 313647 |
| HPCAL1 | 50871 |
| HSF2BP | 499413 |
| ING4 | 297597 |
| IQSEC3 | 404781 |
| IRF6 | 364081 |
| IRX6 | 307715 |
| JAK2 | 24514 |
| JUNB | 24517 |
| KDM1A | 500569 |
| KDM4A | 313539 |
| KLF10 | 81813 |
| KLF11 | 313994 |

| | |
|---|---|
| KLF13 | 499171 |
| KLF14 | 312203 |
| KLF2 | 306330 |
| KLF5 | 84410 |
| KLF7 | 363243 |
| KLF9 | 117560 |
| KLHL14 | 364823 |
| KLHL25 | 293023 |
| KLHL36 | 498957 |
| KLHL5 | 305351 |
| LDB1 | 309447 |
| LEF1 | 161452 |
| LHX2 | 296706 |
| LHX4 | 360858 |
| LMO1///LMO3 | 245979 |
| LMO2 | 362176 |
| LMO4 | 362051 |
| LZTFL1 | 316102 |
| LZTS1 | 266711 |
| MAF1 | 315093 |
| MAFK | 246760 |
| MAML3 | 310405 |
| MAPK3 | 50689 |
| MCM5 | 291885 |
| MED15 | 360743 |
| MED4 | 306030 |
| MEIS2 | 311311 |
| MINK1 | 303259 |
| MKRN1 | 296988 |
| MLL5 | 311968 |
| MLLT3 | 114510 |
| MMS19 | 171124 |
| MNT | 287521 |
| MRRF | 311903 |
| MTERFD3 | 366856 |
| MXD4 | 360961 |
| MXI1 | 25701 |
| MYC | 24577 |
| MYCL1 | 298506 |
| MYT1L | 116668 |
| NAB2 | 314910 |
| NAP1L3 | 170914 |
| NCALD | 553106 |

| | |
|---|---|
| NCOR2 | 360801 |
| NEUROD1 | 29458 |
| NEUROD2 | 54276 |
| NEUROD4 | 288821 |
| NFATC1 | 100361818 |
| NFIB | 29227 |
| NFKBIE | 316241 |
| NFRKB | 315523 |
| NFX1 | 313166 |
| NHLH2 | 295327 |
| NOC3L | 361753 |
| NOP2 | 314969 |
| NOTCH4 | 406162 |
| NR1D1 | 252917 |
| NR1D2 | 259241 |
| NR1I3 | 65035 |
| NR2F1 | 81808 |
| NRF1 | 312195 |
| NRIP1 | 304157 |
| NRIP3 | 361625 |
| OLIG2 | 304103 |
| PAPOLA | 314417 |
| PAX1 | 311505 |
| PBX1 | 304947 |
| PCBP4 | 363133 |
| PCGF2 | 287662 |
| PEX14 | 64460 |
| PHF14 | 500030 |
| PHF20 | 311575 |
| PITX1 | 113983 |
| PKNOX1 | 294322 |
| PKNOX2 | 680549 |
| PLAGL1 | 25157 |
| PLXNA2 | 289392 |
| PLXND1 | 312652 |
| PNRC1 | 286988 |
| POLE4 | 362385 |
| POLR1A | 83581 |
| POLR3E | 361640 |
| POU4F2 | 171355 |
| POU4F3 | 364855 |
| POU6F1 | 116545 |
| PPRC1 | 294007 |

| | |
|---|---|
| PTBP1 | 29497 |
| PURG | 361162 |
| RAD21 | 314949 |
| RAI14 | 294804 |
| RBAK | 288489 |
| RBPMS | 498642 |
| RFXAP | 499617 |
| RGD1563666 | 317617 |
| RGS14 | 114705 |
| ROCK2 | 25537 |
| RPS6KA5 | 314384 |
| RUNX1 | 50662 |
| RUNX1T1 | 362489 |
| RUNX2 | 367218 |
| RXRB | 361801 |
| RXRG | 83574 |
| RYBP | 312603 |
| SAP30BP | 360662 |
| SATB1 | 316164 |
| SCRT2 | 366229 |
| SERTAD1 | 361526 |
| SFMBT1 | 58967 |
| SIM1 | 309888 |
| SIN3A | 363067 |
| SIRT3 | 293615 |
| SIRT5 | 306840 |
| SMAD3 | 25631 |
| SMARCA2 | 361745 |
| SMARCD3 | 296732 |
| SMYD5 | 312503 |
| SORBS2 | 114901 |
| SOX4 | 364712 |
| SP7 | 300260 |
| SRSF9 | 288701 |
| ST18 | 266680 |
| TADA2B | 289717 |
| TAF12 | 682902 |
| TBX21 | 303496 |
| TCEAL1 | 302593 |
| TEAD1 | 361630 |
| TEF | 29362 |
| TFAP4 | 360482 |
| TGFB1 | 59086 |

| | |
|---|---|
| TGIF1 | 316742 |
| THRB | 24831 |
| TOX3 | 291908 |
| TP73 | 362675 |
| TRERF1 | 316219 |
| TRIM23 | 81002 |
| TRIM9 | 155812 |
| TSC22D1 | 498545 |
| TWIST1 | 85489 |
| TXNIP | 117514 |
| USF1 | 83586 |
| VGLL4 | 297523 |
| VPS25 | 681059 |
| WDHD1 | 305827 |
| WDR45L | 360682 |
| ZBTB16 | 353227 |
| ZBTB22 | 309630 |
| ZBTB46 | 311718 |
| ZBTB5 | 298084 |
| ZBTB9 | 294289 |
| ZC3H18 | 292067 |
| ZC3H8 | 311414 |
| ZFA///ZFAT | 362925 |
| ZFAND6 | 293067 |
| ZFP62 | 361500 |
| ZFPM1 | 691504 |
| ZGPAT | 296478 |
| ZMIZ1 | 361103 |
| ZMYND8 | 296374 |
| ZNF217 | 311764 |
| ZNF238 | 64619 |
| ZNF358 | 360754 |
| ZNF395 | 305972 |
| ZNF414 | 299647 |
| ZNF467 | 500110 |
| ZNF513 | 313913 |
| ZNF575 | 308430 |
| ZNF672 | 303165 |
| ZNF697 | 295310 |
| ZNF709///ZNF14 | 266773 |
| ZNF775 | 312309 |
| ZNF786 | 100158223 |
| ZNF90///ZFP90 | 498945 |

| | |
|---|---|
| ZYX | 114636 |

| **Epilepsy Group** | **Entrez ID** |
|---|---|
| ABHD6 | 305795 |
| ADM | 25026 |
| ADRA1A | 29412 |
| ADRA1B | 24173 |
| ADRA1D | 29413 |
| ANKRD6 | 500430 |
| ARC | 54323 |
| BAG3 | 293524 |
| BAIAP2 | 117542 |
| BBC3 | 317673 |
| BDNF | 24225 |
| BEND5 | 362564 |
| CACNB4 | 58942 |
| CALCRL | 25029 |
| CAMKK1 | 60341 |
| CH25H | 309527 |
| CITED2 | 114490 |
| CNTN2 | 25356 |
| CREM | 25620 |
| CRYM | 117024 |
| CXCR4 | 60628 |
| CYR61 | 83476 |
| DBP | 24309 |
| DEPDC5 | 305464 |
| DNAJB5 | 313811 |
| DUSP5 | 171109 |
| DUSP6 | 116663 |
| EGR2 | 114090 |
| EGR3 | 25148 |
| ERRFI1 | 313729 |
| FGF13 | 84488 |
| GABRD | 29689 |
| GABRG2 | 29709 |
| GABRQ | 65187 |
| GADD45B | 299626 |
| GAL | 29141 |
| GFRA1 | 25454 |
| GRASP | 192254 |
| GRIN2A | 24409 |
| HAS1 | 282821 |

| | |
|---|---|
| HCN1 | 84390 |
| HOMER1 | 29546 |
| HSPA1A/HSPA1B | 294254 |
| HTR1B | 25075 |
| HTR1F | 60448 |
| HTR7 | 65032 |
| IFRD1 | 29596 |
| INHBA | 29200 |
| JUNB | 24517 |
| KCNA1 | 24520 |
| KCNB1 | 25736 |
| KCNK1 | 59324 |
| KCNV1 | 60326 |
| KLF10 | 81813 |
| LAMB3 | 305078 |
| LAMP5 | 362220 |
| LGI1 | 252892 |
| MCL1 | 60430 |
| METRNL | 316842 |
| NAMPT | 297508 |
| NGF | 310738 |
| NPTX2 | 288475 |
| NTF3 | 81737 |
| PCDH19 | 317183 |
| PDP1 | 54705 |
| PLK2 | 83722 |
| PNPO | 64533 |
| POLG | 85472 |
| PRICKLE1 | 315259 |
| PRRT2 | 361651 |
| PTGS2 | 29527 |
| RGS16 | 360857 |
| RGS2 | 84583 |
| SCG2 | 24765 |
| SCN1A | 81574 |
| SCN1B | 29686 |
| SCN2A | 24766 |
| SCN3B | 245956 |
| SCN4A | 25722 |
| SERTAD1 | 361526 |
| SLC2A3 | 25551 |
| SOCS3 | 89829 |
| SPNS2 | 100270678 |
| SPRY2 | 306141 |

| | |
|---|---|
| SSTR2 | 54305 |
| ST3GAL3 | 64445 |
| STRADA | 303605 |
| SV2A | 117559 |
| SV2C | 29643 |
| TAC1 | 24806 |
| TEF | 29362 |
| TRIB1 | 78969 |
| TRIM9 | 155812 |
| TRPC7 | 282822 |
| TWNK | 309441 |
| TYRO3 | 25232 |

**Neuroinflammation list of WP+B and RX2+B reversal genes**

| Symbol | Log2 Fold Change |
|---|---|
| KL | -3.603 |
| MMP3 | -3.374 |
| GRIN2A | -3.138 |
| IL6R | -2.765 |
| TGFBR2 | -2.452 |
| PTGS2 | -2.32 |
| FGFR1 | -2.314 |
| IFNG | -2.047 |
| ACVR1C | -2.02 |
| KCNJ5 | -1.993 |
| GDNF | -1.869 |
| NGF | -1.851 |
| PLA2G4A | -1.756 |
| IRS2 | -1.745 |
| BDNF | -1.537 |
| NFATC1 | -1.438 |
| JAK2 | -1.08 |
| CFLAR | -1.05 |
| TGFB1 | -0.946 |
| GABRQ | -0.916 |
| ACVR1 | -0.898 |
| PIK3CB | -0.897 |
| MAP2K4 | -0.896 |
| AKT2 | -0.726 |
| CX3CL1 | -0.675 |
| JAK1 | -0.662 |
| SYK | -0.526 |
| NCSTN | 0.357 |
| FZD1 | 0.792 |
| PPP3CC | 0.793 |
| IFNGR2 | 0.802 |
| PIK3C3 | 1.028 |
| GABRG2 | 1.162 |
| CD200 | 1.176 |
| MAPK3 | 1.215 |
| SNCA | 1.292 |
| PSENEN | 1.339 |
| GABRD | 1.796 |
| PIK3CD | 2.002 |
| IL34 | 2.079 |
| MAPK15 | 2.542 |
